# Jumping plant-lice of the tribe Paurocephalini (Hemiptera: Psylloidea: Liviidae) in Brazil

**DOI:** 10.1101/2024.12.02.626342

**Authors:** Liliya š. Serbina, Igor Malenovský, Dalva L. Queiroz, Daniel Burckhardt

## Abstract

The predominantly tropical tribe Paurocephalini of jumping plant-lice currently consists of seven genera and 94 described species worldwide, of which the genera *Klyveria* Burckhardt *et al*. and *Melanastera* Serbina *et al*. have been recorded from Brazil with two and one species, respectively. Here we review the taxonomy of the Brazilian species based on material collected from extensive fieldwork carried out in 15 states over the last decade. One species of *Klyveria* and 59 species of *Melanastera* are newly described, bringing the number of extant *Klyveria* spp. to three (both in Brazil and worldwide) and that of extant *Melanastera* spp. to 69 (60 in Brazil, 67 in the Neotropical region and one each in the Afrotropical and Oriental regions). The new species are described and illustrated, and identification keys for the Brazilian species are provided for adults and last instar immatures. The most diagnostically important structures are the distal segment of the aedeagus and the paramere, the forewing (shape, venation, surface spinules and colour pattern) and the female terminalia in the adults, and the chaetotaxy, tarsal arolium and shape of the additional pore fields on the caudal plate in the last instar immatures. The species descriptions are complemented by mitochondrial DNA barcodes (*COI* and *cytB*) and information on host plants. *Klyveria* spp. are restricted to *Luehea* (Malvaceae), while in Brazil 28 *Melanastera* spp. develop or are likely to develop on Melastomataceae, 18 spp. on Annonaceae, four spp. each on Asteraceae and Myristicaceae, and one species on Cannabaceae. Only three of the 63 species of Paurocephalini reported here from Brazil, are also known from other countries: two from Paraguay and one from Trinidad. Probably many more species of *Melanastera* are yet to be discovered and described. Priority in fieldwork should be given to areas that are at high risk of destruction or degradation by human activities, such as the Amazon rainforest, the Atlantic Forest and the Cerrado.

## Introduction

Three decades ago, at the Earth Summit in Rio de Janeiro, the Convention on Biological Diversity (CBD) was signed with the aim of sustainably conserving biodiversity, and sharing the benefits from genetic resources fairly and equitably. Habitat destruction is probably the most important cause of biodiversity loss (Dirzo & Raven 2003). Despite the CBD, habitat destruction continues at an alarming rate. The Amazon rainforest, the largest rainforest in the world, has lost more than 820,000 km^2^ (14.5%) of undisturbed humid forest, decreasing from 5,664,024 km^2^ to 4,840,000 km^2^ between 1990 and 2021 (Beuchle *et al*. 2022). In Brazil, which hosts by far the largest part of the Amazon rainforest, other biomes have also suffered drastically from human impact, though hardly noticed internationally. These biomes are the Atlantic Forest (Fig. 1A), which has been reduced by urbanisation, and the Cerrado (Fig. 1B), which has been degraded by intensive agriculture (Alencar *et al*. 2020). How habitat loss affects invertebrate diversity can only be guessed at, due to the uncertainty of the number of species on Earth today and, much less, rates of extinction (May 2011).

**FIGURE 1.**
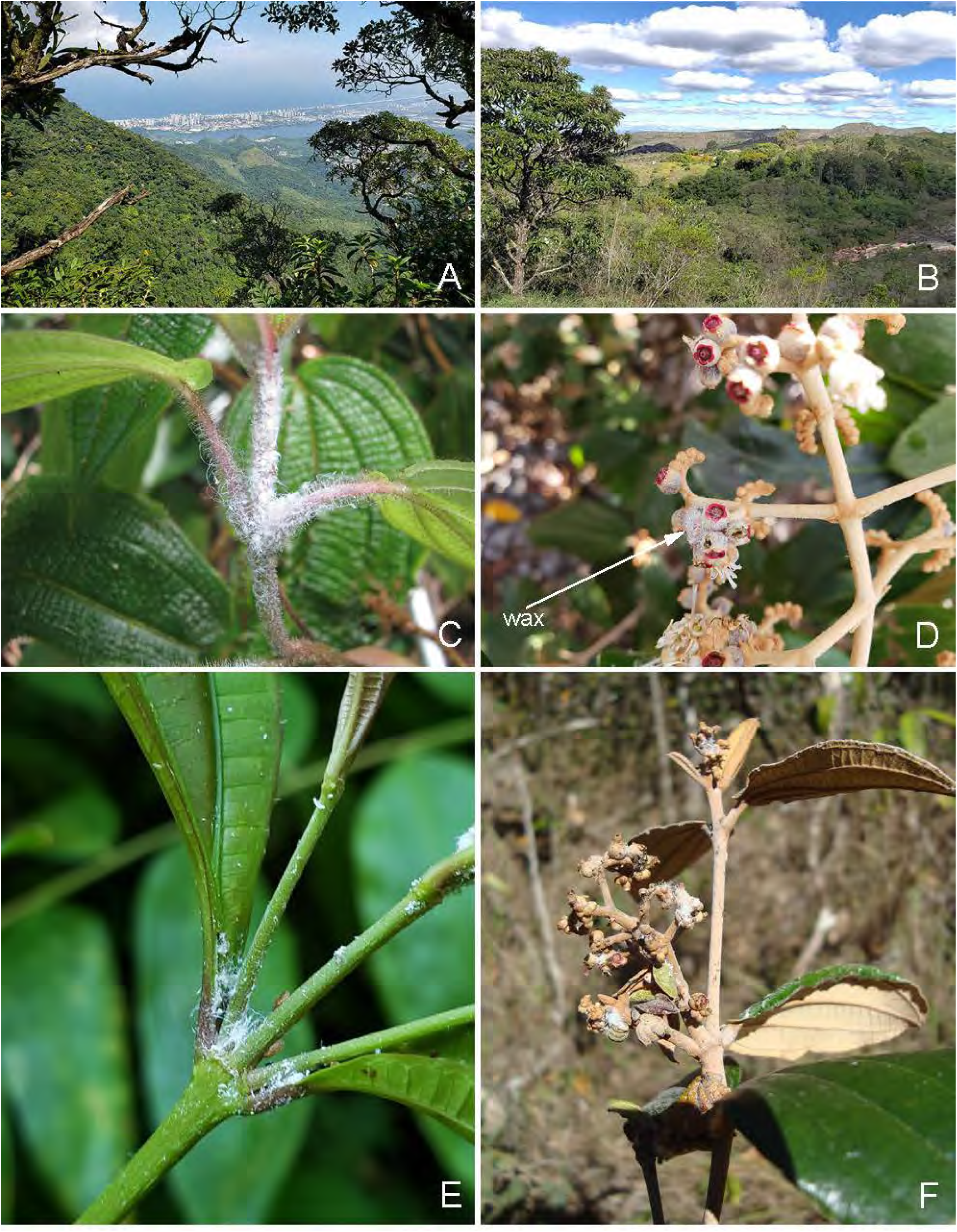
Habitats of Paurocephalini in Brazil. A, Atlantic forest in Rio de Janeiro; B, Cerrado in Minas Gerais; C, immatures of *Melanastera caramambatai* sp. nov. covered with waxy secretions on *Miconia* aff. *neourceolata*; D, immatures of *M. burchellii* sp. nov. on infrutescences of *Miconia burchellii*, visible only through their waxy secretions (arrow); E, adults and immatures of *M. atlantica* sp. nov. on *Miconia* cf. *petropolitana*; F, immatures of *M. melanocephala* sp. nov. covered with waxy secretions on infrutescences on *Miconia albicans*.

Jumping plant lice or psyllids (Insecta: Hemiptera: Psylloidea) form one of these little-known groups of arthropods. Currently, slightly over 4000 described species are known worldwide, representing at most half of the existing diversity (Burckhardt *et al*. 2021). While the Palaearctic psyllids are fairly well studied, this is not the case for the Afrotropical and Neotropical fauna (Burckhardt, unpublished data). Burckhardt & Queiroz (2012) listed 73 valid species from Brazil, but speculated that «considering that Brazil lies to a large extent in the tropics and displays a big diversity of habitats, more than 1000 psyllid species can be expected». A biodiversity project, initiated a decade ago under the leadership of DLQ, investigating the taxonomy, biogeography and host plant associations of the Brazilian Psylloidea with targeted fieldwork in all the biomes and in many states, yielded an estimated 400 undescribed morphospecies (unpublished NHMB data). Some of these have been recently described (Burckhardt *et al*. 2017, 2020; Burckhardt & Queiroz 2017, 2020, 2021, 2024a; Rendón-Mera *et al*. 2020; Serbina & Burckhardt 2017; Burckhardt 2021), bringing the number of psyllid species reported from Brazil to 165 (Burckhardt & Queiroz 2024b), but many are still awaiting taxonomic treatment, such as members of the tribe Paurocephalini (Liviidae: Liviinae). While only two named species of this tribe are known from Brazil so far (Burckhardt & Queiroz 2012; as *Diclidophlebia* Crawford), about 60 species are represented in the available material. According to Burckhardt *et al*. (2023), Paurocephalini is one of two tribes of Liviinae (Liviidae) comprising seven predominantly tropical genera, two of which occur in Brazil, namely *Klyveria* Burckhardt, Serbina & Malenovský and *Melanastera* Serbina, Malenovský, Queiroz & Burckhardt. *Klyveria* currently includes three described Neotropical species (Brazil and Paraguay on *Luehea* spp., Malvaceae, 2 spp.; Dominican amber 1 sp.). *Melanastera* comprises ten Neotropical species (Central America 6 spp., Brazil 1 sp., Mexico 1 sp.; Dominican amber 2 spp.) and one species each in the Afrotropical and Oriental regions (Burckhardt *et al*. 2023, 2024a; He *et al*. 2024). The last two species develop on *Grewia* spp., Malvaceae. One of the Neotropical species is associated with Cannabaceae and seven with Melastomataceae. Two of them are monophagous on *Miconia calvescens* (Melastomataceae), a plant native to Central and South America, and introduced as an ornamental plant to the Pacific region (French Polynesia, Hawaii, New Caledonia and Australia), where it poses a serious threat to local rainforest ecosystems (Morais *et al*. 2013; González-Muñoz *et al*. 2015). *Melanastera lucens* (Burckhardt, Hanson & Madrigal) from Costa Rica and *M. smithi* (Burckhardt, Morais & Picanço) from Brazil have both been considered for biological control of this invasive weed (Burckhardt *et al*. 2005, 2006b; Morais *et al*. 2010, 2013). *Melanastera* species on Melastomataceae species are polyvoltine with overlapping generations; immatures and adults can be found on buds, inflorescences and infrutescences of the host (Morais *et al*. 2008; Barreto *et al*. 2020) (Fig. 1C–F).

This monograph contains the descriptions and illustrations of three previously described and 60 new species of Paurocephalini. Identification keys for adults and fifth instar immatures are provided, as well as DNA barcodes and host plant information where available.

## Material and methods

**Collecting.** Most of the material was collected by D. Burckhardt and D.L. Queiroz following the protocol of Queiroz *et al*. (2017). Immature individuals were often collected directly from the plants. The material was fixed directly in 70% or 96% undenatured ethanol. The following Brazilian states (with acronym) were visited (with years of visit): Amazonas–AM (2014), Ceará–CE (2016), Goiás–GO (2012), Mato Grosso–MT (2012, 2013, 2015), Mato Grosso do Sul–MS (2012, 2013, 2021), Pará–PA (2013), Piauí–PE (2016), Minas Gerais–MG (2011, 2012, 2013, 2014, 2019, 2021), Paraná–PR (2004, 2011, 2012, 2013, 2015, 2017, 2018, 2019, 2021), Rio de Janeiro–RJ (2019), Rio Grande do Sul–RS (2016), Roraima–RR (2015), Santa Catarina–SC (2013) and São Paulo–SP (2019). When necessary, plant specimens were taken for later identification by botanists (see Acknowledgements).

**Depositories.** Psyllid specimens were examined or are cited from the following institutions: BMNH—Natural History Museum, London, UK; FSCA—Florida State Collection of Arthropods, Gainesville, Florida, USA; INPA—Instituto Nacional de Pesquisas da Amazônia, Manaus, Brazil; MHNG—Muséum d’histoire naturelle, Geneva, Switzerland; MMBC—Moravian Museum, Brno, Czech Republic; NHMB—Naturhistorisches Museum, Basel, Switzerland; UFPR—Universidade Federal do Paraná, Curitiba, Brazil. The specimens are dry-mounted on card points, mounted on slides in Canada balsam or preserved in 70% and 96% undenatured ethanol. Plant specimens are dry mounted on herbarium sheets and deposited in the Naturhistorisches Museum, Basel, Switzerland; Embrapa Florestas, Colombo, PR, Brazil; and the Museu Botânico Municipal, Curitiba, PR, Brazil.

**Morphology.** The morphological terminology is detailed in Figs 2–6. It is based on Ossiannilsson (1992), Hollis (2004), Yang *et al*. (2009), Halbert and Burckhardt (2020) and Bastin *et al*. (2023). We use the term “apical spurs” for the sclerotised spurs at the metatibial apex (Fig. 6B) and the term “bristles” for the stiff setae separating the metatibial spurs anteriorly when grouped (Burckhardt *et al*. 2023). Measurements were made using a Leica DM5500B compound microscope and Leica Application Suite software v. 4.8.0 on slide-mounted specimens as indicated in Figs 2–5. The characters and abbreviations used in the keys and descriptions are summarised in Table 1. The dimensions and ratios are given in Tables 2–5. The habitus pictures in Fig. 7 were taken with a Keyence VHX-6000 digital microscope. The photographs of the heads and wings were taken with a Leica DM5500B compound microscope, a Leica DFC320 digital camera and Leica Application Suite software v. 4.8.0. Line drawings of the morphological details were made using the same compound microscope equipped with a drawing tube. For this purpose, the specimens were cleaned in proteinase K or 10% KOH solution, dissected and temporarily mounted on a cavity slide in glycerol or permanently mounted on a normal slide in Canada balsam. The drawings were then digitised using a graphics tablet and Adobe Illustrator CS5.1 software.

**DNA barcodes.** Two mitochondrial DNA regions, cytochrome c oxidase subunit 1 (*COI*) and cytochrome b (*cytb*) gene fragments, were sequenced for most species. Both markers have been shown to be efficient in delimiting psyllid species (Martoni *et al*. 2018; Percy 2018; Cho *et al*. 2020; Pramatarova *et al*. 2024). Specimens for sequencing were preserved in ethanol or dry-mounted. Primer sequences for *COI* and *cytb*, amplicon sizes and PCR conditions used for amplification are detailed in Burckhardt *et al*. (2023). When available, both adults and immatures were sequenced for each species in order to correctly assign them. Whenever possible, specimens were selected from different samples to reflect geographical and host ranges. The uncorrected p-distances between species and populations were calculated in MEGA v. X (Kumar *et al*. 2018). DNA sequences are deposited in GenBank. Information on the voucher specimens and the accession numbers can be found in Appendix 1. Most voucher specimens from which the sequences were extracted are mounted on slides and deposited in NHMB, but some are also in BMNH, INPA, MMBC and UFPR.

**Host plants.** We use the term ʻhost plant’ to refer to those plants on which a psyllid species can complete its development (Burckhardt *et al*. 2014), i.e. plants on which immature psyllids were found. The nomenclature of plant species and genera follows the POWO (2023), the classification for the plant orders and families follows APG IV (2016).

**Conventions.** In the key for adults, one of the most important diagnostic characters is the morphology of the aedeagus, especially the lateral and dorsal views of the distal segment (Fig. 3). To properly examine this structure, material preserved in ethanol or cleared (in KOH or proteinase K) and temporarily mounted in glycerol is best. If the material permits, additional dry specimens and slides are useful. The key for the last instar immatures uses mainly microscopic details for which permanent slides are preferable. Both keys use the abbreviations listed in Table 1 and rely heavily on dimensions and ratios. In Tables 2–5, all values are given to two decimal places. One decimal place is normally given in the keys. Two decimal places (values from Tables 2–5) are given if the difference in the respective character is small. All measurements in the keys and descriptions are given in mm.

Species are listed in order of occurrence in the key for adults and are assigned to species groups for *Melanastera*. *Melanastera granulosi* sp. nov. and *M. sebiferae* sp. nov. for which no males are available are neither included in the key nor assigned to a species group; they are treated at the end. For each species, collection data for the material studied are given, including the number of specimens, geographical information, date of collection, plant on which the sample was collected (not necessarily a host plant), habitat, collector and depository, as well as collection barcode numbers and voucher names for DNA-sequenced specimens. For some species, the uncorrected p-distances are given based on the DNA barcode information in order to compare specimens collected from different localities/plants and to delimit the species from different but morphologically similar psyllid species. The localities are listed alphabetically by state and municipality.

The plant names given under “Host plants” refer to hosts confirmed by the presence of immatures. Additional notes are given under “Host plants and biology”.

**TABLE 1.** Abbreviations of the morphological characters used in the keys and species descriptions.

| **Adults** |  |
| --- | --- |
| HW | head width |
| VL | vertex length |
| VW | vertex width |
| AL | antennal length (including scape and pedicel) |
| RLTS | relative lengths of segment 10, longer terminal seta and shorter terminal seta |
| FL | forewing length |
| FW | forewing width |
| C+ScL | length of vein C+Sc |
| PtL | pterostigma length |
| PtW | pterostigma width |
| ML | length of vein M |
| M<sub>1+2</sub>L | length of vein M<sub>1+2</sub> |
| M<sub>3+4</sub>L | length of vein M<sub>3+4</sub> |
| DisM | distance between apices of veins M<sub>1+2</sub> and M<sub>3+4</sub> |
| Cu<sub>1a</sub>L | length of vein Cu<sub>1a</sub> |
| cu<sub>1</sub>W | width of cell cu<sub>1</sub> |
| MF | metafemur length |
| MT | metatibia length |
| MP | male proctiger length |
| PL | paramere length |
| DL | length of distal segment of aedeagus |
| FP | female proctiger length |
| SP | female subgenital plate length (including length of apical process) |
| CRL | circumanal ring length |
| **Fifth instar immatures** |  |
| BL | body length |
| BW | body width |
| AL | antennal length (including scape and pedicel) |
| FL | forewing pad length |
| MTT | metatibiotarsus length |
| CL | caudal plate length |
| CW | caudal plate width |
| RW | outer circumanal ring width |

**TABLE 2.**
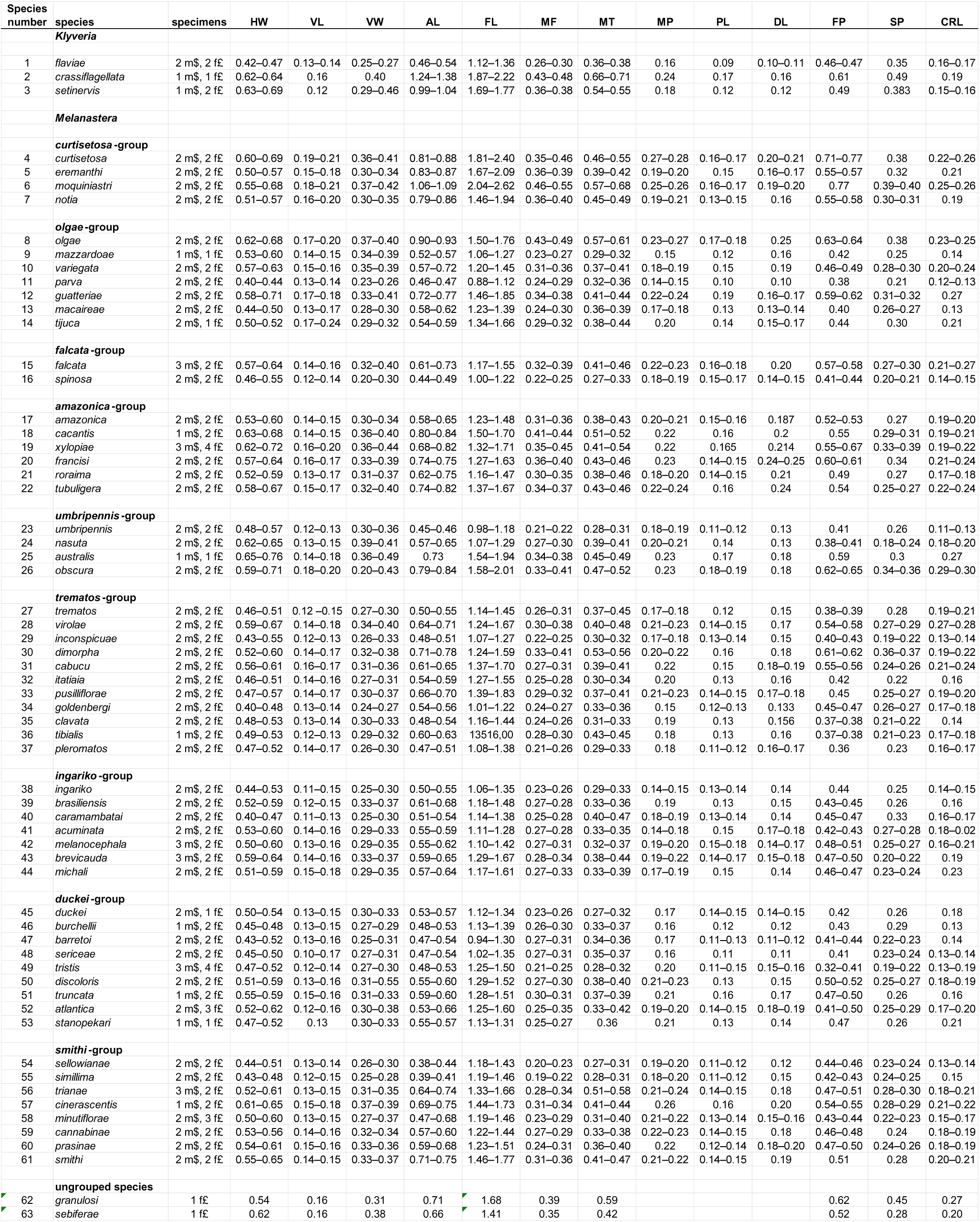
Measurements of adult characters. For abbreviations see Table 1.

**TABLE 3.**
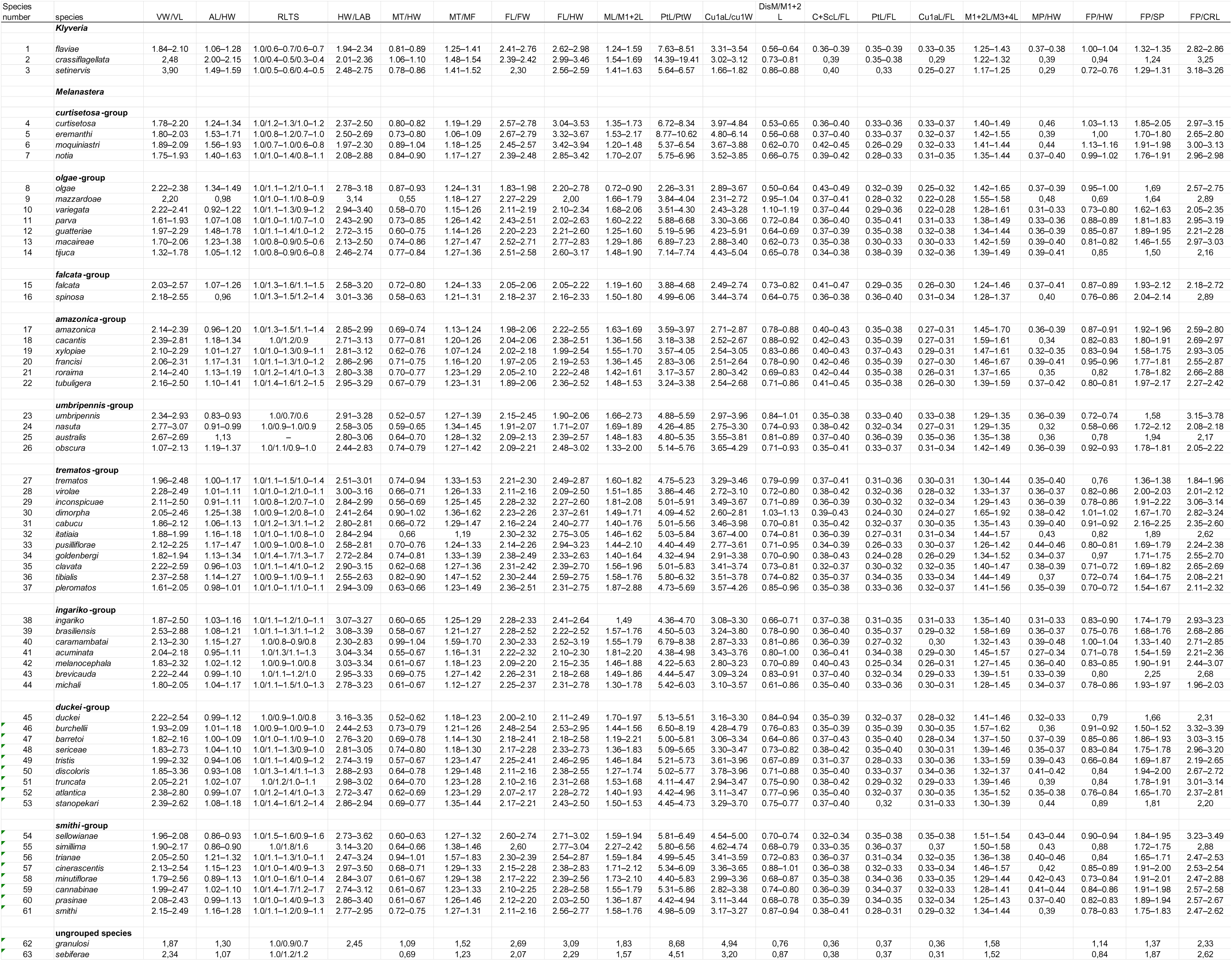
Ratios of adult characters. For abbreviations see Table 1.

**TABLE 4.** Measurements of characters of fifth instar immatures. For abbreviations see Table 1.

| Species number | species | specimens | BL | BW | AL | FL | MTT | CL | CW | RW |
| --- | --- | --- | --- | --- | --- | --- | --- | --- | --- | --- |
|  | ***Klyveria*** |  |  |  |  |  |  |  |  |  |
| 2 | *crassiflagellata* | 2 | 1.08–1.11 | 0.91–0.97 | 0.61–0.6 | 0.49–0.50 | 0.32–0.34 | 0,21 | 0.44–0.45 | 0,09 |
|  | ***Melanastera*** |  |  |  |  |  |  |  |  |  |
|  | ***curtisetos*-group** |  |  |  |  |  |  |  |  |  |
| 5 | *eremanthi* | 3 | 1.32–1.63 | 0.97–1.12 | 0.58–0.61 | 0.59–0.60 | 0.28–0.29 | 0.31–0.34 | 0.60–0.61 | 0.13–1.14 |
| 6 | *moquiniastri* | 3 | 1.69–1.86 | 1.20–1.28 | 0.73–0.77 | 0.68–0.70 | 0,41 | 0.37–0.38 | 0.71–0.73 | 0.12–0.13 |
| 7 | *notia* | 2 | 1.20–1.26 | 0.85–0.95 | 0.53–0.58 | 0.48–0.57 | 0.28–0.31 | 0.29–0.31 | 0.53–0.63 | 0,11 |
|  | ***olgae*-group** |  |  |  |  |  |  |  |  |  |
| 8 | *olgae* | 2 | 1.15–1.49 | 1.04–1.17 | 0.78–0.82 | 0,51 | 0.40–0.41 | 0.33–0.34 | 0,66 | 0,10 |
| 10 | *variegata* | 2 | 1.28–1.39 | 1,03 | 0.59–0.63 | 0.48–0.52 | 0.28–0.29 | 0.36–0.38 | 0.61–0.67 | 0.09–0.10 |
| 11 | *parva* | 1 | 0,80 | 0,61 | 0,40 | 0,35 | 0,22 | 0,20 | 0,39 | 0,10 |
| 12 | *guatteriae* | 2 | 1.25–1.33 | 0.92–0.93 | 0.61–0.66 | 0.49–0.52 | 0.28–0.31 | 0,32 | 0.61–0.62 | 0,13 |
| 13 | *macaireae* | 2 | 1.19–1.33 | 0.83–0.92 | 0.50–0.51 | 0.51–0.52 | 0.27–0.28 | 0.26–0.28 | 0.48–0.50 | 0,08 |
|  | ***falcata*-group** |  |  |  |  |  |  |  |  |  |
| 15 | *falcata* | 3 | 0.12–1.18 | 0.92–1.01 | 0.58–0.59 | 0.40–0.48 | 0.28–0.31 | 0,28 | 0.56–0.59 | 0.09–0.10 |
|  | ***amazonica*-group** |  |  |  |  |  |  |  |  |  |
| 17 | *amazonica* | 5 | 1.21–1.42 | 0.90–1.10 | 0.48–0.63 | 0.47–0.55 | 0.25–0.30 | 0.28–0.31 | 0.54–0.65 | 0.09–0.10 |
| 18 | *cacantis* | 2 | 1.18–1.47 | 0.91–1.09 | 0,74 | 0.45–0.52 | 0.32–0.35 | 0.28–0.30 | 0.59–0.61 | 0,11 |
| 19 | *xylopieae* | 2 | 1.30–1.39 | 1.02–1.12 | 0.65–0.74 | 0.51–0.58 | 0.30–0.34 | 0.29–0.33 | 0.65–0.68 | 0,10 |
| 20 | *francis* | 2 | 1.12–1.62 | 0,86 | 0.67–0.71 | 0.44–0.49 | 0.30–0.32 | 0.26–0.30 | 0.51–0.55 | 0.09–0.10 |
| 22 | *tubuligera* | 2 | 1.27–1.37 | 0.94–0.99 | 0.64–0.65 | 0.48–0.49 | 0.30–0.32 | 0,27 | 0,60 | 0,11 |
|  | ***umbripennis*-group** |  |  |  |  |  |  |  |  |  |
| 24 | *nasuta* | 2 | 1.54–1.72 | 1.17–1.23 | 0.68–0.71 | 0.51–0.54 | 0.33–0.34 | 0.33–0.37 | 0.62–0.69 | 0,13 |
| 25 | *australis* | 2 | 1,30 | 0,85 | 0.45–0.46 | 0.47–0.48 | 0.22–0.23 | 0,28 | 0,47 | 0,10 |
| 26 | *obscura* | 4 | 1.38–1.62 | 1.05–1.22 | 0.73–0.77 | 0.56–0.59 | 0.36–0.37 | 0,35 | 0.64–0.68 | 0.14–0.16 |
|  | ***trematos*-group** |  |  |  |  |  |  |  |  |  |
| 27 | *trematos* | 2 | 1.01–1.18 | 0.80–0.83 | 0.40–0.41 | 0.45–0.46 | 0.26–0.28 | 0,25 | 0.45–0.46 | 0.10–0.13 |
| 28 | *virolae* | 2 | 1.52–1.69 | 1.08–1.69 | 0.65–0.66 | 0.52–0.58 | 0,31 | 0.33–0.34 | 0.62–0.63 | 0.12–0.13 |
| 29 | *inconspicuae* | 2 | 0.92–0.98 | 0.79–0.80 | 0.46–0.47 | 0.38–0.41 | 0.21–0.23 | 0,23 | 0.42–0.45 | 0,09 |
| 30 | *dimorpha* | 2 | 1.03–1.30 | 0.83–0.90 | 0.53–0.55 | 0.45–0.48 | 0.31–0.33 | 0.27–0.29 | 0.52–0.55 | 0,11 |
| 33 | *pusilliflorae* | 2 | 1.06–1.22 | 0.82–0.93 | 0,54 | 0.39–0.46 | 0,27 | 0.26–0.28 | 0,50 | 0.13–0.14 |
| 34 | *goldenbergi* | 2 | 0.95–0.97 | 0.69–0.70 | 0.47–0.49 | 0.38–0.41 | 0,24 | 0.22–0.24 | 0.38–0.42 | 0.11–0.12 |
| 35 | *clavata* | 3 | 0.90–1.04 | 0.68–0.76 | 0.41–0.47 | 0.36–0.41 | 0.19–0.21 | 0.22–0.25 | 0.39–0.44 | 0.10–0.11 |
| 36 | *tibialis* | 1 | 0,92 | 0,71 | 0,55 | 0,43 | 0,30 | 0,24 | 0,45 | 0,11 |
| 37 | *pleromatos* | 2 | 1.00–1.25 | 0.75–0.83 | 0,40 | 0.44–0.45 | 0.20–0.21 | 0.24–0.25 | 0.46–0.47 | 0.10–0.11 |
|  | ***ingariko*-group** |  |  |  |  |  |  |  |  |  |
| 38 | *ingariko* | 2 | 0.93–1.02 | 0.77–0.84 | 0.48–0.53 | 0.40–0.42 | 0,22 | 0,23 | 0,46 | 0,10 |
| 39 | *brasiliensis* | 2 | 0.99–1.12 | 0.86–0.90 | 0.60–0.62 | 0.46–0.48 | 0.25–0.26 | 0.29–0.31 | 0.57–0.58 | 0.10–0.11 |
| 40 | *caramambatai* | 2 | 0.97–1.03 | 0.74–0.78 | 0.48–0.49 | 0.41–0.44 | 0,33 | 0.24–0.26 | 0,43 | 0.09–0.10 |
| 41 | *acuminata* | 2 | 1,29 | 0,76 | 0.44–0.49 | 0,42 | 0.21–0.28 | 0.25–0.27 | 0.42–0.45 | 0.10–0.14 |
| 42 | *melanocephala* | 2 | 1.15–1.31 | 0.96–0.97 | 0,55 | 0.46–0.50 | 0,24 | 0,29 | 0.55–0.56 | 0.10–0.11 |
| 43 | *brevicauda* | 2 | 1,18 | 0.91–0.94 | 0.59–0.64 | 0.49–0.51 | 0.32–0.33 | 0.29–0.30 | 0.54–0.56 | 0.10–0.13 |
| 44 | *Michali* | 3 | 1.10–1.24 | 0.88–0.96 | 0.56–0.57 | 0.48–0.49 | 0,26 | 0.27–0.29 | 0.53–0.56 | 0.13–0.14 |
|  | ***duckei*-group** |  |  |  |  |  |  |  |  |  |
| 46 | *burchellii* | 2 | 1.03–1.07 | 0.82–0.83 | 0.45–0.46 | 0.35–0.40 | 0,23 | 0,23 | 0.40–0.48 | 0.10–0.11 |
| 49 | *tristis* | 2 | 1.15–1.23 | 0.90–0.91 | 0.45–0.46 | 0.47–0.48 | 0.19–0.21 | 0.26–0.27 | 0.44–0.45 | 0.11–0.12 |
| 50 | *discoloris* | 1 | 0,72 | 0,81 |  | 0,37 | 0,25 | 0,19 | 0,43 | 0,14 |
| 52 | *atlantica* | 1 | 1,23 |  | 0,65 | 0,49 | 0,33 | 0,35 | 0,64 | 0,13 |
|  | ***smithi*-group** |  |  |  |  |  |  |  |  |  |
| 54 | *sellowianae* | 2 | 0,97 | 0.73–0.78 | 0.30–0.31 | 0.39–0.43 | 0,17 | 0.21–0.23 | 0.36–0.38 | 0.11–0.12 |
| 55 | *simillima* | 2 | 0.99–1.01 | 0.75–0.76 | 0.35–0.37 | 0.42–0.46 | 0.15–0.18 | 0.20–0.25 | 0.39–0.41 | 0.13–0.14 |
| 56 | *triana* | 3 | 1.18–1.29 | 0.94–1.01 | 0.59–0.65 | 0.51–0.55 | 0.32–0.36 | 0.29–0.31 | 0.54–0.57 | 0.10–0.11 |
| 57 | *cinerascens* | 3 | 1.01–1.26 | 0.96–1.08 | 0.56–0.62 | 0.43–0.50 | 0.26–0.30 | 0.28–0.35 | 0.64–0.67 | 0.11–0.12 |
| 58 | *minutiflorae* | 2 | 1,16 | 0,88 | 0,55 | 0,45 | 0.24–0.25 | 0.27–0.28 | 0.49–0.59 | 0.10–0.11 |
| 59 | *cannabinae* | 2 | 1.15–1.39 | 0.83–1.11 | 0.48–0.55 | 0.43–0.47 | 0.24–0.26 | 0.25–0.30 | 0.52–0.61 | 0,10 |
| 60 | *prasinae* | 2 | 1.08–1.27 | 0.89–0.93 | 0.53–0.58 | 0.40–0.46 | 0.24–0.28 | 0.26–0.27 | 0.52–0.56 | 0.10–0.11 |
| 61 | *smithi* | 3 | 1.00–1.38 | 0.90–1.19 | 0.57–0.63 | 0.44–0.50 | 0.29–0.31 | 0.29–0.32 | 0.58–0.64 | 0.09–0.11 |
|  | **ungrouped species** |  |  |  |  |  |  |  |  |  |
| 62 | *granulosi* | 3 | 1.10–1.40 | 0.63–0.86 | 0.57–0.61 | 0.36–0.55 | 0.32–0.39 | 0.29–0.35 | 0.48–0.51 | 0.09–0.10 |
| 63 | *sebiferae* | 1 | 1,45 | 1,10 | 0,61 | 0,52 | 0,30 | 0,31 | 0,58 | 0,07 |

**TABLE 5.** Measurements of characters of fifth instar immatures. For abbreviations see Table 1.

| Species number | species | BL/BW | AL/FL | BL/AL | MTT/BL | CW/CL | CW/RW |
| --- | --- | --- | --- | --- | --- | --- | --- |
|  | ***Klyveria*** |  |  |  |  |  |  |
| 2 | *crassiflagellata* | 1.14–1.19 | 1.22–1.32 | 1.70–1.83 | 0.29–0.32 | 2.05–2.13 | 4.77–4.85 |
|  | ***Melanastera*** |  |  |  |  |  |  |
|  | ***curtisetosa*-group** |  |  |  |  |  |  |
| 5 | *eremanthi* | 1.36–1.46 | 0.96–1.00 | 2.23–2.82 | 0.17–0.21 | 1.91–1.92 | 4.52–4.85 |
| 6 | *moquiniastri* | 1.41–1.44 | 1.04–1.15 | 2.33–2.41 | 0.22–0.24 | 1.86–1.91 | 5.71–5.73 |
| 7 | *notia* | 1.32–1.41 | 1.03–1.10 | 2.17–2.25 | 0.23–0.25 | 1.87–2.04 | 4.72–5.79 |
|  | ***olgae*-group** |  |  |  |  |  |  |
| 8 | *olgae* | 1.11–1.27 | 1.53–1.62 | 1.41–1.90 | 0.27–0.36 | 1.93–1.98 | 6.43–6.76 |
| 10 | *variegata* | 1.24–1.34 | 1.22–1.24 | 2.04–2.35 | 0.20–0.23 | 1.70–1.75 | 6.08–7.19 |
| 11 | *parva* | 1,32 | 1,14 | 2,03 | 0,27 | 1,97 | 3,93 |
| 12 | *guatteriae* | 1.36–1.44 | 1.19–1.36 | 1.89–2.17 | 0.21–0.24 | 1.88–1.95 | 4.55–4.65 |
| 13 | *macaireae* | 1.42–1.45 | 0.98–0.99 | 2.33–2.65 | 0.21–0.23 | 1.79–1.84 | 6.03–6.30 |
|  | ***falcata*-group** |  |  |  |  |  |  |
| 15 | *falcata* | 1.16–1.27 | 1.22–1.46 | 1.99–2.03 | 0.24–0.26 | 1.98–2.08 | 5.80–6.61 |
|  | ***amazonica*-group** |  |  |  |  |  |  |
| 17 | *amazonica* | 1.25–1.41 | 1.03–1.31 | 2.00–2.67 | 0.19–0.23 | 1.88–2.27 | 5.63–6.82 |
| 18 | *cacantis* | 1.29–1.35 | 1,43 | 2,00 | 0.24–0.27 | 2.02–2.06 | 5.27–5.72 |
| 19 | *xylopieae* | 1.23–1.28 | 1,28 | 1.88–2.00 | 0.23–0.25 | 2.05–2.25 | 6.64–6.86 |
| 20 | *francisi* | 1,31 | 1.46–1.53 | 1.67–2.27 | 0.19–0.27 | 1.83–1.97 | 5.49–5.81 |
| 22 | *tubuligera* | 1.35–1.39 | 1.30–1.36 | 1.95–2.16 | 0.22–0.25 | 2,21 | 5,32 |
|  | ***umbripennis*-group** |  |  |  |  |  |  |
| 24 | *nasuta* | 1.31–1.39 | 1.26–1.41 | 2.27–2.41 | 0.20–0.21 | 1,88 | 4,89 |
| 25 | *australis* | 1,53 | 0.95–0.97 | 2.86–2.88 | 0.17–0.18 | 1.65–1.69 | 4.70–4.84 |
| 26 | *obscura* | 1.25–1.37 | 1.25–1.37 | 1.79–2.20 | 0.22–0.26 | 1.84–2.06 | 4.11–4.71 |
|  | ***trematos*-group** |  |  |  |  |  |  |
| 27 | *trematos* | 1.25–1.43 | 0.86–0.92 | 2.54–2.87 | 0.24–0.26 | 1.79–1.82 | 3.55–4.55 |
| 28 | *virolae* | 1.00–1.41 | 1.14–1.25 | 2.32–2.55 | 0.19–0.21 | 1.84–1.87 | 5.02–5.31 |
| 29 | *inconspicuae* | 1.15–1.23 | 1.14–1.19 | 2.01–2.10 | 0.22–0.23 | 1.88–1.93 | 4.76–4.82 |
| 30 | *dimorpha* | 1.25–1.45 | 1.14–1.17 | 1.96–2.37 | 0.25–0.30 | 1,91 | 4.70–5.20 |
| 33 | *pusilliflorae* | 1,30 | 1.17–1.36 | 1.97–2.26 | 0.22–0.25 | 1,93 | 3,74 |
| 34 | *goldenbergi* | 1.35–1.40 | 1.14–1.30 | 1.99–2.00 | 0.25–0.26 | 1.73–1.75 | 3.42–3.61 |
| 35 | *clavata* | 1.32–1.37 | 1.16–1.19 | 2.13–2.21 | 0.21–0.22 | 1.77–1.85 | 3.68–4.34 |
| 36 | *tibialis* | 1,30 | 1,29 | 1,67 | 0,32 | 1,87 | 4,11 |
| 37 | *pleromatos* | 1.34–1.50 | 0.90–0.91 | 2.48–3.13 | 0.16–0.21 | 1.87–1.93 | 4.36–4.76 |
|  | ***ingariko*-group** |  |  |  |  |  |  |
| 38 | *ingariko* | 1.20–1.22 | 1.21–1.27 | 1,93 | 0.21–0.24 | 1,97 | 4.65–4.82 |
| 39 | *brasiliensis* | 1.14–1.24 | 1.29–1.30 | 1.63–1.81 | 0.23–0.26 | 1.85–2.00 | 5.43–5.73 |
| 40 | *caramambatai* | 1.31–1.33 | 1.09–1.18 | 2.02–2.12 | 0.32–0.34 | 1.65–1.81 | 4.32–4.72 |
| 41 | *acuminata* | 1,69 | 1,05 | 2,92 | 0,16 | 1.66–1.70 | 3.25–4.19 |
| 42 | *melanocephala* | 1.19–1.36 | 1.09–1.20 | 2.09–2.40 | 0.19–0.21 | 1.89–1.96 | 3.98–5.59 |
| 43 | *brevicauda* | 1.26–1.29 | 1.16–1.30 | 1.82–2.02 | 0.27–0.28 | 1.84–1.88 | 4.25–5.42 |
| 44 | *michali* | 1.22–1.30 | 1.15–1.17 | 1.98–2.19 | 0.21–0.24 | 1.85–2.00 | 3.93–4.23 |
|  | ***duckei*-group** |  |  |  |  |  |  |
| 46 | *burchellii* | 1.25–1.31 | 1.13–1.30 | 2.25–2.38 | 0,22 | 1.78–2.06 | 3.59–4.72 |
| 49 | *tristis* | 1.28–1.34 | 0.95–0.97 | 2.53–2.66 | 0,17 | 1.62–1.70 | 3.66–3.98 |
| 50 | *discoloris* | 0,89 |  |  | 0,35 | 2,31 | 3,24 |
| 52 | *atlantica* | 1,28 | 1,31 | 1,89 | 0,27 | 1,84 | 4,94 |
|  | ***smithi*-group** |  |  |  |  |  |  |
| 54 | *sellowianae* | 1,33 | 0.73–0.78 | 3,20 | 0,17 | 1.60–1.80 | 3,14 |
| 55 | *simillima* | 1.29–1.35 | 0.81–0.83 | 2.65–2.91 | 0.15–0.18 | 1.63–1.90 | 2.87–2.95 |
| 56 | *trianae* | 1.19–1.27 | 1.14–1.23 | 1.90–2.02 | 0.27–0.30 | 1.87–1.88 | 5.42–5.68 |
| 57 | *cinerascentis* | 1.12–1.20 | 1.24–1.31 | 1.69–2.07 | 0.23–0.27 | 1.94–2.28 | 5.49–6.39 |
| 58 | *minutiflorae* | 1,32 | 1,21 | 2,12 | 0,21 | 1.86–2.10 | 4.94–5.61 |
| 59 | *cannabinae* | 1.12–1.38 | 1.11–1.16 | 2.27–2.39 | 0,21 | 2.06–2.09 | 5.30–5.88 |
| 60 | *prasinae* | 1.21–1.37 | 1.28–1.34 | 2.02–2.19 | 0.22–0.23 | 1.98–2.09 | 5.15–5.44 |
| 61 | *smithi* | 1.11–1.21 | 1.26–1.30 | 1.74–2.20 | 0.22–0.29 | 1.98–2.04 | 5.57–6.56 |
|  | **ungrouped species** |  |  |  |  |  |  |
| 62 | *granulosi* | 1.49–1.76 | 1.11–1.57 | 1.93–2.32 | 0.28–0.32 | 1.45–1.64 | 4.97–5.61 |
| 63 | *sebiferae* | 1,31 | 1,16 | 2,39 | 0,21 | 1,86 | 7,88 |

**FIGURE 2.**
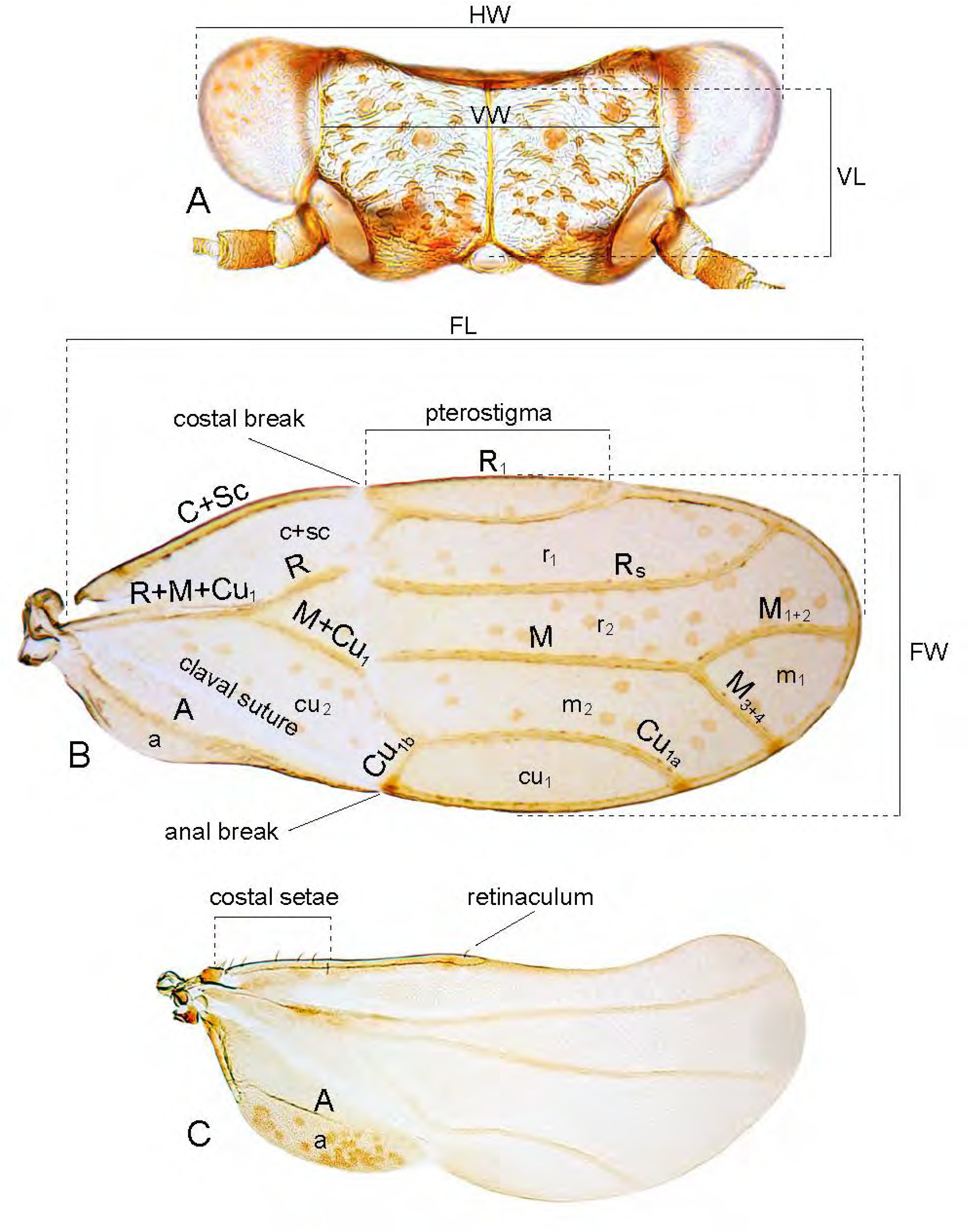
Morphology and measurements of Paurocephalini. A, *Melanastera michali* sp. nov.; B, *M. clavata* sp. nov.; C, *M. tubuligera* sp. nov. A, head, dorsal view; B, forewing; C, hindwing. For abbreviations see Table 1.

**FIGURE 3.**
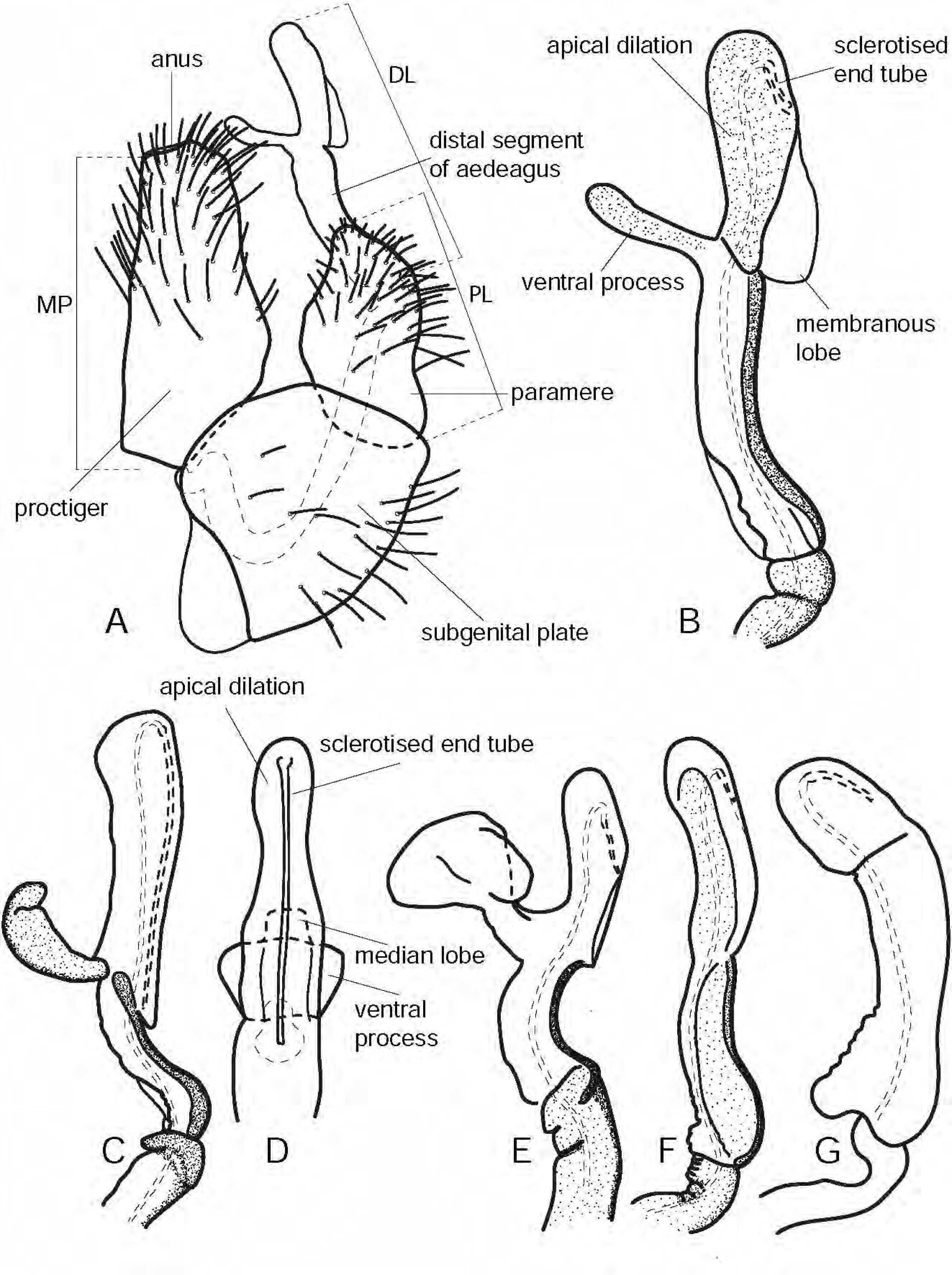
Morphology and measurements of male terminalia of *Melanastera* and *Klyveria* spp. A, B, *M. inconspicuae* sp. nov.; C, D, *M. roraima* sp. nov.; E, *M. sellowianae* sp. nov.; F, *M. curtisetosa* sp. nov.; G, *K. flaviae* sp. nov. A, m$ terminalia, lateral view; B, C, E–G, distal segment of aedeagus, lateral view; D, distal segment of aedeagus, dorsal view. For abbreviations see Table 1.

**FIGURE 4.**
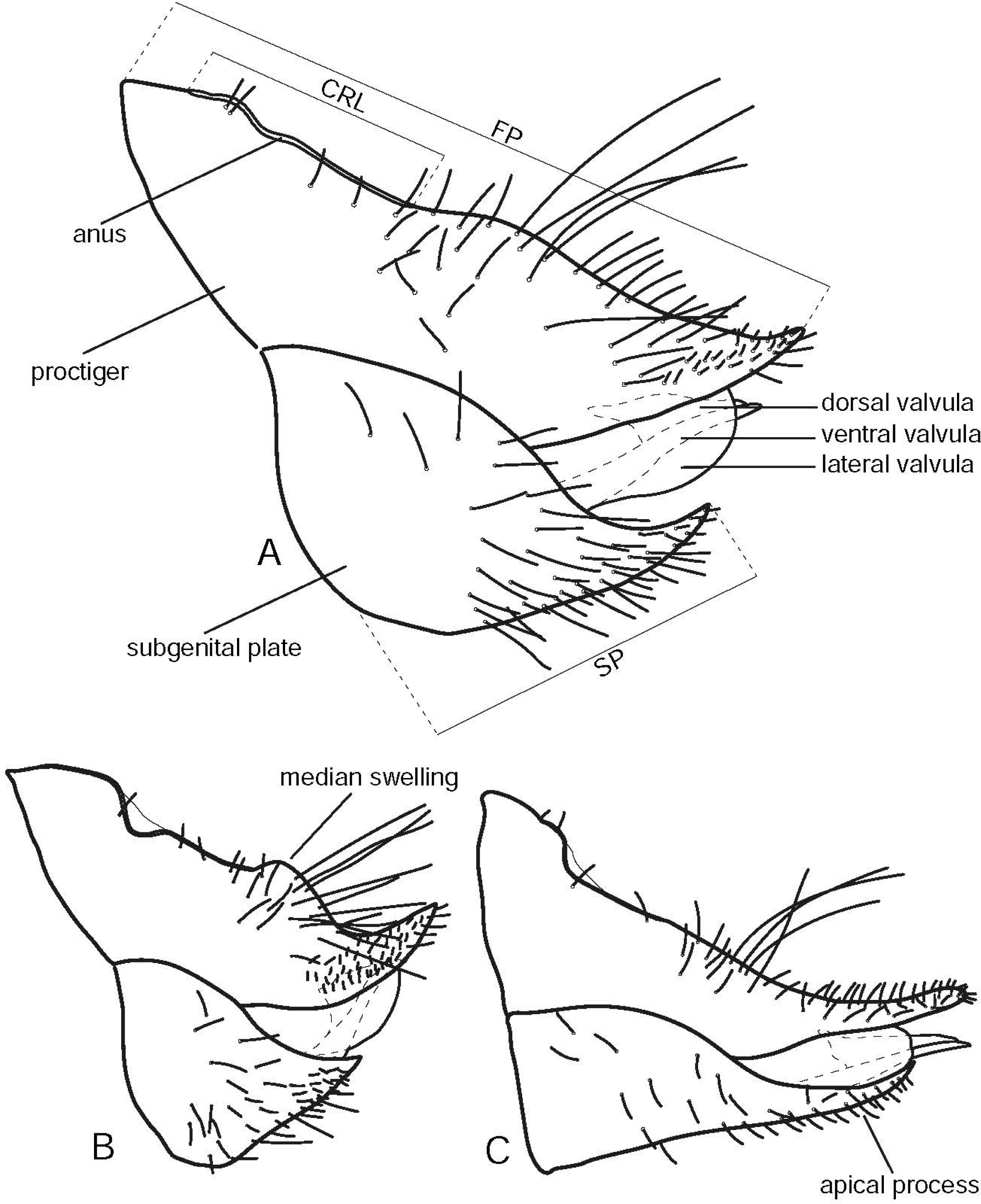
Morphology and measurements of female terminalia of *Melanastera* and *Klyveria* spp. A, *M. discoloris* sp. nov.; B, *M. michali* sp. nov.; C, *K. flaviae* sp. nov. For abbreviations see Table 1.

**FIGURE 5.**
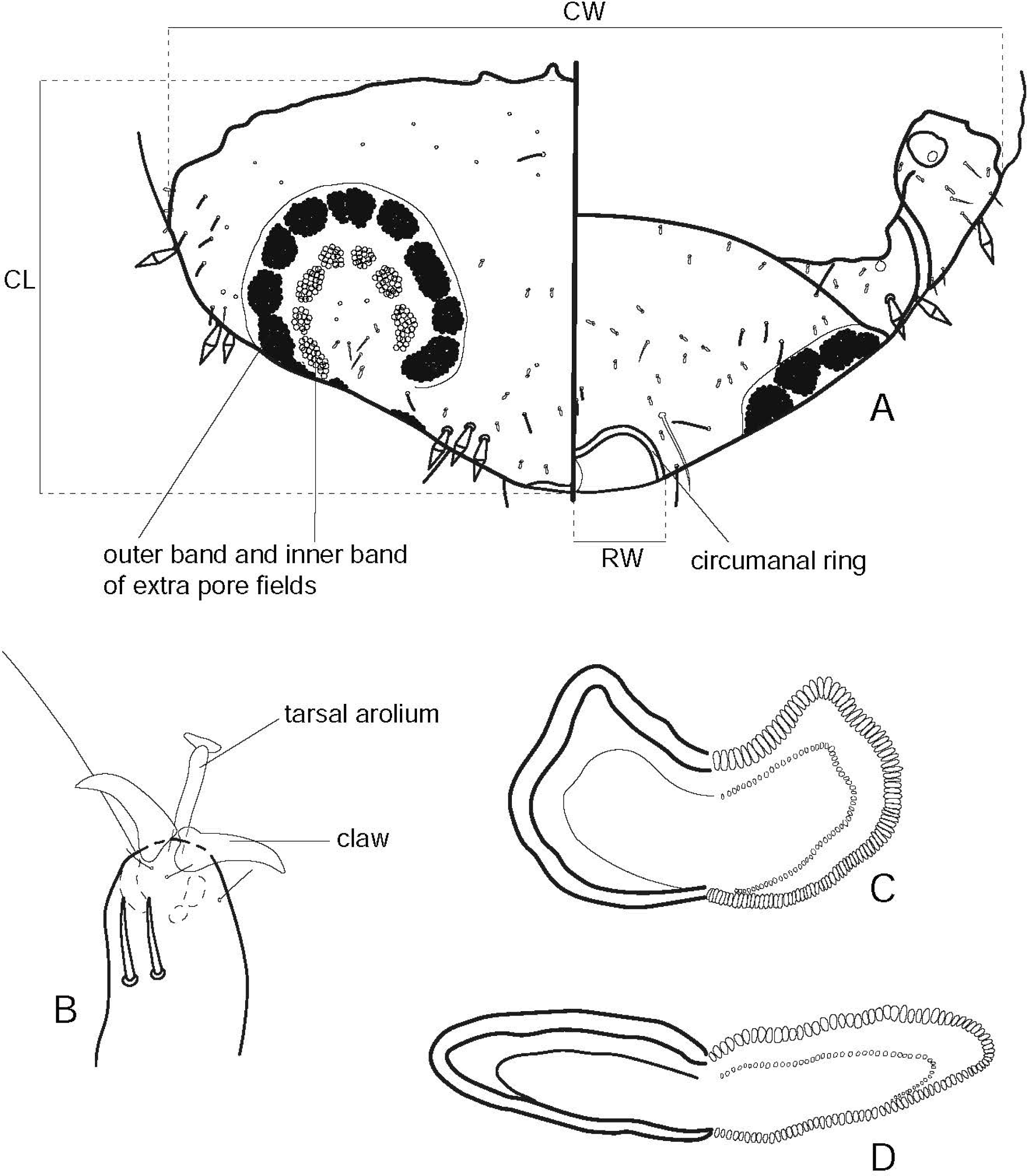
Morphology and measurements of immatures of *Melanastera* spp. A, B, *M. eremanthi* sp. nov.; C, *M. moquiniastri* sp. nov.; D, *M. trematos* sp. nov. A, caudal plate, dorsal (left) and ventral (right) views; B, tarsal arolium with claws, ventral view; C, D, circumanal ring, ventral view. For abbreviations see Table 1. The width of circumanal ring (RW) is indicated only for the ventral side.

**FIGURE 6.**
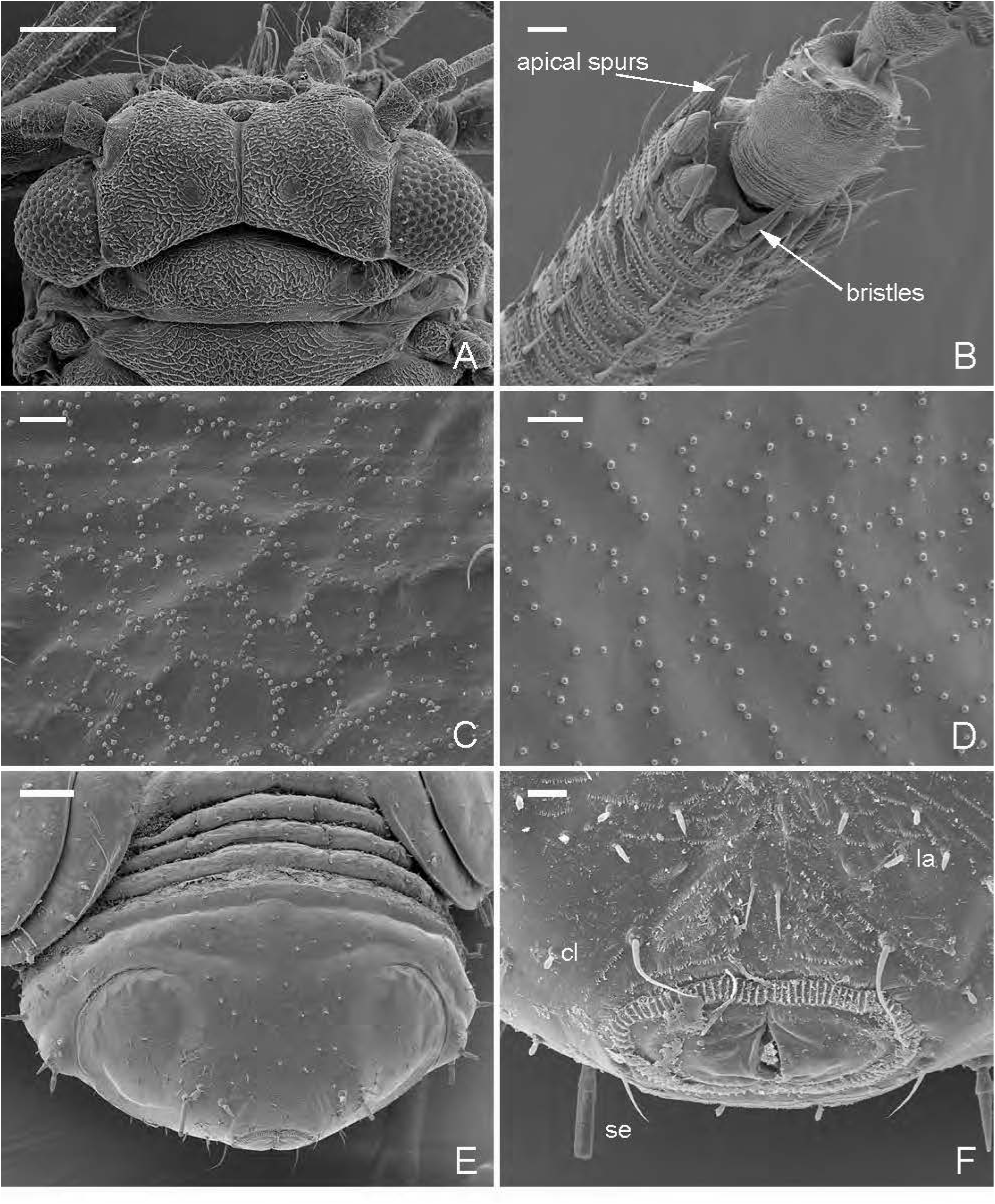
Morphology of *Melanastera* spp. A, head of *M. tubuligera* sp. nov.; B, apex of metatibia of *M. melanocephala* sp. nov.; C, surface spinules of *M. falcata* sp. nov.; D, surface spinules of *M. melanocephala* sp. nov.; E, caudal plate of *M. falcata* sp. nov. with extra pore fields and circumanal ring; F, abdominal apex of *M. falcata* sp. nov. with minute clavate setae (cl), lanceolate setae (la) and sectasetae (se). Scale bars: A = 0.5 mm; B–D, F = 0.01 mm; E = 0.05 mm.

## Taxonomy

## Keys to the Brazilian genera and species of Paurocephalini

Adults

(*Melanastera granulosi* and *M. sebiferae* excluded for the lack of males)

1 Apical metatibial spurs not or only weakly sclerotised, forming a complete, posteriorly open crown. Frons small, rhomboidal. Abdominal tergites lacking median tubercle. Paramere, in lateral view, with short, digitiform apex weakly directed posteriad (Fig. 21B, I, P). Base of distal segment of aedeagus strongly inflated ventrad; apical dilation short, oval (Fig. 21D, K, R). Female terminalia slender and relatively long; apical process on female subgenital plate long and narrow (Fig. 21G, N, U). On *Luehea* spp. (Malvaceae). ***Klyveria*** 2 Apical metatibial spurs strongly sclerotised, often separated by one or several bristles anteriorly (Fig. 6B). Frons large, triangular. At least two abdominal tergites with a median tubercle. Paramere, in lateral view, different or, if apex digitiform, then directed anteriad (Figs 22–35). Base of distal segment of aedeagus not inflated ventrally; apical dilation relatively long (Figs 22–35). Female terminalia robust and relatively short; apical process on female subgenital plate short and broad (Figs 22–35). Not on *Luehea* spp ***Melanastera*** 4 FL < 1.5. Vertex half as long as wide, with distinct imbricate microsculpture (Fig. 8A). Antenna light except for apex, short (AL < 0.8; AL/HW < 1.4). Forewing evenly pale yellow, with subparallel margins, narrowly rounded apically; FL/FW ≥ 2.4 (Fig. 14A). Terminalia as in Figs 21A–G ***K. flaviae*** sp. nov.

2 FL > 1.5. Vertex distinctly shorter than half as wide, with indistinct microsculpture (Fig. 8B, C). Antenna dark brown, long (AL > 0.8; AL/HW > 1.4; Fig. 7A). Forewing with dark brown veins; widest in apical third, broadly rounded apically; FL/FW < 2.4 (Fig. 14B, C). 3 Setae on head (Fig. 8B), thorax and forewing veins short. Forewing with narrow pterostigma (PtL/PtW > 10.0); vein Rs almost straight, slightly curved to fore margin apically; vein Cu_1a_ weakly convex, Cu_1a_L/cu_1_W > 2.5 (Fig. 14B). Paramere, in both dorsal and lateral views, with a broad apex bearing two small sclerotised teeth (Fig. 21I, M). Female proctiger, in dorsal view, with oval circumanal ring. ***K. crassiflagellata*** (Burckhardt) Setae on head (Fig. 8C), thorax and forewing veins long. Forewing with broad pterostigma (PtL/PtW < 10.0); vein Rs sinuate, strongly curved to fore margin apically; vein Cu_1a_ strongly convex, Cu_1a_L/cu_1_W < 2.5 (Fig. 14C). Paramere, in both dorsal and lateral views, with a narrow apex lacking distinct sclerotised teeth (Fig. 21P, T). Female proctiger, in dorsal view, with vaguely cruciform circumanal ring ***K. setinervis*** (Burckhardt) Vertex and thorax covered with long macroscopical setae (Fig. 8D–G). Pterostigma with subparallel margins, weakly or not widening towards apex (Fig. 14D–G). On Asteraceae. … ***curtisetosa*-group** 5

- Vertex and thorax covered with microscopical setae (Fig. 6A). Pterostigma distinctly expanding towards the middle or apical third (Figs 15–20). On other plant families. 8

3 Brown dots on forewing faint or completely lacking (Fig. 14D, E); FL/FW > 2.6. Apical dilation of distal segment of aedeagus narrow in both lateral and dorsal views (Figs 21X, Y, 22C, D). Setae in apical half of female subgenital plate relatively short (Figs 21AA, 22F). 6

- Brown dots on forewing distinct (Fig. 14F, G); FL/FW < 2.6. Apical dilation of distal segment of aedeagus broad in lateral and dorsal views (Fig. 22I, J, O, P). Setae in apical half of female subgenital plate moderately long (Fig. 22L, R). 7 4 Forewing with few, very faint brown dots (Fig. 14D); surface spinules relatively dense. Male proctiger slender (Fig. 21V). Paramere, in lateral view, lamellar (Fig. 21W). Apical dilation of distal segment of aedeagus narrow and, in lateral view, slightly curved (Fig. 21X, Y). Dorsal margin of female proctiger, in lateral view, with small swelling distal to circumanal ring (Fig. 21AA). Probably on Asteraceae ***M. curtisetosa*** sp. nov.

- Forewing membrane lacking brown dots (Fig. 14E); surface spinules relatively sparse. Male proctiger robust (Fig. 22A). Paramere, in lateral view, lanceolate (Fig. 22B). Apical dilation of distal segment of aedeagus wide and, in lateral view, almost straight (Fig. 22C, D). Dorsal margin of female proctiger, in lateral view, with large swelling distal to circumanal ring (Fig. 22F). On *Eremanthus erythropappus **M. eremanthi*** sp. nov.

5 FL ≥ 2.0. Male proctiger, in lateral view, weakly curved posteriorly (Fig. 22G). Paramere, in lateral view, relatively robust (Fig. 22H). Female terminalia relatively long, FP/HW > 1.1; proctiger straight apically; subgenital plate, in lateral view, gradually narrowing towards apex (Fig. 22L). Head usually light (Fig. 8F). On *Moquiniastrum polymorphum **M***.

*moquiniastri* sp. nov.

- FL < 2.0. Male proctiger, in lateral view, distinctly produced posteriorly (Fig. 22M). Paramere, in lateral view, relatively slender (Fig. 22N). Female terminalia relatively short, FP/HW < 1.1; proctiger upturned apically; subgenital plate, in lateral view, in apical third abruptly narrowing towards apex (Fig. 22R). Head usually dark (Fig. 8G). On *Moquiniastrum polymorphum* and *Vernonanthura phosphorica. **M. notia*** sp. nov.

6 Forewing yellow to dark brown, lacking pattern of small brown dots (Figs 14H–J, 15A–F).

… *olgae*-group 9

- Forewing with pattern of small brown dots (Figs 15G–J, 16–19, 20A–F), sometimes confluent (e.g. Figs 16F, 17D). 15

7 Forewing coriaceous, entirely dark brown, FL/FW < 2.0 (Figs 7C, 14H); costal break absent; cell m_1_ long (ML/M_1+2_L < 1.0); surface spinules indistinct. Apical dilation of distal segment of aedeagus long and narrow; ventral process, in lateral view, claw-like apically (Fig. 22U), in dorsal view, broad basally and narrowly truncate apically (Fig. 22V). On *Guatteria punctata **M. olgae*** sp. nov.

- Forewing membranous, pale yellow to brown or partly dark brown with clear patches, FL/FW > 2.0 (Figs 14I, J, 15A–F); cell m_1_ short (ML/M_1+2_L > 1.0); costal break developed; surface spinules distinct. Apical dilation of distal segment of aedeagus short and broad; ventral process different (Figs 23C, G, H, M, N, S, T, 24C–E, J, K). …10

8 Forewing pattern sexually dimorphic: male with three large clear areas at base, along nodal line and at anterior margin subapically (Figs 14I, 15A); female uniformly pale or dark brown with small clear patches at wing base and at apex of cu_2_ cell (Figs 14J, 15B). Vertex smooth, lacking distinct microsculpture (Fig. 8I, J). Paramere, in lateral view, slender, weakly curved anteriad (Fig. 23B, F). … 11

- Forewing in both sexes entirely yellow or pale brown (Fig. 15C–F). Vertex with distinct microsculpture (Fig. 9A–D). Paramere, in lateral view, robust, relatively straight (Figs 23L, R, 24B, I). … 12

11 Coloration light: body orange, forewing pale brown (Fig. 14I, J). Forewing of male with narrow light band along nodal line and small light patch at anterior margin subapically (Fig. 14I). Teminalia as in Fig. 23 A–D. Matto Grosso. ***M. mazzardoae*** sp. nov.

- Coloration dark: body and forewing dark brown (Fig. 15A, B). Forewing of male with broad clear band along nodal line and large clear patch at anterior wing margin subapically (Fig. 14A). Teminalia as in Fig. 23E–J. Amazonas and Roraima. On *Guatteria* sp. ***M. variegata***

sp. nov.

- FL < 1.2. Ventral process of distal segment of aedeagus, in lateral view, about as long as apical dilation (Fig. 23M). Female terminalia relatively elongate (FP/HW > 0.87). Perhaps on *Xylopia* sp. ***M. parva*** sp. nov.

- FL > 1.2. Ventral process of distal segment of aedeagus, in lateral view, shorter than apical dilation (Figs 23S, 24C, J). Female terminalia relatively short (FP/HW ≤ 0.87) 13

13 Forewing broad (FL/FW < 2.4), membrane yellow to pale brown, lacking clear bands along veins (Fig. 15D). Metatibia with ca. 5 bristles separating apical spurs in two groups. Paramere, in lateral view, strongly narrowing apically (Fig. 23R). On *Guatteria sellowiana.* …*M. guatteriae* sp. nov.

- Forewing narrow (FL/FW > 2.4), membrane bright yellow, with clear bands along veins (Fig. 15E, F). Metatibia lacking bristles between apical spurs. Paramere, in lateral view, weakly narrowing apically (Fig. 24B, I). 14

14 Median suture of vertex fully developed (Fig. 9C). Forewing with short pterostigma (PtL/FL < 0.35) and broad cell cu_1_ (Cu_1a_L/cu_1_W < 4.0) (Fig. 15E). Paramere, in lateral view, relatively slender with weakly curved fore margin (Fig. 24B). Apical dilation of distal segment of aedeagus, in both lateral and dorsal views, cylindrical with subparallel margins (Fig. 24C–E); ventral process, in dorsal view, three-lobed. Circumanal ring of female proctiger small (FP/CRL > 2.5) (Fig. 24G). On *Macairea radula. **M. macaireae*** sp. nov. -Median suture of vertex mostly reduced, developed only at apex (Fig. 9D). Forewing with long pterostigma (PtL/FL > 0.35) and narrow cell cu_1_ (Cu_1a_L/cu_1_W > 4.0) (Fig. 15F). Paramere, in lateral view, relatively massive with strongly curved fore margin (Fig. 24I). Apical dilation of distal segment of aedeagus, in lateral view, falcate (Fig. 24J); both ventral process and apical dilation, in dorsal view, rhomboid with curved margins (Fig. 24K). Circumanal ring of female proctiger large (FP/CRL < 2.5) (Fig. 24M). ***M. tijuca*** sp. nov. - Distal segment of aedeagus lacking ventral process (Fig. 24P, W). … ***falcata*-group** 16 -Distal segment of aedeagus bearing ventral process (Figs 25–35, 36C), which can be small and button-like (Figs 28J, O, 30I: *M. dimorpha*, *M. cabucu*, *M. ingariko*) or fused with apical dilation (Fig. 26U: *M. nasuta*). 17

16 Paramere, in lateral view, lanceolate, lacking stout setae on inner face (Fig. 24O); apex, in dorsal view, subacute (Fig. 24R). Apical portion of distal aedeagal segment, in lateral view, slender, falcate (Fig. 24P). Dorsal margin of female proctiger, in lateral view, weakly concave distal to circumanal ring; apex not upturned (Fig. 24S). On *Virola sebifera **M. falcata*** sp.nov.

- Paramere, in lateral view, digitiform, bearing a group of six stout setae on inner face near posterior margin in basal half (Fig. 24U, V); apex, in dorsal view, truncate (Fig. 24Y). Apical portion of distal aedeagal segment, in lateral view, straight, cylindrical (Fig. 24W). Dorsal margin of female proctiger, in lateral view, sinuate distal to circumanal ring, apex upturned (Fig. 24Z). ***M. spinosa*** sp. nov.

17 Forewing with pterostigma shorter than 4.0 times as wide; vein R_1_ strongly convex medially relative to costal wing margin (Figs 15J, 16A–E). Ventral process of distal aedeagal segment, in lateral view, claw-like and directed dorsad (Figs 25C, J, P, V, 26C, I). On Annonaceae. … ***amazonica*-group** 18

- Forewing with pterostigma longer than 4.0 times as wide; vein R_1_ weakly convex or straight medially (Figs 16F–J, 17–20). Ventral process of distal aedeagal segment not claw-shaped (Figs 26O, U, 27C, I, 35C, I, O, U, 36C). On Annonaceae and other plant families. 23

18 Surface spinules in cell r_2_ of forewing above bifurcation of M forming distinct hexagons of a double row of spinules; spinules always absent from within the cells. Distal segment of aedeagus with ventral process situated distal of the middle of the segment (Fig. 25C, D). On *Annona foetida* and *Guatteria megalophylla **M. amazonica*** sp. nov.

- Surface spinules in cell r_2_ of forewing above bifurcation of M forming at most indistinct hexagons; always with a few spinules in the cells. Position of ventral process of distal segment of aedeagus variable. …19

19 Ventral process of distal segment of aedeagus situated distal of the middle or in the middle of the segment (Fig. 25J, P). …20

- Ventral process of distal segment of aedeagus situated proximal of the middle of the segment (Figs 25V, 26C, I). 21

20 Brown dots on forewing relatively dense, usually three and more confluent (Fig. 16A); body relatively dark. Paramere, in lateral view, robust (Fig. 25H). Sclerotised end tube of aedeagus short (Fig. 25J, L). On *Annona cacans **M. cacantis*** sp. nov.

- Brown dots on forewing relatively sparse, usually only two confluent (Fig. 16B); body relatively light. Paramere, in lateral view, elongate (Fig. 25O). Sclerotised end tube of aedeagus very long (Fig. 25P, Q). On *Xylopia aromatica*, *X. nitida* and *Guatteria* sp. … ***M. xylopiae*** sp. nov.

21 Paramere, in lateral view, hardly indented antero-subapically (Fig. 25U). FP/HW > 0.9. On

*Guatteria* sp. and *Xylopia aromatica **M. francisi*** sp. nov.

- Paramere, in lateral view, distinctly indented antero-subapically (Fig. 26B, H). FP/HW <

0.9. … 22

22 Basal part of distal segment of aedeagus strongly curved dorsally; apical dilation, in lateral view, slightly concave ventrally and relatively broad apically (Fig. 26C). Dorsal margin of female proctiger, in lateral view, weakly concave in apical half (Fig. 26F); FP/SP < 1.9.

Probably on *Guatteria* sp ***M. roraima*** sp. nov.

- Basal part of distal segment of aedeagus weakly curved dorsally; apical dilation, in lateral view, almost straight ventrally and with subparallel margins apically (Fig. 26I). Dorsal margin of female proctiger, in lateral view, strongly concave in apical half (Fig. 26L); FP/SP > 1.9. *Guatteria schomburgkiana*, perhaps also *Xylopia aromatica **M. tubuligera*** sp. nov. 23 Forewing proximal to nodal line conspicuously lighter than distally (Fig. 16F–J). …

***umbripennis*-group** 24

- Forewing relatively uniformly coloured (Figs 17–20). 27

24 Forewing pattern strongly sexually dimorphic (Fig. 16F, G). Vertex almost smooth, lacking distinct microsculpture (Fig. 10C). Paramere, in lateral view, wide (Fig. 26N). Ventral process of distal segment of aedeagus large and broad, well separated from apical dilation (Fig. 26O, P). FP/SP < 1.6. ***M. umbripennis*** sp. nov.

- Forewing pattern not sexually dimorphic (Fig. 16H–J). Vertex with distinct microsculpture (Fig. 10D–F). Paramere, in lateral view, slender (Figs 26T, 27B, H). Ventral process of distal segment of aedeagus small and narrow, well separated from apical dilation (Fig. 27C, D, I, J) or large, broad and fused with apical dilation (Fig. 26U, V). FP/SP > 1.6. On Annonaceae. … 25

25 AL/HW < 1.0; FL/HW < 2.2. Paramere, in lateral view, strongly narrowed in apical quarter, forming digitiform process which is directed anteriad (Fig. 26T, W). Ventral process of distal segment of aedeagus large, fused with apical dilation (Fig. 26U); in dorsal view, large, triangular with small button-like lobes directed apicad (Fig. 26V). FP/HW < 0.7. On *Annona* sp. and perhaps *Guatteria* sp ***M. nasuta*** sp. nov.

- AL/HW > 1.0; FL/HW > 2.2. Paramere, in lateral view, gradually narrowing in apical quarter towards apex, which is directed upwards (Fig. 27B, H). Ventral process of distal segment of aedeagus short, well separated from apical dilation (Fig. 27C, I). FP/HW > 0.7. … 26

26 Paramere relatively short (MP/PL > 1.3) and narrow (Fig. 27B). Ventral process of distal segment of aedeagus, in lateral view, distinctly shorter than apical dilation (Fig. 27C). FP/HW < 0.9. On *Guatteria australis*, perhaps also on *Annona dolabripetala **M. australis*** sp. nov.

- Paramere relatively long (MP/PL < 1.3) and broad (Fig. 27H). Ventral process of distal segment of aedeagus, in lateral view, only slightly shorter than apical dilation (Fig. 27I). FP/HW > 0.9. On *Annona cacans **M. obscura*** sp. nov.

27 Ventral process of distal segment of aedeagus, in dorsal view, narrower than apical dilation (e.g. Figs 27P, V, 30D). … ***trematos*-group. …**28

- Ventral process of distal segment of aedeagus, in dorsal view, at least slightly wider than apical dilation (e.g. Fig. 30P, V). …38

28 Surface spinules in cell r_2_ of forewing above bifurcation of vein M forming mostly single-row hexagons (at least in males) (Fig. 6D). …29

- Surface spinules in cell r_2_ of forewing above bifurcation of vein M forming mostly double-row hexagons (Fig. 6C). …31

29 Brown dots on forewing indistinct, absent from cell c+sc (Fig. 17A). Ventral process of distal aedeagal segment, in lateral view, shorter than half length of apical portion (Fig. 27O). Female proctiger, in lateral view, with straight or weakly concave dorsal margin (Fig. 27R); FP/SP < 1.5. On *Trema micranthum **M. trematos*** sp. nov.

- Brown dots on forewing conspicuous, present also in cell c+sc (Figs 17B, C). Ventral process of distal aedeagal segment, in lateral view, longer than half length of apical portion (Figs 27U, 28C). Female proctiger, in lateral view, with flat swelling distal to circumanal ring (Figs 27X, 28F); FP/SP > 1.5. …30

30 AL > 0.6. FL/FW < 2.2. Paramere apex, in dorsal view, truncate, bearing two small teeth (Fig. 27W). Ventral process of distal segment of aedeagus relatively broad (Fig. 27V).

FP/CRL < 2.5. On *Virola bicuhyba **M. virolae*** sp. nov.

- AL < 0.6. FL/FW > 2.2. Paramere apex, in dorsal view, pointed, bearing a single small tooth (Figs 28E). Ventral process of distal segment of aedeagus relatively narrow (Fig. 28D). FP/CRL > 2.5. On *Miconia inconspicua **M. inconspicuae*** sp. nov.

31 Ventral process of distal segment of aedeagus small, button-like (Fig. 28J, O). …32

- Ventral process of distal segment of aedeagus large (e.g. Figs 28U, 29C, I, O, U). …33

32 Coloration and microsculpture on vertex sexually dimorphic: male with head mostly dark, vertex lacking distinct microsculpture (Fig. 10J); forewing dark brown with pattern consisting of confluent dots leaving almost no clear space (Fig. 17D); female with head light, vertex with distinct microsculpture (Fig. 11A); forewing with dense, rarely confluent brown dots (Fig. 17E). PtL/FL < 0.3; Cu_1a_L/FL < 0.3. Paramere slender (Fig. 28H). FP/HW > 1.0, FP/SP < 2.0, FP/CRL > 2.7; dorsal margin of female proctiger, in lateral view, posterior to circumanal ring almost straight (Fig. 28L). On *Guatteria sellowiana. **M. dimorpha*** sp. nov.

- Coloration and vertex microsculpture not sexually dimorphic: both male and female vertex with distinct microsculpture (Fig. 11B); forewing covered with sparse brown dots (Fig. 17F). PtL/FL > 0.3; Cu_1a_L/FL ≥ 0.3. Paramere robust (Fig. 28N). FP/HW < 1.0, FP/SP > 2.0, FP/CRL < 2.7; dorsal margin of female proctiger, in lateral view, slightly sinuate distal to circumanal ring, with an upturned apex (Fig. 28R). Perhaps on *Miconia formosa **M. cabucu*** sp. nov.

33 Paramere broadly lanceolate (Figs 28T, 29B). …34

- Paramere acuminate (Fig. 29H, N) or narrowly lanceolate and then blunt apically (Figs 29T, 30B). …35

34 Head and thorax yellow. Forewing pale yellow (Fig. 17G); FL/FW ≥ 2.3. Ventral process of distal segment of aedeagus relatively short (Fig. 28U). Apex of female proctiger, in lateral view, pointed and slightly upturned (Fig. 28X). On *Pleroma granulosum **M. itatiaia*** sp. nov.

- Head and thorax dark brown. Forewing amber-coloured (Fig. 17H); FL/FW < 2.3. Ventral process of distal segment of aedeagus relatively long (Fig. 29C). Apex of female proctiger, in lateral view, truncate and almost straight (Fig. 29F). On *Miconia pusilliflora **M. pusilliflorae*** sp. nov.

35 PtL/FL < 0.3, Cu_1a_L/FL < 0.3; veins slightly lighter than wing membrane which is irregularly light yellow (Fig. 17I). Paramere, in lateral view, narrow (Fig. 29H), with a narrow apex, in dorsal view (Fig. 29K). Ventral process of distal segment of aedeagus short (Fig. 29I, J). FP/HW > 0.9. On *Miconia manauara* and *M. phanerostila **M. goldenbergi*** sp. nov.

- PtL/FL ≥ 0.3, Cu_1a_L/FL > 0.3; veins concolourous with wing membrane which is uniformly dark yellow (Figs 17J, 18A, B). Paramere, in lateral view, broad (Figs 29N, T, 30B), with broad apex, in dorsal view (Figs 29Q, W, 30E). Ventral process of distal segment of aedeagus long (Figs 29O, U, 30C). FP/HW < 0.9. 36

36 Apical metatibial spurs grouped, the two groups anteriorly separated by several bristles (Fig. 6B). Paramere narrow apically, lacking sclerotised tooth (Fig. 29N, Q). Apical dilation of distal segment of aedeagus, in lateral view, narrow subapically (Fig. 29O). Apex of female proctiger, in lateral view, slightly upturned (Fig. 29R). On *Miconia pusilliflora* and *M. sellowiana **M. clavata*** sp. nov.

- Metatibia with a posteriorly open crown of apical spurs, lacking bristles. Paramere broad apically, with a small sclerotised tooth (Figs 29T, W, 30B, E). Apical dilation of distal segment of aedeagus, in lateral view, broad subapically (Figs 29U, 30C). Apex of female proctiger, in lateral view, not upturned (Figs 29X, 30F). 37

37 MT > 0.4, MT/HW > 0.8. Anterior margin of paramere, in lateral view, strongly convex (Fig. 29T). Dorsal margin of female proctiger, in lateral view, slightly convex distal to circumanal ring (Fig. 29X). On *Tibouchina* sp. ***M. tibialis*** sp. nov.

- MT < 0.4, MT/HW < 0.8. Anterior margin of paramere, in lateral view, weakly convex (Fig. 30B). Dorsal margin of female proctiger, in lateral view, almost straight distal to circumanal ring (Fig. 30F). On *Pleroma raddianum* and *P. sellowianum **M. pleromatos*** sp. nov.

38 Ventral process of distal aedeagal segment less than half as long as apical dilation (e.g. Fig. 30I, O, U). … ***ingariko*-group** 39

- Ventral process of distal aedeagal segment more than half as long as apical dilation (e.g. Fig. 32C, I, O, U) 45

39 Surface spinules in cell r_2_ of forewing above bifurcation of M arranged in single-row hexagons. ….40

- Surface spinules in cell r_2_ of forewing above bifurcation of M arranged in double-row hexagons. …42

40 Ventral process of distal segment of aedeagus very short, in lateral view, contiguous with apical dilatation; sclerotised end tube inserted at midlength of apical dilation (Fig. 30I). Apex of female proctiger, in lateral view, upturned (Fig. 30L). On *Miconia chrysophylla. **M. ingariko*** sp. nov.

- Ventral process of distal segment of aedeagus long, in lateral view, separated from apical dilatation; sclerotised end tube inserted in apical third of apical dilation (Fig. 30O, U). Apex of female proctiger, in lateral view, straight (Fig. 30R, X). …41

41 Dark pattern on body and forewing strongly contrasted (Fig. 18D). PtL/PtW < 6.0). MT < 0.4; MT/HW < 0.8; MT/MF < 1.4. Paramere, in lateral view, relatively broad in basal half, strongly narrowing towards apex in apical half (Fig. 30N). Basal portion of distal segment of aedeagus, in lateral view, sinuate along dorsal margin (Fig. 30O). FP/HW < 0.9; FP/SP > 1.5; dorsal margin of female proctiger, in lateral view, slightly sinuate (Fig. 30R). On *Xylopia brasiliensis* and *X. emarginata **M. brasiliensis*** sp. nov.

- Dark pattern on body and forewing weakly contrasted (Fig. 18E). PtL/PtW > 6.0. MT ≥ 0.4; MT/HW > 0.8; MT/MF > 1.4. Paramere, in lateral view, narrow in basal half, weakly narrowing towards apex in apical half (Fig. 30T). Basal portion of distal segment of aedeagus, in lateral view, almost straight along dorsal margin (Fig. 30U). FP/HW > 0.9; FP/SP < 1.5; dorsal margin of female proctiger, in lateral view, straight (Fig. 30X). On *Miconia* aff. *neourceolata **M. caramambatai*** sp. nov.

42 Paramere, in lateral view, narrow (Fig. 31B, H); apex, in dorsal view, subacute (Fig. 31E, K). Apex of female proctiger, in lateral view, almost straight (Fig. 31F, L). …43

- Paramere, in lateral view, broad (Fig. 31N, U); apex, in dorsal view, blunt or truncate (Fig. 31Q, X). Apex of female proctiger, in lateral view, upturned (Fig. 31R, Y). …44

43 Forewing yellow; dark dots dense, confluent and covering most of the surface apically (Fig. 18F). Paramere, in lateral view, with apex directed anteriad (Fig. 31B); in dorsal view with a single small tooth (Fig. 31E). Apical part of proximal aedeagal segment much longer than wide; apical dilation of distal segment of aedeagus, in lateral view, widening towards apex (Fig. 31C). Setae on apical third of female proctiger long (Fig. 31F); FP/SP < 1.7. On *Guatteria* sp ***M. acuminata*** sp. nov.

- Forewing yellow to almost colourless, dark dots spaced, rarely confluent, leaving surface mostly clear (Fig. 18G). Paramere, in lateral view, with apex directed upwards (Fig. 31H); in dorsal view, with two minute teeth (Fig. 31K). Apical part of proximal aedeagal segment about as long as wide; apical dilation of distal segment of aedeagus, in lateral view, with subparallel margins (Fig. 31I). Setae on apical third of female proctiger very short (Fig. 31L); FP/SP > 1.7. On *Miconia albicans* and *M. rubiginosa. **M. melanocephala*** sp. nov.

44 Paramere, in lateral view, broadest in apical third (Fig. 31N). Ventral process of distal segment of aedeagus short and broad; apical dilation, in lateral view, narrow (Fig. 31O, P). Dorsal margin of female proctiger, in lateral view, weakly sinuate distal to circumanal ring; FP/CRL > 2.4; FP/SP > 2.1; apex of female subgenital plate, in lateral view, broad, directed posteriad (Fig. 31R); in ventral view, broadly truncate (Fig. 31S). Perhaps on *Miconia trianae*

… *M. brevicauda* sp. nov.

- Paramere, in lateral view, broadest in the middle (Fig. 31U). Ventral process of distal segment of aedeagus long and slender; apical dilation, in lateral view, broad (Fig. 31V, W). Dorsal margin of female proctiger, in lateral view, strongly sinuate distal to circumanal ring; FP/CRL < 2.4; FP/SP < 2.1; apex of female subgenital plate, in lateral view, narrow, upturned (Fig. 31Y); in ventral view, narrowly truncate. On *Miconia* cf. *cephaloides* … ***M. michali*** sp. nov.

45 Ventral process of distal aedeagal segment, in lateral view, straight or hardly curved and with short apico-median lobe (e.g. Fig. 32C, I, O, U); in dorsal view, apico-median lobe mostly visible (Fig. 32D, J, P, V). ***duckei*-group** 46

- Ventral process of distal aedeagal segment, in lateral view, distinctly curved and with long apico-median lobe (e.g. Fig. 35C, I, O, U); in dorsal view, apico-median lobe mostly hidden below the plane of lateral lobes (Fig. 35D, J, P, V). … ***smithi*-group** 54

46 Paramere, in lateral view, slender, not widened medially (Fig. 32B, H, N, T). …47

- Paramere, in lateral view, robust, widened medially (Figs 33B, I, O, U, 34B). …50

47 FL/FW ≤ 2.1. MT/HW < 0.65. Paramere, in lateral view, with posterior margin sinuate and apex directed anteriad (Fig. 32B); in dorsal view, apex bearing small tooth (Fig. 32E). Basal portion of distal aedeagal segment, in lateral view, distinctly sinuate dorsally; sclerotised end tube relatively long (Fig. 32C, D). FP/HW < 0.8. ***M. duckei*** sp. nov.

- FL/FW > 2.1. MT/HW > 0.65. Paramere, in lateral view, with posterior margin straight and apex directed upwards (Fig. 32H, N, T); in dorsal view, apex blunt, lacking tooth (Fig. 32K, Q, W). Basal portion of distal segment of aedeagus, in lateral view, weakly sinuate dorsally; sclerotised end tube relatively short (Fig. 32I, J, O, P, U, V). FP/HW > 0.8. …48

48 FL/FW > 2.4; PtL/PtW > 6; Cu_1a_L/cu_1_W > 4; forewing pattern consisting of sparse dots, lacking antero-basally (Fig. 19A). Distal segment of aedeagus, in lateral view, relatively broad in basal half (Fig. 32I). FP/HW > 0.9; FP/SP < 1.7. On *Miconia burchellii **M***.

*burchellii* sp. nov.

- FL/FW ≤ 2.4; PtL/PtW < 6; Cu_1a_L/cu_1_W < 4; forewing pattern consisting of relatively dense dots, present also antero-basally (Fig. 19B, C). Distal segment of aedeagus, in lateral view, relatively narrow in basal half (Fig. 32O, U). FP/HW < 0.9; FP/SP > 1.7. …49

49 Paramere, in lateral view, relatively robust in apical half (Fig. 32N). FP/SP > 1.8. Perhaps on *Guatteria schomburgkiana **M. barretoi*** sp. nov.

- Paramere, in lateral view, relatively slender in apical half (Fig. 32T). FP/SP < 1.8. Perhaps on *Xylopia sericea **M. sericeae*** sp. nov.

50 FL/FW > 2.3, FL/HW > 2.7; forewing pattern as in Fig. 19D. Ventral process of distal aedeagal segment, in dorsal view, suboval, weakly produced laterally; apical dilation, in lateral view, weakly curved (Fig. 33D–F). FP/HW < 0.8. On *Miconia* cf. *paniculata* and *M. tristis* subsp. *australis. **M. tristis*** sp. nov.

- FL/FW < 2.3, FL/HW < 2.7. Ventral process of distal segment of aedeagus, in dorsal view, strongly produced laterally; apical dilation, in lateral view, straight (Figs 33J, K, P, Q, V, W, 34D, E). FP/HW > 0.8. …51

51 Paramere, in lateral view, broad subapically, bearing posterior lobe in basal half (Fig. 33I). Ventral process of distal aedeagal segment, in lateral view, situated slightly proximal of the middle of segment (Fig. 33J). On *Miconia discolor*, perhaps also on *M. buddlejoides **M***.

***discoloris*** sp.nov.

- Paramere, in lateral view, narrow subapically; lacking posterior lobe in basal half (Figs 33O, U, 34B). Ventral process of distal aedeagal segment, in lateral view, situated distal of the middle of segment (Figs 33P, V, 34D). …52

52 Paramere narrow (Fig. 33O). Ventral process of distal aedeagal segment almost without apico-median lobe (Fig. 33P, Q). FP/HW < 0.9; dorsal margin of female proctiger, in lateral view, upturned apically (Fig. 33S). Perhaps on *Miconia cinerascens **M. truncata*** sp. nov.

- Paramere robust (Figs 33U, 34B). Ventral process of distal aedeagal segment bearing a distinct apico-median lobe (Figs 33V, W, 34D, E). FP/HW > 0.9 or dorsal margin of female proctiger, in lateral view, straight apically. ….53

53 Ventral process of distal aedeagal segment, in lateral view, short relative to apical dilation (Fig. 33V); in dorsal view, only slightly wider than apical dilation (Fig. 33W). Dorsal margin of female proctiger, in lateral view, weakly concave in apical third, straight apically; setae on female subgenital plate long (Fig. 33Y). On *Miconia cinnamomifolia*, perhaps also on *Miconia* cf. *petropolitana **M. atlantica*** sp. nov.

- Ventral process of distal aedeagal segment, in lateral view, long relative to apical dilation (Fig. 34D); in dorsal view, much wider than apical dilation (Fig. 34E). Dorsal margin of female proctiger, in lateral view, concave in apical third, upturned apically; setae on female subgenital plate short (Fig. 34F). On cf. *Virola* sp ***M. stanopekari*** sp. nov.

54 FL/FW > 2.5; Cu_1a_L/cu_1_W > 4.0; pterostigma and vein Cu_1a_ relatively straight along most of their lengths (Fig. 19I, J). Basal part of distal aedeagal segment, in lateral view, distinctly curved (Fig. 34I, O); ventral process, in dorsal view, cruciform, with narrow wing-like lateral lobes (Fig. 34J, P). ….55

- FL/FW < 2.5; Cu_1a_L/cu_1_W < 4.0; pterostigma and vein Cu_1a_ relatively convex in apical two thirds (Fig. 20A–F). Basal part of distal aedeagal segment, in lateral view, straight (Figs 34U, 35C, I, O, U, 36C); ventral process, in dorsal view, ovoid, with broadly rounded lateral lobes (Figs 34V, 35D, J, P, V, 36D). …56

55 ML/M_1+2_L < 2.1. Anterior margin of paramere, in lateral view, concave subapically (Fig. 34H). Ventral process of distal aedeagal segment short with large lateral lobes (Fig. 34I, J). FP/SP > 1.8. On *Miconia sellowiana **M. sellowianae*** sp. nov.

- ML/M_1+2_L > 2.1. Anterior margin of paramere, in lateral view, straight subapically (Fig. 34N). Ventral process of distal aedeagal segment long with small lateral lobes (Fig. 34O, P). FP/SP < 1.8. On *Pleroma raddianum*, perhaps also on *Miconia pusilliflora* and *P. reitzii*. … *M. simillima* sp. nov.

56 FL/FW ≥ 2.3. MT > 0.5, MT/HW > 0.8, MT/MF > 1.5. Male and female terminalia as in Fig. 34S–X. On *Miconia trianae **M. trianae*** sp. nov.

- FL/FW < 2.3. MT < 0.5, MT/HW < 0.8, MT/MF < 1.5. …57

57 Dark dots on forewing relatively dense; apical part of cell c+sc with many dots (Fig. 20B, C). …58

- Dark dots on forewing relatively sparse, almost completely absent from cell c+sc (Fig. 20D– F). …59

58 DisM/M_1+2_L > 0.87. AL ≥ 0.69. Paramere, in lateral view, broad subapically (Fig. 35B). FP/HW ≥ 0.85. On *Miconia cinerascens* var. *robusta* and *M. pepericarpa **M. cinerascentis***

sp. nov.

- DisM/M_1+2_L ≤ 0.87. AL < 0.69. Paramere, in lateral view, narrower subapically (Fig. 35H). FP/HW < 0.85. On *Miconia crenata*, *M. minutiflora* and *M. pusilliflora **M. minutiflorae***

sp. nov.

59 Brown dots on forewing relatively sparse (Fig. 20D). Paramere, in lateral view, strongly convex posteriorly, wide medially (Fig. 35N). Ventral process of distal aedeagal segment, in dorsal view, only slightly broader than apical dilation (Fig. 35P). On *Miconia cannabina*. … *M. cannabinae* sp. nov.

- Brown dots on forewing relatively dense (Fig. 20E, F). Paramere, in lateral view, weakly convex posteriorly, narrow medially (Figs 35T, 36B). Ventral process of distal segment of aedeagus, in dorsal view, much wider than apical dilation (Figs 35V, 36D). ...60

60 AL < 0.7, AL/HW < 1.15. Surface spinules on forewing forming mostly single-row hexagons (Fig. 6D). Paramere, in lateral view, with slightly narrower apex (Fig. 35T). SP < 0.27; FP/SP > 1.86. On *Miconia alata*, *M. calvescens*, *M. prasina* and *M. splendens/umbrosa*.

… ***M. prasinae*** sp. nov.

- AL > 0.7, AL/HW > 1.15. Surface spinules on forewing forming double-row hexagons. Paramere, in lateral view, relatively robust apically (Fig. 36B). SP > 0.27; FP/SP < 1.86. On *Miconia calvescens. **M. smithi*** (Burckhardt, Morais & Picanço).

Fifth instar immatures

(Following species are not included in the key for the lack of material: *Klyveria flaviae*, *K. setinervis*, *Melanastera barretoi*, *M. cabucu*, *M. curtisetosa*, *M. duckei*, *M. itatiaia*, *M. mazzardoae*, *M. michali*, *M. roraima*, *M. sericeae*, *M. spinosa*, *M. stanopekari*, *M. tijuca*, *M. truncata*, and *M. umbripennis*)

1 Antenna 9-segmented. On Asteraceae. ***Melanastera*** (part) 2

- Antenna 10-segmented. On other plant families. 4

2 AL > 0.7. Metatibiotarsus lacking pointed sectasetae, only with simple setae laterally; MTT

≥ 0.4. On *Moquiniastrum polymorphum **M. moquiniastri*** sp. nov.

- AL < 0.7. Metatibiotarsus with 2–3 pointed sectasetae laterally; MTT < 0.4 3

3 BL > 1.3. Forewing pad lacking dorsal sectasetae, with 5–8 marginal pointed sectasetae. Abdomen anterior to caudal plate with one pointed sectaseta on either side laterally, lacking sectasetae dorsally; caudal plate on either side with 1 + 2 pointed sectasetae laterally and 3 sectasetae dorsally close to circumanal ring (Fig. 36K). On *Eremanthus erythropappus **M***.

*eremanthi* sp. nov.

- BL < 1.3. Forewing pad and abdomen covered with many pointed sectasetae and lanceolate setae dorsally (Fig. 36L). On *Moquiniastrum polymorphum* and *Vernonanthura phosphorica*…. ***M. notia*** sp. nov.

4 Circumanal ring on ventral body side indented medially, distinctly produced antero- sublaterally (cf. Figs 5C, 36I, 38A). On *Luehea candicans*, *L. divaricata*, *L. microcarpa* and *L. paniculata* (Malvaceae). ***Klyveria crassiflagellata*** (Burckhardt)

- Circumanal ring transversely elliptical, anterior margin on ventral body side relatively straight (Fig. 5D). On Annonaceae, Cannabaceae, Melastomataceae or Myristicaceae. ***Melanastera*** (part) 5

5 Caudal plate lacking extra pore fields (Fig. 37A). CW/RW < 3.2. ...6

- Caudal plate with extra pore fields (Fig. 6E, 38). CW/RW > 3.2. ...7

6 AL < 0.33, BL/AL > 3.0. Forewing pad with less than 8 marginal pointed sectasetae. Lanceolate setae on dorsum of wing pads and abdomen narrowly truncate apically. On *Miconia sellowiana. **M. sellowianae*** sp. nov.

- AL > 0.33, BL/AL < 3.0. Forewing pad with more than 8 marginal pointed sectasetae. Lanceolate setae on dorsum of wing pads and abdomen pointed apically. On *Pleroma raddianum*, perhaps also on *Miconia pusilliflora* and *P. reitzii **M. simillima*** sp. nov.

7 Pointed sectasetae or relatively large lanceolate setae present on the abdominal dorsum in addition to lateral sectasetae. ...8

- Pointed sectasetae or relatively large lanceolate setae absent from abdominal dorsum, present only laterally 14

8 Anterior margin of caudal plate with a row of 4 dorsal sectasetae. ...9

- Anterior margin of caudal plate with a row of at least 10 dorsal sectasetae. ...12

9 Antenna with a single sectaseta or a simple seta on segment 8 subapically. Caudal plate with strongly sinuate lateral margins. 10

- Antenna with two sectasetae on segment 8 subapically. Caudal plate with almost straight lateral margins. 11

10 BL > 1.0. Antenna with sectasetae on segments 2, 4, 6, 7 and 8. Perhaps on *Miconia trianae **M. brevicauda*** sp. nov.

- BL < 1.0. Antenna lacking sectasetae, only with simple setae, segment 4 with one slender lanceolate seta. Perhaps on *Xylopia* sp. ***M. parva*** sp. nov.

11 Forewing pad with 9–10 relatively large pointed sectasetae or lanceolate setae marginally and 9–10 similar setae scattered dorsally. MTT < 0.3, MTT/BL < 0.3. On *Miconia burchellii*.

… *M. burchellii* sp. nov.

- Forewing pad with 3–4 relatively large pointed sectasetae marginally and a single sectaseta dorsally, otherwise covered only in minute lanceolate setae. MTT > 0.3, MTT/BL > 0.3. On *Miconia* aff. *neourceolata **M. caramambatai*** sp. nov.

12 AL < 0.53, AL/FL < 1.0, BL/AL > 2.32; MTT < 0.3, MTT/BL < 0.27. 13

- AL > 0.53, AL/FL > 1.0, BL/AL ≤ 2.32; MTT ≥ 0.3, MTT/BL > 0.27. 14

13 Antennal segment 8 with two subapical sectasetae. Forewing pad with 8 marginal sectasetae (in addition to some smaller lanceolate setae). Caudal plate with 4–6 sectasetae near circumanal ring on either side dorsally. CW/RW > 5.5. On *Macairea radula. **M***.

*macaireae* sp. nov.

- Antennal segment 8 with a single subapical sectaseta. Forewing pad with at least 10 marginal sectasetae. Caudal plate with 9–12 sectasetae near circumanal ring on either side dorsally. CW/RW < 5.5. On *Trema micranthum **M. trematos*** sp. nov.

14 BL/BW > 1.4, CW/CL < 1.7. Forewing pad with 9–11 marginal sectasetae and ca. 30 large sectasetae or smaller lanceolate setae dorsally. On *Pleroma granulosum. **M. granulosi*** sp.

nov.

- BL/BW < 1.4, CW/CL > 1.7. Forewing pad with 6–7 marginal sectasetae and ca. 20 large sectasetae or smaller lanceolate setae dorsally. On *Tibouchina* sp. ***M. tibialis*** sp. nov.

15 Antennal segment 8 with two subapical sectasetae. On *Miconia pusilliflora **M***.

*pusilliflorae* sp. nov.

- Antennal segment 8 with a single subapical sectaseta. 16

16 Caudal plate with a large pointed sectaseta medially close to extra pore field on each side of midline. 17

- Caudal plate lacking large sectasetae medially close to extra pore fields (a smaller narrowly lanceolate seta is often present on each side of midline) 20

17 Tarsal arolium long, approximately 1.5 times longer than claws. On *Guatteria* or *Virola*

spp. 18

- Tarsal arolium short, only slightly longer than claws. On *Miconia* spp. 19

18 AL < 0.57, AL/FL < 1.2. Antennal segment 7 lacking a sectaseta, bearing only a fine simple seta subapically. Forewing pad with 5 marginal and 1 dorsal pointed sectasetae; hindwing pad with 2 marginal pointed sectasetae and 1 small dorsal pointed sectaseta. CW/RW < 5.5. On *Guatteria sellowiana **M. dimorpha*** sp. nov.

- AL > 0.57, AL/FL > 1.2. Antennal segment 7 with a sectaseta subapically. Forewing pad with 6 marginal and 4–5 dorsal pointed sectasetae; hindwing pad with 3 marginal and 1–2 dorsal pointed sectasetae. CW/RW > 5.5. On *Virola sebifera **M. falcata*** sp. nov.

19 Forewing pad with 7–8 marginal sectasetae. MTT < 0.3, MTT/BL < 0.25. Caudal plate with 5 pointed sectasetae on either side laterally, and 4 pointed sectasetae on either side subapically. On *Miconia albicans* and *M. rubiginosa. **M. melanocephala*** sp. nov.

- Forewing pad with 5–6 marginal sectasetae. MTT > 0.3, MTT/BL > 0.25. Caudal plate with 4 pointed sectasetae on either side laterally, and 3 pointed sectasetae on either side subapically. On *Miconia trianae. **M. trianae*** sp. nov.

20 On Myristicaceae: *Virola* spp. 21

- On other host families. ...22

21 BL > 1.5. Forewing pad with 8–10 marginal and 0–1 dorsal pointed sectasetae. Caudal plate lacking pointed sectasetae at anterior margin; lateral margins almost straight, bearing 5 pointed sectasetae laterally and 4 pointed sectasetae subapically near circumanal ring on either side dorsally. RW > 0.1, CW/RW < 6.6. On *Virola bicuhyba. **M. virolae*** sp. nov.

- BL < 1.5. Forewing pad with 9–11 marginal and 2 dorsal pointed sectasetae. Caudal plate with 2 pointed sectasetae situated laterally and sublaterally at anterior margin on either side; lateral margins strongly sinuate, bearing 6 pointed sectasetae on two tubercles laterally and 5 pointed sectasetae subapically near circumanal ring on either side dorsally. RW < 0.1, CW/RW > 6.6. On *Virola sebifera. **M. sebiferae*** sp. nov.

22 On Annonaceae. …23

- On Melastomataceae. …35

23 BL/BW > 1.5; AL < 0.5, AL/FL < 1.1, BL/AL > 2.6; MTT/BL ≤ 0.18. Caudal plate

strongly produced posteriorly; CW < 0.5, CW/CL ≤ 1.7. …24

- BL/BW < 1.5; AL > 0.5, AL/FL > 1.1, BL/AL < 2.6; MTT/BL > 0.18. Caudal plate weakly

produced posteriorly; CW > 0.5, CW/CL ≥ 1.7. …25

24 Fore- and hindwing pads only with marginal but lacking dorsal sectasetae. Caudal plate with 4 lateral and 3 subapical sectasetae on either side. On *Guatteria* sp. ***M. acuminata*** sp.

nov.

- Fore- and hindwing pads with 2–4 and 1 dorsal sectasetae, respectively, in addition to marginal sectasetae. Caudal plate with 5 lateral and usually 4 subapical sectasetae on either side. On *Guatteria australis*, perhaps also on *Annona dolabripetala **M. australis*** sp. nov.

25 AL ≥ 0.78, AL/FL ≥ 1.53, MTT ≥ 0.40, MTT/BL ≥ 0.27. Tarsal arolium about twice longer than claws. On *Guatteria punctata **M. olgae*** sp. nov.

- AL ≤ 0.78, AL/FL ≤ 1.53, MTT < 0.40, MTT/BL ≤ 0.27. Tarsal arolium at most 1.5 times longer than claws. …26

26 Forewing pad with 5–7 marginal sectasetae; hindwing pad with 1 dorsal sectaseta close to posterior margin, in addition to 2 marginal sectasetae. On *Xylopia aromatica*, *X. nitida* and *Guatteria* sp ***M. xylopiae*** sp. nov.

- Forewing pad with 4–6 (usually 5) marginal sectasetae; hindwing pad lacking dorsal sectasetae, only with 2 marginal sectasetae. …27

27 Ratio of CL and distance between anterior margin of caudal plate and anterior margin of extra pore fields < 3.5. …28

- Ratio of CL and distance between anterior margin of caudal plate and anterior margin of extra pore fields > 3.5. …31

28 CW/CL > 2.1, CW/RW > 5.0. On *Guatteria schomburgkiana*, perhaps also *Xylopia aromatica **M. tubuligera*** sp. nov.

- CW/CL < 2.1, CW/RW < 5.0. …29

29 BL < 1.35, BW < 1.0, MTT < 0.32. Tarsal arolium about 1.5 times longer than claws. On *Guatteria sellowiana **M. guatteriae*** sp. nov.

- BL > 1.35, BW > 1.0, MTT > 0.32. Tarsal arolium only slightly longer than claws. On *Annona* spp. …30

30 AL < 0.72, BL/AL > 2.2, MTT < 0.35, MTT/BL < 0.22. On *Annona* sp. and perhaps on *Guatteria* sp ***M. nasuta*** sp. nov.

- AL > 0.72, BL/AL ≤ 2.2, MTT > 0.35, MTT/BL ≥ 0.22. On *Annona cacans. **M. obscura*** sp. nov.

31 BL ≤ 1.12, BL/BW ≤ 1.24. On *Xylopia brasiliensis* and *X. emarginata **M. brasiliensis*** sp. nov.

- BL ≥ 1.12, BL/BW ≥ 1.24. …32

32 Caudal plate angular apically, with 4 (2+2) pointed sectasetae on either side laterally and 3 pointed sectasetae subapically. On *Guatteria* sp. ***M. variegata*** sp. nov.

- Caudal plate irregularly rounded apically; with 5–6 pointed sectasetae on either side laterally and 3 pointed sectasetae subapically. …33

33 AL < 0.65, AL/FL < 1.4, MTT ≤ 0.30. On *Annona foetida* and *Guatteria megalophylla*. …

1. *M. amazonica* sp. nov.

- AL > 0.65, AL/FL > 1.4, MTT ≥ 0.30. …34

34 CW > 0.57, CW/CL > 2.0. On *Annona cacans **M. cacantis*** sp. nov.

- CW < 0.57, CW/CL < 2.0. On *Guatteria* sp. and *Xylopia aromatica **M. francisi*** sp. nov.

35 AL/FL < 1.0, BL/AL > 2.4, MTT/BL ≤ 0.21. …36

- AL/FL > 1.0, BL/AL < 2.4, MTT/BL ≥ 0.21. …37

36 BW < 0.85, BL/BW ≥ 1.34. Forewing pad with 5 marginal and 1 dorsal sectasetae, hindwing pad with 2 marginal and 1 dorsal sectasetae, only sparsely covered with inconspicuous minute clavate and lanceolate setae. CW/CL > 1.8, CW/RW > 4.2, caudal plate angular antero-laterally, not tubercular. On *Pleroma raddianum* and *P. sellowianum **M***.

*pleromatos* sp. nov.

- BW > 0.85, BL/BW ≤ 1.34. Forewing and hindwing pads with 5 and 2 marginal sectasetae, respectively, lacking dorsal sectasetae but densely covered with conspicuous small clavate and lanceolate setae. CW/CL < 1.8, CW/RW < 4.2; caudal plate with an antero-lateral tubercle on either side. On *Miconia* cf. *paniculata* and *M. tristis* subsp. *australis **M. tristis***

sp. nov.

37 Tarsal arolium longer than 1.5 times as claws. CW/CL ≤ 1.75. On *Miconia manauara* and *M. phanerostila **M. goldenbergi*** sp. nov.

- Tarsal arolium shorter than 1.5 times as claws. CW/CL > 1.75. …38

38 Semicircular ribbon of extra pore fields on caudal plate, on one side, narrower than a quarter of caudal plate width. …39

- Semicircular ribbon of extra pore fields on caudal plate, on one side, wider than a quarter of caudal plate width. …42

39 BL < 0.8, BL/BW < 1.0, CW/CL > 2.2. Caudal plate with 2 subapical sectasetae on either side of circumanal ring. CW/RW < 3.5. On *Miconia discolor*, perhaps also on *M. buddlejoides **M. discoloris*** sp. nov.

- BL > 0.8, BL/BW > 1.0, CW/CL < 2.2. Caudal plate with 3 subapical sectasetae on either side of circumanal ring. CW/RW > 3.5. …40

40 Body dorsum including wing pads sparsely covered with microscopic clavate and lanceolate setae. On *Miconia chrysophylla **M. ingariko*** sp. nov.

- Body dorsum including wing pads densely covered with small clavate macroscopic and lanceolate setae. …41

41 BL/BW > 1.3, CW/CL < 1.86, CW/RW < 4.5. Hindwing pad with only 2 marginal sectasetae at apex, lacking sectaseta close to posterior margin medio-dorsally. On *Miconia pusilliflora* and *M. sellowiana **M. clavata*** sp. nov.

- BL/BW < 1.3, CW/CL > 1.86, CW/RW > 4.5. Hindwing pad with 2 marginal sectasetae at apex and another one close to posterior margin medio-dorsally. On *Miconia inconspicua*. … *M. inconspicuae* sp. nov.

42 CW/CL < 1.85; caudal plate with only 3 sectasetae on either side basally. On *Miconia cinnamomifolia*, perhaps also on *M.* cf. *petropolitana **M. atlantica*** sp. nov.

- CW/CL > 1.85; caudal plate mostly with 4 sectasetae on either side basally. …43

43 Forewing pad with 1 dorsal sectaseta in addition to marginal sectasetae. On *Miconia cinerascens* var. *robusta* and *M. pepericarpa **M. cinerascentis*** sp. nov.

- Forewing pad lacking dorsal sectasetae. …44

44 AL/FL < 1.2, BL/AL > 2.25. On *Miconia cannabina **M. cannabinae*** sp. nov.

- AL/FL > 1.2, BL/AL < 2.25. …45

45 Small lanceolate and clavate setae on dorsum of body including forewing pads relatively conspicuous and dense. On *Miconia calvescens. **M. smithi*** (Burckhardt, Morais & Picanço)

- Small lanceolate and clavate setae on dorsum of body including forewing pads relatively inconspicuous and sparse. …46

46 On *Miconia crenata*, *M. minutiflora* and *M. pusilliflora **M. minutiflorae*** sp. nov.

- On *Miconia alata*, *M. calvescens*, *M. prasina* and *M. splendens*/*umbrosa **M. prasinae*** sp.

nov.

## *Klyveria* Burckhardt, Serbina & Malenovský

*Klyveria* Burckhardt, Serbina & Malenovský in Burckhardt *et al*., 2023: 409. Type species:

*Paurocephala crassiflagellata* Burckhardt, 1996, by original designation.

**Diagnosis.** Burckhardt *et al*. (2023, 2024a).

### 1 Klyveria flaviae sp. nov

(Figs 3G, 4C, 8A,14A, 21A–G)

**Type material. Holotype m$: Brazil: MATO GROSSO DO SUL:** Rio Verde, MS-427, South of Rio Verde, S19.0330, W54.8583, 470 m, 14.xi.2012, *Luehea grandiflora* (D. Burckhardt & D.L. Queiroz) #69(3) (UFPR; dry).

**Paratypes. Brazil: MATO GROSSO DO SUL:** 11 m$, 10 f£, same as holotype but (UFPR, NHMB; dry, slide, 70% ethanol; NMB-PSYLL0008173, NMB-PSYLL0008611, NMB- PSYLL0008239 [LSKlyfla-5]); 1 f£, Campo Grande, near BR-163, S20.8907/8948, W54.5713/6599, 440–450 m, 16.xi.2012, *Luehea grandiflora* (D. Burckhardt & D.L. Queiroz) #74(5) (NHMB; 70% ethanol; NMB-PSYLL0008176); 9 m$, 7 f£, Jardim, near BR- 267, S21.4494/4683, W55.7919/8027, 380–440 m, 18–20.xi.2012, *Luehea divaricata* (D. Burckhardt & D.L. Queiroz) #76(5) (MMBC, NHMB, UFPR; dry, slide, 70% ethanol; NMB- PSYLL0008178, NMB-PSYLL0008174, NMB-PSYLL0008612, NMB-PSYLL0008613, NMB-PSYLL0008614, NMB-PSYLL0008615, NMB-PSYLL0008616, NMB- PSYLL0008617, NMB-PSYLL0008536); 2 m$, Rochedo, MS-080, S19.9683, W54.6500, 350 m, 15.xi.2012, *Luehea grandiflora* (D. Burckhardt & D.L. Queiroz) #73(3) (NHMB; 70% ethanol; NMB-PSYLL0008175); 1 m$, 6 f£, Sidrolândia, South of Sidrolândia, MS-162, S21.0789, W54.9560, 440 m, 17.xi.2012, *Luehea grandiflora* (D. Burckhardt & D.L. Queiroz) #75(9) (NHMB; 70% ethanol; NMB-PSYLL0008177).—**Minas Gerais:** 1 m$, Lavras, S21.2672, W44.9353, 900 m, 1–6.vi.2010, edge of Atlantic forest around coffee plantation mixed with pastures (D. Burckhardt) #1(0) (NHMB; dry; NMB-PSYLL0008580); 1 f£, Vargem Bonita, Parque Nacional da Serra da Canastra, Cachoeira Casca d’Anta, near waterfall, S20.3090, W46.5231, 860 m, 5.ix.2014 (D. Burckhardt & D.L. Queiroz) #143 (NHMB; 70% ethanol; NMB-PSYLL0008180).—**Paraná:** 1 m$, Curitiba, Parque Barigui, S25.4269, W49.3134, 910 m, 4.xii.2012 (D. Burckhardt & D.L. Queiroz) #85(0) (NHMB; 70% ethanol; NMB-PSYLL0008537).

**Description. *Adult.*** Coloration. General body colour whitish to pale yellow. Antenna pale yellow to brownish, gradually becoming darker towards apex, segments 9–10 entirely dark brown. Forewing (Fig. 14A) with pale yellow veins and clear to pale yellow membrane lacking pattern. Legs pale yellow. Abdomen including terminalia whitish, distinctly paler than thorax.

Structure. Vertex trapezoidal, about half as long as wide, with distinct imbricate microsculpture and short setae (Fig. 8A). Metatibia slender, bearing posteriorly open crown of 8–9 hardly sclerotised apical spurs. Forewing (Fig. 14A) oblong-oval, with subparallel costal and anal margins, widest in the middle, evenly rounded apically; wing apex situated in cell r_2_;

C+Sc moderately curved along most of its length of its section between base and costal break; pterostigma long and narrow, in the middle distinctly narrower than cell r_1_, with subparallel margins in basal two thirds; Rs almost straight, hardly curved to costal margin apically; M longer than M_1+2_ and M_3+4_; Cu_1a_ weakly curved, ending at bifurcation of M; cell cu_1_ short and broad; surface spinules present in all cells, irregularly spaced, leaving narrow spinule-free stripes along the veins; spinules absent from basal two third of cell c+sc. Hindwing with 9 ungrouped costal setae.

Terminalia (Fig. 21A–G). Male proctiger long, distinctly produced posteriad medially; covered with dense, moderately long setae in apical two thirds. Subgenital plate subglobular, with posterior margin evenly convex; with moderately long setae. Paramere shorter than proctiger; in lateral view, lamellar, with anterior margin almost straight in basal two thirds and concave subapically and posterior margin straight to weakly convex in basal half and distinctly concave in apical half, hence irregularly narrowing in apical third, with blunt, upwards directed apex; in dorsal view, apex narrow, directed inwards, and slightly upwards and posteriad, with small apical tooth; in posterior view, outer face strongly convex medially and relatively straight in basal and apical thirds, apex directed upwards; outer and inner face with sparse, moderately long setae, posterior margin with long setae. Proximal segment of aedeagus apically narrow and strongly sinuate, not subdivided. Distal segment of aedeagus thick; dorsal magin almost straight in basal two thirds, ventral margin strongly inflated basally; apical dilation, in lateral view, oval, broadly rounded apically, subdivided from basal portion; in dorsal view, apical dilation narrow, with subparallel margins, narrowly rounded apically; sclerotised end tube short and slightly curved. – Female terminalia cuneate, long; covered with short setae. Dorsal margin of proctiger, in lateral view, distinctly concave in apical half, apex slightly upturned, narrow, blunt; in dorsal view, apex narrowly rounded; circumanal ring, in dorsal view, oval. Subgenital plate slightly shorter than proctiger, with long apical process; in lateral view, pointed, upturned apically; in ventral view, apex narrowly rounded.

*Fifth instar immature.* Unknown.

**Host plant.** Adults were collected on *Luehea divaricata* Mart. and *L. grandiflora* Mart. (Malvaceae) which are likely hosts.

**Distribution.** Brazil (MG, MS, PR).

**Derivation of name.** Dedicated to Flavia Queiroz Sant’Anna, daughter of D.L. Queiroz.

**Comments.** *Klyveria flaviae* sp. nov. differs from both extant species known from Brazil,

*K. crassiflagellata* (Burckhardt) and *K. setinervis* (Burckhardt), in the smaller body size (FL < 1.4 versus > 1.6); distinct imbricate microsculpture on the vertex; the short, light-coloured antenna (AL < 0.6 versus > 0.9; AL/HW < 1.3 versus > 1.4); the evenly pale yellow and elongate forewing (FL/FW > 2.4 versus ≤ 2.4), with subparallel margins, and narrowly rounded apex; and details of the paramere shape bearing a single small apical tooth. *Klyveria flaviae* sp. nov. differs from *K. crassiflagellata* in the broader pterostigma (PtL/PtW < 9.0 versus > 14.0), and from *K. setinervis* in the much shorter setae on head, thorax and forewing veins; the longer vertex relative to the width; the narrower pterostigma (PtL/PtW > 7.0 versus < 7.0), and longer (Cu_1a_L/FL > 0.3 versus < 0.3) and narrower (Cu_1a_L/cu_1_W > 3.0 versus < 2.0) cell cu_1_; and the oval female circumanal ring.

*Klyveria flaviae* sp. nov. differs from the fossil species *K. succina* Burckhardt & Drohojowska described from Dominican amber (Burckhardt *et al*. 2024a) in the smaller body size (HW < 0.5 vs. = 0.6; FL < 1.4 versus = 1.5; MT < 0.4 vs. = 0.5), more elongate forewing with the cell c+sc narrow (vs. broad), vein M longer than M_1+2_ (vs. shorter), vein Cu longer than Cu_1b_ (vs. shorter) and cell cu_1_ broader (vs. narrower).

### 2 Klyveria crassiflagellata (Burckhardt, 1996)

(Figs 7A, 8B, 14B, 21H–N, 36I, 38A)

*Paurocephala crassiflagellata* Burckhardt, 1996: 79.

*Diclidophlebia crassiflagellata*, Burckhardt & Mifsud (2003): 13; Burckhardt & Queiroz (2012): 36.

*Klyveria crassiflagellata*, Burckhardt *et al*. (2023): 410.

**Material examined. Brazil: MATO GROSSO DO SUL:** 4 m$, 8 f£, Campo Grande, near BR- 163, S20.8907/8948, W54.5713/6599, 440–450 m, 16.xi.2012, *Luehea divaricata* (D. Burckhardt & D.L. Queiroz) #74(12) (MMBC, NHMB; dry, 70% ethanol; NMB- PSYLL0008182, NMB-PSYLL0008609); 1 f£, Dourados, Embrapa campus, S22.2738, W54.8182,400 m, 8.x.2021, park vegetation (D. Burckhardt & D.L. Queiroz) #459 (NHMB; 70% ethanol; NMB-PSYLL0008325); 6 m$, 8 f£, 2 immatures, 1 skin, Jardim, near BR-267, S21.4494/4683, W55.7919/8027, 440–450 m, 18–20.xi.2012, *Luehea microcarpa* (D. Burckhardt & D.L. Queiroz) #76(11) (NHMB; 70% ethanol; NMB-PSYLL0008183); 4 m$, 12 f£, 3 immatures, Naviraí, S23.0298, W54.2785, 320 m, 7.x.2021, edge of cerrado, *Luehea divaricata* (D. Burckhardt & D.L. Queiroz) #456(2) (NHMB; 70% ethanol; NMB- PSYLL0008324).—**Minas Gerais:** 6 m$, 4 f£, Belo Horizonte, Parque Estadual Serra Verde, S19.7866, W439518, 790 m, iv.2013, *Luehea candicans* (A.P. Santana) #1 (NHMB; 70% ethanol; NMB-PSYLL0008189).—**Paraná:** 2 m$, 3 f£, Colombo, Embrapa campus, S28.2288, W52.4065, 920 m, 8.x.2019, *Luehea* cf. *divaricata* (D. Burckhardt & D.L. Queiroz) #382(9) (NHMB; dry; NMB-PSYLL0008188, NMB-PSYLL0008610); 7 m$, 12 f£, 8 immatures, 2 skins, Curitiba, Parque Barigui, S25.4152/4255, W49.3072/3092, 900–910 m, 18.ix.2011, *Luehea paniculata* (D. Burckhardt & D.L. Queiroz) #10(5) (NHMB, UFPR; dry, slide, 70% ethanol; NMB-PSYLL0008118, NMB-PSYLL0008181, NMB-PSYLL0008206 [LSWolcrass], NMB-PSYLL0008212 [LSWolcrass], NMB-PSYLL0008323, NMB- PSYLL0008606–NMB-PSYLL0008608); 1 f£, same but S25.4433, W49.3133, 910 m, 9.ii.2016 (D. Burckhardt & D.L. Queiroz) #195 (NHMB; dry; NMB-PSYLL0008186); 3 m$, 2 f£, 15 immatures, same but Parque Passaúna, S25.4750, W49.3783, 930 m, 27–30.xi.2012, *Luehea candicans* (D. Burckhardt & D.L. Queiroz) #78(4) (NHMB; slide, 70% ethanol; NMB-PSYLL0005759, NMB-PSYLL0008184); 1 m$, same but Jardim Botânico, S25.4416/4433, W49.2383/2833, 930 m, 15.ii.2013 (D. Burckhardt & D.L. Queiroz) #94A (NHMB; 70% ethanol; NMB-PSYLL0008185).—**Rio Grande di Sul:** 1 m$, 5 f£, Passo Fundo, Embrapa campus, S28.2288, W52.4065, 640 m, 20.ix.2018, *Luehea* sp. (D. Burckhardt & D.L. Queiroz) #306(11) (NHMB; 70% ethanol; NMB-PSYLL0008187).

Description. *Adult.* Burckhardt (1996).

***Fifth instar immature*** (Fig. 38A). Coloration. Pale yellow; antenna with dark brown apical segment.

Structure. Eye with one ocular pointed sectaseta dorsally. Antennal segments with following numbers of pointed sectasetae: 1(0), 2(1), 3(1), 4(2), 5(0–1), 6(2), 7(1), 8(1), 9(0), 10(0). Forewing pad with 11–13 pointed marginal sectasetae; hindwing pad with 2 marginal and 6 dorsal pointed sectasetae. Tarsal arolium broadly fan-shaped apically, 1.2 times as long as claws. Abdomen with five transverse rows of pointed sectasetae dorsally. Anterior margin of caudal plate (Fig. 36I) distant from anterior margin of extra pore fields; with two rows of pointed sectasetae in the middle and along anterior margin dorsally, and four or five pointed sectasetae near circumanal ring on either side dorsally. Extra pore fields forming continuous outer and inner bands, consisting of small oval and round patches; outer band short medially, end pointing outwards. Each half of circumanal ring distinctly bent forwards antero-ventrally.

**Host plants.** Luehea candicans Mart., *L. divaricata* Mart., *L. microcarpa* R.E. Fr. and *L. paniculata* Mart. (Malvaceae).

**Distribution.** Reported from Brazil (PR) (Burckhardt & Ouvrard 2012, as *Diclidophlebia crassiflagellata*); Paraguay (Burckhardt 1996, as *Paurocephala crassiflagellata*). Brazil (MG, MS, PR, RS).

**Comments.** *Klyveria crassiflagellata* differs from *K. flaviae* sp. nov. and *K. setinervis* (Burckhardt) in the larger body size (FL > 1.8 versus < 1.8), the very narrow pterostigma of the forewing (PtL/PtW > 14.0 versus 9.0), and in the paramere, in both lateral and dorsal views, with a broad apex bearing two small sclerotised teeth (versus narrow apex lacking distinct sclerotised teeth). It differs from *K. flaviae* sp. nov. as detailed under the latter species, and from *K. setinervis* in the following characters: the shorter setae on head, thorax and forewing veins; the relatively shorter vertex (VW/VL < 3.0 versus > 3.0); the narrower cell cu_1_ of forewing (Cu_1a_L/cu_1_W > 3.0 versus < 2.0); the oval circumanal ring on the female proctiger (versus cruciform).

### 3 *Klyveria setinervis* (Burckhardt, 1996)

(Figs 8C, 14C, 21O–U)

*Paurocephala setinervis* Burckhardt, 1996: 78.

*Diclidophlebia setinervis*, Burckhardt & Mifsud (2003): 14.

*Klyveria setinervis*, Burckhardt *et al*. (2023): 410.

**Material examined. Brazil: MINAS GERAIS:** 12 m$, 6 f£, Vazante, Fazenda Bainha, near source of Curtume river, S17.8891/8942, W46.9196/9205, 640–690 m, 11.ix.2014 (D. Burckhardt & D.L. Queiroz) #145 (NHMB; dry; NMB-PSYLL0004028, NMB- PSYLL0004033, NMB-PSYLL0004030, NMB-PSYLL0004029, NMB-PSYLL0004031, NMB-PSYLL0004032); 27 m$, 27 f£, same but S17.8913/8841, W46.9163/9221, 660–670

m, 29–30.x.2012, *Luehea divaricata* (D. Burckhardt & D.L. Queiroz) #50(13) (NHMB; slide, 70% ethanol; NMB-PSYLL0008190, NMB-PSYLL0008179, NMB-PSYLL0008618, NMB- PSYLL0008535 [LSWolset], NMB-PSYLL0008211); same but S17.88815, W46.92126, 630 m, 7.iii.2019 (D.L. Queiroz) #922(4) (NHMB; 70% ethanol; NMB-PSYLL0008192).—**Paraná:** 8 m$, 1 f£, Curitiba, Parque Passaúna, S25.4750, W49.3783, 930 m, 27–30.xi.2012, *Luehea divaricata* (D. Burckhardt & D.L. Queiroz) #78(24) (NHMB; 70% ethanol; NMB- PSYLL0008191).—**Rio Grande do Sul:** 35 m$, 53 f£, Passo Fundo, Embrapa campus, S28.2288, W52.4065, 640 m, 23.x.2013, *Luehea divaricata* (A.L. Marsaro Jr.) (MMBC, NHMB, UFPR; dry, 70% ethanol; NMB-PSYLL0003908, NMB-PSYLL0008193, NMB- PSYLL0008619, NMB-PSYLL0008620); 5 m$ 4 f£, same but 23.x.2013 (A.L. Marsaro Jr.) (NHMB; 70% ethanol; NMB-PSYLL0003908).

Description. *Adult.* Burckhardt (1996).

*Fifth instar immature.* Unknown.

**Host plant.** Adults were collected on *Luehea divaricata* Mart. (Malvaceae) which is a likely host.

**Distribution.** Reported from Brazil (RS) (Marsaro *et al*. 2021, as *Diclidophlebia setinervis*); Paraguay (Burckhardt 1996, as *Paurocephala setinervis*). Brazil (MG, PR, RS).

**Comments.** Klyveria setinervis differs from *K. crassiflagellata* (Burckhardt) and *K*. *flaviae*

sp. nov. as detailed under these species.

## *Melanastera* Serbina, Malenovský, Queiroz & Burckhardt

*Melanastera* Serbina, Malenovský, Queiroz & Burckhardt in Burckhardt *et al*., 2023: 411. Type species: *Diclidophlebia lucens* Burckhardt, Hanson & Madrigal, 2005, by original designation.

**Diagnosis.** Burckhardt *et al*. (2023).

**Comments.** *Melanastera* includes twelve previously described species of which six are from Central America, one each from Brazil, Mexico, East Africa and China, and two are fossils from Dominican amber (Burckhardt *et al*. 2023, 2024a; He *et al*. 2024). Here, 59 new species are added, bringing the number of Brazilian species to 60 (71 spp. worldwide).

*Melanastera* is a morphologically homogeneous genus with often small differences between closely related species. In order to keep species descriptions as brief as possible and avoid unnecessary repetition, nine species groups are defined here on the basis of morphology, most of which are artificial and reflect the structure of the key for adults rather than phylogenetic relationships. Brown & Hodkinson (1988) defined the *paucipunctata*-group for three Panamanian species and one each from Mexico and Puerto Rico. They did not define any other species groups and left one additional Panamanian species without group affiliation. The *paucipunctata*-group differs in definition and content from all species groups defined here.

For eight of the twelve previously described species we have material at hand. These species are assigned to one of our species groups and are discussed in the current study under species descriptions. For four species we lack material and the original descriptions (Caldwell & Martorell 1952, Conconi 1972, Brown & Hodkinson 1988) do not allow us to classify them with certainty into one of the species groups. The four species develop on the following hosts not occurring in Brazil, suggesting that these psyllid taxa are not conspecific with any of the new Brazilian species described here: *Miconia argentea* (Sw.) DC., host of *M. fava* (Brown & Hodkinson) and *M. paucipunctata* (Brown & Hodkinson), *Miconia umbellata* (Mill.) Judd & Ionta (= *Heterotrichum cymosum* (Wendl. ex Spreng.) Urb., host of *M. heterotrichi* (Caldwell & Martorell); and *Miconia xalapensis* (Bonpl.) M.Gómez (= *Conostegia xalapensis* (Bonpl.) D.Don ex DC.), host of *M. paucipunctata* (Brown & Hodkinson) and *M. tuxtlaensis* (Conconi).

## The curtisetosa-group

**Description. *Adult*.** General body colour light (Fig. 7B). Head (Fig. 8D–G) and thorax covered with long setae. Head, in lateral view, inclined at < 45° from longitudinal body axis. Vertex trapezoidal, 1.8–2.2 times as long as wide, with imbricate microsculpture and long macroscopical setae. AL 0.8–11, AL/HW 1.2–1.9. Thorax weakly to moderately arched.

Forewing (Fig. 14D–G) oblong-oval, with subparallel costal and anal margins, widest in the middle; wing apex in cell r_2_ near apex of M_1+2_; pterostigma with subparallel margins, weakly or not widening towards apex (in one Oriental species, the forewing is broadly oval, with divergent costal and anal margins, widest in apical third, with wing apex in the middle of cell r_2_ and pterostigma strongly widening to middle). Distal segment of aedeagus lacking ventral process. Dorsal margin of female proctiger, in lateral view, weakly or strongly sinuate distal to circumanal ring. ***Immature*.** Antenna 9-segmented (in Brazilian species) or 10-segmented (in Afrotropical and Oriental species).

**Comments.** In addition to the four Brazilian species described here, the *curtisetosa*-group includes *Melanastera pilosa* (Burckhardt, Aléné, Ouvrard, Tamesse & Messi) from Kenya and Tanzania, developing on *Grewia bicolor* Juss. (Malvaceae) (Burckhardt *et al*. 2006a), *M. sinica* He & Burckhardt from China (Hainan island), developing on *Grewia* cf. *chuniana* Burret (Malvaceae) (He *et al*. 2024) and two fossil species from Miocene Dominican amber:

*M. casca* Burckhardt & Drohojowska and *M. vetus* Burckhardt & Drohojowska (Burckhardt *et al*. 2024a). Adults of *M. pilosa* and *M. sinica* differ from the Brazilian taxa in having a shorter and broader forewing with much more numerous and denser dark dots (in *M. pilosa*) or dark bands (in *M. sinica*), which are also present in cell c+sc. *Melanastera sinica* also differs from the Brazilian species in the broader, medially widened pterostigma and broader, subrectangular paramere (He *et al*. 2024). The fifth instar immatures of *M. pilosa* and *M*.

*sinica* differ from the Brazilian species of the *curtisetosa-*group by having a 10-segmented antenna, a forewing pad with 3–4 (in *M. pilosa*) or 5 (in *M. sinica*) marginal sectasetae and a tarsal arolium about twice as long as the claws (versus 9-segmented antenna, forewing pad with 5 or more marginal sectasetae and arolium about as long as claws or slightly longer). The two fossil species differ from the extant Brazilian species as follows: *M. casca* by the relatively broader forewings (FL/FW = 2.2 versus ≥ 2.4) and the relatively short female proctiger with a nearly straight dorsal outline (versus relatively long with a distinctly sinuate or concave dorsal outline), and *M. vetus* by the forewing with dark dots restricted to the apices of the veins along the margin but otherwise absent on the cells (versus with dark dots either absent altogether or present on most cells) (Burckhardt *et al*. 2024a). The species of the *curtisetosa*-group are associated with Asteraceae (in Brazil) and Malvaceae (in Africa and Asia). The four Brazilian species and the two Dominican amber species probably form a monophyletic group (Burckhardt *et al*. 2024a).

### 4 Melanastera curtisetosa sp. nov

(Figs 3F, 8D, 14D, 21V–AA)

**Type material. Holotype m$: Brazil: RIO GRANDE DO SUL:** Passo Fundo, S28.2288, W52.4061, 670 m, 25.x.2013 (A.L. Marsaro Jr.) (UFPR; dry).

**Paratypes. Brazil: RIO GRANDE DO SUL:** 4 f£, Passo Fundo, Embrapa campus, S24.5621, W50.2581, 28.vi.2013, degraded vegetation (D.L. Queiroz) #520 (NHMB; dry; NMB- PSYLL0007806); 3 m$, 4 f£, same as holotype but (NHMB, MMBC, UFPR; dry, slide, 70% ethanol; NMB-PSYLL0003909, NMB-PSYLL0007838 [LSMelcur-68], NMB- PSYLL0007839 [LSMelcur-68], NMB-PSYLL0007840 [LSMelcur-68], NMB- PSYLL0007841 [LSMelcur-68]).

**Description. *Adult.*** Coloration. Body pale yellow to pale orange. Antenna pale yellow; segments 3–8 with dark brown apices, segment 9 at least in apical half and segment 10 entirely dark brown. Forewing (Fig. 14D) veins and membrane pale yellow with pale brown dots sparsely scattered in apical two thirds of wing (sometimes indistinct); apices of veins Rs, M_1+2_, M_3+4_ and Cu_1a_ at wing margin pale brown, apex of vein Cu_1b_ brown. Legs pale yellow, metacoxa brown with pale yellow meracanthus; metafemur with pale brown spots. Abdomen pale yellow. Terminalia pale yellow to pale brown; female proctiger with brown apex.

Structure. Setae on thorax moderately long. Forewing (Fig. 14D) evenly, narrowly rounded apically; C+Sc distinctly curved in apical third; pterostigma relatively long and narrow, with subparallel margins, in the middle distinctly narrower than cell r_1_; Rs almost straight along most of its length, hardly curved to fore margin apically; M longer than M_1+2_ and M_3+4_; Cu_1a_ weakly curved in apical quarter, ending distad of M fork; cell cu_1_ long and narrow; surface spinules present in all cells, dense, leaving narrow spinule-free stripes along the veins, not forming distinct hexagons at least in apical part of the wing, spinules absent in basal part of cell c+sc. Costal setae of hindwing grouped (6 + 4–5). Metatibia bearing 7–8 grouped apical spurs, arranged as 3 + 4–5, anteriorly separated by 4–5 bristles.

Terminalia (Fig. 21V–AA). Male. Proctiger long, broad, posterior margin broadly convex; densely covered with moderately long setae in apical two thirds. Subgenital plate, in lateral view, irregularly ovoid; dorsal margin curved anteriorly and posteriorly, straight in the middle, posterior margin almost straight; beset with moderately long setae in posterior half. Paramere shorter than proctiger; in lateral view, lamellar, with mostly subparallel margins; anterior margin straight in basal four fifths, strongly bent in apical fifth, posterior margin weakly, irregularly curved; apex, in lateral view, narrowly rounded, directed upwards, in dorsal view, broadly rounded, directed slightly inwards, lacking a distinct sclerotised tooth; outer face with dense, moderately long setae in apical half; inner face with dense, moderately long setae medially and short setae apically; posterior margin with long setae. Proximal segment of aedeagus with apical part moderately subdivided. Distal segment of aedeagus almost straight with dorsal margin sinuate; apical dilation, in lateral view, narrowly elongate, evenly rounded apically; sclerotised end tube short and weakly curved. – Female terminalia cuneate, long; with dense setae. Dorsal margin of proctiger, in lateral view, distinctly sinuate distal to circumanal ring; apex upturned, pointed; in dorsal view, apex pointed; circumanal ring, in dorsal view, distinctly cruciform. Subgenital plate distinctly shorter than proctiger, covered with numerous short setae in apical half; in lateral view, abruptly narrowing in apical third, with pointed apex; in ventral view, apex truncate, incised medially.

*Fifth instar immature.* Unknown.

**Host plant.** Adults were collected on an unidentified species of Asteraceae, which is possibly a host.

**Distribution.** Brazil (RS).

**Derivation of name.** From the Latin adjectives *curtus* = shortened and *setosus* = coarsely hairy, referring to the short setae on the female subgenital plate and apex of the proctiger.

### 5 Melanastera eremanthi sp. nov

(Figs 5A, B, 8E, 14E, 22A–F, 36K, 39A, B)

**Type material. Holotype m$: Brazil: MINAS GERAIS:** Ouro Preto, Parque Estadual do Itacolomi, S20.4326, W43.5119, 1300 m, 16.xi.2018, *Eremanthus erythropappus* (D.L. Queiroz) #912(3) (UFPR; dry).

**Paratypes. Brazil: MINAS GERAIS:** 23 m$, 15 f£, 62 immatures, 13 skins, Diamantina, Clube Campestre, S18.1451, W43.3889, 1290 m, 12.ix.2021, *Eremanthus* cf. *erythropappus* (D.L. Queiroz) #1008(2) (NHMB; 70% ethanol; NMB-PSYLL0008533); 7 m$, 9 f£, 24 immatures, 2 skins, same as holotype but (MMBC, NHMB, UFPR; dry, slide, 70% ethanol; NMB-PSYLL0007860, NMB-PSYLL0007993, NMB-PSYLL0007868 [LSMelere-103], NMB-PSYLL0007869 [LSMelere-103], NMB-PSYLL0007870 [LSMelere-103], NMB- PSYLL0007871 [LSMelere-103], NMB-PSYLL0007872, NMB-PSYLL0007873); 1 f£, same but S20.4326, W43.5119, 1300 m, 9.i.2019, *Eremanthus erythropappus* (R. Caffaro & N. Jorge) PF1 (NHMB; 70% ethanol; NMB-PSYLL0007859); 2 immatures, same but S20.4326, W43.5119, 1300 m, 27.iii.2019, *Eremanthus erythropappus* (R. Caffaro & N. Jorge) PF1 (NHMB; 70% ethanol; NMB-PSYLL0007858); 2 f£, same but Trevo da Chapada, Cachoeira do Moinho, S20.4530, W43.5523, 1240 m, 16.xi.2018, *Eremanthus erythropappus* (D.L. Queiroz) #914 (NHMB; 70% ethanol; NMB-PSYLL0007829).

**Description. *Adult.*** Coloration. Body pale yellow. Head (Fig. 8E) brown. Antennal segments 4–8 with dark brown apices; segments 9–10 entirely dark brown. Thorax pale yellow to orange. Forewing (Fig. 14E) with whitish to pale yellow veins and pale yellow, almost colourless membrane lacking dark dots. Legs pale yellow, metacoxa dark brown with pale yellow meracanthus. Abdominal sclerites brown; female proctiger with brown apex. Younger specimens with less expanded dark colour.

Structure. Setae on thorax moderately long. Forewing (Fig. 14E) narrowly and evenly rounded apically; C+Sc evenly curved; pterostigma short to long and narrow, distinctly narrower in the middle than r_1_ cell; Rs almost straight, hardly curved to costal margin apically; M longer than M_1+2_ and M_3+4_; Cu_1a_ weakly evenly curved, ending slightly distad of M fork; cell cu_1_ narrow; surface spinules present in all cells, faint, leaving narrow spinule-free stripes along the veins, forming indistinct hexagons of a single row of spinules, spinules absent from basal half of cell c+sc. Costal setae of hindwing grouped (5–6 + 4). Metatibia bearing 5–6 grouped apical spurs, arranged as 2–3 and 2–3, anteriorly separated by 5–7 bristles.

Terminalia (Fig. 22A–F). Male. Proctiger broad, with posterior margin widely curved; densely covered with moderately long setae in apical two thirds. Subgenital plate, in lateral view, subtrapezoidal, dorsal margin strongly angular in proximal third, irregularly curved in distal two thirds; posterior margin relatively straight; with long setae, mostly posteriorly. Paramere shorter than proctiger; in lateral view, irregularly lanceolate, anterior margin sinuate, concave submedially, convex in apical half, posterior margin convex in apical three quarters; paramere apex, in lateral view, blunt, directed upwards and slightly anteriad, in dorsal view, apex subacute, directed inwards, slightly anteriad, lacking sclerotised tooth; outer face with dense, moderately long setae, mostly in apical half; inner face with dense, short setae; posterior margin with long setae. Proximal segment of aedeagus with apical part moderately subdivided. Distal segment of aedeagus almost straight in basal half, with dorsal margin sinuate; apical dilation, in lateral view, narrow, elongate, with subparallel margins and broadly rounded apically, with a small membranous extension at base dorsally; in dorsal view, apical dilation relatively broad, broadest subapically, rounded apically, sclerotised end tube short and curved. – Female terminalia cuneate; with dense, mostly short setae. Dorsal margin of proctiger, in lateral view, with a large hump distal to circumanal ring, strongly concave subapically, apex distinctly upturned, pointed; in dorsal view, apex subacute; circumanal ring, in dorsal view, vaguely cruciform. Subgenital plate shorter than proctiger, covered with numerous short setae in apical half; in lateral view, abruptly narrowing towards apex in apical third, with pointed apex; in ventral view, apex truncate.

***Fifth instar immature.*** Coloration. Pale yellow, sometimes with pale to dark brown pattern; antenna pale yellow, gradually becoming darker towards apex.

Structure. Eye with one long, simple ocular seta dorsally. Antennal segments with following numbers of pointed sectasetae: 1(0), 2(1), 3(2), 4(0), 5(2), 6(1), 7(1), 8(0), 9(0). Forewing pad with 5–8 marginal pointed sectasetae; hindwing pad with two marginal pointed sectasetae; fore- and hindwing pads lacking dorsal sectasetae. Metatibiotarsus long, with 2–3 pointed sectasetae on outer side; tarsal arolium broadly triangular apically, about as long as claws (Fig. 5B, 37D). Abdomen lacking dorsal pointed sectasetae proximal to caudal plate, with one pointed sectaseta on either side laterally. Caudal plate (Fig. 5A) with anterior margin close to anterior margin of extra pore fields; with 1 + 2 pointed sectasetae on either side laterally, and three pointed sectasetae on either side of circumanal ring dorsally. Extra pore fields forming continuous outer and inner bands, consisting of several large oval and rounded patches; outer band long medially, end pointing outwards. Circumanal ring broad, with anterior margin distinctly produced cephalad.

**Host plant and biology.** *Eremanthus erythropappus* (DC.) MacLeish (Asteraceae).

Immatures induce leaf roll galls in which they develop (Fig. 39A, B).

**Distribution.** Brazil (MG).

**Derivation of name.** Named after its host, *Eremanthus*.

### 6 Melanastera moquiniastri sp. nov

(Figs 5C, 7B, 8F, 14F, 22G–L, 36J, 39C–F)

**Type material. Holotype m$: Brazil: SANTA CATARINA:** Urubici, Pousada Kiriri-Eté, S27.9183, W49.5700, 1180 m, 1.v.2013, *Moquiniastrum polymorphum* (D. Burckhardt & D.L. Queiroz) #120(3) (UFPR; dry).

**Paratypes. Brazil: MINAS GERAIS:** 1 f£, Barroso, Mata do Baú, S21.1833/2000, W43.9167/9667, 940–1140 m, 13–14.vi.2010, Cerrado semideciduous forest, along forest edge (D. Burckhardt) #7(0) (NHMB; dry; NMB-PSYLL0008584); 2 f£, Lavras, S21.2672, W44.9353, 900 m, 1–6.vi.2010, edge of Atlantic forest around coffee plantation mixed with pastures (D. Burckhardt) #1(0) (NHMB; dry; NMB-PSYLL0008581); 1 m$, 4 f£, Lavras, Universidade Federal de Lavras, UFLA, S21.2308, W44.9914, 900 m, 20.vi.2010, degraded forest margin (D. Burckhardt) #11(0) (NHMB; dry, 70% ethanol; NMB-PSYLL0007513, NMB-PSYLL0008582, NMB-PSYLL0008583).—**Paraná:** 2 m$, 1 f£, Araucária, Chácara do Paulo, S2.8921, W59.9537, 870 m, 5.iv.2014 (D.L. Queiroz) #619 (NHMB; 70% ethanol; NMB-PSYLL0007555); 1 m$, 2 f£, 3 immatures, 2 skins, Castro, Parque Caxambu, S24.6775, W50.0142, 1010 m, 11.vii.2013, Atlantic forest (D.L. Queiroz) #535 (NHMB; 70% ethanol; NMB-PSYLL0007552); 7 m$, 14 f£, 12 immatures, Colombo, Embrapa campus, S25.3200/3350, W49.1566/1683, 920 m, 21.ix.2017 (J.T. Cremonese) #1 (NHMB; 70% ethanol; NMB-PSYLL0007556); 3 m$, 4 f£, 1 immature, same but S25.3226, W49.15686, 930 m, 6.v.2014, *Moquiniastrum polymorphum* (D. Burckhardt & D.L. Queiroz) #135(3) (NHMB; 70% ethanol; NMB-PSYLL0007523); 4 m$, 9 f£, 2 immatures, same but 30.v.2023, *Moquiniastrum polymorphum* (D.L. Queiroz) (NHMB; 70% ethanol; NMB-PSYLL0003249); 1 m$, 2 f£, same but S25.3196/3353, W49.1562/1677, 920 m, 6.i.2012 (D. Burckhardt & D.L. Queiroz) #30 (NHMB; 70% ethanol; NMB-PSYLL0007514); 5 m$, 8 f£, 6 immatures, 1 skin, same but 1.v.2019, *Moquiniastrum polymorphum* (D. Burckhardt & D.L. Queiroz) #339(14) (NHMB; 70% ethanol; NMB-PSYLL0008534); 1 m$, 4 f£, 3 immatures, 5 skins, same but 1– 5.iv.2013, *Moquiniastrum polymorphum* (D. Burckhardt & D.L. Queiroz) #96(5) (NHMB; 70% ethanol; NMB-PSYLL0007520); 12 m$, 18 f£, 113 immatures, 8 skins, same but 28.ix.2017, *Moquiniastrum polymorphum* (I. Malenovský) (MMBC; 70% ethanol); 1 m$, 4 immatures, same but 21.ix.2017 (L. Serbina) (MMBC; 70% ethanol); 7 m$, 12 f£, 4 immatures, Curitiba, Jardim Botânico, S25.4417, W49.2394, 920 m, 12.iii.2013 (D.L. Queiroz) #462 (NHMB; 70% ethanol; NMB-PSYLL0007550, NMB-PSYLL0007512); 2 m$, same but S25.4416/4433, W49.2366/2383, 930 m, 15.ii.2013, *Moquiniastrum polymorphum* (D. Burckhardt & D.L. Queiroz) #94A(1) (NHMB; 70% ethanol; NMB-PSYLL0007518); 2 f£, 1 immature, same but Parque Passaúna, S25.4750, W49.3783, 930 m, 8.ii.2016, *Moquiniastrum polymorphum* (D. Burckhardt & D.L. Queiroz) #195(12) (NHMB; 70% ethanol; NMB-PSYLL0007547); 17 m$, 14 f£, 5 immatures, same but S25.4750, W49.3783, 930 m, 27–30.xi.2012, *Moquiniastrum polymorphum* (D. Burckhardt & D.L. Queiroz) #78(9) (NHMB; 70% ethanol; NMB-PSYLL0007516); 7 m$, 9 f£, 18 immatures, same but Parque Tingui, S25.3950, W49.3052, 870 m, 26.xi.2012, *Moquiniastrum polymorphum* subsp. *floccosum* (D. Burckhardt & D.L. Queiroz) #77(3) (NHMB; 70% ethanol; NMB- PSYLL0007515); 1 m$, 1 f£, 2 immatures, same but S25.3950, W49.3066, 890 m, 31.iii.2013, *Moquiniastrum polymorphum* (D. Burckhardt & D.L. Queiroz) #95(8) (NHMB; 70% ethanol; NMB-PSYLL0007519); 4 m$, 6 f£, 2 immatures, same but Centro Politécnico, UFPR, S25.4474/4485, W49.2312/2383, 890–920 m, 3–7.xii.2012, *Moquiniastrum polymorphum* (D. Burckhardt & D.L. Queiroz) #84(3) (NHMB; 70% ethanol; NMB- PSYLL0007517); 2 m$, 2 f£, 14 immatures, 2 skins, Ibaiti, Fazendinha Nova Esperança, road BR-153, S23.9116, W50.2400, 635 m, 18.ix.2014, *Moquiniastrum polymorphum* (D. Burckhardt & D.L. Queiroz) #148(4) (NHMB; 70% ethanol; NMB-PSYLL0007524); 3 m$, 7 f£, Jaguariaíva, Parque Estadual do Cerrado, S24.1685/1833, W49.6603/6670, 780–820 m, 15–16.ii.2016, *Moquiniastrum polymorphum* (D. Burckhardt & D.L. Queiroz) #197(13) (NHMB; 70% ethanol; NMB-PSYLL0007548); 5 m$, 4 f£, Ponta Grossa, Parque Estadual de Vila Velha, S25.23078/2484, W49.9925/50.0298, 860‒900 m, 17‒18.ii.2016, *Moquiniastrum polymorphum* (D. Burckhardt & D.L. Queiroz) #198(12) (NHMB; 70% ethanol; NMB- PSYLL0007546); 1 m$, 4 f£, 4 immatures, same but S25.22384/24645, W49.99272/50.03994, 750‒870 m, 12‒14.vii.2017, *Moquiniastrum polymorphum* (D. Burckhardt & D.L. Queiroz) #246(14) (MMBC, NHMB; 70% ethanol; NMB- PSYLL0007549); 23 immatures, same but (L. Serbina) (MMBC; 70% ethanol); 1 f£, same but S25.2458, W50.0200, 968 m, 12.vii.2013, Cerrado vegetation (D.L. Queiroz) #539 (NHMB; 70% ethanol; NMB-PSYLL0007553); 1 f£, same but S25.3183, W49.1517, 970 m, 12.vii.2013, Cerrado vegetation (D.L. Queiroz) #540 (NHMB; 70% ethanol; NMB- PSYLL0007554); 1 immature, 1 skin, Tibagi, Parque Estadual Guartelá, S24.5623/5666, W50.2589, 920–950 m, 23–25. vi.2015, *Moquiniastrum polymorphum* (D. Burckhardt & D.L. Queiroz) #171(18) (NHMB; 70% ethanol; NMB-PSYLL0007545).—**Santa Catarina:** 15 m$, 18 f£, 19 immatures, same as holotype but (BMNH, MMBC, MHNG, NHMB, UFPR, USNM; dry, slide, 70% ethanol; NMB-PSYLL0007521, NMB-PSYLL0007540, NMB- PSYLL0007541, NMB-PSYLL0007542, NMB-PSYLL0007598, NMB-PSYLL0007543, NMB-PSYLL0007544, NMB-PSYLL0008285, NMB-PSYLL0007561, NMB- PSYLL0007562, NMB-PSYLL0007563, NMB-PSYLL0007564, NMB-PSYLL0007565, NMB-PSYLL0007566, NMB-PSYLL0007567 [LSMelmoq-1]); 2 m$, 4 f£, 1 immature, Urubici, S27.9416/9566, W49.5750/9560, 920–1140 m, 2.v.2013, *Moquiniastrum polymorphum* (D. Burckhardt & D.L. Queiroz) #121(4) (NHMB; 70% ethanol; NMB- PSYLL0007522).—**São Paulo:** 1 f£, Campinas, 28.xi.1966, ex water trap (C.L. Costa) #12/67 (BMNH; slide).

**Description. *Adult.*** Coloration. Body (Fig. 7B) pale to brownish yellow. Antenna pale yellow, segment 3 brownish at apex, segments 4–8 with dark brown apices, segments 9–10 entirely dark brown. Forewing (Fig. 14F) with pale yellow to brownish veins and clear membrane with distinct brownish scattered dots in the middle of cell c+sc and in apical two thirds of the wing; apices of pterostigma and veins Rs, M_1+2_, M_3+4_, Cu_1a_ and Cu_1b_ brown at wing margin. Legs pale yellow; apical tarsal segments greyish brown.

Structure. Setae on thorax long. Forewing (Fig. 14F) broadly and unevenly rounded apically; C+Sc evenly curved; pterostigma short and narrow, narrower in the middle than r_1_ cell; Rs almost straight, weakly curved to costal margin apically; M slightly longer than M_1+2_ and M_3+4_; Cu_1a_ curved in apical third, ending more or less at the same level with M fork; cell cu_1_ narrow; surface spinules present in all cells, leaving narrow spinule-free stripes along veins, forming indistinct hexagons of a single or double row of spinules, spinules absent from basal half of cell c+sc. Hindwing with 11 ungrouped costal setae. Metatibia bearing 5–8 apical spurs, arranged as 2–4 + 3+5, anteriorly separated by 5–7 bristles.

Terminalia (Fig. 22G–L). Male. Proctiger broad, posterior margin broadly convex; densely covered with long setae in apical two thirds. Subgenital plate irregularly ovoid, in lateral view; dorsal margin curved in proximal third, almost straight in apical two thirds, posterior margin slightly sinuate; with moderately long setae in apical half. Paramere distinctly shorter than proctiger; in lateral view, lamellar, with subparallel margins, except for apical fifth which is narrowing to apex; paramere apex, in lateral view, blunt, directed inwards and slightly caudad, in dorsal view, apex blunt, directed upwards and slightly inwards, lacking sclerotised tooth; outer face with sparse, moderately long setae mostly posteriorly; inner face with dense, moderately long setae; posterior margin with long setae. Proximal segment of aedeagus with apical part moderately subdivided. Distal segment of aedeagus almost straight in basal half with dorsal margin distinctly sinuate; apical dilation, in lateral view, oblong with subparallel margins, with lateral groove on each side forming lateral lobes, rounded apically; in dorsal view, apical dilation broad, with wing-like lateral lobes, obovate, broadest in apical third; sclerotised end tube, in lateral view, short and weakly curved. – Female terminalia cuneate, long; densely covered with setae. Dorsal margin of proctiger, in lateral view, regularly concave distal to circumanal ring, apex upturned, pointed; in dorsal view, apex pointed; circumanal ring, in dorsal view, cruciform. Subgenital plate shorter than proctiger; in lateral view, gradually narrowing towards apex, pointed apically; in ventral view, apex blunt.

***Fifth instar immature.*** Coloration. Pale yellow (Figs 38E, F); antenna pale yellow, segments 6–8 with brown apices, segment 9 entirely brown; base of wing pads and caudal plate brownish.

Structure. Eye with one long ocular seta dorsally. Antennal segments with following numbers of pointed sectasetae: 1(0), 2(1), 3(2), 4(0), 5(2), 6(1), 7(1), 8(0), 9(0); first segment with several long simple setae. Forewing pad with at least 20 marginal and ca. 40 dorsal lanceolate setae and pointed sectasetae densely spread over entire surface; hindwing pad with two marginal and ca. 10 dorsal pointed sectasetae and lanceolate setae. Metatibiotarsus, lacking pointed sectasetae; tarsal arolium broadly fan-shaped apically, about 1.2 times as long as claws. Abdomen with transverse rows of densely spaced pointed sectasetae and lanceolate setae on each sclerite dorsally. Caudal plate (Fig. 36J) with anterior margin close to anterior margin of extra pore fields; with many (ca. 75) pointed sectasetae and lanceolate setae arranged in several groups dorsally. Extra pore fields forming continuous outer and inner bands, consisting of several large oval patches; outer band long medially, end pointing outwards. Circumanal ring with anterior margin strongly concave medially (Fig. 5C).

**Host plant and biology.** *Moquiniastrum polymorphum* (Less.) G. Sancho (Asteraceae), including *M. polymorphum* subsp. *floccosum* (Cabrera) G. Sancho. Immatures develop on the adaxial surface of young leaves or abaxial surface of older leaves (Fig. 39D–F) where they secrete long, white filamentous wax threads. Immatures producing large amounts of honeydew were observed to be attended by two ant species, *Camponotus* sp. and *Crematogaster crinosa* (Mayr) (Hymenoptera, Formicidae) (I. Malenovský, pers. obs.).

**Distribution.** Brazil (MG, PR, SC, SP).

**Derivation of name.** Named after its host, *Moquiniastrum*.

### 7 Melanastera notia sp. nov

(Figs 8G, 14G, 22M–R, 36L)

**Type material. Holotype m$: Brazil: RIO GRANDE DO SUL:** 1 m$, Não-Me-Toque, S28.4666, W52.8027, 510 m, 8.iii.2013, *Vernonanthura phosphorica* (A.L. Marsaro Jr.) (UFPR; dry).

**Paratypes. Brazil: MINAS GERAIS:** 4 m$, 10 f£, 18 immatures, 5 skins, São Gonçalo do Rio Preto, Parque Estadual do Rio Preto, Vau das Éguas, S18.0995, W43.3301, 740 m, 15.iv.2021, indet. Asteraceae, Cerrado vegetation on rocky slope (D. Burckhardt & D.L. Queiroz) #407(4) (NHMB; 70% ethanol; NMB-PSYLL0008540, NMB-PSYLL0008541,

NMB-PSYLL0008542).—**Paraná:** 1 m$, Colombo, Embrapa campus, S25.3196/3353, W49.1562/1677, 920 m, 1–5.iv.2013, *Moquiniastrum polymorphum* (D. Burckhardt & D.L. Queiroz) #96(5) (NHMB; 70% ethanol; NMB-PSYLL0008539); 1 m$, Curitiba, Bosque Zaninelli, S253967, W492833, 950 m, 11.ii.2013, *Moquiniastrum polymorphum* (D. Burckhardt & D.L. Queiroz) #91(3) (NHMB; dry; NMB-PSYLL0007954); 1 m$, Ponta Grossa, Parque Estadual Vila Velha, S25.3146, W49.15197, 810 m, 12.vii.2013, Cerrado vegetation (D.L. Queiroz) #541 (NHMB; dry; NMB-PSYLL0007958); 1 m$, 1 immature, same but S25.2500, W500124, 830 m, 9.xii.2012, *Moquiniastrum polymorphum* (D. Burckhardt & D.L. Queiroz) #87(3) (NHMB; 70% ethanol; NMB-PSYLL0007953); 17 m$, 12 f£, 30 immatures, 7 skins, Tibagi, Parque Guartelá, S24.5638, W50.25586, 930 m, 8.xii.2013, Cerrado vegetation (D.L. Queiroz) #522 (NHMB; 70% ethanol; NMB- PSYLL0007551).—**Rio Grande do Sul:** 21 m$, 21 f£, 7 immatures, Não-Me-Toque, S28.4666, W52.8027, 510 m, 8.iii.2013, *Vernonanthura phosphorica* (A.L. Marsaro Jr.) (MMBC, NHMB, UFPR; dry, slide, 70% ethanol; NMB-PSYLL0007960, NMB- PSYLL0007995, NMB-PSYLL0008012, NMB-PSYLL0008013, NMB-PSYLL0008014

[LSMelver-60], NMB-PSYLL0008015 [LSMelver-60]); 2 m$, 4 f£, 6 immatures, Passo Fundo, Embrapa campus, S28.2288, W52.4065, 640 m, 20.ix.2018, *Moquiniastrum polymorphum* subsp. *ceanothifolium* (D. Burckhardt & D.L. Queiroz) #306(8) (NHMB; 70% ethanol; NMB-PSYLL0007956); 1 m$, 2 f£, 2 immatures, same but 27.vi.2016, degraded

vegetation (D.L. Queiroz) #517 (NHMB; 70% ethanol; NMB-PSYLL0007957); 1 f£, 1 immature, same but 28.vi.2013, degraded vegetation (D.L. Queiroz) #519 (NHMB; 70% ethanol; NMB-PSYLL0007959); 2 m$, 1 f£, Passo Fundo, Reserva Maragato, S28.2333, W52.4501, 640 m, 28.vi.2016 (A.L. Marsaro Jr.) 02/16PF (NHMB; dry; NMB- PSYLL0005203, NMB-PSYLL0007994); 4 m$, 10 f£, Passo Fundo, S28.2288, W52.4061, 670 m, 20.xi.2013 (A.L. Marsaro Jr.) (BMNH, MMBC, NHMB; dry, 70% ethanol; NMB- PSYLL0003912, NMB-PSYLL0007998); 2 m$, 8 f£, 5 immatures, same but 4.xii.2013 (A.L. Marsaro Jr.) (NHMB; 70% ethanol; NMB-PSYLL0003902, NMB-PSYLL0003900); 2 m$, 6 f£, 1 immature, same but 7.iv.2014 (A.L. Marsaro Jr.) (MHNG, NHMB; dry; NMB- PSYLL0007997, NMB-PSYLL0003890); 4 m$, 4 f£, 31 immatures, 8 skins, Santa Tecla, Tupanciretã, RS-377, S28.8777, W54.1274, 450 m, 19.ix.2018, *Moquiniastrum polymorphum* subsp. *ceanothifolium* (D. Burckhardt & D.L. Queiroz) #305(3) (NHMB; 70% ethanol; NMB- PSYLL0007955).—**São Paulo:** 1 f£, Matão, Fazendo Cambuhy, S21.6010, W48.4331, 550 m, 27.viii.2007 (P. Yamamoto) (FSCA; 70% ethanol).

**Description. *Adult.*** Coloration. Head (Fig. 8G) and clypeus dark brown. Antenna pale yellow, segments 4–9 with dark brown apices, segment 10 entirely dark brown. Thorax brown. Forewing (Fig. 14G) with pale yellow veins and clear membrane with distinct brown dots (in females more numerous than in males), spaced over entire wing, becoming denser towards wing apex; apices of veins Rs, M_1+2_, M_3+4_ and Cu_1a_ pale brown, apex of vein Cu_1b_ conspicuously brown. Legs pale brown; metacoxa dark brown with pale yellow meracanthus. Abdomen including terminalia brown. Younger specimens with less expanded dark colour.

Structure. Setae on thorax long. Forewing (Fig. 14G) narrowly, evenly rounded apically; C+Sc irregularly convex in median third; pterostigma moderately narrow, slightly narrower than cell r_1_, with; Rs almost straight in basal two thirds, weakly curved to costal margin apically; M much longer than M_1+2_ and M_3+4_; Cu_1a_ straight in basal two thirds, curved in apical third, ending distal to M fork; surface spinules present in all cells, relatively dense, leaving narrow to broad spinule-free stripes along the veins, forming indistinct hexagons, spinules absent from base of cell c+sc. Hindwing with grouped costal setae as 4 + 4–5.

Metatibia bearing 4–7 apical spurs, arranged as 2–3 + 2–4, anteriorly separated by 5–6 bristles.

Terminalia (Fig. 22M–R). Male. Proctiger broad, posterior margin convex; densely covered with long setae in apical two thirds. Subgenital plate irregularly ovoid, in lateral view; dorsal margin strongly curved in distal third, posterior margin almost straight; with moderately long setae in apical half. Paramere shorter than proctiger; in lateral view, lamellar, with subparallel margins, narrowing in apical fifth; paramere apex, in lateral view, blunt, directed slightly posteriad, in dorsal view, apex blunt, directed upwards and slightly inwards, lacking sclerotised tooth; outer face with sparse, short setae mostly in apical third; inner face with dense, moderately long setae submedially and short setae in apical half; anterior and posterior margins with longer setae. Proximal segment of aedeagus with apical part moderately subdivided. Distal segment of aedeagus almost straight in basal half with dorsal margin sinuate; apical dilation, in lateral view, broad, widest in the middle, with a lateral groove on each side forming lateral lobes, unevenly rounded apically; in dorsal view, apical dilation with wing-like lateral lobes, obovate, broadest in apical third, sclerotised end tube short and weakly curved. – Female terminalia cuneate, long; densely covered with setae.

Dorsal margin of proctiger, in lateral view, almost straight distal to circumanal ring, with a small bulge in the middle, concave subapically, apex pointed, upturned; in dorsal view, apex narrowly rounded; circumanal ring, in dorsal view, distinctly cruciform. Subgenital plate shorter than proctiger; in lateral view, abruptly narrowing towards apex in apical third, apex pointed; in ventral view, apex blunt.

***Fifth instar immature.*** Coloration. Brownish yellow; antenna gradually becoming darker towards apex; cephalothoracic sclerite, wing pads, legs and caudal plate brown.

Structure. Eye with one long, simple ocular seta dorsally. Antennal segments with following numbers of pointed sectasetae: 1(0), 2(1), 3(2), 4(0), 5(2), 6(1), 7(1), 8(0), 9(0). Forewing pad with 15–19 pointed sectasetae or lanceolate setae marginally and ca. 40 pointed lanceolate setae scattered over entire dorsal surface; hindwing pad with three marginal and one dorsal pointed sectasetae, and several (ca. 10) dorsal lanceolate setae. Metatibiotarsus with 2–3 pointed sectasetae on outer margin; tarsal arolium broadly fan-shaped apically, about

1.2 times as long as claws. Abdomen proximal to caudal plate with a transverse row of densely spaced lanceolate setae on dorsum of each segment, and several (5–6) pointed sectasetae on either side laterally. Caudal plate (Fig. 36L) with anterior margin close to anterior margin of extra pore fields; with at least seven pointed sectasetae arranged in two groups (3 + 4 or 4 + 3) on either side laterally, a number of lanceolate setae and few pointed sectasetae scattered over entire surface dorsally, and with 6–7 pointed sectasetae and several lanceolate setae on either side of circumanal ring dorsally. Extra pore fields forming continuous outer and inner bands, consisting of large oval patches; outer band long medially, end pointing outwards. Circumanal ring medially strongly indented anteriorly.

**Host plants.** *Moquiniastrum polymorphum* (Less.) G. Sancho, including *M. polymorphum* subsp. *ceanothifolium* (Less.) G. Sancho, *Vernonanthura phosphorica* (Vell.) H.Rob. (Asteraceae).

**Distribution.** Brazil (MG, PR, RS, SP).

**Derivation of name.** From the Ancient Greek adjective νοτιος = southern, referring to its southern distribution.

## The *olgae*-group

**Description. *Adult*.** Head, in lateral view, inclined at 45° or more from longitudinal body axis (Fig. 7C). Vertex trapezoidal, with faint to strong imbricate microsculpture and microscopic setae; median suture fully developed (Figs 8H–J, 9A–C) except for *M. tijuca* (Fig. 9D) where it is reduced in basal three quarters of vertex. VW/VL 1.6–2.4. AL 0.5–0.9; AL/HW 0.9–1.8. Thorax moderately arched, with microscopic setae. Forewing pale yellow to dark brown, lacking small brown dots; wing apex in cell r_2_ near M_1+2_ or in the middle of cell margin; pterostigma expanding towards the middle or apical third. Distal segment of aedeagus bearing ventral process.

***Immature*.** Antenna 10-segmented.

**Comments.** The *olgae-*group includes seven species in Brazil. In addition, *Melanastera granulosi* sp. nov. resembles *M. macaireae* sp. nov. and *M. tijuca* sp. nov. in the narrow, yellow forewing with light stripes along the veins, but is not placed in the *olgae-*group due to a lack of male material (see comments under *M. granulosi*).

Brown & Hodkinson (1988) figured a species, *Haplaphalara* sp. A from Panama, which resembles the male of *M. variegata* sp. nov. It differs from the latter species in having a less extensive forewing pattern and less curved veins Rs and Cu_1a_.

*Melanastera lucens* (Burckhardt, Hanson & Madrigal) from Costa Rica resembles the *olgae-*group by the absence of dark dots on the forewings. It differs from the Brazilian species by a smaller size, a bright orange body and details of the male and female terminalia (Burckhardt *et al*. 2005). See also the comment under *duckei*-group.

The species of the *olgae-*group are associated with Annonaceae and Melastomataceae. Its members show great diversity in forewing shape, venation and colour pattern, as well as in the structure of the paramere and aedeagus. The group is probably polyphyletic.

### 8 Melanastera olgae sp. nov

(Figs 7C, 8H, 14H, 22S–X, 40A, B)

**Type material. Holotype m$: Brazil: MATO GROSSO:** 1 m$, Sinop, Embrapa campus, S22.0085, W55.5828, 360 m, 30.viii.2013 (D.L. Queiroz) #571 (UFPR; dry).

**Paratypes. Brazil: MATO GROSSO:** 2 f£, Lucas do Rio Verde, S13.2490, W56.0275, 400 m, 12.6.2016 (L.A. Pezzini) #794 (NHMB; 70% ethanol; NMB-PSYLL0005792); 31 m$, 39 f£, 40 immatures, 9 skins, same as holotype but (BMNH, MHNG, MMBC, NHMB, UFPR, USNM; dry, slide, 70% ethanol; NMB-PSYLL0007703, NMB-PSYLL0007717 [LSMelolg- 16], NMB-PSYLL0007718 [LSMelolg-16], NMB-PSYLL0007720, NMB-PSYLL0007721, NMB-PSYLL0007722, NMB-PSYLL0007723, NMB-PSYLL0007654, NMB- PSYLL0007668, NMB-PSYLL0007669, NMB-PSYLL0007670, NMB-PSYLL0007671, NMB-PSYLL0007672); 1 m$, Sinop, S11.7477, W55.3668, 360 m, 5.iii.2014 (L.A. Pezzini & A.D.B. Sedano) #288(NHMB; 70% ethanol; NMB-PSYLL0005587); 2 m$, 2 f£, same but S11.8662, W55.5186, 360 m, 25.viii.2012 (D.L. Queiroz) #336 (NHMB; 70% ethanol; NMB-PSYLL0007715); 32 m$, 50 f£, 20 immatures, 3 skins, same but Embrapa campus, S11.8675, W55.6526, 350 m, 28.viii.2013 (D.L. Queiroz) #566 (MMBC, NHMB; 70% ethanol; NMB- PSYLL0007712); 5 m$, 4 f£, 1 skin, same but S22.0071, W55.5776, 360 m, 30.viii.2013 (D.L. Queiroz) #572 (NHMB; 70% ethanol; NMB-PSYLL0007714); 2 m$, 4 f£, 3 immatures, same but S25.2346, W49.1059, 355 m, 27.ii.2015, *Guatteria punctata* (D.L. Queiroz) #680(5) (NHMB; 70% ethanol; NMB-PSYLL0007716); 1 f£, Sorriso, Fazenda Mazzardo II, S11.8714, W55.5957, 390 m, 26.ii.2015 (D.L. Queiroz) #678(3) (NHMB; 70% ethanol; NMB-PSYLL0007713); 2 m$, same but S12.4276, W55.7959, 350 m, 26.vii.2014 (T. Mazzardo) #P013 (NHMB; dry; NMB-PSYLL00003016); 1 m$, 1 f£, same but S12.4233, W55.8502, 400 m, 26.vii.2014 (T. Mazzardo) #P017 (NHMB; dry; NMB-PSYLL00003029); 1 f£, same but S12.4253, W55.8505, 390 m, 26.vii.2014 (T. Mazzardo) #P021 (NHMB; dry; NMB-PSYLL00003014); 1 f£, same but S12.4222, W55.8500, 390 m, 15.viii.2013 (T. Mazzaro) (NHMB; 70% ethanol; NMB-PSYLL0007702).

**Description. *Adult.*** Coloration. Body (Figs 7C, 39B) dark brown. Antenna whitish, scape and pedicel and base of segment 3 brown, segments 3–8 with dark brown apices, segment 10 entirely dark brown. Forewing (Fig. 14H) dark brown, membrane sometimes slightly paler in the middle of cells; claval suture light. Metacoxa brown, metafemur brown in basal half, yellowish in apical half; metatibia and metatarsus yellowish. Younger specimens with less expanded and intense dark colour.

Structure. Forewing (Fig. 14H) semicoriaceous, strongly convex; oviform, widest in the middle, broadly, unevenly rounded apically; C+Sc curved in distal third; pterostigma long, slightly wider than cell r_1_, evenly convex; Rs moderately and evenly curved; M straight, slightly shorter than M_1+2_, slightly longer than M_3+4_; Cu_1a_ weakly and evenly curved, ending distad of M fork; cell cu_1_ short and narrowly triangular; surface spinules indistinct. Hindwing with 10–13 ungrouped costal setae. Metatibia bearing 7–8 grouped apical spurs, arranged as 3–4 + 4, anteriorly separated by 3–4 bristles.

Terminalia (Fig. 22S–X). Male. Proctiger weakly produced posteriorly; densely covered with long setae in apical two thirds. Subgenital plate large, subtrapezoidal; dorsal margin mostly straight, downcurved in distal third, posterior margin slightly concave; with moderately long setae. Paramere shorter than proctiger; in lateral view, lamellar, anterior margin straight, slightly indented subapically, posterior margin bearing small lobe in basal half and evenly curved in apical fifth; paramere apex, in lateral view, broad, blunt, directed slightly anteriad, in dorsal view, apex long, narrow, subacute, directed inwards, laking a sclerotised tooth; both outer and inner faces with dense moderately long setae; posterior margin with long setae. Proximal segment of aedeagus with apical part strongly subdivided. Distal segment of aedeagus distinctly sinuate in basal half; ventral process situated slightly proximad of the middle of segment, shovel-like, in lateral view, with a narrow base, broadest at apical third, with narrow, claw-like apex strongly curved dorsad; in dorsal view, ventral process much broader than apical dilation and subtrapezoidal basally, with long, narrow, apically truncate median lobe arising from the base ventrally; apical dilation, in lateral view, long, narrow, with subparallel margins, slightly inflated apically, with narrowly rounded apex, base bearing a small membranous extension at dorsal margin, in dorsal view, apical dilation distinctly constricted medially; sclerotised end tube long, hardly sinuate. – Female terminalia cuneate, long; densely covered with setae. Dorsal margin of proctiger, in lateral view, weakly concave distal to circumanal ring, apex relatively straight, pointed; in dorsal view, apex narrowly rounded; circumanal ring, in dorsal view, distinctly cruciform. Subgenital plate slightly shorter than proctiger; in lateral view, pointed apically; in ventral view, apex narrowly rounded.

***Fifth instar immature.*** Coloration. Light; antenna gradually becoming darker towards apex; cephalothoracic sclerite, wing pads, legs and caudal plate pale brown.

Structure. Eye with one short, simple ocular seta dorsally. Antennal segments with following numbers of pointed sectasetae: 1(0), 2(1), 3(0), 4(2), 5(0), 6(2), 7(1), 8(1), 9(0), 10(0). Forewing pad with five marginal pointed sectasetae; hindwing pad with two marginal pointed sectasetae; both pads lacking sectasetae dorsally. Metatibiotarsus very long; tarsal arolium broadly fan-shaped apically, 1.5–2.0 times as long as claws. Abdomen with three lateral pointed sectasetae on either side anterior to caudal plate. Caudal plate with anterior margin in distance of anterior margin of extra pore fields; on either side laterally, with 2+2 pointed sectasetae, in some specimens with one moderately large lanceolate seta associated with the posterior pair, and three pointed sectasetae subapically on either side of circumanal ring dorsally. Extra pore fields forming continuous outer and inner bands, consisting of small rounded patches; outer band relatively short medially, end pointing backwards. Circumanal ring small.

**Host plant and biology.** *Guatteria punctata* (Aubl.) R.A.Howard (Annonaceae).

Immatures secrete long wax filaments. Adults were observed on the flowers of its host plant (Fig. 40A, B) (D.L. Queiroz, pers. observation).

**Distribution.** Brazil (MT).

**Derivation of name.** Dedicated to Olga Yarmolik, mother of LS, for her support and unconditional love.

**Comments.** *Melanastera olgae* sp. nov. resembles the species of the *amazonica*-group (*M. amazonica* sp. nov., *M. cacantis* sp. nov., *M. francisi* sp. nov., *M. roraima* sp. nov., *M. tubuligera* sp. nov. and *M. xylopiae* sp. nov.) in the broadly, unevenly rounded apex of the forewing with a broad pterostigma (PtL/PtW < 4.0) and convex R_1_, as well as details of the distal segment of aedeagus: ventral process bearing a distinct apico-median lobe, which is, in lateral view, claw-like and directed dorsad, and, in dorsal view, with narrow, truncate lobe; long apical dilation (short in *M. amazonica*), and long sclerotised end tube (short in *M. amazonica*). *Melanastera olgae* differs from these species in the relatively uniform dark body and forewing colour (versus variegated), the semicoriaceous, strongly convex forewing (versus semitransparent, flat); the indistinct microsculpture on the vertex (versus distinct); the long antenna (AL > 0.9 versus < 0.8); the long metatibia (MT/HW > 0.8 versus < 0.8).

### 9 Melanastera mazzardoae sp. nov

(Figs 8I, 14I, J, 23A–D)

**Type material. Holotype m$: Brazil: MATO GROSSO:** Sorriso, Fazenda Mazzardo II, S12.4217, W55.8432, 370 m, 15.viii.2013 (T. Mazzardo) (UFPR; slide; [LSMelmazz-23]).

**Paratype. Brazil: MATO GROSSO:** 1 f£, same as holotype but (NHMB; slide; NMB- PSYLL0007929 [LSMelmazz-23]); 1 m$, same but S12.4536, W55.8071, 370 m, 25.vii.2014 (T. Mazzardo) #P008 (NHMB; dry; NMB-PSYLL00003023).

**Description. *Adult.*** Coloration. Body orange. Antenna pale yellow, segments 6 and 8 with brown apices, segments 9–10 entirely brown. Forewing (Fig. 14I, J) yellow, in male with colourless areas: a patch at base, a band at nodal line, and a circular area subapically at costal margin. Legs pale yellow to orange, meracanthus of metacoxa pale yellow.

Structure. Forewing (Fig. 14I, J) oval, widest in the middle, broadly, evenly rounded apically; C+Sc almost straight in basal two thirds, weakly curved in apical third; pterostigma slightly narrower than r_1_ cell in the middle; Rs moderately convex; M much longer than M_1+2_ and M_3+4_; Cu_1a_ unevenly curved, ending slightly proximad of M fork; cell cu_1_ short and wide; surface spinules present in all cells, leaving very narrow spinule-free stripes along veins, forming hexagons of a single row of spinules, spinules absent from basal half of cell c+sc. Hindwing with 5–7 ungrouped costal setae. Metatibia bearing 7 apical spurs, arranged as 3 + 4, anteriorly separated by 2 or more bristles.

Terminalia (Fig. 23A–D). Male. Proctiger strongly produced in basal half; densely covered with long setae in apical two thirds. Subgenital plate irregularly ovoid; dorsal margin hardly curved, posterior margin evenly convex; with sparse, moderately long setae. Paramere, in lateral view, sickle-shaped: broadest basally, gradually narrowing to apex; paramere apex, in lateral view, narrow, directed anteriad, with a small tooth; outer face with short setae in apical two thirds; inner face with moderately long setae; posterior margin with long setae. Proximal segment of aedeagus with apical part strongly subdivided. Distal segment of aedeagus with strongly sinuate dorsal margin; ventral process situated slightly distad of the middle of the segment, in lateral view, narrow, simple, tongue-shaped, lacking lateral lobes; apical dilation elongate, narrow, with weakly convex ventral margin and narrowly, unevenly rounded apex; sclerotised end tube short and almost straight. – Female terminalia cuneate; covered with setae. Dorsal margin of proctiger, in lateral view, slightly convex posterior to circumanal ring, almost straight in apical third, apex subacute, in dorsal view, apex blunt; circumanal ring, in dorsal view, distinctly cruciform. Subgenital plate, in lateral view, unevenly narrowing to apex, apex pointed, slightly upturned; in ventral view, apex broadly blunt.

*Fifth instar immature.* Unknown.

**Host plant.** Unknown.

**Distribution.** Brazil (MT).

**Derivation of name.** Dedicated to Tatiana Mazzardo, the collector of the species.

**Comments.** *Melanastera mazzardoae* sp. nov. resembles *M. acuminata* sp. nov., *M. duckei* sp. nov. and *M. variegata* sp. nov. in the forewing shape and venation; the more or less sickle-shaped paramere and the distal segment of the aedeagus with a narrow, simple, tongue-shaped ventral process lacking lateral lobes. It differs from *M. acuminata* sp. nov. and *M. duckei* sp. nov. in the sexually dimorphic forewing, lacking brown dots (versus not dimorphic, bearing brown dots), with broader pterostigma (PtL/PtW < 4.0 versus > 4.4) and broader cell cu_1_ (Cu_1a_L/cu_1_W < 2.7 versus > 3.2). It differs from *M. variegata* as indicated in the key and in DNA barcoding sequences: the uncorrected p-distance between *M. mazzardoae* and *M. variegata* is 17.9% for *COI* and 24.7% for *cytb*.

### 10 Melanastera variegata sp. nov

(Figs 8J, 15A, B, 23E–J)

**Type material. Holotype m$: Brazil: AMAZONAS:** Novo Airão, Parque Nacional de Anavilhanas, S2.5366/6217, W60.8327/9559, 20 m, 16.iv.2014, *Guatteria* sp. (D. Burckhardt & D.L. Queiroz) #128(1) (UFPR; dry).

**Paratypes. Brazil: AMAZONAS:** 2 f£, Manaus, Sede Embrapa, Campo Experimental, AM- 010, km 29, S2.8950, W59.9733, 100 m, 27–30.iv.2014 (D. Burckhardt & D.L. Queiroz) #133 (NHMB; dry, slide; NMB-PSYLL0008139, NMB-PSYLL0008171 [LSMelvar-56A]); 1 m$, 13 immatures, same as holotype but (NHMB; slide, 70% ethanol; NMB-PSYLL0008135, NMB-PSYLL0008167 [LSMelvar-56A], NMB-PSYLL0008165, NMB-PSYLL0008166); 1 m$, 1 f£, 20 immatures, same but *Guatteria* sp. (D. Burckhardt & D.L. Queiroz) #128(3) (NHMB; dry, slide, 70% ethanol; NMB-PSYLL0008135, NMB-PSYLL0008167 [LSMelvar- 56A], NMB-PSYLL0008165, NMB-PSYLL0008166, NMB-PSYLL0008136, NMB- PSYLL0008137, NMB-PSYLL0008168 [LSMelvar-56A], NMB-PSYLL0008169, NMB-PSYLL0008170 [LSMelvar-56A]); 1 f£, same but along road from Novo Airão to Manacapuru, S2.7033/3.1366, W60.7033/7316, 60 m, 22.iv.2014 (D. Burckhardt & D.L. Queiroz) #130 (NHMB; 70% ethanol; [LSMelvar-56A]).—**Roraima:** 1 f£, Mucajaí, km 442–440 along BR-174, S Mucujaí, N2.3583/3850, W60.9116/9166, 70–80 m, 18–19.iv.2015

Burckhardt & D.L. Queiroz) #165 (NHMB; 70% ethanol; NMB-PSYLL0008316).

**Description. *Adult.*** Coloration. Body dark brown. Genae and clypeus pale brown. Antenna pale yellow, segments 4–8 with dark brown apices, segments 9–10 entirely dark brown. Forewing dark brown, sexually dimorphic: in male with three large colourless areas: a patch at base, a wide transverse band along nodal line and an oval subapical patch at costal wing margin (Fig. 15A); in female with slightly lighter basal third and nodal line and a colourless small patch at the apex of clavus (Fig. 15B). Femora, tibiae and tarsi pale brown. Younger specimens with less extended dark colour.

Structure. Forewing (Fig. 15A, B) oval, widest in the middle, broadly and evenly rounded apically; C+Sc almost straight except for apical fifth which is curved; pterostigma short in males, long in females, in the middle about as wide as cell r_1_, convex; Rs strongly convex; M longer than M_1+2_ and M_3+4_; Cu_1a_ unevenly curved, ending proximal of M fork; cell cu_1_ short and wide; surface spinules present in all cells, leaving narrow spinule-free stripes along the veins, spinules absent in basal half of cell c+sc, forming hexagons of a single row of spinules. Hindwing with 4–5 + 3–4 grouped costal setae. Metatibia bearing 7 apical spurs, arranged as 3 + 4, anteriorly separated by 3–4 bristles.

Terminalia (Fig. 23E–J). Male. Proctiger relatively narrow; densely covered with long setae in apical two thirds. Subgenital plate subglobular, in lateral view, with dorsal margin straight, rounded down caudally, posterior margin convex; with sparse, long setae, mostly in apical half. Paramere, in lateral view, weakly falcate; paramere apex, in lateral view, subacute, directed anteriad, in dorsal view, apex digitiform, narrow, with a small sclerotised tooth; both outer and inner faces with relatively sparse, moderately long setae; posterior margin with long setae. Proximal segment of aedeagus with apical part strongly subdivided. Distal segment of aedeagus with dorsal margin strongly sinuate; ventral process situated in the middle of the segment, in lateral view, elongate, relatively short, narrow, simple; in dorsal view, ventral process short, wider than apical dilation, widest in apical third, indistinctly three-lobed: median lobe longest; apical dilation, in lateral view, narrow, slightly widening to unevenly rounded apex, in dorsal view, apical dilation narrow, broadest medially, broadly rounded apically; sclerotised end tube moderately long and almost straight. – Female terminalia cuneate; densely covered with setae. Dorsal margin of proctiger, in lateral view, almost straight, with a small hump distal to circumanal ring, apex subacute; in dorsal view, apex blunt; circumanal ring, in dorsal view, distinctly cruciform. Subgenital plate in lateral view, with pointed apex; in ventral view, blunt apex.

***Fifth instar immature.*** Coloration. Yellow; antenna gradually darkening to apex; cephalothoracic sclerite, wing pads, legs and caudal plate brown.

Structure. Eye with one short, simple ocular seta dorsally. Antennal segments with following numbers of pointed sectasetae: 1(0), 2(1), 3(0), 4(2), 5(0), 6(2), 7(1), 8(1), 9(0), 10(0). Forewing pad with five marginal pointed sectasetae; hindwing pad with two marginal pointed sectasetae; both pads lacking sectasetae or larger lanceolate setae dorsally. Metatibiotarsus long; tarsal arolium broadly fan-shaped apically, 1.5 times as long as claws. Abdomen with 2–3 lateral pointed sectasetae on either side anterior to caudal plate. Caudal plate with anterior margin relatively close to anterior margin of extra pore fields; with 2+2 pointed sectasetae on either side laterally, and three pointed sectasetae subapically, on either side of circumanal ring dorsally. Extra pore fields forming continuous outer and inner bands, consisting of small oval patches; outer band long medially, end pointing outwards. Circumanal ring small.

**Host plant.** *Guatteria* sp. (Annonaceae).

**Distribution.** Brazil (AM, RR).

**Derivation of name.** From the Latin adjective *variegatus* = marked with a variety of colours, referring to the patchy forewing pattern of the male.

**Comments.** See comments under *M. mazzardoae* sp. nov.

### 11 Melanastera parva sp. nov

(Figs 9A, 15C, 23K–P)

**Type material. Holotype m$: Brazil: MATO GROSSO:** Sinop, S11.8713, W55.3970, 380 m, 4.iii.2014, *Xylopia* sp. (L.A. Pezzini & A.D.B. Sedano) #281 (UFPR; dry).

**Paratypes. Brazil: MATO GROSSO:** 6 m$, 8 f£, 1 skin, same as holotype but (MMBC, NHMB, UFPR; dry, slide, 70% ethanol; NMB-PSYLL0005582, NMB-PSYLL0005665, NMB-PSYLL0005672, NMB-PSYLL0005673); 6 m$, 7 f£, same but S11.8510, W55.3694, 380 m, 4.iii.2014, *Xylopia* sp. (L.A. Pezzini & A.D.B. Sedano) #272 (NHMB; 70% ethanol; NMB-PSYLL0007968); 6 m$, 11 f£, same but S55.3975, W11.8696, 380 m, 4.iii.2014, *Xylopia* sp. (L.A. Pezzini & A.D.B. Sedano) #283 (NHMB, MMBC; slide, 70% ethanol; NMB-PSYLL0005583, NMB-PSYLL0008053); 4 m$, 2 f£, 7 immatures, 9 skins, same but S11.9328, W55.48970, 390 m, 27.viii.2012 (D.L. Queiroz) #343 (NHMB; slide, 70% ethanol; NMB-PSYLL0008040, NMB-PSYLL0008049 [LSMelpar-22-1], NMB-PSYLL0008050 [LSMelpar-22], NMB-PSYLL0008051 [LSMelpar-22], NMB-PSYLL0008052).

**Description. *Adult.*** Coloration. Body pale yellow to orange. Antenna yellow to orange, apices of segments 6 and 8 brown, segments 9–10 entirely dark brown. Mesopraescutum with two pale orange patches along fore margin; mesoscutum with five longitudinal pale orange stripes, the median one narrower than the remainder. Forewing (Fig. 15C) evenly pale to bright yellow; apex of Cu_1b_ brown. Legs pale yellow to orange, meracanthus pale yellow.

Structure. Forewing (Fig. 15C) oblong-oval, widest in apical third, broadly, evenly rounded apically; C+Sc weakly curved in apical third; pterostigma long, slightly narrower in the middle than cell r_1_, weakly convex; Rs weakly and evenly convex; M longer than M_1+2_ and M_3+4_; Cu_1a_ weakly and irregularly convex, ending slightly proximad of M fork; cell cu_1_ short and relatively wide; surface spinules present in all cells, leaving narrow spinule-free stripes along the veins, absent from basal half of cell c+sc, forming distinct hexagons of a single row of spinules. Hindwing with 6 ungrouped costal setae. Metatibia bearing 4–5 apical spurs, arrangd as 2 + 2–3, anteriorly separated by 3–4 bristles.

Terminalia (Fig. 23K–P). Male. Proctiger with posterior lobe; densely covered with long setae in apical two thirds. Subgenital plate, in lateral view, irregularly ovoid; dorsal margin curved, posterior margin almost straight; with few moderately long setae. Paramere shorter than proctiger, in lateral view, narrowly lanceolate; paramere apex, in lateral view, blunt, in dorsal view, rounded, directed inwards and slightly upwards, lacking distinct sclerotised tooth; outer and inner face with relatively dense, moderately long setae; posterior margin with long setae. Proximal segment of aedeagus with apical part moderately subdivided. Distal segment of aedeagus, in lateral view, weakly sinuate in basal half, in dorsal view, very broad; ventral process situated slightly distad of the middle of segment, in lateral view, broad, slightly widening to apex which is broadly truncate except for a small point produced, with small lateral lobes; in dorsal view, ventral process broadly ovoid, broadest in apical third, much broader than diameter of apical dilation, with a short, median apical point; apical dilation relatively small, in lateral view, narrow, with subparallel margins and rounded apex, in dorsal view, elongate, with subparallel margins and rounded apex; sclerotised end tube short and weakly sinuate. – Female terminalia cuneate; densely covered with setae. Dorsal margin of proctiger, in lateral view, weakly convex distal to circumanal ring, almost straight in apical third, apex blunt; in dorsal view, apex subacute; circumanal ring, in dorsal view, vaguely cruciform. Subgenital plate, in lateral view, abruptly narrowing in apical third, pointed apically; in ventral view, apex blunt.

***Fifth instar immature.*** Coloration. Brownish yellow; antenna with dark apical segments; cephalothoracic sclerite, wing pads, legs and caudal plate brownish yellow to pale brown.

Structure. Eye with one simple ocular seta dorsally. Antenna lacking pointed sectasetae but segments 1, 2, 3, 5, 6, 7 and 8 bearing simple setae, and segment 4 with one relatively large slender lanceolate seta. Forewing pad with 8–11 marginal and 3–4 dorsal pointed sectasetae; hindwing pad with 3 marginal and 2 dorsal pointed sectasetae. Metatibiotarsus short; tarsal arolium broadly fan-shaped apically, at least 1.5 times as long as claws. Abdominal tergites anterior to caudal plate with three dorsal rows of 6, 2 and 4 pointed sectasetae, respectively. Caudal plate dorsally with one row of four pointed sectasetae along anterior margin, two rows each of four pointed sectasetae submedially, and with two groups of four pointed sectasetae on each side subapically, anterior to circumanal ring; with a few long simple setae on either side ventrally. Extra pore fields forming continuous outer and inner band, consisting of small rounded patches; outer band very short, end pointing outwards. Circumanal ring large.

**Host plant.** Adults and a single damaged immature were collected on *Xylopia* sp. (Annonaceae), which is a likely host. The description of the immature is based on the sample DLQ#343 lacking a host identification.

**Distribution.** Brazil (MT).

**Derivation of name.** From the Latin adjective *parvus* = little, small, referring to the small size of the species.

**Comments.** *Melanastera parva* sp. nov. resembles *M. barretoi* sp. nov. and *M. sericeae* sp. nov. in the narrowly lanceolate paramere with blunt apex, the broad ventral process of the distal segment of the aedeagus with an apico-median point or lobe, and FP/HW > 0.8. It differs from the two species in the absence of brown dots on the forewing (versus presence) and the ventral process of the distal segment of the aedeagus with an apico-median point (versus lobe). *Melanastera parva* also resembles *M. macaireae* sp. nov. in the small body size (FL < 1.5 mm), the light body colour and yellow forewing lacking brown dots, the narrowly lanceolate paramere with blunt apex; the broad ventral process of the distal segment of the aedeagus and the relatively straight dorsal margin of the female proctiger. It differs from *M. macaireae* as indicated in the key. Finally, *M. parva* resembles *M. granulosi* sp. nov. in the pale yellow forewing with surface spinules forming hexagons of a single row of spinules. It differs from *M. granulosi* in the shorter metatibia (MT/HW < 0.9 versus > 1.0), the shorter female proctiger (FP/HW < 0.9 versus > 1.1) and the shorter female subgenital plate (FP/SP > 1.8 versus < 1.4).

### 12 Melanastera guatteriae sp. nov

(Figs 9B, 15D, 23Q–V)

**Type material. Holotype m$: Brazi**l: CEARÁ: Ubajara, Parque Nacional de Ubajara, Planalto, S3.8433, W40.9100, 840 m, 5‒6.vii.2016, *Guatteria sellowiana* (D. Burckhardt & D.L. Queiroz) #218(3) (UFPR; dry).

**Paratypes. Brazil: CEARÁ:** 4 m$, 12 f£, 2 immatures, same as holotype but (NHMB; dry, slide, 70% ethanol; NMB-PSYLL0005919, NMB-PSYLL0007979, NMB-PSYLL0005920, NMB-PSYLL0007917, NMB-PSYLL0007918, NMB-PSYLL0007919, NMB- PSYLL0007920 [LSMelguat-17], NMB-PSYLL0007921 [LSMelguat-17]); 1 immature, Ubajara, Planalto, S3.8433, W40.9100, 840 m, 5‒6.vii.2016, *Guatteria sellowiana* (D. Burckhardt & D.L. Queiroz) #219(3) (NHMB; slide; NMB-PSYLL0007922); 1 m$, same but S3.8368/8371, W40.8980/9096, 780‒820 m, 1‒7.vii.2016 (D. Burckhardt & D.L. Queiroz) #211 (NHMB; dry; NMB-PSYLL0005916).

**Description. *Adult.*** Coloration. Head (Fig. 9B) and clypeus bright yellow to orange; vertex dark brown posteriorly. Antenna pale yellow, segments 3–8 with brown apices, segments 9–10 entirely brown. Pronotum dark brown, distinctly darker than vertex; mesopraescutum brown, often darker anteriorly; mesoscutum pale brown, brown posteriorly; mesoscutellum dark brown along margins, lighter brown in the middle. Metanotum dark brown. Forewing (Fig. 15D) yellow, pale brown at apex and in cell a_2_. Legs yellow to orange, metafemur dark brown at base. Abdomen yellow to orange, with dark brown sclerites; in female the first two tergites dark brown. Terminalia dark brown. Young specimens entirely pale yellow.

Structure. Forewing (Fig. 15D) oviform, widest in the middle, broadly rounded apically; C+Sc weakly curved in distal third; pterostigma slightly narrower than r_1_ cell, weakly convex in apical third; Rs straight in basal three quarters, weakly curved to fore margin apically; M longer than M_1+2_ and M_3+4_; Cu_1a_ mostly weakly curved to anal margin apically, ending at M fork; cell cu_1_ narrow; surface spinules present in all cells, forming distinct hexagons of a single row of spinules, leaving narrow spinule-free stripes along the veins, spinules becoming sparse in basal half of the wing and completely absent in basal half of cell c+sc. Hindwing with 3–5 + 4 grouped costal setae. Metatibia bearing 6–8 apical spurs, arranged as 3 + 3–5, anteriorly separated by ca. 5 bristles.

Terminalia (Fig. 23Q–V). Male. Proctiger moderately produced posteriorly; densely covered with moderately long setae in apical two thirds. Subgenital plate subtriangular, in lateral view, dorsal margin sinuate, posterior margin relatively straight; with moderately long setae. Paramere, in lateral view, narrowly lanceolate; paramere apex, in lateral view, blunt, directed upwards, in dorsal view, apex blunt, directed upwards and slightly inwards, lacking distinct sclerotised tooth; both outer and inner faces with dense, moderately long setae; posterior margin with long setae. Proximal segment of aedeagus with apical part weakly subdivided. Distal segment of aedeagus straight in basal two thirds, with dorsal margin weakly sinuate; ventral process inserted in apical third of the segment, in lateral view, simple, broadly rounded apically, with dorsal and ventral margins subparallel, lacking lateral lobes; in dorsal view, ventral process broad, gradually widening towards apex, broadest subapically; apical dilation, in lateral view, moderately broad, rounded apically, in dorsal view, apical dilation narrowly oval, broadest medially, rounded apically; sclerotised end tube relatively short and weakly curved. – Female terminalia cuneate; densely covered with setae. Dorsal margin of proctiger, in lateral view, almost straight posterior to circumanal ring, weakly concave subapically, apex pointed, slightly turned upwards; in dorsal view, apex blunt; circumanal ring, in dorsal view, cruciform. Subgenital plate, in lateral view, abruptly narrowing in apical half, pointed apically; in ventral view, apex blunt.

***Fifth instar immature.*** Coloration. Light; cephalothoracic sclerite, antenna, wing pads, legs and caudal plate pale brown.

Structure. Eye with one short, simple ocular seta dorsally. Antennal segments with following numbers of pointed sectasetae: 1(0), 2(1), 3(0), 4(2), 5(0), 6(2), 7(1), 8(1), 9(0),

10(0). Forewing pad with 4–5 marginal pointed sectasetae; hindwing pad with two marginal pointed sectasetae; both pads lacking sectasetae or larger lanceolate setae dorsally. Metatibiotarsus long; tarsal arolium broadly triangular apically, 1.2–1.5 times as long as claws. Abdomen with three lateral pointed sectasetae on either side anterior to caudal plate. Caudal plate with anterior margin close to anterior margin of extra pore fields; with 2+2 or 1+2 pointed sectasetae on either side laterally, and three pointed sectasetae subapically, on either side of circumanal ring dorsally. Extra pore fields forming continuous outer and inner bands, consisting of small oval patches; outer band long medially, end pointing outwards. Circumanal ring small.

**Host plant.** *Guatteria sellowiana* Schltdl. (Annonaceae).

**Distribution.** Brazil (CE).

**Derivation of name.** Named after its host, *Guatteria*.

**Comments.** *Melanastera guatteriae* sp. nov. shares with *M. australis* sp. nov., *M. dimorpha* sp. nov. and *M. obscura* sp. nov. similar male terminalia, i.e. the narrowly lanceolate paramere and the simple ventral process which is inserted in apical third of the distal segment of the aedeagus. It differs from these species in the absence of brown dots on the forewing.

### 13 Melanastera macaireae sp. nov

(Figs 9C, 15E, 24A–G)

**Type material. Holotype m$: Brazil: MINAS GERAIS:** Buenópolis, Parque Estadual da Serra do Cabral, Vereda do Jacaré, S17.9212, W44.2372, 1000 m, 20.ix.2019, *Macairea radula* (D. Burckhardt & D.L. Queiroz) #370(1) (UFPR; dry).

**Paratypes. Brazil: MINAS GERAIS:** 9 m$, 11 f£, 18 immatures, 1 skin, same as holotype but (BMNH, MMBC, NHMB, UFPR; dry, 70% ethanol; NMB-PSYLL0007946, NMB- PSYLL0007980, NMB-PSYLL0007981, NMB-PSYLL0007982); 1 f£, same but S17.8853, W44.2636, 1000 m, 8.iv.2021 (D. Burckhardt & D.L. Queiroz) #384(0) (NHMB; 70% ethanol; NMB-PSYLL0008560); 1 m$, 2 f£, same as holotype but S17.9212, W44.2371, 1030 m, 9.iv.2021, riparian vegetation, *Miconia* sp. (D. Burckhardt & D.L. Queiroz) #393(1) (NHMB; 70% ethanol; NMB-PSYLL0008500); 2 m$, 7 f£, 3 immatures, 1 skin, Diamantina, Parque Biribiri, S18.10986, W43.37106, 1080 m, 14.ix.2021, *Macairea radula* (D.L. Queiroz) #1010(1) (NHMB; 70% ethanol; NMB-PSYLL0008562); 2 m$, 1 f£, same but Poço do Estudante, S18.2040, W43.6216, 1110 m, 16.ix.2019 (D. Burckhardt & D.L. Queiroz) #359 (NHMB; dry; NMB-PSYLL0007945); 4 f£, same but Cachoeira dos Cristais, S18.1569, W43.5998, 1070 m, 12.iv.2021, Cerrado vegetation along river, waterfalls and on rocks (D. Burckhardt & D.L. Queiroz) #399(0) (NHMB; 70% ethanol; NMB-PSYLL0008561); 1 m$, 5 f£, 6 immatures, São Gonçalo do Rio Preto, Parque Estadual do Rio Preto, Prainha, S18.1170, W43.3407, 760 m, 11‒12.ix.2019, *Macairea radula* (D. Burckhardt & D.L. Queiroz) #351(4) (NHMB; 70% ethanol; NMB-PSYLL0007943); 1 m$, same but, heliponto, S18.0910, W43.3421, 830 m, 12.ix.2019, *Macairea radula* (D. Burckhardt & D.L. Queiroz) #354(3) (NHMB; dry; NMB-PSYLL0007944); 1 f£, same but S18.0882, W43.3366, 790 m, 13.iv.2021, shrubby Cerrado vegetation and gallery forest (D. Burckhardt & D.L. Queiroz) #403(0) (NHMB; 70% ethanol; NMB-PSYLL0008501); 1 f£, same but S18.0882, W43.3366, 790 m, 13.iv.2021, shrubby Cerrado vegetation and gallery forest (D. Burckhardt & D.L. Queiroz) #404(0) (NHMB; 70% ethanol; NMB-PSYLL0008502); 1 m$, 1 f£, 1 skin, same but Casa de Hóspedes, S18.1263, W43.3570, 850 m, 16.iv.2021, planted trees (D. Burckhardt & D.L. Queiroz) #411(0) (NHMB; 70% ethanol; NMB-PSYLL0008503); 1 m$, Vargem Bonita, Parque Nacional da Serra da Canastra, Cachoeira Casca d’Anta, around park entrance, S24.8541/8573, W48.6982/7121, 850 m, 4–8.ix.2014, *Xylopia sericea* (D. Burckhardt & D.L. Queiroz) #141(10) (NHMB; slide; NMB-PSYLL0007933 [LSMelmac-64]); 3m$, 3 f£, 7 immatures, same but Cachoeira Casca d’Anta, plateau, S24.8541/8573, W48.6982/7121, 1160–1250 m, 6.ix.2014, *Macairea radula* (D. Burckhardt & D.L. Queiroz) #144(4) (MMBC, NHMB; slide, 70% ethanol; NMB-PSYLL0007926 [LSMelmac-64], NMB-PSYLL0007927 [LSMelmac-64], NMB-PSYLL0007930, NMB-PSYLL0007931, NMB-PSYLL0007932, NMB-PSYLL0007934, NMB-PSYLL0007942 [LSMelmac-64]).

**Description. *Adult.*** Coloration. Body pale to bright yellow. Antenna yellow, scape and pedicel pale brown, segments 4–8 with dark brown apices, segments 9–10 entirely dark brown. Mesopraescutum with two pale orange patches along the fore margin; mesoscutum with four pale orange longitudinal stripes. Forewing (Fig. 15E) bright yellow with white stripes along veins. Coxae pale brown, meracanthus pale yellow, distal tarsal segment brown. Abdomen and terminalia pale yellow. Younger specimens with less dark colour.

Structure. Forewing (Fig. 15E) oblong-oval, widest in the middle, narrowly, evenly rounded apically; C+Sc curved in apical third; pterostigma slightly narrower in the middle than r_1_ cell, mostly parallel-sided; Rs straight in basal two thirds, curved to fore margin apically; M longer than M_1+2_ and M_3+4_; Cu_1a_ curved in apical third, ending proximal of M fork; cell cu_1_ moderately wide; surface spinules present in all cells, leaving narrow spinule- free stripes along the veins, spinules absent in basal half of cell c+sc, dense, not forming distinct hexagons. Hindwing with 3 + 3 grouped costal setae. Metatibia bearing 8–10 apical spurs, arranged as 4–5 + 4–6, anteriorly separated by ca. 2 bristles.

Terminalia (Fig. 24A–G). Male. Proctiger weakly produced posteriorly; densely covered with long setae in apical two thirds. Subgenital plate, in lateral view, irregularly ovoid; dorsal margin curved, posterior margin slightly indented; with dense, long setae. Paramere, in lateral view, lanceolate; paramere apex, in lateral view, blunt, directed upwards, in dorsal view, directed inwards and slightly upwards, lacking a distinct sclerotised tooth; outer face with dense, moderately long setae in apical two thirds; inner face with dense, relatively short setae; posterior margin with longer setae. Proximal segment of aedeagus with apical part strongly subdivided. Distal segment of aedeagus with sinuous ventral margin and sinuate dorsal margin in basal half; ventral process, in lateral view, short, simple, situated in the middle of segment, truncate apically, without distinct lateral lobes; ventral process, in dorsal view, broad, shovel-shaped, widening to apex, indistinctly three-lobed; apical dilation relatively large, oblong-oval, slightly widening to apex which is rounded; sclerotised end tube short and sinuate. – Female terminalia cuneate; densely covered with setae. Dorsal margin of proctiger, in lateral view, weakly sinuate distal to circumanal ring, apex blunt; in dorsal view, apex blunt; circumanal ring, in dorsal view, vaguely cruciform. Subgenital plate, in lateral view, pointed apically; in ventral view, apex blunt.

***Fifth instar immature.*** Coloration. Yellow; antenna gradually becoming darker towards apex; legs pale to dark brown with darker tarsal segment; cephalothoracic sclerite, wing pads and caudal plate pale to bright brown.

Structure. Eye with one short simple ocular seta dorsally. Antennal segments with following numbers of pointed sectasetae: 1(0), 2(1), 3(0), 4(2), 5(0), 6(2), 7(1), 8(2), 9(0), 10(0). Forewing pad with 8 pointed sectasetae marginally and 13–35 sectasetae or smaller pointed lanceolate setae dorsally; hindwing pad with 3 marginal and 5–7 pointed sectasetae or lanceolate setae dorsally. Tarsal arolium broadly fan-shaped apically, ca. 1.5 times as long as claws. Abdominal tergites anterior to caudal plate with four irregular rows of 12–18, 12–20, 14–25 and 10–12 pointed lanceolate setae dorsally, respectively. Caudal plate dorsally with one row of 12–14 pointed lanceolate setae along anterior margin, a row of 16–20 pointed setae submedially, one group of 6–7 pointed sectasetae on each side in the middle and one group of 4–6 pointed sectasetae on each side subapically, anterior to circumanal ring. Caudal plate with large tubercules posteriorly; anterior margin remote from anterior margin of extra pore fields; extra pore fields forming continuous outer and inner bands, consisting of large oval patches; outer band very short medially, end pointing backwards. Circumanal ring small.

**Host plant.** *Macairea radula* (Bonpl.) DC. (Melastomataceae).

**Distribution.** Brazil (MG).

**Derivation of name.** Named after its host, *Macairea*.

**Comments.** *Melanastera macaireae* sp. nov. resembles *M. granulosi* sp. nov. and *M. tijuca* sp. nov. in the narrow yellow forewing with light stripes along the veins. From the former it differs in the shorter metatibia (MT 0.4 versus 0.6; MT/HW < 0.9 versus 1.1) and female terminalia (FP 0.4 versus 0.6). From the latter it differs in the fully developed median suture of the vertex (versus partly reduced) and the shape of the apical dilation of the distal segment of the aedeagus (sickle-shaped versus oblong-oval, in lateral view).

### 14 Melanastera tijuca sp. nov

(Figs 9D, 15F, 24H–M)

**Type material. Holotype m$: Brazil: RIO DE JANEIRO:** Parque Nacional da Tijuca, pico da Tijuca, S22.9429/9438, W43.2851/2862, 960‒1030 m, 11.iv.2019 (D. Burckhardt & D.L.

Queiroz) #325 (UFPR; slide; [LSMelsut-100]).

**Paratypes. Brazil: RIO DE JANEIRO:** 1 m$, 1 f£, same as holotype but (NHMB; slide; NMB-PSYLL0008150 [LSMelsut-100], NMB-PSYLL0008095 [LSMelsut-100]).

**Description. *Adult.*** Coloration. Body yellowish. Antenna brownish yellow, segments 4–7 with dark brown apices, segment 8 dark brown in its apical half, segments 9–10 entirely dark brown. Mesopraescutum with two orange patches along the fore margin; mesoscutum with four orange longitudinal stripes. Forewing (Fig. 15F) bright yellow, paler along veins, lacking dark pattern. Coxae pale brown, meracanthus pale yellow.

Structure. Median suture of vertex reduced in basal three quarters of vertex. Forewing (Fig. 15F) oblong-oval, widest in apical third, narrowly, evenly rounded apically; C+Sc curved in apical quarter; pterostigma slightly narrower in the middle than cell r_1_, weakly convex in apical third; Rs straight, curved in apical fifth; M longer than M_1+2_ and M_3+4_; Cu_1a_ curved in apical third, ending at M fork; cell cu_1_ narrow; surface spinules present in all cells, absent from basal half of cell c+sc, leaving narrow spinule-free stripes along the veins, forming hexagons, of a single row of spinules. Hindwing with 4 + 1–2 grouped costal setae. Metatibia bearing a posteriorly open crown of 9–11 apical spurs.

Terminalia (Fig. 24H–M). Male. Proctiger long, weakly produced posteriorly; densely covered with long setae in apical two thirds. Subgenital plate subglobular; dorsal margin strongly curved, posterior margin irregularly curved; with dense, long setae posteriorly. Paramere, in lateral view, irregularly sublanceolate; paramere apex, in lateral view, rounded, in dorsal view, directed inwards and slightly posteriad, with a small sclerotised tooth; outer and inner faces with dense, moderately long setae in apical two thirds; posterior margin with long setae. Proximal segment of aedeagus with apical part subdivided. Distal segment of aedeagus with sinuate ventral margin and almost straight dorsal margin in basal half; ventral process large and broad, simple, lacking lateral lobes, situated slightly distal of the middle of the segment, in lateral view, ventral process tongue-shaped, rounded apically; in dorsal view, ventral process narrow, oval, widest in basal third, rounded apically; apical dilation, in lateral view, sickle-shaped, apex subacute, directed ventrad, dorsal margin evenly convex, with a membranous lobe basally, apical dilation, in dorsal view, lanceolate, widest medially, blunt apically; sclerotised end tube short, weakly sinuate. – Female terminalia cuneate; densely covered with setae. Dorsal margin of proctiger, in lateral view, straight distal to circumanal ring, apex blunt; in dorsal view, apex blunt; circumanal ring, in dorsal view, cruciform. Subgenital plate, in lateral view, abruptly narrowing to apex in apical half, with ventral margin angular medially, pointed apically; in ventral view, blunt apically.

*Fifth instar immature.* Unknown.

**Host plant.** Unknown.

**Distribution.** Brazil (RJ).

**Derivation of name.** Named after the Parque Nacional da Tijuca, where the species was collected.

**Comments.** *Melanastera tijuca* sp. nov. resembles *M. macaireae* sp. nov. and *M. granulosi* sp. nov. in the narrow, yellow forewing with clear stripes along veins, from which it differs in the partly reduced median suture of the vertex and the structure of the male and female terminalia.

## The *falcata*-group

**Description. *Adult*.** Vertex and thorax covered with microscopical setae. Forewing (Fig. 15G–I) with small brown dots; pterostigma distinctly expanding towards the middle or apical third. Distal segment of aedeagus lacking ventral process.

***Immature.*** Antenna 10-segmented.

**Comments.** The group includes two species in Brazil. One is associated with Myristicaceae, while the host family of the other species is unknown.

### 15 Melanastera falcata sp. nov

(Figs 9E, 15G, H, 24N–S)

**Type material. Holotype m$: Brazil: MINAS GERAIS:** Coromandel, Fazenda Laje, S18.5748, W46.8881, 1050 m, 5.iii.2014 (D.L. Queiroz) #605(7) (UFPR; dry).

**Paratypes. Brazil: GOIÁS:** 1 m$, 3 f£, Mossâmedes, 7 km NE Mossâmedes, Fazenda Ribeirão Bonito, S16.1150, W50.1972, 640 m, 20.ii.2018, *Virola sebifera* (D. Burckhardt & D.L. Queiroz) #274(4) (NHMB; slide, 70% ethanol; NMB-PSYLL0007590, NMB- PSYLL0007583 [LSMelfal-18-4], NMB-PSYLL0008623 [LSMelfal-18-4]); 4 m$, 1 f£, 1 immature, 5 skins, Rio Verde, ca. 40 km E Jatai, BR-060, S17.8316, W51.3716, 820 m, 31.x.2012, *Miconia rubiginosa* (D. Burckhardt & D.L. Queiroz) #52(2) (NHMB; 70% ethanol; NMB-PSYLL0008558).—**Mato Grosso:** 6 m$, 2 f£, Chapada dos Guimarães, S12.4736, W55.7693, 800 m, 14.iv.2016 (L.A. Pezzini) #582 (NHMB; 70% ethanol; NMB-PSYLL0005737, NMB-PSYLL0005716); 3 m$, same but road MT-251, Casa do mel, S15.2246, W55.5033, 520 m, 25.viii.2018, *Virola sebifera* (D.L. Queiroz) #884(2) (NHMB; 70% ethanol; NMB-PSYLL0007597); 1 m$, 1 f£, Nova Mutum, S13.7991, W56.0955, 440 m, 13.vi.2016 (L.A. Pezzini) #824 (NHMB; 70% ethanol; NMB-PSYLL0005785); 1 m$, Sorriso, S12.4022, W55.8323, 390 m, 15.viii.2013 (L.A. Pezzini) #178 (NHMB; 70% ethanol; NMB-PSYLL0005622); 2 m$, 4 f£, same but S12.4022, W55.8323, 390 m, 15.viii.2013 (L.A. Pezzini) #179 (NHMB; 70% ethanol; NMB-PSYLL0005638); 2 m$, 4 f£, same but S12.4648, W55.7856, 340 m, 25.vii.2014 (L.A. Pezzini) #374 (NHMB; 70% ethanol; NMB-PSYLL0005704); 2 m$, 1 f£, same but S12.4284, W55.7963, 340 m, 26.vii.2014 (L.A. Pezzini) #381 (NHMB; 70% ethanol; NMB-PSYLL0005692); 2 f£, same but S12.4235, W55.8502, 400 m, 26.vii.2014 (L.A. Pezzini) #389 (NHMB; 70% ethanol; NMB-PSYLL0005683); 13 m$, same but S12.4261, W55.7856, 340 m, 27.ii.2015 (L.A. Pezzini) #480 (MMBC, NHMB; 70% ethanol; NMB-PSYLL0005687); 3 m$, 4 f£, 1 immature, same but S12.4261, W55.7944, 340 m, 27.ii.2015 (L.A. Pezzini) #485 (NHMB; 70% ethanol; NMB-PSYLL0005712); 6 m$, 2 f£, same but S12.4251, W55.8550, 400 m, 27.ii.2015 (L.A. Pezzini) #487 (NHMB; 70% ethanol; NMB-PSYLL0005610); 1 m$, 6 f£, 1 immature, same but S12.4259, W55.8507, 400 m, 27.ii.2015 (L.A. Pezzini) #488 (MMBC, NHMB; 70% ethanol; NMB-PSYLL0005613); 3 m$, 4 f£, 2 immatures, same but S12.4222, W55.8507, 390 m, 27.ii.2015 (L.A. Pezzini) #489 (NHMB; 70% ethanol; NMB-PSYLL0007589); 2 m$, 8 f£, 10 immatures, same but Fazenda Mazzardo II, S11.8714, W55.5957, 390 m, 26.ii.2015 (D.L. Queiroz) #678(2) (NHMB; slide, 70% ethanol; NMB- PSYLL0007571, NMB-PSYLL0007572, NMB-PSYLL0007574 [LSMelfal-18-3], NMB- PSYLL0007594, NMB-PSYLL0007595, NMB-PSYLL0007596); 1 m$, 1 f£, same but S11.8714, W55.5957, 350 m, 26.ii.2015 (D.L. Queiroz) #679 (NHMB; 70% ethanol; NMB-PSYLL0006043).—**Mato Grosso do Sul:** 3 m$, 2 f£, 9 immatures, Bataguassu, MS-395, near Rio Pardo, S21.6751, W52.4017, 300 m, 23.x.2021, riverine vegetation, *Virola sebifera* (D. Burckhardt & D.L. Queiroz) #485(2) (NHMB; 70% ethanol; NMB-PSYLL0008538).— **Minas Gerais:** 36 m$, 33 f£, 4 immatures, 3 skins, same as holotype but (MHNG, MMBC, NHMB, UFPR, USNM; dry, slide, 70% ethanol; NMB-PSYLL0007559, NMB- PSYLL0007602–NMB-PSYLL0007607, NMB-PSYLL0007581 [LSMelfal-18-1], NMB- PSYLL0007591, NMB-PSYLL0008207); 3 m$, 6 f£, same but S18.5677, W46.9034, 1060 m, 5.iii.2014 (D.L. Queiroz) #606 (NHMB; 70% ethanol; NMB-PSYLL0007568); 2 m$, 4 f£, Patos de Minas, Fazenda Barreiro, S18.6303, W46.6211, 880 m, 5.iii.2014 (D.L. Queiroz) #610(3) (NHMB; 70% ethanol; NMB-PSYLL0007570); 2 m$, 5 f£, 4 immatures, 2 skins, Uberlândia, Panga, S19.1837, W48.3961, 790 m, 8.ii.2018, *Virola sebifera* (D. Burckhardt & D.L. Queiroz) #261(1) (NHMB; slide, 70% ethanol; NMB-PSYLL0007573, NMB- PSYLL0007586 [voucher 63-3], NMB-PSYLL0007587 [voucher 63-2], NMB-PSYLL0007588 [voucher #261(1)]); 3 m$, 3 f£, 5 immatures, Vargem Bonita, Parque Nacional da Serra da Canastra, Cachoeira Casca d’Anta, around park entrance, S24.8541/8573, W48.6982/7121, 850–860 m, 4–8.ix.2014, *Guatteria ferruginea* (D. Burckhardt & D.L. Queiroz) #141(11) (NHMB; dry, slide, 70% ethanol; NMB- PSYLL00003109, NMB-PSYLL0007579 [LSMelfal-18-69], NMB-PSYLL0007580 [LSMelfal-18-69], NMB-PSYLL00003112, NMB-PSYLL00003111); 1 f£, same but near waterfall, S20.3090, W46.5231, 860 m, 5.ix.2014, *Miconia calvescens* (D. Burckhardt & D.L. Queiroz) #143(8) (NHMB; slide, 70% ethanol; NMB-PSYLL0007577 [LSMelfal-18-71], NMB-PSYLL0007557).—**Pará:** 74 m$, 60 f£, 42 immatures, 1 skin, Belém, Jardim Botânico Bosque Rodrigues Alves, S1.4300, W48.4533, 30 m, 14.iv.2013, *Virola sebifera* (D. Burckhardt & D.L. Queiroz) #103(3) (NHMB; dry, 70% ethanol; NMB-PSYLL0008543, NMB-PSYLL0008544, NMB-PSYLL0008482, NMB-PSYLL0008483, NMB-PSYLL0008577–NMB-PSYLL0008579); 6 m$, 5 f£, 9 immatures, 3 skins, same but *Guatteria schomburgkiana* (D. Burckhardt & D.L. Queiroz) #103(4) (NHMB; 70% ethanol; NMB-PSYLL0008496, NMB-PSYLL0008497); 1 m$, 2 f£, 1 immature, Belém, Embrapa Campus, S1.4150/4433, W48.4216/4433, 20 m, 8–15.iv.2013 (D. Burckhardt & D.L. Queiroz) #99 (NHMB; slide, 70% ethanol; NMB-PSYLL0007558, NMB-PSYLL0007584 [voucher 28-1], NMB-PSYLL0007585 [voucher #99]).

**Material not included in type series. Brazil: MINAS GERAIS:** 2 f£, Vazante, Fazenda Bocaina, S17.8913/8925, W46.9108/9112, 670–690 m, 22.ix.2011 (D. Burckhardt & D.L.Queiroz) #17 (NHMB; 70% ethanol; NMB-PSYLL0007557); 2 f£, Coromandel, Fazenda Laje, S18.6303, W46.6211, 1066 m, 6.iii.2014, *Miconia* sp. (D.L. Queiroz) #609(6) (NHMB; 70% ethanol; NMB-PSYLL0007569).—**Mato Grosso:** 2 f£, Nobres, Bom Jardim, S14.5504, W55.8366, 230 m, 13.iv.2016 (L.A. Pezzini) #568 (NHMB; 70% ethanol; NMB- PSYLL0005735); 1 f£,same but S14.5502, W55.8366, 240 m, 13.iv.2016 (L.A. Pezzini) #569 (NHMB; 70% ethanol; NMB-PSYLL0005714); 4 m$, 5 f£, Sinop, Embrapa Campus, S25.2346, W49.1059, 360 m, 27.ii.2015 (D.L. Queiroz) #680(4) (NHMB; slide, 70% ethanol; NMB-PSYLL0008624, NMB-PSYLL0007582 [LSMelfal-18-2], NMB-PSYLL0007592, NMB-PSYLL0007593); 2 f£,Sorriso, S12.4235, W55.8502, 400 m, 26.vii.2014 (L.A. Pezzini) #389 (NHMB; 70% ethanol; NMB-PSYLL0005683); 1 f£, same but Fazenda Mazzardo II, S12.4217, W55.8433, 15.viii.2013 (T. Mazzardo) (NHMB; slide; NMB-PSYLL0007576 [voucher M1-3]).—**Roraima:** 1 m$, Amajari, Tepequém, outside village, N3.76000, W61.7183, 720 m, 3.iv.2015 (D. Burckhardt & D.L. Queiroz) #155 (NHMB; slide; NMB- PSYLL0007575 [voucher 45-2A]).

**Description. *Adult.*** Coloration. Ochreous. Vertex (Fig. 9E) with dark brown dots. Antenna ochreous, scape and pedicel brown ventrally, segments 3–8 with dark brown apices, segments 9–10 entirely dark brown. Thorax ochreous with dark irregular dots; pronotum orange anteriorly, with each a submedian large and sublateral small white patch on either side which are linked with a white line; mesopraescutum with two orange brown patches along fore margin; mesoscutum with four broad orange brown longitudinal stripes margined by dark brown dots, medially a longitudinal row of irregularly spaced dark dots; meso- and metascutellum whitish with dark brown dots; metapostnotum with whitish keel. Forewing (Fig. 15G, H) almost colourless, slightly whitish or greyish with relatively large brown dots sparsely to moderately densely scattered over entire surface; with each a conspicuous dark brown dot at the base and apex of pterostigma, and at the apices of veins Rs, M_1+2_, M_3+4_, Cu_1a_ and Cu_1b_ as well as the apex of clavus. Large dark brown dots on femora and a dark subbasal ring on pro- and mesotibiae. Abdominal sclerites brown. Younger specimens with less expanded dark colour. Structure. Head, in lateral view, inclined at > 45° from longitudinal body axis. Vertex (Fig. 9E) trapezoidal, with distinct imbricate microsculpture and microscopical setae. Thorax distinctly arched, with microscopical setae. Forewing (Fig. 15G, H) oviform, widest distal of the middle, broadly rounded apically; wing apex situated in the middle of the margin of cell r_2_; C+Sc weakly curved in apical third; pterostigma slightly narrower than cell r_1_, widest in apical third; Rs weakly curved, slightly less in the middle than basally and apically; M longer than M_1+2_ and M_3+4_; Cu_1a_ moderately curved, ending at M fork; cell cu_1_ wide; surface spinules present in all cells, spinules absent or sparse in basal half of cell c+sc, leaving broad spinule- free stripes along the veins, forming hexagons of a single row of spinules. Hindwing with 3–5 + 3–5 grouped costal setae. Metatibia bearing 8–9 grouped apical spurs, arranged as 4 + 4–5, anteriorly separated by 2 bristles.

Terminalia (Fig. 24N–S). Male. Proctiger weakly curved posteriorly; densely covered with long setae in apical half. Subgenital plate subglobular, in lateral view, with dorsal margin strongly curved, posterior margin almost straight; with moderately long setae. Paramere, in lateral view, narrowly lanceolate; paramere apex, in lateral as well as dorsal views, blunt, directed upwards and slightly inwards, lacking a sclerotised tooth; both outer and inner faces with dense, short to moderately long setae; posterior margin with long setae. Proximal segment of aedeagus with apical part moderately subdivided. Distal segment of aedeagus straight in basal portion, with dorsal margin weakly sinuate, distinctly longer than apical dilation; ventral process absent; apical dilation, in lateral view, relatively narrow, sickle-shaped, with a minute membranous lobe basally at dorsal margin, in dorsal view, apical dilation oblong-oval with subparallel margins, rounded apically; sclerotised end tube short, weakly curved. – Female terminalia cuneate; densely covered with setae. Dorsal margin of proctiger, in lateral view, weakly concave posterior to circumanal ring, apex slender, pointed; in dorsal view, apex blunt; circumanal ring, in dorsal view, vaguely cruciform. Subgenital plate, in lateral view, abruptly narrowing to apex in apical half, forming long apical process, apex pointed; in ventral view, apex blunt.

***Fifth instar immature.*** Coloration. Light; cephalothoracic sclerite, antenna, wing pads, legs and caudal plate pale to dark brown.

Structure. Eye with one tiny ocular seta dorsally. Antennal segments with following numbers of pointed sectasetae: 1(0), 2(1), 3(0), 4(2), 5(0), 6(2), 7(1), 8(1), 9(0), 10(0). Forewing pad with 6 marginal and 4–5 dorsal pointed sectasetae; hindwing pad with 3 marginal and 1–2 dorsal pointed sectasetae. Metatibiotarsus relatively long, with 3 + 4 sectasetae in two rows on outer side; tarsal arolium broadly fan-shaped apically, 1.5 times as long as claws. Abdomen with four lateral pointed sectasetae on each side anterior to caudal plate. Caudal plate with anterior margin relatively close to anterior margin of extra pore fields; with five pointed sectasetae on either side laterally, one sectaseta in the middle on inner side of each extra pore field dorsally (absent in the sample #605(7)), and three pointed subapical sectasetae near circumanal ring on either side dorsally. Extra pore fields forming continuous outer and inner bands, consisting of small oval patches; outer band long medially, end pointing outwards. Circumanal ring small.

**Host plant.** *Virola sebifera* Aubl. (Myristicaceae).

**Distribution.** Brazil (GO, MG, MS, MT, PA, RR).

**Derivation of name.** From the Latin adjective *falcatus* = hooked, sickle-shaped, referring to the characteristic shape of the apical aedeagal dilation.

**Comments.** The specimens referred here to *M. falcata* sp. nov. are morphologically homogeneous but there are relatively large differences between some samples in the DNA sequences. Therefore, some samples are not included in the type series. The uncorrected p-distance between samples DLQ#605(7) (type material) and DLQ#680(4) (non-type material) is 7.13% for *COI* and 10% for *cytb* and between samples DLQ#605(7) and DLQ#678(5) (non-type material) 10% for *cytb*. The uncorrected p-distance between the samples DB- DLQ#141(11), DB-DLQ#143(8) from Minas Gerais, DB-DLQ#274(4) from Goiás and DLQ#680(4) (non-type material) is 7.40% for *COI* and 9.7–10% for *cytb*; between the samples DB-DLQ#141(11), DB-DLQ#143(8), DB-DLQ#274(4) and DLQ#678(5) (non-type material) it is 9.7–10% for *cytb*. This would suggest that *M. falcata* represents a complex of cryptic species.

### 16 Melanastera spinosa sp. nov

(Figs 9F, 15I, 24T–Z)

**Type material. Holotype m$: Brazil: AMAZONAS:** Manaus, Adolpho Ducke Forest Reserve, S2.9645, W59.9202, 100 m, 18.vi.1996, fogging of *Eschweilera rodriguesiana* (J.C.H. Guerrero, P.O.I. de Lima & G.P.L. Mesquita) F3-30(8) (INPA; dry).

**Paratypes. Brazil: AMAZONAS:** 1 f£, same as holotype but 17.xi.1995, fogging of *Eschweilera pseudodecolorans* (J.C.H. Guerrero, P.O.I. de Lima & G.P.L. Mesquita) F2-32(7) (INPA; dry); 1 m$, same but 21.vii.1995, fogging of *Eschweilera wachenheimii*, F2-35(2) (INPA; slide; [LSMelspin-95]); 1 m$, same but 21.vii.1995, F2-68 (BMNH; dry); 1 f£, same but 15.xi.1995, fogging of *Eschweilera pseudodecolorans*, F2-129(3) (NHMB; dry; NMB- PSYLL0008149); 1 m$, same but 16.viii.1995, fogging of *Pouteria glomerata*, F2-146(3) (INPA; dry); 1 m$, same but 18.vi.1996, fogging of *Eschweilera rodriguesiana*, F3-30(3) (MMBC; dry); 1 f£, same but 18.vi.1996, fogging of *Eschweilera rodriguesiana*, F3-30(6) (MMBC; dry); 1 m$, same but 18.vi.1996, fogging of *Eschweilera rodriguesiana*, F3-30(8) (NHMB; dry; NMB-PSYLL0008148); 1 m$, same but 18.vi.1996, fogging of *Eschweilera rodriguesiana*, F3-30(9) (INPA; dry); 1 f£, same but 18.vi.1996, fogging of *Eschweilera rodriguesiana*, F3-30(10) (INPA; slide; [LSMelspin-95]); 1 f£, same but 22.vi.1996, fogging of *Eschweilera pseudodecolorans*, F3-129(8) (BMNH; dry).

**Description. *Adult.*** Coloration. Body brownish yellow; head (Fig. 9F) and thorax with small dark brown dots. Clypeus brown at base. Antennal segments 3–8 with dark brown apices, segments 9–10 entirely brown. Mesopraescutum with two orange patches at fore margin; mesoscutum with four broad and, in the middle, one narrow longitudinal orange stripes; mesoscutellum and metanotum with dark brown stripe in the middle. Forewing (Fig. 15I) brownish yellow with scattered contrasting dark brown dots; apex of pterostigma and apices of Rs, M_1+2_, M_3+4_, Cu_1a_ and Cu_1b_ dark brown. Femora with brown dots. Abdominal sternites dark brown. Young specimens with less extended dark colour.

Structure. Head, in lateral view, inclined at < 45° from longitudinal body axis. Vertex (Fig. 9F) trapezoidal, with distinct imbricate microsculpture and microscopical setae. Thorax weakly arched, with microscopical setosity. Forewing (Fig. 15I) oval, widest in the middle, evenly, broadly rounded apically; with wing apex in the middle of the margin of cell r_2_; C+Sc weakly curved in apical third; pterostigma as wide as or slightly narrower as r_1_ cell in the middle, weakly convex in apical third; Rs almost straight, weakly curved to fore margin apically; M longer than M_1+2_ and M_3+4_; Cu_1a_ weakly convex, ending at level of M fork; cell cu_1_ relatively narrow; surface spinules present in all cells, spinules absent at least in basal third of cell c+sc, leaving relatively broad spinule-free stripes along the veins, forming hexagons of a single or double row of spinules. Hindwing with 4–5 + 2–3 grouped costal setae. Metatibia bearing 6–7 apical spurs, arranged as 3 + 3–4, anteriorly separated by ca. 5 bristles.

Terminalia (Fig. 24T–Z). Male. Proctiger narrowly cylindrical, almost straight posteriorly; densely covered with long setae in apical two thirds. Subgenital plate, in lateral view, narrowly ovoid, dorsal margin strongly angular in apical thirds, posterior margin slightly convex; with sparse, short setae. Paramere, in lateral view, digitiform, widest basally, narrowing to apex; paramere apex, in lateral view, rounded, directed upwards, in dorsal view, truncate with anterior angle sharp, directed inwards, bearing two spur-like setae on its inner face; outer face with dense, short and long setae in apical two thirds; inner face with sparse, short setae and a row of six spur-like setae in basal third near posterior margin, well-visible in posterior view; with very long setae posteriorly. Proximal segment of aedeagus with apical part moderately subdivided. Distal segment of aedeagus slender in basal half, with dorsal margin sinuate; ventral process absent, in lateral view, ventral margin sinuate, in dorsal view, slightly expanded in apical third; apical dilation moderately long and narrow, in lateral view, fusiform, rounded apically, with distinct membranous extension basally at dorsal margin, in dorsal view, apical dilation with subparallel margins, only slightly widening to apex; sclerotised end tube moderately long and straight. – Female terminalia cuneate; covered with setae. Dorsal margin of proctiger, in lateral view, sinuate posterior to circumanal ring, apex slightly upturned, pointed; in dorsal view, apex subacute or blunt; circumanal ring, in dorsal view, vaguely cruciform. Subgenital plate, in lateral view, abruptly narrowing to apex in apical third, slightly upturned and pointed apically; in ventral view, apex blunt.

*Fifth instar immature.* Unknown.

**Host plant.** Unknown. The type material was fogged from *Eschweilera pseudodecolorans* S.A.Mori, *E. rodriguesiana* S.A.Mori, *E. wachenheimii* (Benoist) Sandwith and *Pouteria glomerata* (Miq.) Radlk. (Lecythidaceae), which are unlikely hosts.

**Distribution.** Brazil (AM).

**Derivation of name.** From the Latin noun *spina* = thorn, referring to the spur-like setae on the inner face of the paramere.

**Comments.** *Melanastera spinosa* sp. nov. resembles *M. cabucu* sp. nov. in the forewing shape, venation and colour but differs in the details of the male and female terminalia.

### The *amazonica*-group

**Description. *Adult*.** Head, in lateral view, inclined at 45° or more from longitudinal axis of body (Fig. 7D). Vertex (Figs 9G–J, 10A, B) trapezoidal, with distinct imbricate microsculpture and microscopical setae. Thorax weakly to moderately arched, with microscopical setae. Forewing (Figs 15J, 16A–E) beset with brown dots; pterostigma strongly expanding towards the middle or apical third, shorter than 4.0 times as wide; vein R_1_ strongly curved medially relative to costal wing margin. Paramere lanceolate. Ventral process of the distal aedeagal segment, in lateral view, claw-like and directed dorsad.

***Immature.*** Antenna 10-segmented.

**Comments.** The six Brazilian species are associated with Annonaceae. Due to a similar forewing shape and venation as well as the structure of the distal segment of the the aedeagus, the six species and also *M. olgae* sp. nov. from the *olgae*-group could be closely related.

### 17 Melanastera amazonica sp. nov

(Figs 9G, 15J, 25A–F)

**Type material. Holotype m$: Brazil: AMAZONAS:** Manaus, Bairro Tarumã-Açu, BR-174 km 1, S2.9466, W60.0333, 100 m, 2.v.2014, *Annona foetida* (D. Burckhardt & D.L. Queiroz) #134(2) (UFPR; dry).

**Paratypes. Brazil: AMAZONAS:** 5 m$, 4 f£, 13 immatures, 2 skins, same as holotype but (MMBC, NHMB; dry, slide, 70% ethanol; NMB-PSYLL0007735, NMB-PSYLL0007673, NMB-PSYLL0007750–NMB-PSYLL0007754, NMB-PSYLL0007757 [LSMelama-13]); 1 f£, Rio Preto da Eva, Embrapa, Fazenda Rio Urubu, S2.4300/4683, W59.5616/5683, 50–100 m, 23–24.iv.2014 (D. Burckhardt & D.L. Queiroz) #131 (NHMB; dry; NMB- PSYLL0007674); 1 m$, 1 f£, 13 immatures, same but S2.4716, W59.5900, 120 m, 25.iv.2014, *Guatteria* sp. (D. Burckhardt & D.L. Queiroz) #132(3) (NHMB; 70% ethanol; NMB-PSYLL0007733).—**Mato Grosso:** 2 m$, Sinop, road to Cláudia, S11.7468, W55.3674, 360 m, 5.vi.2014 (L.A. Pezzini & A.M.B. Sedano) #287 (NHMB; dry, slide; NMB-PSYLL0005588 [voucher 53-1]).

**Material not included in type series. Brazil: AMAZONAS:** 4 m$, 4 f£, 6 immatures, Manaus, Sede Embrapa, Campo Experimental, AM-10 km 29, S2.8950, W59.9733, 100 m, 27–30.iv.2014, *Guatteria megalophylla* (D. Burckhardt & D.L. Queiroz) #133(7) (NHMB; 70% ethanol; NMB-PSYLL0007732 [LSMelama-57]); 2 m$, 5 f£, 18 immatures, 1 skin, Iranduba, Embrapa, Campo Experimental, Caldeirão, S3.2516/2566, W60.2216/2233, 30–50 m, 14–17.iv.2014, *Guatteria* sp. (D. Burckhardt & D.L. Queiroz) #125(7) (NHMB; slide, 70% ethanol; NMB-PSYLL0007730, NMB-PSYLL0007763–NMB-PSYLL0007765 [LSMelama- 29]).

**Description. *Adult.*** Coloration. Body whitish with dark brown dots. Vertex in the middle of either half with orange patch. Antenna yellowish, scape and pedicel light brown, segments 3–8 with brown apices, segments 9–10 entirely dark brown. Pronotum with transverse orange-brown patch on either side; mesopraescutum with two orange patches at fore margin; mesoscutum with four broad orange longitudinal stripes, margined by brown dots, and a row of dots in the middle; mesoscutellum with three and metanotum with one narrow longitudinal stripes, consisting of brown dots. Forewing (Fig. 15J) yellow with brown, moderately dense dots, covering the whole wing except for nodal line which appears slightly lighter; base and apex of pterostigma and apices of veins Rs, M_1+2_, M_3+4_, Cu_1a_ and Cu_1b_ dark brown. Femora with brown dots. Abdomen and terminalia brown. Younger specimens with less expanded dark colour.

Structure. Forewing (Fig. 15J) ovoid, broadest in apical third, broadly, slightly unevenly rounded apically; wing apex situated in cell r_2_ close to apex of M_1+2_; C+Sc curved in apical third; pterostigma broader than cell r_1_ in apical third, strongly convex; Rs weakly convex in basal two thirds, stronger curved apically; M longer than M_1+2_ and M_3+4_; Cu_1a_ moderately irregularly curved, ending at M fork; cell cu_1_ moderately wide; surface spinules present in all cells, dense, leaving narrow spinule-free stripes along the veins, forming hexagons of a double row of spinules, sparse in basal half of cell c+sc. Hindwing with 7–10 ungrouped or 7 + 3 grouped costal setae. Metatibia bearing 5–6 grouped apical spurs, arranged as 2–3 + 3–4, anteriorly separated by 4–6 bristles.

Terminalia (Fig. 25A–F). Male. Proctiger weakly produced posteriorly; densely covered with long setae in apical two thirds. Subgenital plate irregularly ovoid; dorsal margin slightly curved, posterior margin sinuate; with long setae on most of the surface. Paramere, in lateral view, subtriangular, narrowing from base to apex; paramere apex, in lateral view, blunt, directed upwards and slightly anteriad, in dorsal view, apex subacute, directed upwards, inwards and slightly anteriad, bearing a small sclerotised tooth; outer and inner faces with dense, long setae; posterior margin with longer setae. Proximal segment of aedeagus with apical part moderately subdivided. Distal segment of aedeagus with dorsal margin weakly sinuate in basal half; ventral process situated slightly distal of the middle of the segment, shovel-like, in lateral view, relatively short and broad, with a sclerotised base ventrally and with claw-like apex directed dorsad; in dorsal view, ventral process slightly broader than apical dilation, subcircular, and with short, broad, apically truncate median lobe arising from the base ventrally; apical dilation, in lateral view, slightly widening towards rounded apex on ventral side, with a small membranous sack at dorsal margin basally; in dorsal view, apical dilation elongate, with subparallel margins, rounded apically; sclerotised end tube short and weakly curved. – Female terminalia cuneate; densely covered with setae. Dorsal margin of proctiger, in lateral view, weakly concave distal to circumanal ring, apex subacute; in dorsal view, apex pointed; circumanal ring, in dorsal view, distinctly cruciform. Subgenital plate, in lateral view, gradually narrowing to apex in apical half, pointed apically; in ventral view, apex blunt.

***Fifth instar immature.*** Coloration. Pale yellow; antenna brownish yellow to pale brown, gradually becoming darker towards apex; cephalothoracic sclerite, wing pads, legs and caudal plate brownish yellow to pale brown.

Structure. Eye with one short, simple ocular seta dorsally. Antennal segments with following numbers of pointed sectasetae: 1(0), 2(1), 3(0), 4(2), 5(0), 6(2), 7(1), 8(1), 9(0), 10(0). Forewing pad with five marginal pointed sectasetae; hindwing pad with two marginal pointed sectasetae; both wing pads lacking sectasetae dorsally. Tarsal arolium broadly triangular apically, 1.5 times as long as claws. Abdomen with 2–3 lateral pointed sectasetae on each side anterior to caudal plate. Caudal plate with anterior margin close to anterior margin of extra pore fields; with 5–6 pointed sectasetae on either side laterally, and three pointed sectasetae subapically, on either side of circumanal ring dorsally. Extra pore fields forming continuous outer and inner bands, consisting of small oval patches; outer band long medially, end pointing outwards. Circumanal ring small.

**Host plants.** *Annona foetida* Mart., *Guatteria megalophylla* Diels (Annonaceae).

**Distribution.** Brazil (AM, MT).

**Derivation of name.** Latinised form of the name Amazonas, the State in Brazil, where the species was collected.

**Comments.** *Melanastera amazonica* sp. nov. resembles *M. cacantis* sp. nov., *M. francisi* sp. nov., *M. olgae* sp. nov., *M. roraima* sp. nov., *M. tubuligera* sp. nov. and *M. xylopiae* sp. nov. in the apically broadly, unevenly rounded forewing with broad pterostigma, the paramere apex bearing a small sclerotised tooth, and the claw-like ventral process of the distal segment of the aedeagus. It differs from these species in the absence of a posterior lobe on the paramere. From *M. olgae* it differs in the dotted forewings (versus uniformly coloured), from *M. cacantis*, *M. francisi*, *M. roraima* and *M. tubuligera* in the less densely spaced dark dots on the forewing, and from *M. xylopiae* in the triangular (versus narrowly lanceolate) paramere and the shorter apical dilation of the distal segment of the aedeagus.

Some specimens from sample DB-DLQ#134(2) collected on *A. foetida* differ considerably from the remainder of the sample, as well as from other samples (DB&DLQ#125(7) and DB&DLQ#133(7), from *Guatteria* spp.) in the molecular markers: the uncorrected p-distance between them being 14% for *COI* and 13.3% for *cytb*. Although no morphological differences have been found between these specimens, the high genetic distance would suggest that *M. amazonica* sp. nov. may represent a complex of cryptic species.

### 18 Melanastera cacantis sp. nov

(Figs 9H, 16A, 25G–M)

**Type material. Holotype m$: Brazil: MINAS GERAIS:** Vazante, Fazenda Bocaina, S17.8930, W46.9111, 660 m, 28‒31.viii.2019, *Annona cacans* (D. Burckhardt & D.L. Queiroz) #348(1) (UFPR; dry).

**Paratypes. Brazil: MINAS GERAIS:** 1 m$, 5 f£, 9 immatures, Vazante, Fazenda Bocaina, S17.8930, W46.9111, 660 m, 28‒31.viii.2019, *Annona cacans* (D. Burckhardt & D.L. Queiroz) #348(1) (NHMB; dry, slide, 70% ethanol; NMB-PSYLL0007800, NMB- PSYLL0007923, NMB-PSYLL0007983, NMB-PSYLL0007814, NMB-PSYLL0007815, NMB-PSYLL0007816 [LSMelcac-117], NMB-PSYLL0007817 [LSMelcac-117], NMB- PSYLL0007818).

**Description. *Adult.*** Coloration. Similar to *M. amazonica* sp. nov. but with denser dark brown dots on forewing.

Structure. Forewing (Fig. 16A) ovoid, broadest in apical third, broadly, unevenly rounded apically, with wing apex cell r_2_ close to apex of M_1+2_; C+Sc curved in distal third; pterostigma slightly wider in the apical third than r_1_ cell, strongly convex; Rs strongly convex apart from median third which is relatively straight; M longer than M_1+2_ and M_3+4_; Cu_1a_ unevenly curved, ending at M fork; cell cu_1_ wide; surface spinules present in all cells, leaving narrow spinule- free stripes along the veins, spinules densely spaced, not forming distinct hexagons, sparse at base of cell c+sc. Hindwing with 4–5 + 3–4 grouped costal setae. Metatibia bearing 7–9 grouped apical spurs, arranged as 3–4 + 4–5, anteriorly separated by 3–4 bristles.

Terminalia (Fig. 25G–M). Male. Proctiger evenly produced posteriorly; densely covered with long setae in apical two thirds. Subgenital plate irregularly ovoid; dorsal margin weakly sinuate, posterior margin weakly convex; with dense, long setae. Paramere, in lateral view, lamellar, irregularly tapering to apex, posterior margin bearing a lobe in basal half; paramere apex, in lateral view, subacute, directed anteriad, in dorsal view, apex subacute, directed inwards, slightly upwards and slightly anteriad, with a small sclerotised tooth; outer and inner faces with dense, moderately long setae; posterior margin with long setae. Proximal segment of aedeagus with apical part moderately subdivided. Distal segment of aedeagus almost straight in basal half, with weakly sinuate dorsal margin; ventral process situated slightly distal of the middle of the segment, shovel-like, in lateral view, relatively short and broad, with a sclerotised base ventrally and with claw-like apex directed dorsad; in dorsal view, ventral process about as wide as apical dilation, subrectangular, constricted medially and subapically, apex broad, truncate, with a distinct sclerotised bar; apical dilation, in lateral view, narrow, with subparallel margins, rounded apically, in dorsal view, apical dilation oblong-oval, with subparallel lateral margins, apex rounded; sclerotised end tube short, weakly sinuate. – Female terminalia cuneate; densely covered with setae. Dorsal margin of proctiger, in lateral view, straight, apex subacute, in dorsal view, apex blunt; circumanal ring, in dorsal view, distinctly cruciform. Subgenital plate, in lateral view, gradually narrowing to apex in apical half, pointed apically; in ventral view, apex blunt.

***Fifth instar immature.*** Coloration. Pale yellow; cephalothoracic sclerite, antenna, wing pads, legs and caudal plate pale brown.

Structure. Eye with one short, simple ocular seta dorsally. Antennal segments with following numbers of pointed sectasetae: 1(0), 2(1), 3(0), 4(2), 5(0), 6(2), 7(1), 8(1), 9(0), 10(0). Forewing pad with five marginal pointed sectasetae; hindwing pad with two marginal pointed sectasetae; both wing pads lacking sectasetae or large lanceolate setae dorsally. Metatibiotarsus long; tarsal arolium broadly fan-shaped apically, 1.5 times as long as claws. Abdomen with 3–4 lateral pointed sectasetae on either side anterior to caudal plate. Caudal plate with anterior margin relatively close to anterior margin of extra pore fields; with 2+3 pointed sectasetae on either side laterally, and three pointed sectasetae subapically, on either side of circumanal ring dorsally. Extra pore fields forming continuous outer and inner bands, consisting of relatively small oval and rounded patches; outer band long medially, end pointing outwards. Circumanal ring small.

**Host plant.** *Annona cacans* Warm. (Annonaceae).

**Distribution.** Brazil (MG).

**Derivation of name.** Named after its host, *A. cacans* Warm. (Annonaceae).

**Comments.** *Melanastera cacantis* sp. nov. resembles *M. francisi* sp. nov., *M. olgae* sp. nov., *M. roraima* sp. nov., *M. tubuligera* sp. nov. and *M. xylopiae* sp. nov. in the posteriorly lobed paramere and the claw-like ventral process of the distal aedeagal segment but differs in the short sclerotised end tube of the ductus ejaculatorius and details of the the paramere and the distal segment of the aedeagus.

### 19 Melanastera xylopiae sp. nov

(Figs 7D, 9I, 16B, 25N–S, 40C, D)

**Type material. Holotype m$: Brazil: RORAIMA:** Boa Vista, Embrapa campus, N2.7550, W60.7300, 80 m, 31.iii.2015, *Xylopia aromatica* (D. Burckhardt & D.L. Queiroz) #151(1) (UFPR; dry).

**Paratypes. Brazil: AMAZONAS:** 2 m$, 7 f£, 2 immatures, Novo Airão, along road from Novo Airão to Manacapuru, S2.7033/3.1366, W60.7033/7316, 60 m, 22.iv.2014, *Xylopia nitida* (D. Burckhardt & D.L. Queiroz) #130(2) (NHMB; 70% ethanol; NMB- PSYLL0007386).—**BAHIA:** 2 m$, 2 f£, Luís Eduardo Magalhães, S11.9877 W45.1728, 740 m, 25.ix.2012 (D.L. Queiroz) #357(-) (MMBC, NHMB; slide, 70% ethanol; NMB- PSYLL0007397, NMB-PSYLL0007415).—**GOIÁS:** 2 m$, 1 f£, Alto Paraíso do Goiás, near São Jorge, Parque Nacional da Chapada dos Veadeiros, around researchers’ accommodations, S18.5610, W46.9020, 1060 m, 15.ii.2018, *Xylopia aromatica* (D. Burckhardt & D.L. Queiroz) #265(2) (NHMB; 70% ethanol; NMB-PSYLL0007392); 1 m$, 10 f£, 19 immatures, 1 skin, same but park headquarters, S18.5610, W46.9020, 970 m, 17.ii.2018, *Xylopia aromatica* (D. Burckhardt & D.L. Queiroz) #268(2) (NHMB; 70% ethanol; NMB-PSYLL0007393); 2 m$, same but São Jorge, S14.1783, W478097, 1010 m, 17‒18.ii.2018, *Xylopia aromatica* (D. Burckhardt & D.L. Queiroz) #269(3) (NHMB; 70% ethanol; NMB-PSYLL0007394); 10 m$, 4 f£, Mineiros, ca. 15 km NW of Mineiros, BR-364, S17.8310, W51.3709, 860 m, 1.xi.2012, *Xylopia aromatica* (D. Burckhardt & D.L. Queiroz) #53(2) (NHMB; 70% ethanol; NMB- PSYLL0007376); 2 m$, 1 f£, Rio Verde, ca. 20 km W Rio Verde, BR-060, S17.8146, W51.0918, 830 m, 31.x.2012 (D. Burckhardt & D.L. Queiroz) #51(-) (NHMB; 70% ethanol; NMB-PSYLL0007375).—**MATO GROSSO DO SUL:** 9 m$, 11 f£, Corguinho, S19.8203, W54.8203, 300 m, 15.xi.2012, *Xylopia aromatica* (D. Burckhardt & D.L. Queiroz) #72(3) (NHMB; 70% ethanol; NMB-PSYLL0007382); 1 m$, 4 f£, Campo Grande, near BR-163, S20.8907/8948, W54.5713/6599, 440–450 m, 16.xi.2012, *Xylopia aromatica* (D. Burckhardt & D.L. Queiroz) #74(9) (NHMB; 70% ethanol; NMB-PSYLL0007385); 2 m$, Rio Verde, MS-427, South of Rio Verde, S19.0183, W54.8583, 470 m, 14.xi.2012, *Xylopia aromatica* (D. Burckhardt & D.L. Queiroz) #69(4) (NHMB; 70% ethanol; NMB-PSYLL0007383); 3 m$, 2 f£, Rio Verde do Mato Grosso, BR-163, S18.9281/9519, W54.8357/9339, 350–440 m, 13.xi.2012, *Xylopia aromatica* (D. Burckhardt & D.L. Queiroz) #68(2) (NHMB; 70% ethanol; NMB-PSYLL0005632); 2 m$, 4 f£, Rochedo, MS-080, S19.9683, W54.6500, 350 m, 15.xi.2012, *Xylopia aromatica* (D. Burckhardt & D.L. Queiroz) #73(1) (NHMB; 70% ethanol; NMB-PSYLL0007384).—**Mato Grosso:** 4 m$, 7 f£, Acorizal, MT-010, S15.1950, W56.2566, 200 m, 4.xi.2012 (D. Burckhardt & D.L. Queiroz) #58(0) (NHMB; 70% ethanol; NMB-PSYLL0007377); 2 m$, 4 f£, Bataguassu, MS-395, near Rio Pardo, S21.6751, W52.4017, 300 m, 23.x.2021, riverine vegetation, *Xylopia aromatica* (D. Burckhardt & D.L. Queiroz) #485(1) (NHMB; 70% ethanol; NMB-PSYLL0008553); 1 m$, 1 f£, Chapada dos Guimarães, road MT-251, Restaurante caminho das Aguas, S15.3931, W55.9870, 170 m, 25.viii.2018, *Xylopia aromatica* (D.L. Queiroz) #883(2) (NHMB; 70% ethanol; NMB- PSYLL0007404); 4 m$, 4 f£, Itiquira, 112 km S Rondonópolis, BR-163, S17.5139, W54.7401, 410 m, 12.xi.2012, *Xylopia aromatica* (D. Burckhardt & D.L. Queiroz) #66(1) (NHMB; 70% ethanol; NMB-PSYLL0007381); 9 m$, 15 f£, Nobres/Rosario do Oeste, Bom Jardim, S14.5282/5766, W55.8047/9174, 250–300 m, 10.xi.2012, *Xylopia aromatica* (D. Burckhardt & D.L. Queiroz) #65(1) (NHMB; 70% ethanol; NMB-PSYLL0007378); 5 m$, 7 f£, Nova Mutum, S13.7023, W56.0712, 410 m, 12.vi.2016, *Guatteria* sp. (L.A. Pezzini) #808 (NHMB; slide, 70% ethanol; NMB-PSYLL0005778, NMB-PSYLL0007426 [voucher 31/351]); 2 m$, 1 f£, same but S13.8078, W56.1128, 490 m, 13.vi.2016 (L.A. Pezzini) #834 (NHMB; 70% ethanol; NMB-PSYLL0005801); 2 m$, 4 f£, same but S13.8091, W56.1134, 490 m, 13.vi.2016 (L.A. Pezzini) #836 (NHMB; 70% ethanol; NMB-PSYLL0005809); 1 f£, Ribas do Rio Pardo, Ribas do Rio Pardo, S20.4559, W53.7426, 390 m, 22.x.2021, edge of Cerrado, *Xylopia aromatica* (D. Burckhardt & D.L. Queiroz) #481(6) (NHMB; 70% ethanol; NMB-PSYLL0008550); 8 m$, 4 f£, Ribas do Rio Pardo, BR-262, S20.4145, W53.2245, 430 m, 22.x.2021, edge of Cerrado, *Xylopia aromatica* (D. Burckhardt & D.L. Queiroz) #483(2) (NHMB; 70% ethanol; NMB-PSYLL0008551); 1 f£, Sorriso, Fazenda Mazzardo II, S12.4288, W55.7965, 350 m, 26.vii.2014 (T. Mazzardo) #P009 (NHMB; dry; NMB- PSYLL00003012); 1 f£, same but S12.4281, W55.7964, 350 m, 26.vii.2014 (T. Mazzardo) #P011 (NHMB; dry; NMB-PSYLL00003013); 1 m$, 3 f£, same but S12.4235, W55.8502, 400 m, 26.vii.2014 (T. Mazzardo) #P014 (NHMB; dry; NMB-PSYLL00003008, NMB- PSYLL00003011, NMB-PSYLL00003010); 1 m$, same but S12.4217, W55.8432, 350 m, 15.viii.2013 (T. Mazzardo) (NHMB; slide; NMB-PSYLL0007407); 1 m$, Tabaporã, Fazenda Crestani, S11.3133/3366, W55.9616/9750, 330–380 m, 6–8.xi.2012 (D. Burckhardt & D.L. Queiroz) #62(0) (NHMB; 70% ethanol; NMB-PSYLL0007379); 4 f£, same but S11.3133/3366, W55.9616/9750, 330–380 m, 6–8.xi.2012, *Xylopia aromatica* (D. Burckhardt & D.L. Queiroz) #62(9) (NHMB; 70% ethanol; NMB-PSYLL0007380); 1 m$, 2 f£, Terra Nova do Norte, S10.4788, W55.0549, 340 m, 28.v.2016, *Xylopia* sp. (L.A. Pezzini) #711 (NHMB; 70% ethanol; NMB-PSYLL0005760); 11 m$, 19 f£, 7 immatures, Três Lagoas, BR- 158, Lotrans Armazem, S20.9779, W51.8548, 410 m, 23.x.2021, edge of cerrado, *Xylopia aromatica* (D. Burckhardt & D.L. Queiroz) #484(2) (NHMB; 70% ethanol; NMB- PSYLL0008552).—**Minas Gerais:** 1 f£, Buenópolis, Parque Estadual da Serra do Cabral, Cascalheira, S17.9109, W44.2000, 740 m, 9.iv.2021, cerrado vegetation, *Xylopia aromatica* (D. Burckhardt & D.L. Queiroz) #390(4) (NHMB; 70% ethanol; NMB-PSYLL0008549); 2 m$, 2 f£, Buritizeiro, along route BR-365, S17.5169, W45.2356, 860 m, 23.ix.2019, *Xylopia aromatica* (D. Burckhardt & D.L. Queiroz) #379(2) (NHMB; 70% ethanol; NMB- PSYLL0007400); 2 m$, 1 f£, Chapada Gaúcha, Parque Nacional Grande Sertão Veredas, confluence of the rivers Rio Preto and Rio da Onça, S15.1188, W45.7314, 700 m, 2.v.2021, cerrado vegetation, *Xylopia emarginata* (D. Burckhardt & D.L. Queiroz) #449(7) (NHMB; 70% ethanol; NMB-PSYLL0008499); 1 m$, 2 f£, Coromandel, Fazenda Laje, S18.5610, W46.9020, 1060 m, 12.ii.2018, *Xylopia aromatica* (D. Burckhardt & D.L. Queiroz) #263(6) (NHMB; 70% ethanol; NMB-PSYLL0007391); 1 m$, 7 f£, same but 23.viii.2019, *Xylopia aromatica* (D. Burckhardt & D.L. Queiroz) #346(3) (NHMB; 70% ethanol; NMB-PSYLL0007399); 5 m$, 16 f£, 23 immatures, 6 skins, same but S18.57477, W46.88815, 1050 m, 5.iii.2014, *Xylopia aromatica* (D.L. Queiroz) #605(3) (NHMB; slide, 70% ethanol; NMB- PSYLL0007403, NMB-PSYLL0007408, NMB-PSYLL0007409, NMB-PSYLL0007410, NMB-PSYLL0007417, NMB-PSYLL0007428 [LSMelverg-21]); 1 f£, same but S18.56773, W46.90342, 1060 m, 5.iii.2014 (D.L. Queiroz) #606 (NHMB; 70% ethanol; NMB- PSYLL0007402); 2 f£, same but S18.55631, W46.89574, 1060 m, 4.iii.2019 (D.L. Queiroz) #921 (NHMB; 70% ethanol; NMB-PSYLL0007406); 4 m$, 2 f£, São Gonçalo do Rio Preto, Parque Estadual do Rio Preto, Prainha, S18.1170, W43.3407, 760 m, 11‒12.ix.2019, *Xylopia aromatica* (D. Burckhardt & D.L. Queiroz) #351(9) (NHMB; 70% ethanol; NMB- PSYLL0007398); 1 m$, 2 f£, same but Poço do Veado, S18.1099, W43.3385, 760 m, 11‒ 12.ix.2019, *Xylopia aromatica* (D. Burckhardt & D.L. Queiroz) #352(1) (NHMB; 70% ethanol; NMB-PSYLL0007396); 2 f£, Vanzante, Fazenda Bainha, S17.88815, W46.92126, 630 m, 12.vii.2018 (D.L. Queiroz) (NHMB; 70% ethanol; NMB-PSYLL0007405); 1 m$, 3 immatures, 1 skin, same but Guariba, S17.8791/8795, W46.9187/9202, 640–660 m, 20.ix.2011, *Xylopia aromatica* (D. Burckhardt & D.L. Queiroz) #13(5) (NHMB; 70% ethanol; NMB-PSYLL0008545); 2 f£, same but near source of Curtume river, S17.8891/8942, W46.9196/9205, 640–690 m, 11.ix.2014, *Xylopia aromatica* (D. Burckhardt & D.L. Queiroz) #145(2) (NHMB; dry; NMB-PSYLL0004044); 6 m$, 9 f£, same but around the house and eucalypt plantation, S17.8930, W46.9111, 640 m, 12.ix.2014, *Xylopia aromatica* (D. Burckhardt & D.L. Queiroz) #147(1) (NHMB; 70% ethanol; NMB-PSYLL0007388); 5 m$, 10 f£, 4 immatures, 11 skins, same but Curtume river, S17.8822/8863, W46.9213/9229, 650– 660 m, 21–22.ix.2011, *Xylopia aromatica* (D. Burckhardt & D.L. Queiroz) #16(8) (NHMB; 70% ethanol; NMB-PSYLL0008546); 10 m$, 12 f£, 1 immature, same but S17.8913/8925, W46.9108/9112, 670–690 m, 22.ix.2011, *Xylopia aromatica* (D. Burckhardt & D.L. Queiroz) #17(6) (NHMB; slide, 70% ethanol; NMB-PSYLL0008158, NMB-PSYLL0008159, NMB- PSYLL0008161, NMB-PSYLL0008547); 3 m$, 3 f£, 1 immature, same but S17.8828/8852, W46.9174/9181, 670–690 m, 23.ix.2011, *Xylopia aromatica* (D. Burckhardt & D.L. Queiroz) #20(2) (NHMB; dry, slide; NMB-PSYLL0008300–NMB-PSYLL0008304, NMB- PSYLL0008572, NMB-PSYLL0008573); 1 f£, 2 immatures, same but S17.8817/8832, W46.9164/9165, 660–670 m, 26.xii.2011, *Xylopia aromatica* (D. Burckhardt & D.L. Queiroz) #23(6) (NHMB; 70% ethanol; NMB-PSYLL0008548); 1 m$, 5 f£, 1 immature, same but S17.8908, W469247, 660 m, 28‒31.viii.2019, *Xylopia aromatica* (D. Burckhardt & D.L. Queiroz) #347(9) (NHMB; 70% ethanol; NMB-PSYLL0007395); 23 m$, 25 f£, 3 immatures, 2 skins, same but S17.8913/8841, W46.9163/9221, 670–690 m, 29–30.x.2012, *Xylopia aromatica* (D. Burckhardt & D.L. Queiroz) #50(3) (NHMB; slide, 70% ethanol; NMB- PSYLL0007374, NMB-PSYLL0008205); 4 f£, same but S17.88034, W46.92282, 690 m, 26.xii.2014 (D.L. Queiroz) #662 (NHMB; 70% ethanol; NMB-PSYLL0006023); 2 m$, 2 f£, same but S28.1420, W49.6353, 570 m, 9.vi.2018 (D.L. Queiroz) #9.vi.2018(4) (NHMB; 70% ethanol; NMB-PSYLL0007401); 2 m$, 10 f£, same but, Fazenda Bocaina, S17.8930, W46.9111, 640 m, 12.ix.2014, *Xylopia aromatica* (D. Burckhardt & D.L. Queiroz) #146(2) (NHMB; dry, 70% ethanol; NMB-PSYLL0007387, NMB-PSYLL0008574–NMB-PSYLL0008576); 1 f£, Vazante, Votorantim Florestal, Fazenda Rio Escuro, S17.61941, W46.71853, 570 m, 16.iii.2015 (D.L. Queiroz) #685(0) (NHMB; slide; NMB- PSYLL0007427 [voucher 21-2]); 1 m$, 2 f£, same but S17.62858, W46.70022, 540 m, 16.iii.2015 (D.L. Queiroz) #686 (NHMB; 70% ethanol; NMB-PSYLL0006074).—**Pará:** 1 f£, Belém, Jardim Botânico Bosque Rodrigues Alves, S1.4300, W48.4533, 30 m, 14.iv.2017 (D. Burckhardt & D.L. Queiroz) #103(0) (NHMB; 70% ethanol; NMB-PSYLL0008484).— **Roraima:** 40 m$, 20 f£, 3 immatures, 1 skin, Boa Vista, Embrapa campus, N2.7550, W60.7300, 80 m, 31.iii.2015, *Xylopia aromatica* (D. Burckhardt & D.L. Queiroz) #151(1) (NHMB; 70% ethanol; NMB-PSYLL0007368); 9 m$, 10 f£, same but N2.7550, W60.7300, 80 m, 31.iii.2015, *Xylopia aromatica* (D. Burckhardt & D.L. Queiroz) #151(1) (BMNH, MHNG, MMBC, NHMB, UFPR, USNM; dry; NMB-PSYLL0007369–NMB-PSYLL0007373); 5 m$, 11 f£, 1 immature, Boa Vista, 41 km E Boa Vista, Campo Experimental Água Boa, N2.6716, W60.8400, 80 m, 2.iv.2015, *Guatteria* sp. (D. Burckhardt & D.L. Queiroz) #154(4) (NHMB; slide, 70% ethanol; NMB-PSYLL0007389, NMB- PSYLL0007418, NMB-PSYLL0007419 [voucher 41-1], NMB-PSYLL0007420 [voucher 7-5]); 29 m$, 14 f£, 1 immature, Boa Vista, Embrapa campus, N2.7550, W60.7300, 80 m, 23.iv.2015, *Xylopia aromatica* (D. Burckhardt & D.L. Queiroz) #169(1) (MMBC, NHMB; dry, slide, 70% ethanol; NMB-PSYLL0007390, NMB-PSYLL0007413, NMB- PSYLL0007421 [LSMelxyl-7], NMB-PSYLL0007422 [voucher 7-2], NMB-PSYLL0007423 [voucher 7-4], NMB-PSYLL0007424 [voucher 7], NMB-PSYLL0007425 [voucher 7]).— **São Paulo:** 2 m$, Matão, Fazenda Marchesan, 23.x.2007, short suction trap, citrus grove (P. Yamamoto) (NHMB; slide, 70% ethanol; [voucher 48-1]); 1 m$, same but 3.viii.2006, short suction trap, citrus grove (P. Yamamoto) (FSCA; 70% ethanol).

**Description. *Adult.*** Coloration as in *M. amazonica* sp. nov.

Structure. Forewing (Fig. 16B) ovoid, broadest in apical third, broadly and unevenly rounded apically; wing apex in the middle of cell r_2_ margin; C+Sc curved in distal third; pterostigma in the apical third broader than r_1_ cell, strongly convex; Rs unevenly convex, mostly in apical third; M longer than M_1+2_ and M_3+4_; Cu_1a_ moderately convex, ending at M fork; cell cu_1_ short and wide; surface spinules present in all cells, very dense, leaving at most very narrow spinule-free stripes along the veins, forming hexagons of a double row of spinules, spinules sparse in basal half of cell c+sc. Hindwing with 8–9 ungrouped or 4–5 + 4 grouped costal setae. Metatibia bearing 6–7 grouped apical spurs, arranged as 3–4 + 3–4, anteriorly separated by 4–5 bristles.

Terminalia (Fig. 25N–S). Male. Proctiger evenly produced posteriorly; densely covered with long setae in apical two thirds. Subgenital plate irregularly ovoid; dorsal margin relatively straight, curved proximally and distally, posterior margin weakly convex; with dense, long setae. Paramere, in lateral view, irregularly lanceolate, posteriorly with a small lobe in basal third; paramere apex, in lateral view, blunt, directed anteriad, in dorsal view, forming short digitiform process, directed upwards, slightly inwards and slightly anteriad, with a small tooth; outer and inner faces with dense, moderately long setae; posterior margin with long setae. Proximal segment of aedeagus with apical part strongly subdivided. Distal segment of aedeagus almost straight in basal half, with weakly sinuate dorsal margin; ventral process situated in the middle of the segment, spatulate, in lateral view, broadest medially and with narrow, claw-like apex slightly curved dorsad; in dorsal view, ventral process broad (distinctly broader than apical dilation) and subtrapezoidal basally, with a long and narrow, apically truncate median lobe arising from the base ventrally; apical dilation, in lateral view, relatively narrow, with subparallel margins, weakly widening towards rounded apex, in dorsal view, apical dilation elongate, widening towards apex in apical half, apex evenly rounded; sclerotised end tube long and almost straight. – Female terminalia cuneate; densely covered with setae. Dorsal margin of proctiger, in lateral view, concave medially, apex subacute, in dorsal view, apex pointed; circumanal ring, in dorsal view, distinctly cruciform. Subgenital plate, in lateral view, gradually narrowing to apex in apical half, pointed apically; in ventral view, apex blunt.

***Fifth instar immature.*** Coloration. Light; cephalothoracic sclerite, antenna, wing pads, legs and caudal plate brown.

Structure. Eye with one short, simple ocular seta dorsally. Antennal segments with following numbers of pointed sectasetae: 1(0), 2(1), 3(0), 4(2), 5(0), 6(2), 7(1), 8(1), 9(0), 10(0). Forewing pad with 5–7 marginal pointed sectasetae, lacking sectasetae dorsally; hindwing pad with two marginal and one dorsal pointed sectasetae. Tarsal arolium broadly fan-shaped apically, 1.5 times as long as claws. Abdomen with four lateral pointed sectasetae on either side anterior to caudal plate. Caudal plate with anterior margin relatively distant from anterior margin of extra pore fields; with 3 + 3 pointed sectasetae on either side laterally, and three pointed sectasetae subapically, on either side of circumanal ring dorsally. Extra pore fields forming continuous outer and inner bands, consisting of relatively small oval and rounded patches; outer band long medially, end pointing outwards. Circumanal ring small.

**Host plants and biology.** *Xylopia aromatica* (Lam.) Mart., *X. nitida* Dunal, *Guatteria* sp. (Annonaceae). Immatures were found in galls induced by the fungus *Aecidium xylopiae* Henn. (Fungi, Pucciniales) on *Xylopia aromatica* where they secrete white flocculent wax (Fig. 40C, D).

**Distribution.** Brazil (AM, BA, GO, MG, MS, MT, PA, RR, SP).

**Derivation of name.** Named after its host, *Xylopia*.

**Comments.** *Melanastera xylopiae* sp. nov. differs from the other species of the *amazonica*-group as indicated in the key. In the structure of the male and female terminalia, *M. xylopiae* is also similar to *M. olgae* sp. nov., but differs in the dotted (versus uniformly coloured) forewing.

### 20 Melanastera francisi sp. nov

(Figs 9J, 16C, 4)

**Type material. Holotype m$: Brazil: RORAIMA:** Mucajaí, Campo experimental Serra da Prata, Embrapa, N2.3983, W60.9800, 90 m, 16–17.iv.2015, *Xylopia aromatica* (D. Burckhardt & D.L. Queiroz) #163(2) (UFPR; dry).

**Paratypes. Brazil: MATO GROSSO:** 1 m$, 1 f£, Sorriso, Fazenda Mazzardo II, S12.4645, W55.7840, 390 m, 25.vii.2014 (T. Mazzardo) #P001 (NHMB; dry; NMB-PSYLL00003007); 1 m$, same but S12.4244, W55.8503, 390 m, 26.vii.2014 #P020 (NHMB; dry; NMB- PSYLL00003009); 2 m$, 8 f£, same but S12.4217, W55.8432, 390 m, 15.viii.2013 (NHMB; dry, 70% ethanol; NMB-PSYLL0007874, NMB-PSYLL0003232, NMB-PSYLL0003233).—**Roraima:** 18 m$, 16 f£, Boa Vista, 28 km E Boa Vista, Campo Experimental Água Boa, N2.6717, W60.8400, 90 m, 2.iv.2015, *Xylopia aromatica* (D. Burckhardt & D.L. Queiroz) #154(1) (NHMB; 70% ethanol; NMB-PSYLL0007875); 7 m$, 1 f£, 15 immatures, 3 skins, same but *Guatteria* sp. (D. Burckhardt & D.L. Queiroz) #154(4) (NHMB; 70% ethanol; NMB-PSYLL0007876, NMB-PSYLL0008317); 12 m$, 17 f£, Boa Vista, Embrapa campus, N2.7550, W60.7300, 80–100 m, 20.iv.2015, *Xylopia aromatica* (D. Burckhardt & D.L. Queiroz) #169(1) (NHMB; 70% ethanol; NMB-PSYLL0008495); 7 m$, 14 f£, 7 immatures, same as holotype but (NHMB, UFPR; dry, slide, 70% ethanol; NMB-PSYLL0007877, NMB- PSYLL0008258, NMB-PSYLL0008259, NMB-PSYLL0008260, NMB-PSYLL0008261, NMB-PSYLL0008262, NMB-PSYLL0007888, NMB-PSYLL0007889 [LSMelmac-12], NMB-PSYLL0007890 [LSMelmac-12]); 21 m$, 21 f£, 6 immatures, same but km 442–440 along BR-174 S Mucujaí, N2.3583/3850, W60.9133/9167, 100 m, 17.iv.2015 (D. Burckhardt & D.L. Queiroz) #165(1) (MMBC, NHMB; dry, slide, 70% ethanol; NMB-PSYLL0007878, NMB-PSYLL0008625, NMB-PSYLL0008626, NMB-PSYLL0007879 [LSMelmac-12], NMB-PSYLL0007880 [LSMelmac-12], NMB-PSYLL0007881 [LSMelmac-12], NMB- PSYLL0007882 [LSMelmac-12], NMB-PSYLL0007883, NMB-PSYLL0007886, NMB- PSYLL0007887).

**Description. *Adult.*** Coloration as in *M. cacantis* sp. nov.

Structure. Forewing (Fig. 16C) ovoid, widest in the middle, broadly and unevenly rounded apically; wing apex situated in cell r_2_ near apex of M_1+2_; C+Sc curved in distal third; pterostigma wider than cell r_1_ in apical third, strongly convex; Rs irregularly convex, strongly curved in apical third; M longer than M_1+2_ and M_3+4_; Cu_1a_ irregularly convex, ending at M fork; cell cu_1_ short and wide; surface spinules present in all cells, very dense, leaving no spinule-free stripes along the veins. Hindwing with 9–12 ungrouped costal setae. Metatibia bearing 5–6 apical spurs, arranged as 2–3 + 3–4, anteriorly separated by 4 bristles. Terminalia (Fig. 25T–Y). Male. Proctiger long, moderately produced in basal third posteriorly; densely covered with long setae in apical two thirds. Subgenital plate irregularly ovoid; dorsal margin weakly curved, posterior margin almost straight; with densely spaced long setae. Paramere, in lateral view, irregularly lamellar, with irregular posterior lobe in basal half; paramere apex, in lateral view, subacute, directed anteriad, in dorsal view, forming short digitiform process, pointed, directed inwards and slightly anteriad, with a sclerotised tooth; both outer and inner faces with dense, long setae; posterior margin with very long setae. Proximal segment of aedeagus with apical part sclerotised and strongly subdivided. Distal segment of aedeagus weakly sinuate in basal half; ventral process situated slightly proximad of the middle of the segment, in lateral view, spatulate, with a narrow base, broadest slightly posterior of the middle and with narrow, claw-like apex strongly curved dorsad; in ventral view, ventral process broad, distinctly broader than apical dilation, and subtrapezoidal basally, with a short and narrow, apically rounded median lobe arising from the base ventrally; apical dilation, in lateral view, long, narrow, with subparallel margins and rounded apex, base bearing small membranous extension at dorsal margin, in dorsal view, apical dilation slightly constricted submedially or subparallel-sided between ventral process and apex; sclerotised end tube long and almost straight. – Female terminalia cuneate; densely covered with setae.

Dorsal margin of proctiger, in lateral view, concave distal to circumanal ring, apex narrow and subacute, in dorsal view, apex pointed; circumanal ring, in dorsal view, cruciform.

Subgenital plate, in lateral view, gradually narrowing to pointed apex; in ventral view, apex blunt.

***Fifth instar immature.*** Coloration. Yellow; antenna brownish yellow to pale brown, segments 6–8 with dark brown apices, segments 9–10 entirely dark brown; cephalothoracic sclerite, wing pads, legs and caudal plate pale brown.

Structure. Eye with one short, simple ocular seta dorsally. Antennal segments with following numbers of pointed sectasetae: 1(0), 2(1), 3(0), 4(2), 5(0), 6(2), 7(1), 8(1), 9(0), 10(0). Forewing pad with five marginal pointed sectasetae; hindwing pad with two marginal pointed sectasetae; both pads lacking sectasetae or larger lanceolate setae dorsally. Tarsal arolium broadly fan-shaped apically, 1.5 times as long as claws. Abdomen with one lateral pointed sectaseta on either side anterior to caudal plate. Caudal plate with anterior margin remote from anterior margin of extra pore fields; with 2 + 3 pointed sectasetae on either side laterally, and three pointed sectasetae near circumanal ring on either side dorsally. Extra pore fields forming continuous outer and inner bands, consisting of relatively small oval and round patches; outer band relatively long medially, end pointing outwards. Circumanal ring small.

**Host plant.** *Guatteria* sp., *Xylopia aromatica* (Lam.) Mart. (Annonaceae).

**Distribution.** Brazil (MT, RR).

**Derivation of name.** Dedicated to Francis Halley Queiroz Sant’Anna, son of D.L. Queiroz.

**Comments.** *Melanastera francisi* sp. nov. differs from the other species of the *amazonica*- group as indicated in the keys. It also shares a similar forewing venation and similar type of terminalia with *M. olgae* sp. nov. but differs from the latter in the dotted forewing (versus uniformly coloured).

### 21 Melanastera roraima sp. nov

(Figs 3C, D, 10A, 16D, 26A–F)

**Type material. Holotype m$: Brazil: RORAIMA:** Amajari, Tepequém, Cachoeiras do Paiva, N3.7566, W61.7183, 610 m, 4.iv.2015, *Guatteria* sp. (D. Burckhardt & D.L. Queiroz) #157(2) (UFPR; dry).

**Paratypes. Brazil: RORAIMA:** 2 m$, 3 f£, same as holotype but (NHMB, UFPR; dry, slide; NMB-PSYLL0005575, NMB-PSYLL0008592, NMB-PSYLL0008593, NMB-PSYLL0008078 [LSMelror-36]); 1 f£, same but outside village, N3.7566, W61.7183, 720 m, 3.iv.2015 (D. Burckhardt & D.L. Queiroz) #155 (NHMB; slide; NMB-PSYLL0008079 [voucher 45-1]); 2 m$, 6 f£, same but along river, N3.7566, W61.7183, 720 m, 3.iv.2015, *Guatteria* sp. (D. Burckhardt & D.L. Queiroz) #156(1) (MMBC, NHMB; slide, 70% ethanol; NMB-PSYLL0008055, NMB-PSYLL0008076 [LSMelror-36], NMB-PSYLL0008077 [LSMelror-36]).

**Description. *Adult.*** Coloration similar to *M. cacantis* sp. nov.

Structure. Forewing (Fig. 16D) oval, widest in the middle, broadly and unevenly rounded apically, wing apex situated in the middle of the margin of cell r_2_; C+Sc curved in in apical third; pterostigma wider in the middle than r_1_ cell, distinctly convex in apical two thirds; Rs slightly convex along most of its length, strongly curved; M longer than M_1+2_ and M_3+4_; Cu_1a_ weakly curved, ending at M fork; cell cu_1_ short and wide; surface spinules present in all cells, very dense, hexagons hardly distinct, leaving very narrow spinule-free stripes along the veins. Hindwing with 7–9 ungrouped or 5 + 3 grouped costal setae. Metatibia bearing 6–7 grouped apical spurs, arranged as 3 + 3–4, anteriorly separated by 3–4 bristles.

Terminalia (Fig. 26A–F). Male. Proctiger posteriorly weakly produced in basal third; densely covered with long setae in apical two thirds. Subgenital plate subtriangular; dorsal margin slightly sinuate, posterior margin almost straight; with dense, moderately long setae. Paramere, in lateral view, irregularly lanceolate, posterior margin bearing conspicuous lobe in basal half; paramere apex, in lateral view, subacute, directed slightly anteriad, in dorsal view, apex forming short narrow process, acute, directed inwards and slightly anteriad, with sclerotised tooth; outer and inner faces with dense, moderately long setae; posterior margin with long setae. Proximal segment of aedeagus with apical part strongly sclerotised and subdivided. Distal segment of aedeagus strongly sinuate in basal third; ventral process situated slightly proximal of the middle of the segment, spatulate, in lateral view, with a narrow base, broadest slightly distal of the middle, with narrow, claw-like apex strongly curved dorsad; in dorsal view, ventral process broader than apical dilation, subtrapezoidal basally, with short and narrow, apically truncate median lobe arising from the base ventrally; apical dilation, in lateral view, long, moderately thick, slightly inflated apically, with rounded apex, base bearing a small membranous extension at dorsal margin, in dorsal view, distinctly constricted submedially between ventral process and apex; sclerotised end tube long and almost straight. – Female terminalia cuneate; densely covered with setae. Dorsal margin of proctiger, in lateral view, slightly concave posterior to the circumanal ring, apex pointed and slightly upturned; in dorsal view, apex pointed; circumanal ring, in dorsal view, cruciform.

Subgenital plate, in lateral and ventral views, pointed apically.

*Fifth instar immature.* Unknown.

**Host plant.** Adults were collected on *Guatteria* sp. (Annonaceae) which is a probable host.

**Distribution.** Brazil (RR).

**Derivation of name.** Named after Roraima, the State in Brazil, where the species was collected.

**Comments.** *Melanastera roraima* sp. nov. differs from the other members of the *M. amazonica*-group as indicated in the keys. From *M. olgae* sp. nov. with which it shares similarities in the forewing venation and terminalia, it differs in the dotted forewing (versus uniformly coloured).

### 22 Melanastera tubuligera sp. nov

(Figs 2C, 10B, 16E, 26G–L)

**Type material. Holotype m$: Brazil: MATO GROSSO:** Tabaporã, S11.5816, W55.7666, 370 m, 8.xi.2012, *Xylopia aromatica* (D. Burckhardt & D.L. Queiroz) #64(2) (UFPR; dry).

**Paratypes. Brazil: MATO GROSSO:** 3 m$, 1 f£, Juína, S11.5631, W58.9516, 340 m, 17.vi.2014 (L.A. Pezzini) #319 (NHMB; 70% ethanol; NMB-PSYLL0005637); 4 m$, 1 f£, Lucas do Rio Verde, S13.1171, W55.8978, 390 m, 21.iv.2015 (L.A. Pezzini) #502 (NHMB; slide, 70% ethanol; NMB-PSYLL0005754, NMB-PSYLL0008225); 1 m$, 1 skin, Nova Mutum, S13.7045, W56.0715, 420 m, 12.vi.2016, *Xylopia* sp. (L.A. Pezzini) #807 (NHMB; 70% ethanol; NMB-PSYLL0008120); 1 m$, same but S13.7898, W56.2901, 390 m, 13.vi.2016 #845 (NHMB; 70% ethanol; NMB-PSYLL0005808); 2 m$, 3 f£, Sinop, S11.3783, W55.3783, 350 m, 5.iii.2014 (L.A. Pezzini & A.M.B. Sedano) #286 (MMBC, NHMB; dry, slide; NMB-PSYLL0005675, NMB-PSYLL0008235 [voucher 75-1]); 2 m$, 1 f£, Sinop, Embrapa campus, S11.8949, W55.6555, 340 m, 29.viii.2013 (D.L. Queiroz) #568 (NHMB; 70% ethanol; NMB-PSYLL0008114); 3 m$, 1 f£, Sorriso, S12.4645, W55.7839, 340 m, 25.vii.2014 (L.A. Pezzini) #372 (NHMB; 70% ethanol; NMB-PSYLL0005646); 2 m$, 2 f£, 1 immature, same S12.4536, W55.8071, 370 m, 25.vii.2014 #379 (NHMB; 70% ethanol; NMB- PSYLL0005690); 1 m$, 2 f£, 2 immatures, same but S12.4261, W55.7943, 340 m, 27.ii.2015 #485 (NHMB; 70% ethanol; NMB-PSYLL0008121); 5 m$, 7 f£, 10 immatures, 4 skins, same but Comunidade Navegantes, S12.4054, W55.8238, 380 m, 26.ii.2015 (D.L. Queiroz) #676(2) (NHMB; slide, 70% ethanol; NMB-PSYLL0008115, NMB-PSYLL0008219, NMB- PSYLL0008257, NMB-PSYLL0005462, NMB-PSYLL0008224 [voucher 78-1], NMB- PSYLL0008221 [LSMeltub-25], NMB-PSYLL0008220 [LSMeltub-25]); 11 m$, 12 f£, 1 skin, same but Fazenda Mazzardo II, S11.8714, W55.5957, 390 m, 26.ii.2015, *Guatteria schomburgkiana* (D.L. Queiroz) #678(3) (NHMB; slide, 70% ethanol; NMB-PSYLL0008284, NMB-PSYLL0008116, NMB-PSYLL0008227 [voucher 25-3], NMB-PSYLL0008222 [LSMeltub-25]); 6 m$, 5 f£, same but S11.8714, W55.5957, 390 m, 15.viii.2013 (T. Mazzardo) (NHMB; slide, 70% ethanol; NMB-PSYLL0008122, NMB-PSYLL0008230 [voucher 25-3], NMB-PSYLL0008231 [voucher 25-4], NMB-PSYLL0008234); 2 m$, 1 f£, Tabaporã, Fazenda Crestani, S11.3133/3366, W55.9616/9750, 330–380 m, 6–8.xi.2012, *Xylopia aromatica* (D. Burckhardt & D.L. Queiroz) #62(9) (NHMB; 70% slide, ethanol; NMB-PSYLL0008299, NMB-PSYLL0008233 [voucher 115-3], NMB-PSYLL0008228 [voucher 115-4]); 3 m$, 4 f£, same as holotype but (NHMB; slide, 70% ethanol; NMB- PSYLL0008113, NMB-PSYLL0008223, NMB-PSYLL0008226, NMB-PSYLL0008229 [voucher 115-1]); 1 m$, Terra Nova do Norte, S10.4695, W54.9913, 310 m, 28.v.2016 (L.A. Pezzini) #717 (NHMB; dry; NMB-PSYLL0005777).—**Minas Gerais:** 1 m$, Paula Cândido, S20.8080, W42.9839, 780 m, 22.viii.2013, coffee plantation with *Inga* (D.L. Queiroz) #565 (NHMB; slide; NMB-PSYLL0008232).

**Description. *Adult.*** Coloration similar to *M. francisi* sp. nov. but darker.

Structure. Forewing (Fig. 16E) ovoid, widest in the middle, broadly and unevenly rounded apically, wing apex situated at the margin of cell r_2_ near apex of M_1+2_; C+Sc curved in distal third; pterostigma in the middle wider than cell r_1_, strongly convex; Rs irregularly convex, strongly curved in apical third; M longer than M_1+2_ and M_3+4_; Cu_1a_ moderately convex, ending at M fork; cell cu_1_ short and wide; surface spinules present in all cells, very dense, leaving no spinule-free stripes along the veins, rarely spinules forming hexagons of a double row of spinules. Hindwing with 5 + 3–4 grouped costal setae. Metatibia bearing 6–7 grouped apical spurs, arranged as 3 + 3–4, anteriorly separated by ca. 5 bristles.

Terminalia (Fig. 26G–L). Male. Proctiger weakly produced produced posteriorly; densely covered with very long setae in apical two thirds. Subgenital plate irregularly ovoid; dorsal margin indented in proximal third, posterior margin almost straight; with dense, long setae. Paramere, in lateral view, irregularly lanceolate with flattened lobe in basal half of posterior margin; paramere apex, in lateral view, subacute, directed slightly anteriad, in dorsal view, apex subacute, directed upwards and slightly inwards, with small sclerotised tooth; outer face with dense, moderately long setae in apical two thirds; inner face with dense, moderately long setae in apical half and dense, long setae submedially; posterior margin with very long setae. Proximal segment of aedeagus with apical part strongly subdivided. Distal segment of aedeagus with dorsal margin distinctly sinuate in basal third; ventral process situated slightly proximal of the middle of segment, spatulate, in lateral view, with narrow base, widest medially and with narrow, claw-like apex slightly curved dorsad; in dorsal view, ventral process broad and subtrapezoidal basally, with a long and narrow, apically truncate median lobe arising from the base ventrally; apical dilation, in lateral view, long, narrow, subparallel- sided, slightly inflated apically, with broadly rounded apex, base bearing a large membranous extension at dorsal margin, in dorsal view, apical dilation weakly constricted or parallel-sided medially; sclerotised end tube very long and almost straight. – Female terminalia cuneate; densely covered with setae. Dorsal margin of proctiger, in lateral view, moderately concave, apex upturned, subacute, in dorsal view, apex blunt; circumanal ring, in dorsal view, cruciform. Subgenital plate, in lateral view, abruptly narrowing to apex in apical half, apex pointed; in ventral view, apex blunt.

***Fifth instar immature.*** Coloration. Entirely yellow; antenna gradually darkening to apex.

Structure. Eye with one short simple ocular seta dorsally. Antennal segments with following numbers of pointed sectasetae: 1(0), 2(1–2), 3(0), 4(2), 5(0), 6(2), 7(1), 8(1), 9(0), 10(0). Forewing pad with 5–6 marginal pointed sectasetae; hindwing pad with 2 marginal pointed sectasetae; both pads lacking sectasetae dorsally. Tarsal arolium broadly fan-shaped apically, 1.5 times as long as claws. Abdomen with 2–3 lateral pointed sectasetae on each side anterior to caudal plate. Caudal plate with anterior margin close to anterior margin of extra pore fields; with five pointed sectasetae on either side laterally, and three pointed sectasetae subapically, on either side of circumanal ring dorsally. Extra pore fields forming continuous outer and inner bands, consisting of small oval patches; outer band long medially, end pointing outwards. Circumanal ring small.

**Host plant.** *Guatteria schomburgkiana* Mart. (Annonaceae); adults were also collected on *Xylopia aromatica* (Lam.) Mart. which is a possible host.

**Distribution.** Brazil (MG, MT).

**Derivation of name.** From the Latin noun *tubulus* = a small tube and the verb *gerere* = to carry, referring to the very long sclerotised end tube of the ductus ejaculatorius.

**Comments.** *Melanastera tubuligera* sp. nov. differs from the members of the *amazonica*- group as indicated in the keys. From *M. olgae* sp. nov. with which it shares similarities in the forewing venation and terminalia, it differs in the dotted forewing (versus uniformly coloured).

### The umbripennis-group

**Description. *Adult*.** Vertex (Fig. 10C–F) trapezoidal, covered with microscopical setae. Thorax weakly to moderately arched, with microscopical setae. Forewing (Fig. 16F–J) conspicuously lighter proximal to nodal line, with irregularly spaced brown dots, oviform, widest in the middle, broadly, evenly rounded apically; wing apex situated in cell r_2_ close to apex of M_1+2_; C+Sc almost straight, weakly curved subapically; pterostigma distinctly expanding towards the middle or apical third, in the middle narrower than ajacent part of cell r_1_; R_1_ irregularly curved, strongest in apical third; M longer than M_1+2_ and M_3+4_; Cu_1a_ weakly or moderately convex, ending at or distal of M fork; cell cu_1_ long and relatively wide. Ventral process of the distal aedeagal segment present, relatively broad, curved ventrad (Fig. 26O, P, U, V) or relatively narrow, straight, oriented obliquely apicad (Fig. 27C, D, I, J).

***Immature.*** Antenna 10-segmented.

**Comments.** The group includes four species in Brazil. Hosts are known for three of them and belong to Annonaceae.

### 23 Melanastera umbripennis sp. nov

(Figs 10C, 16F, G, 26M–R)

**Type material. Holotype m$: Brazil: AMAZONAS:** 1 m$, Manaus, Adolpho Ducke Forest Reserve, S2.9645, W59.9202, 100 m, 18.vi.1996, fogging of *Eschweilera rodriguesiana* (J.C.H. Guerrero, P.O.I. de Lima & G.P.L. Mesquita) F3-30(8) (INPA; dry).

**Paratypes: Brazil: AMAZONAS:** 3 m$, 1 f£, same as holotype but F3-30(3) (MMBC; dry, slide; [LSMelumb]); 1 m$, 2 f£, same but (MMBC, NHMB; dry, slide; NMB- PSYLL0008162); 1 m$, 2 f£, same but F3-30(5) (BMNH, INPA; dry, slide; [LSMelumb]); 1 m$, 1 f£, same but (NHMB; slide; NMB-PSYLL0008163 [LSMelumb], NMB- PSYLL0008164 [LSMelumb]); same but F3-30(9) (INPA; dry); 1 m$, same but fogging of *Micropholis guyanensis*, F3-50A(10) (BMNH; dry); 2 f£, same but 8.iii.1996, fogging of *Ecclinusa guianensis* F3-150(3) (INPA; dry).

**Description. *Adult.*** Coloration. Body brown. Antenna pale yellow, rarely segments 3–10 with brown apices. Mesoscutum with four broad light brown longitudinal stripes; thorax laterally ochreous. Forewing (Fig. 16F) mostly clear in basal third, dark brown in apical two thirds; sometimes with dark brown dots (Fig. 16G); apices of pterostigma and Rs dark brown, sometimes also apices of M_1+2_, M_3+4_, Cu_1a_ and Cu_1b_. Legs pale yellow, pro- and mesocoxae and basal two thirds of femora dark brown. Younger specimens with much less expanded dark colour.

Structure. Head, in lateral view, inclined at < 45° from longitudinal body axis. Vertex (Fig. 10C) with indistinct microsculpture. Forewing (Fig. 16F, G) with Rs strongly curved apically; surface spinules present in all cells, forming hexagons of a single row of spinules, spinules leaving broad spinule-free stripes along the veins, absent in basal two thirds of cell c+sc. Hindwing with 6 ungrouped costal setae. Metatibia bearing 5–7 apical spurs, arranged as 2–3 + 3–4, anteriorly separated by 4 bristles.

Terminalia (Fig. 26M–R). Male. Proctiger strongly produced in basal half; covered with moderately long setae in apical half. Subgenital plate, in lateral view, subglobular; dorsal margin weakly curved, posterior margin evenly convex; with sparse, long setae. Paramere, in lateral view, irregularly lanceolate, widest medially; paramere apex, in lateral view, blunt, directed slightly anteriad, in dorsal view, apex blunt, directed upwards as well as slightly inwards and anteriad, lacking a sclerotised tooth; outer face with moderately long setae in apical two thirds; inner face with dense, moderately long setae; posterior margin with long setae. Proximal segment of aedeagus with apical part not subdivided. Distal segment of aedeagus with dorsal margin strongly sclerotised and strongly sinuate in basal half; ventral process slightly proximal to the middle, large, in lateral view, thick, distinctly curved ventrad, broadly rounded apically, in dorsal view, ventral process much wider than apical dilation, with broad wing-like lateral lobes on each side and a relatively small button-like median lobe directed slightly ventrad, apical dilation, in lateral view, moderately elongate, subparallel- sided, rounded apically, with a small membranous extension postero-basally, in dorsal view, apical dilation widest in apical third, rounded apically; sclerotised end tube moderately long and weakly sinuate. – Female terminalia cuneate; covered with short setae. Dorsal margin of proctiger, in lateral view, weakly convex distal to circumanal ring, apex upturned, subacute; in dorsal view, apex pointed; circumanal ring, in dorsal view, vaguely cruciform. Subgenital plate, in lateral view, abruptly narrowing in apical half, pointed apically, not upturned; in ventral view, apex blunt.

*Fifth instar immature.* Unknown.

**Host plant.** Adults were fogged from *Eschweilera atropetiolata* S.A.Mori, *E. rodriguesiana* S.A.Mori (Lecythidaceae), *Ecclinusa guianensis* Eyma and *Micropholis guyanensis* (A.DC.) Pierre (Sapotaceae) which are unlikely hosts.

**Distribution.** Brazil (AM).

**Derivation of name.** From the Latin nouns *umbra* = shade, shadow, and *penna* = feather, wing, referring to the dark brown forewing.

### 24 Melanastera nasuta sp. nov

(Figs 10D, 16H, 26S–X)

**Type material. Holotype m$: Brazil: AMAZONAS:** Iranduba, Embrapa, Campo Experimental, Caldeirão, S3.2516/2566, W60.2216/2233, 30–50 m, 14–17.iv.2014, *Annona* sp. (D. Burckhardt & D.L. Queiroz) #125(10) (UFPR; dry).

**Paratypes. Brazil: AMAZONAS:** 3 m$, 3 f£, 8 immatures, same as holotype but (NHMB; slide, 70% ethanol; NMB-PSYLL0007969, NMB-PSYLL0007961 [LSMelinas-15], NMB- PSYLL0007962 [LSMelnas-15], NMB-PSYLL0007963, NMB-PSYLL0007964, NMB- PSYLL0007965).—**MATO GROSSO:** 10 m$, 11 f£, Colíder, S10.5839, W55.4862, 290 m, 29.v.2016, *Guatteria* sp. (L.A. Pezzini) #769 (NHMB; dry, slide, 70% ethanol; NMB- PSYLL0005768, NMB-PSYLL0007928, NMB-PSYLL0007948, NMB-PSYLL0007966); 1 m$, Lucas do Rio Verde, S13.2490, W56.0275, 400 m, 12.vi.2016 (L.A. Pezzini) #794 (NHMB; dry; NMB-PSYLL0005771); 1 f£, Nova Mutum, S13.5930, W56.0611, 430 m, 12.vi.2016 (L.A. Pezzini) #801 (NHMB; dry; NMB-PSYLL0005790); 4 m$, 8 f£, Terra Nova do Norte, S10.4369, W55.1532, 300 m, 28.v.2016, *Guatteria* sp. (L.A. Pezzini) #704 (BMNH, MMBC, NHMB, USNM; dry, 70% ethanol; NMB-PSYLL0007970, NMB-PSYLL0005743).

**Description. *Adult.*** Coloration. Head (Fig. 10D) and clypeus dark brown; genae and vertex laterally yellow. Antenna pale yellow, segments 3–8 with brown apices, segments 9–10 entirely brown. Thorax brown to dark brown; mesoscutum with four broad and one median narrow dark brown longitudinal stripes. Forewing (Fig. 16H) pale yellow to pale brown with distinct, dense brown dots scattered over entire surface, becoming darker, denser and often confluent towards wing apex; apical third of cell cu_2_ whitish lacking dark dots; cell a_2_ brown; base and apex of pterostigma and apices of Rs, M_1+2_, M_3+4_, Cu_1a_ and Cu_1b_ dark brown. Legs pale yellow, base of femora dark brown and distal tarsal segments brownish. Abdomen with dark brown sclerites. Terminalia brown; female terminalia with pale brown apex.

Structure. Head, in lateral view, inclined at > 45° from longitudinal body axis. Vertex with distinct imbricate microsculpture. Forewing (Fig. 16H) with Rs obliquely curved to fore margin apically; surface spinules present in all cells, leaving narrow spinule-free stripes along veins, forming hexagons of a single row of spinules; spinules sparse at apex of cell cu_2_ and absent from basal half of cell c+sc. Hindwing with 5–6 ungrouped costal setae. Metatibia bearing 6 apical spurs, arranged as 3 + 3, anteriorly separated by 3 bristles.

Terminalia (Fig. 26S–X). Male. Proctiger weakly produced posteriorly; densely covered with short setae in apical two thirds. Subgenital plate, in lateral view, subtriangular; dorsal margin rounded in proximal half, straight in apical half, posterior margin irregularly convex; with moderately long setae. Paramere, in lateral view, lamellar, with abruptly narrowed digitiform, subacute apical process which is weakly curved anteriad; in dorsal view, apex subacute, directed obliquely anteriad and upwards; outer face with sparse, short setae in apical two thirds; inner face with dense, short setae in apical half and dense, long setae arranged in an oblique row in basal half. Proximal segment of aedeagus with apical part strongly subdivided. Distal segment of aedeagus almost straight in basal half, with dorsal margin weakly sinuate; ventral process in distal third of the segment; in lateral view, ventral process large, nose-shaped, broadly fused with apical dilation poximally, bearing a small button-like lobe on either side apically; in dorsal view, ventral process triangular, broader than apical dilation, with small button-like lobes only partly covered by apical dilation; apical dilation, in lateral view, almost half as long as segment, with rounded apex and membranous basal lobe; in dorsal view, apical dilation oblong, with rounded apex; sclerotised end tube short and almost straight. – Female terminalia cuneate; densely covered with setae. Dorsal margin of proctiger, in lateral view, weakly sinuate distal to circumanal ring, apex blunt; in dorsal view, apex blunt; circumanal ring, in dorsal view, distinctly cruciform. Subgenital plate, in lateral view, pointed apically; apex, in ventral view, blunt.

***Fifth instar immature.*** Coloration. Pale yellow, apical two antennal segments pale brown; cephalothoracic sclerite, wing pads, legs and caudal plate yellow to orange.

Structure. Eye with one short, simple ocular seta dorsally. Antennal segments with following numbers of pointed sectasetae: 1(0), 2(1), 3(0), 4(2), 5(0), 6(2), 7(1), 8(1), 9(0), 10(0). Forewing pad with 5–6 marginal pointed sectasetae; hindwing pad with two marginal pointed sectasetae; both wing pads lacking sectasetae or larger lanceolate setae dorsally. Tarsal arolium broadly triangular apically, slightly longer than claws. Abdomen with three lateral pointed sectasetae on either side anterior to caudal plate. Caudal plate with anterior margin close to anterior margin of extra pore fields; with 3–4 (1+2 or 2+2) pointed sectasetae on either side laterally, and three pointed sectasetae subapically, on either side of circumanal ring dorsally. Extra pore fields forming continuous outer and inner bands, consisting of relatively small rounded patches; outer band long medially, end pointing outwards. Circumanal ring small.

**Host plant.** *Annona* sp. (Annonaceae); adults were also collected on *Guatteria* sp. which is a possible host.

**Distribution.** Brazil (AM, MT).

**Derivation of name.** From the Latin adjective *nasutus* = big-nosed, referring to the nose- shaped apical dilation of the aedeagus.

**Comments.** *Melanastera nasuta* sp. nov. differs from the other species of the genus in the unique shape of the distal segment of aedeagus.

### 25 Melanastera australis sp. nov

(Figs 10E, 16I, 27A–F)

**Type material. Holotype m$: Brazil: PARANÁ:** Antonina, Usina Parigot de Souza, S25.2518, W48.7752, 480 m, 19.vii.2017 (D. Burckhardt & D.L. Queiroz) #249 (UFPR; dry).

**Paratypes. Brazil: PARANÁ:** 1 m$, 3 immatures, same as holotype but S25.2557, W48.7787, 720 m, 19.vii.2017, *Guatteria australis* (D. Burckhardt & D.L. Queiroz) #250(3) (MMBC, NHMB; slide, 70% ethanol; NMB-PSYLL0007742, NMB-PSYLL0007782, NMB- PSYLL0007783 [LSMelaus-31]); 1 f£, same as but S25.2500, W48.7700, 300 m, 20.vii.2017, *Guatteria australis*, Atlantic forest (I. Malenovský) (MMBC; slide).—**Santa Catarina:** 2 m$, 1 f£, Timbó, SC-416, 4 km from Timbó, Pousada Paraíso da Pesca, S26.7716/7800, W49.2183/2250, 270–380 m, 29.iv.2013, *Annona dolabripetala* (D. Burckhardt & D.L. Queiroz) #117(7) (NHMB; 70% ethanol; NMB-PSYLL0008498).

**Description. *Adult.*** Coloration. Head, genae and vertex pale brown; vertex with dark brown pattern; clypeus dark brown. Antenna orange, apices of segments 4–9 and entire segment 10 brown. Thorax brownish yellow to pale brown; mesopraescutum with dark brown patches along fore margin; mesoscutum with four broad brown longitudinal stripes, margined by rows of dark brown dots, and one or two median rows of dark brown dots. Metanotum brown, sometimes with dark brown pattern. Forewing (Fig. 16I) with brown veins and yellow membrane covered with brown dots, relatively sparse in basal third and along wing margins, very dense and often confluent otherwise; nodal line irregularly whitish. Legs pale yellow to orange, pro- and mesocoxae dark brown, metacoxa pale yellow, basal half of femora pale to dark brown, distal tarsal segment pale brown. Abdomen yellow with dark brown sclerites; terminalia dark brown.

Structure. Head, in lateral view, inclined at > 45° from longitudinal body axis. Vertex (Fig. 10E) with imbricate microsculpture and short setae. Forewing (Fig. 16I) with Rs weakly curved to fore margin apically; surface spinules present in all cells, leaving narrow to broad spinule-free stripes along the veins, forming hexagons of a single row of spinules; spinules very sparse in cell cu_2_ and in basal half of cells r_1_, r_2_ and m_2_, and absent in basal part of cell c+sc. Hindwing with 4 + 3 grouped or 7 ungrouped costal setae. Metatibia bearing 5–7 grouped apical spurs, arranged as 2–3 + 3–4, anteriorly separated by 4–5 bristles.

Terminalia (Fig. 27A–F). Male. Proctiger weakly produced posteriorly; densely covered with long setae in apical half. Subgenital plate, in lateral view, triangular; dorsal margin straight basally, curved in apical quarter; posterior margin weakly curved; with moderately long setae. Paramere lamellar, irregularly narrowing to apex; apex, in lateral view, blunt, directed upwards, in dorsal view, subacute, directed upwards and slightly inwards, lacking distinct sclerotised tooth; outer face with dense, moderately long setae in apical two thirds; inner face with dense, moderately long setae in apical two thirds and longer setae at base; posterior margin with long setae. Proximal segment of aedeagus with apical part weakly subdivided. Distal segment of aedeagus with weakly sinuate dorsal margin in basal two thirds; ventral process situated in apical third of the segment, in lateral view, short, broad, simple, truncate apically, lacking lateral lobes; in dorsal view, ventral process tongue-shaped with rounded apex, slightly broader than apical dilation; apical dilation short, in lateral view, subparallel-sided, with almost straight dorsal margin, apex irregularly rounded, base bearing small membranous angular lobe at dorsal margin; sclerotised end tube short and weakly sinuate. – Female terminalia cuneate; densely covered with setae. Dorsal margin of proctiger, in lateral view, slightly concave in apical half, apex straight, pointed; in dorsal view, apex blunt; circumanal ring, in dorsal view, distinctly cruciform. Subgenital plate, in lateral view, strongly narrowing to apex in apical half, pointed apically; in ventral view, apex blunt.

***Fifth instar immature.*** Coloration. Yellow; antenna pale yellow to pale brown, three apical segments entirely dark brown; cephalothoracic sclerite, wing pads, legs and caudal plate brown.

Structure. Eye with one short, simple ocular seta dorsally. Antenna with following numbers of pointed sectasetae per segment: 1(0), 2(1), 3(0), 4(2), 5(0), 6(2), 7(1), 8(1), 9(0), 10(0). Forewing pad with five large marginal pointed sectasetae and 2–3 smaller sectasetae or lanceolate setae dorsally; hindwing pad with two large marginal pointed sectasetae and one smaller sectaseta dorsally. Tarsal arolium broadly triangular apically, slightly longer than claws. Abdomen with 3–4 lateral pointed sectasetae on each side anterior to caudal plate. Anterior margin of caudal plate relatively distant from anterior margin of extra pore fields; with five (2+3) pointed sectasetae on either side laterally, and 3–4 pointed sectasetae subapically, on either side of circumanal ring dorsally. Extra pore fields forming continuous outer and inner bands, consisting of moderate oval patches; outer band relatively short medially, end pointing outwards. Circumanal ring small.

**Host plant.** *Guatteria australis* A.St.-Hil. (Annonaceae); adults were also collected on

*Annona dolabripetala* Raddi which is a possible host.

**Distribution.** Brazil (PR, SC).

**Derivation of name.** Named after its host, *G. australis*.

**Comments.** *Melanastera australis* sp. nov. resembles *M. dimorpha* sp. nov., *M. guatteriae* sp. nov. and *M. obscura* sp. nov. in the paramere shape, and *M. dimorpha*, *M. nasuta* sp. nov. and *M. obscura* in the forewing very light basally, contrasting with a much darker pattern in its apical two thirds. It can be separated from these species as indicated in the keys.

### 26 Melanastera obscura sp. nov

(Figs 10F, 16J, 27G–L)

**Type material. Holotype m$: Brazil: MINAS GERAIS:** Barroso, Mata do Baú, Rio das Mortes, S21.2140, W43.9347, 930 m, 12–17.vi.2010, *Annona* sp., gallery forest along river (D. Burckhardt) #5(13) (UFPR; dry).

**Paratypes. Brazil: MINAS GERAIS:** 6 m$, 7 f£, same as holotype but (BMNH, MMBC, NHMB, UFPR; dry, slide; NMB-PSYLL0008008, NMB-PSYLL0008586, NMB- PSYLL0008590, NMB-PSYLL0008591, NMB-PSYLL0008589, NMB-PSYLL0008587, NMB-PSYLL0008588); 2 f£, same but *Miconia chartacea* (D. Burckhardt) #5(18) (NHMB; dry; NMB-PSYLL0008589); 7 m$, 7 f£, same but *Annona cacans* (D. Burckhardt) #5(4) (NHMB; dry, 70% ethanol; NMB-PSYLL0008003–NMB-PSYLL0008005, NMB-PSYLL0008571); 1 f£, 11 immatures, 2 skin, same but *Annona cacans* (D. Burckhardt) #5(5) (MMBC, NHMB; slide, 70% ethanol; NMB-PSYLL0008023, NMB-PSYLL0008024, NMB-PSYLL0008019, NMB-PSYLL0008020); 7 m$, 3 immatures, same but *Miconia flammea* (D. Burckhardt) #5(8) (MMBC, NHMB; dry, slide, 70% ethanol; NMB-PSYLL0008006, NMB- PSYLL0008007, NMB-PSYLL0008025, NMB-PSYLL0008036–NMB-PSYLL0008038); 2 m$, 2 f£, same but 21.1833/2000, W43.9167/9667, 940–1140 m, 13–14.vi.2010, Cerrado semideciduous forest, along forest edge (D. Burckhardt) #7 (NHMB; dry, slide; NMB- PSYLL0008031 [voucher 74-7], NMB-PSYLL0008021 [voucher 74-8], NMB-PSYLL0008011 [voucher 74-7]); 3 m$, 2 f£, 3 immatures, same but S21.1833/2000, W43.9167/9667, 940–1140 m, 13.vi.2010, *Annona cacans*, Cerrado semideciduous forest (D. Burckhardt) #9(1) (NHMB; dry, slide, 70% ethanol; NMB-PSYLL0008030, NMB- PSYLL0008026, NMB-PSYLL0008035 [voucher 74-3], NMB-PSYLL0008034, NMB- PSYLL0008033); 1 f£, same but S21.1833/2000, W43.9167/9667, 940–1140 m, 3.vi.2010, cleared area with low vegetation and some shrubs and small trees prepared for eucalypt plantations (D. Burckhardt) #6 (NHMB; dry; NMB-PSYLL0008010); 2 f£, 5 immatures, 1 skin, Catas Altas da Noruega, roadside, S20.5806, W43.7218, 720 m, 20.viii.2013, degraded Atlantic forest (D.L. Queiroz) #559 (NHMB; slide, 70% ethanol; NMB-PSYLL0008028, NMB-PSYLL0008018 [LSMelobs-59], NMB-PSYLL0008016 [LSMelobs-59], NMB-PSYLL0008017 [LSMelobs-59]); 3 m$, 1 f£, Lavras, Universidade Federal de Lavras, UFLA, S21.2308, W44.9914, 900 m, 2–11.vi.2010, park trees, forest edge, hedges and plantations (D. Burckhardt) #2 (NHMB; dry; NMB-PSYLL0008000–NMB-PSYLL0008002; 1 m$, Paula Cândido, S20.8060, W42.9798, 790 m, 22.viii.2013, experimental plot coffee and *Inga* plantation (D.L. Queiroz) #565 (NHMB; 70% ethanol; NMB-PSYLL0008029).

**Material not included in type series. Brazil: MINAS GERAIS:** 2 m$, 4 f£, 2 immatures, Lavras, S21.2672, W44.9353, 900 m, 1–6.vi.2010, *Annona* cf. *cacans*, edge of Atlantic forest around coffee plantation mixed with pastures (D. Burckhardt) #1(1) (NHMB; dry, slide, 70% ethanol; NMB-PSYLL0007996, NMB-PSYLL0008080, NMB-PSYLL0008032 [LSMelobs- 74], NMB-PSYLL0007999, NMB-PSYLL0008022).

**Description. *Adult.*** Coloration. Sexually dimorphic. Male dark brown to black. Antenna ochreous, apices of segments 3–8 and entire segments 9–10 brown to dark brown.

Meracanthus, tibiae, tarsi and apex of femora greyish white. Forewing (Fig. 16J) greyish white in basal third, light brown in apical two thirds; bearing indistinct brown dots, sparse basally, dense and often confluent in apical two thirds. Subgenital plate reddish brown apically; paramere light brown. Intersegmental membranes reddish. Female as male but head ochreous with fine, almost black dots on vertex. Thorax brown with dark dots and patches, and longitudinal stripes on mesoscutum. Apices of proctiger and subgenital plate brown.

Structure. Head, in lateral view, inclined inclined at > 45° from longitudinal body axis. Vertex (Fig. 10F) in basal half with imbricate microsculpture, bearing short setae. Forewing (Fig. 16J) with Rs weakly curved to fore margin apically; surface spinules present in all cells, leaving narrow spinule-free stripes along the veins, forming hexagons of a single row of spinules; spinules sparse in basal half of the wing and absent from basal half of cell c+sc. Hindwing with 9 ungrouped costal setae. Metatibia bearing 7–9 grouped apical spurs, arranged as 3–4 + 4–5, anteriorly separated by a few bristles.

Terminalia (Fig. 27G–L). Male. Proctiger weakly produced posteriorly; densely covered with long setae in apical two thirds. Subgenital plate, in lateral view, subtriangular; dorsal margin straight proximally, curved in apical third; posterior margin almost straight in the middle, curved ventrally; with long setae. Paramere, in lateral view, lanceolate, apex subacute; in dorsal view, apex blunt, directed upwards and slightly inwards, lacking distinct sclerotised tooth; both outer and inner faces with dense, long setae; posterior margin with longer setae. Proximal segment of aedeagus with apical part weakly subdivided. Distal segment of aedeagus almost straight in basal two thirds, dorsal margin weakly concave; ventral process situated in apical third of the segment, lacking lateral lobes; in lateral view, relatively long, simple, rounded apically; in dorsal view, ventral process tongue-shaped, slightly wider than apical dilation, with irregularly rounded apex; apical dilation, in lateral view, short, subparallel-sided, with irregularly rounded apex, basally with relatively large membranous lobe at dorsal margin, in dorsal view, apical dilation narrow, with narrowly rounded apex; sclerotised end tube moderately long and almost straight. – Female terminalia cuneate; densely covered with setae. Dorsal margin of proctiger, in lateral view, with imperceptible hump distal to circumanal ring, almost straight otherwise; in dorsal view, apex subacute; circumanal ring, in dorsal view, distinctly cruciform. Subgenital plate, in lateral view, irregularly narrowing to pointed apex, which is slightly upturned; in ventral view, apex rounded.

***Fifth instar immature.*** Coloration. Brownish yellow to pale brown; cephalothoracic sclerite, antenna, wing pads, legs and caudal plate brown.

Structure. Eye with one short, simple ocular seta dorsally. Antennal segments with following numbers of pointed: 1(0), 2(1), 3(0), 4(2), 5(0), 6(2), 7(1), 8(1), 9(0), 10(0). Forewing pad with five marginal sectasetae, in some specimens one relatively large lanceolate is present dorsally; hindwing pad with two marginal pointed sectasetae, lacking sectasetae or larger lanceolate setae dorsally. Tarsal arolium broadly triangular apically, slightly longer than claws. Abdomen with 1–2 lateral pointed sectasetae on either side anterior to caudal plate. Caudal plate with anterior margin close to anterior margin of extra pore fields; with 3–5 (1+2 or 2+3) pointed sectasetae on either side laterally, and three pointed sectasetae subapically, on either side of circumanal ring dorsally. Extra pore fields forming continuous outer and inner bands, consisting of small oval patches; outer band long medially, end pointing outwards. Circumanal ring small.

**Host plant.** *Annona cacans* Warm. (Annonaceae). According to the field notes (D. Burckhardt, unpublished data), sample DB#5(8) with adults and immatures was collected on *Miconia flammea* (Melastomataceae). This is a very unlikely host and possibly due to a mixup of the samples in the field.

**Distribution.** Brazil (MG).

**Derivation of name.** From the Latin adjective *obscurus* = dark, referring to the dark body colour.

**Comments.** *Melanastera obscura* sp. nov. resembles *M. australis* sp. nov., *M. dimorpha* sp. nov., *M. guatteriae* sp. nov. and *M. nasuta* sp. nov. in the lanceolate paramere. It differs from the other species as indicated in the key.

The specimens of the sample DB#1(1) differ considerably from those of samples DB#9(1) and DLQ#559 in the DNA barcoding sequences; the uncorrected p-distances for *cytb* between them are 8.10% and 8.97%, respectively. Although no morphological differences have been found between DB#1(1) and the other samples, the high genetic distance would suggest that *M. obscura* sp. nov. may represent a complex of cryptic species.

## The *trematos*-group

**Description. *Adult*.** Head, in lateral view, inclined at ≥ 45° from longitudinal axis of body. Vertex (Figs 10G–I, 11A–H) trapezoidal, covered with distinct imbricate microsculpture (partly reduced in m$ of *M. dimorpha* sp. nov.: Fig. 10J) and microscopical setae. Thorax weakly or moderately arched, covered with microscopical setae. Forewing (Figs 17, 18A, B) relatively uniformly coloured, with small brown dots; pterostigma distinctly expanding towards the middle or apical third, longer than 4.0 times as wide; vein R_1_ weakly convex or straight medially. Paramere, in lateral view, irregularly spatulate or lanceolate. Distal segment of aedeagus bearing ventral process (small in *M. cabucu* sp. nov. and *M. dimorpha*) which is, in dorsal view, narrower than apical dilation.

***Immature.*** Antenna 10-segmented.

**Comments.** The group includes eleven species from Brazil described here and *Melanastera maculipennis* (Brown & Hodkinson) from Panama. Eight species develop on Melastomataceae (*Miconia*, *Pleroma*, *Tibouchina*), two on Cannabaceae (*Trema*) and one each on Annonaceae (*Guatteria*) and Myristicaceae (*Virola*).

### 27 Melanastera trematos sp. nov

(Figs 5D, 10G, 17A, 27M–R)

**Type material. Holotype m$: Brazil: MINAS GERAIS:** Patos de Minas, Parque Mocambo, S18.5836, W46.5053, 840 m, 27.x.2012, *Trema micranthum* (D. Burckhardt & D.L. Queiroz) #48(6) (UFPR; dry).

**Paratypes. Brazil: MATO GROSSO DO SUL:** 2 f£, Ponta Porã, Fazenda Mariana, Conflora, S22.0027, W55.5658, 590 m, 12.ix.2013 (D.L. Queiroz) #573 (NHMB; 70% ethanol; NMB- PSYLL0007695); 1 f£, same but S22.0325, W55.518, 560 m, #575 (NHMB; 70% ethanol; NMB-PSYLL0007696).—**Minas Gerais:** 2 m$, 14 f£, 4 immatures, same as holotype but (MMBC, NHMB, UFPR; dry, slide, 70% ethanol; NMB-PSYLL0007666, NMB- PSYLL0007698, NMB-PSYLL0007706 [LSMeltre-8], NMB-PSYLL0007707 [LSMeltre-8], NMB-PSYLL0007708, NMB-PSYLL0007704); 3 m$, 8 f£, 8 immatures, same but 21.ii.2018, *Trema micranthum* (D. Burckhardt & D.L. Queiroz) #278(1) (NHMB; dry, 70% ethanol; NMB-PSYLL0007690, NMB-PSYLL0007667, NMB-PSYLL0008298).—**Paraná:** 2 m$, 1 f£, Amaporã, Parque Estadual de Amaporã, S23.0835, W52.7938, 340 m, 10.viii.2019, cf. *Ossaea* sp., border of river (D. Burckhardt & D.L. Queiroz) #341(2) (NHMB; 70% ethanol; NMB-PSYLL0007692); 1 m$, 1 f£, Campina da Lagoa, Beira de estrada, S24.9727, W51.5517, 630 m, 7.vi.2013, degraded vegetation (D.L. Queiroz) #512 (NHMB; 70% ethanol; NMB-PSYLL0007694); 5 m$, 6 f£, 10 immatures, Cerro Azul, BR-476 km 69, S25.0685, W49.0877, 1080 m, 18–19.iv.2013, *Trema micranthum* (D. Burckhardt & D.L. Queiroz) #106(3) (NHMB; dry, slide, 70% ethanol; NMB-PSYLL0007685, NMB- PSYLL0007699, NMB-PSYLL0007700, NMB-PSYLL0007701); 1 f£, Colombo, Embrapa campus, S25.3215, W49.1579, 920 m, 1.v.2019 (D. Burckhardt & D.L. Queiroz) #339 (NHMB; 70% ethanol; NMB-PSYLL0007691); 1 m$, 2 f£, 3 immatures, Foz do Iguaçu, Parque Nacional do Iguaçu, Trilha do poço preto, S25.5969, W54.3921, 180 m, 7.vii.2021, *Trema micranthum* (D.L. Queiroz) #1002(1) (NHMB; 70% ethanol; NMB-PSYLL0008326); 4 f£, same but Parque das Aves, sede até trilha da onça, S25.6138, W54.4822, 225 m, 9.vii.2021, *Trema* sp. (D.L. Queiroz) #1006(3) (NHMB; 70% ethanol; NMB- PSYLL0008327); 1 m$, 6 f£, 1 immature, Irati, S25.4278/4883, W50.6096/6451, 770 m, 14.vi.2017, *Trema micranthum* (D. Burckhardt & D.L. Queiroz) #230(1) (NHMB; 70% ethanol; NMB-PSYLL0007687); 1 m$, 1 f£, 3 immatures, Londrina, Parque Estadual Mata dos Godoy, S23.4428, W51.2423, 630 m, 12‒14.viii.2019, *Trema micranthum* (D. Burckhardt & D.L. Queiroz) #343(5) (NHMB; 70% ethanol; NMB-PSYLL0007693); 2 m$, 4 f£, 4 immatures, Mallet, ARIE Serra do Tigre, S25.9465, W50.8069, 1010 m, 13‒14.vi.2017, *Trema micranthum* (D. Burckhardt & D.L. Queiroz) #228(6) (NHMB; 70% ethanol; NMB- PSYLL0007688); 1 m$, Morretes, Embrapa campus, S25.4515, W48.8723, –30 m, 28.viii.2018 (D.L. Queiroz) #885 (NHMB; dry; NMB-PSYLL0007697); 5 m$, 4 f£, 18immatures, 3 skins, Ponta Grossa, Parque Estadual de Vila Velha, S25.22384/24645, W49.99272/W50.03994, 750‒870 m, 12‒14.vii.2017, *Trema micranthum* (D. Burckhardt & D.L. Queiroz) #246(7) (NHMB; 70% ethanol; NMB-PSYLL0007689); 1 m$, same but S25.22384/24645, W49.99272/W50.03994, 750‒870 m, 12‒14.vii.2017, *Trema micranthum* (L. Serbina) (MMBC; slide).—**Santa Catarina:** 2 m$, 4 f£, 1 immature, Florianópolis, Praia da solidão, S27.7911, W48.5320, 30 m, 13.i.2019 (D.L. Queiroz) #920(3) (NHMB; 70% ethanol; NMB-PSYLL0007705); 1 m$, 6 f£, Indaial, S26.9316, W49.2883, 70 m, 30.iv.2013, *Trema micranthum* (D. Burckhardt & D.L. Queiroz) #118(2) (NHMB; slide, 70% ethanol; NMB-PSYLL0007709, NMB-PSYLL0007710 [LSMeltre-8], NMB-PSYLL0007711 [LSMeltre-8], NMB-PSYLL0007686); 1 m$, 4 f£, Nova Teutônia, S27.1650, W52.4211, 320 m, 8.vii.1943 (F. Plaumann) 1957-341 (BMNH; dry, slide).—**São Paulo:** 1 f£, São Paulo, 1.iv.2007 (P. Yamamoto) (FSCA; 70% ethanol); 1 f£, same but 29.ix.2006 (FSCA; 70% ethanol); 1 f£, same but 1.x.2007 (FSCA; 70% ethanol); 1 m$, same but 23.x.2007 (FSCA; slide).

**Description. *Adult.*** Coloration. Orange; vertex brownish yellow. Antennal segments 1–3 yellow, segments 4–8 brown, segments 3–8 with dark brown apices, segments 9–10 entirely dark brown. Brown patches on pronotum and mesopraescutum; mesoscutum with brown longitudinal stripes. Forewing (Fig. 17A) amber-coloured, transparent, often with only a few scattered indistict brown dots; apex of pterostigma and apices of veins Rs, M_1+2_, M_3+4_, Cu_1a_ and Cu_1b_ more or less distinctly dark brown. Hindwing pale yellow. Abdominal sternites and basal half of female subgenital plate brown.

Structure. Forewing (Fig. 17A) oval, broadest in the middle, broadly, relatively evenly rounded apically; wing apex in cell r_2_ near apex M_1+2_; C+Sc weakly curved in distal portion; pterostigma distinctly narrower in the middle than cell r_1_, slightly convex in apical two thirds; Rs relatively straight in basal two thirds, obliquely curved to fore margin apically; M longer than M_1+2_ and M_3+4_; Cu_1a_ weakly and irregularly curved, ending proximal of M fork; surface spinules present in all cells, forming hexagons of a single row of spinules, almost absent in basal half of c+sc; spinules leaving narrow spinule-free stripes along the veins in apical part. Hindwing with 6–8 costal setae, somtimes grouped 3–4 + 3–4. Metatibia bearing posteriorly grouped 5–6 apical spurs, arranged as 2–3 + 3, anteriorly separated by 4–5 bristles.

Terminalia (Fig. 27M–R). Male. Proctiger narrowly tubular; densely covered with long setae in apical two thirds. Subgenital plate, in lateral view, irregularly ovoid; dorsal margin strongly curved, posterior margin weakly curved; with long setae. Paramere, in lateral view, irregularly spatular, rounded apically; apex with small, strongly sclerotised tooth, in dorsal view, subacute, directed inwards and slightly posteriad; outer face with dense long setae in apical half; inner face with dense, moderately long setae; posterior margin with long setae. Proximal segment of aedeagus with apical part strongly subdivided. Distal segment of aedeagus weakly curved in basal two thirds; ventral process situated in apical two fifths of the segment, in lateral view, short, simple, lacking lateral lobes, in dorsal view, ovoid, broadest in basal third; apical dilation broad, in lateral view, with subparallel margins, curved, bean-shaped, apex irregularly rounded, dorsal part slightly expanded and membranous at base; sclerotised end tube moderately long, weakly sinuous. – Female terminalia cuneate; densely covered with setae. Dorsal margin of proctiger, in lateral view, almost straight, apex pointed; in dorsal view, apex blunt; circumanal ring, in dorsal view, cruciform. Subgenital plate, in lateral view, irregularly narrowing to apex, pointed apically; in ventral view, apex blunt.

***Fifth instar immature.*** Coloration. Bright yellow; caudal plate pale brown.

Structure. Eye with one long simple ocular seta dorsally. Antennal segments with following numbers of pointed sectasetae: 1(0), 2(1), 3(0), 4(2), 5(0), 6(2), 7(0), 8(1), 9(0), 10(0). Forewing pad with 10–14 marginal pointed sectasetae and ca. 25–30 pointed sectasetae or lanceolate setae dorsally; hindwing pad with 3 marginal and 4–5 dorsal pointed sectasetae. Metatibiotarsus long; tarsal arolium broadly triangular apically, > 1.5 times as long as claws. Abdominal tergites anterior to caudal plate with four rows of 8–10, 6, 8 and 16 pointed sectasetae or lanceolate setae dorsally, respectively. Caudal plate dorsally with one row of 16– 22 pointed sectasetae along anterior margin, a row of 15–16 pointed setae submedially, one group of 3–4 pointed sectasetae on each side in the middle and one group of 9–12 pointed sectasetae on each side subapically, anterior to circumanal ring, and with several long, normal setae on either side ventrally. Extra pore fields forming continuous outer and inner bands, consisting of several large oval and rounded patches; outer band relatively long medially, end pointing outwards. Circumanal ring (Fig. 5D) moderately elongate.

**Host plant.** *Trema micranthum* (L.) Blume (Cannabaceae).

**Distribution.** Brazil (MG, MS, PR, SC, SP).

**Derivation of name.** Named after its host, *Trema*.

**Comments.** *Melanastera trematos* sp. nov. and *M. virolae* sp. nov. differ from other members of the *trematos*-group in the spatulate (versus lanceolate) paramere. *Melanastera trematos* differs from *M. virolae* in the following characters: body and forewing with faint (versus distinct) pattern; dark dots absent (versus present) from antero-basal part of the forewing; PtL/PtW > 4.6 (versus < 4.6); Cu_1a_L/cu_1_W > 3.2 (versus < 3.2); paramere, in lateral view, strongly (versus weakly) convex anteriorly; distal segment of aedeagus with short (versus long) ventral process; apical dilation, in lateral view, curved (versus straight); dorsal margin of female proctiger, in lateral view, almost straight (versus sinuate); FP/SP < 1.7 (versus > 1.7).

*Melanastera trematos* sp. nov. shares the host plant species, *Trema micranthum*, with *M. maculipennis* (Brown & Hodkinson), which was described and is so far only known from Panama. Both species are also similar in size, the forewing shape, venation and pattern, and the structure of the female terminalia (Brown & Hodkinson 1988). *Melanastera trematos* differs from *M. maculipennis* by a much broader, spatulate paramere (as opposed to narrow, elongate) and the distal segment of the aedeagus, which is relatively slender, elongate, only weakly sinuate dorsally in its basal portion and with a ventral process well-separated from the apical dilation (versus relatively thick, short, strongly sinuate dorsally in its basal portion and with ventral process that lies button-like against the base of apical dilation).

### 28 Melanastera virolae sp. nov

(Figs 7E, 10H, 17B, 27S–X)

**Type material. Holotype m$: Brazil: PIAUÍ:** Brasileira/Piracuruca, Parque Nacional de Sete Cidades, S4.0727/0991, W41.6797/7291, 130‒210 m, 21‒24.vi.2016, *Virola bicuhyba* (D. Burckhardt & D.L. Queiroz) #201(4) (UFPR; dry).

**Paratypes. Brazil: PIAUÍ:** 5 m$, 4 f£, 11 immatures, 3 skins, same as holotype but (MMBC, NHMB; dry, slide, 70% ethanol; NMB-PSYLL0008146, NMB-PSYLL0005826, NMB-PSYLL0008197 [LSMelvir-62], NMB-PSYLL0008199 [LSMelvir-62], NMB- PSYLL0008198 [LSMelvir-62], NMB-PSYLL0008200 [LSMelvir-62], NMB- PSYLL0008195, NMB-PSYLL0008196).

**Description. *Adult.*** Coloration. Yellow; head and thorax with dark brown dots. Antennal segments 4–9 with dark brown apices, segment 10 entirely dark brown. Forewing (Fig. 17B) yellow with relatively large dark brown dots, sparse at base, becoming denser towards apex, sometimes confluent at apex; base and apex of pterostigma, apices of veins Rs, M_1+2_, M_3+4_, Cu_1a_, Cu_1b_ and spots on vein A dark brown. Femora with two brown tranverse bands. Abdominal sternites brown laterally.

Structure. Forewing (Fig. 17B) oviform, broadest in the middle, broadly and evenly rounded apically; wing apex situated in cell r_2_ near apex of M_1+2_; C+Sc weakly curved in distal third; pterostigma slightly narrower in the middle than cell r_1_, convex; Rs weakly curved in basal two thirds, obliquely curved to fore margin in apical third; M longer than M_1+2_ and M_3+4_; Cu_1a_ weakly irregularly curved, ending at level of M fork; surface spinules present in all cells, leaving narrow spinule-free stripes along the veins, forming distinct hexagons of a single row of spinules, spinules absent in basal half of cell c+sc. Hindwing with 10 ungrouped or 5 + 5 grouped costal setae. Metatibia bearing 8‒9 grouped apical spurs, arranged as 3‒4 + 5, anteriorly separated by 3 or more bristles.

Terminalia (Fig. 27S–X). Male. Proctiger long, tubular, weakly convex posteriorly; densely covered with long setae in apical half. Subgenital plate irregularly ovoid; dorsal margin irregularly curved, posterior margin straight; with moderately dense, long setae. Paramere, in lateral view, irregularly spatulate; apex, in lateral view, blunt, directed upwards, in dorsal view, truncate, directed upwards and inwards, bearing two small teeth; outer face with dense, moderately long setae mostly posteriorly; inner face with dense, moderately long setae; posterior margin with long setae. Proximal segment of aedeagus with apical part strongly subdivided. Distal segment of aedeagus with dorsal margin distinctly sinuate in basal half; ventral process situated slightly distal of the middle of the segment, in lateral view, long, narrow, simple; in dorsal view, ventral process oblong-oval; apical dilation, in lateral view, elongate, subparallel-sided, with narrowly rounded apex, at base with angular membranous lobe at dorsal margin; in dorsal view, apical dilation narrowly obovate, broadest in apical third; sclerotised end tube short and almost straight. – Female terminalia cuneate; densely covered with setae. Dorsal margin of proctiger, in lateral view, with a distinct hump distal to circumanal ring, straight apically, apex pointed; in dorsal view, apex blunt; circumanal ring, in dorsal view, distinctly cruciform. Subgenital plate, in lateral view, abruptly narrowing in apical half, pointed apically; in ventral view, apex blunt.

***Fifth instar immature.*** Coloration. Light; antenna pale yellow with pale brown apical segment; cephalothoracic sclerite, wing pads, legs and caudal plate yellow.

Structure. Eye with one short simple ocular seta dorsally. Antennal segments with following numbers of pointed sectasetae: 1(0), 2(1), 3(0), 4(2), 5(0), 6(2), 7(1), 8(1), 9(0), 10(0). Forewing pad with 8–10 marginal pointed sectasetae and 0–1 pointed sectaseta dorsally; hindwing pad with 3 large marginal pointed sectasetae and 1–2 smaller pointed sectasetae situated close to pad posterior margin but more dorsally. Metatibiotarsus long; tarsal arolium broadly triangular apically, 1.2–1.5 times as long as claws. Abdomen with three lateral pointed sectasetae on each side anterior to caudal plate. Caudal plate with anterior margin relatively close to anterior margin of extra pore fields; with five pointed sectasetae on either side laterally, and four pointed sectasetae subapically near circumanal ring on either side dorsally. Extra pore fields forming continuous outer and inner bands, consisting of small

oval and rounded patches; outer band long medially, end pointing outwards. Circumanal ring small.

**Host plant.** *Virola bicuhyba* (Schott) Warb. (Myristicaceae).

**Distribution.** Brazil (PI).

**Derivation of name.** Named after its host, *Virola*.

**Comments.** Within the the *trematos*-group, *Melanastera virolae* sp. nov. and *M. trematos* sp. nov. are the only species with a spatulate (versus lanceolate) paramere. They differ from each other as indicated under the latter species.

### 29 Melanastera inconspicuae sp. nov

(Figs 3A, B, 10I, 17C, 28A–F)

**Type material. Holotype m$: Brazil: PARANÁ:** Bocaiuva do Sul, Sítio do irmão do Arnaldo Soares, S17.6439, W46.7007, 980 m, 3.xii.2015, *Miconia inconspicua* (D.L. Queiroz) #682(1) (UFPR; dry).

**Paratypes. Brazil: Paraná:** 4 m$, 10 f£, 3 immatures, 1 skin, same as holotype but (MMBC, NHMB, UFPR; dry, slide, 70% ethanol; NMB-PSYLL0006055, NMB- PSYLL0007978, NMB-PSYLL0007903, NMB-PSYLL0007908, NMB-PSYLL0007913 [LSMelinc-97], NMB-PSYLL0007914 [LSMelinc-97], NMB-PSYLL0007915, NMB- PSYLL0007916).

**Description. *Adult.*** Coloration. Yellow, head and thorax with irregular brown patches, mesoscutum with longitudinal brown bands. Antennal segments 3–8 with brown apices, segments 9–10 entirely brown. Forewing (Fig. 17C) yellow with sparse, faint brown dots mostly in apical half of wing; apices of veins Rs, M_1+2_, M_3+4_, Cu_1a_ and Cu_1b_ dark brown, sometimes indistinct. Femora with a subapical transverse brown band.

Structure. Forewing (Fig. 17C) oval, widest in the middle, broadly evenly rounded apically; wing apex situated in cell r_2_, slightly remote from apex of M_1+2_; C+Sc distinctly bent in distal third; pterostigma distinctly narrower than cell r_1_ in the middle, weakly widening to apical third; Rs relatively straight in proximal three quarters, moderately curved to fore margin apically; M longer than M_1+2_ and M_3+4_; Cu_1a_ irregularly curved, ending at about same level with M fork; surface spinules present in all cells, leaving relatively narrow spinule-free stripes along the veins, forming hexagons of a single and/or double rows of spinules.

Hindwing with 4 + 2–3 grouped costal setae. Metatibia bearing 6–9 grouped apical spurs, arranged as 3–5 + 3–5, anteriorly separated by 1–2 bristles.

Terminalia (Fig. 28A–F). Male. Proctiger tubular, weakly produced posteriorly; densely covered with long setae in apical two thirds. Subgenital plate, in lateral view, irregularly ovoid; dorsal margin curved; posterior margin relatively straight; with dense, long setae mostly posteriorly. Paramere irregularly lanceolate, strongly narrowed in apical third; apex, in lateral view, blunt, directed upwards, in dorsal view, narrowly blunt, directed upwards and inwards, lacking distinct sclerotised tooth; both outer and inner faces with dense, moderately long setae; posterior margin with longer setae. Proximal segment of aedeagus with apical part moderately subdivided. Distal segment of aedeagus with dorsal margin sinuate in basal half; ventral process situated slightly distal of the middle of the segment, in both lateral and dorsal views, simple, long and narrow, lamellar, in dorsal view, subparallel-sided; apical dilation, in lateral view, relatively narrow, subparallel-sided, rounded apically, with a large membranous sack at dorsal margin basally; in dorsal view, apical dilation broad, subrectangular, truncate apically; sclerotised end tube short and weakly curved. – Female terminalia cuneate; covered with setae. Dorsal margin of proctiger, in lateral view, with a small hump distal to circumanal ring, weakly concave in apical third, apex subacute, turned slightly upwards; in dorsal view, apex blunt; circumanal ring, in dorsal view, vaguely cruciform. Subgenital plate, in lateral view, abruptly narrowing in apical third, apex pointed; in ventral view, apex blunt.

***Fifth instar immature.*** Coloration. Pale brown; antenna gradually becoming darker towards apex; cephalothoracic sclerite, wing pads, legs and caudal plate brown.

Structure. Body sparsely covered with minute clavate and narrowly lanceolate setae. Eye with one short simple ocular seta dorsally. Antennal segments with following numbers of pointed sectasetae: 1(0), 2(2 – one large and one small), 3(0), 4(2), 5(0), 6(2), 7(1), 8(1), 9(0), 10(0). Forewing pad with 4–5 marginal pointed sectasetae, lacking sectasetae dorsally; hindwing pad with 1–2 marginal and 1–2 dorsal pointed sectasetae. Metatibiotarsus short; tarsal arolium broadly fan-shaped apically, 1.3 times as long as claws. Abdomen with 2–3 pointed sectasetae laterally on either side anterior to caudal plate. Caudal plate with anterior margin relatively remote from anterior margin of extra pore fields; with 4–5 (1 or 2 + 3) pointed sectasetae on either side laterally, and with three pointed sectasetae subapically, on either side of circumanal ring dorsally. Extra pore fields forming continuous outer and inner bands, consisting of large and small oval and round patches; outer band short medially, end pointing outwards. Circumanal ring small.

**Host plant.** *Miconia inconspicua* Miq. (Melastomataceae).

**Distribution.** Brazil (PR).

**Derivation of name.** Named after its host, *M. inconspicua*.

**Comments.** Within the the *trematos*-group, *Melanastera inconspicuae* sp. nov. resembles *M. clavata* sp. nov. in the long, simple, lamellar ventral process of the distal aedeagal segment, the sinuate dorsal margin of the female proctiger and the minute narrowly lanceolate setae covering the body of the immature. *Melanastera inconspicuae* differs from *M. clavata* in the following characters: paramere, in lateral view, broader and with more strongly sinuate anterior margin; ventral process of the distal aedeagal segment narrower, with subparallel margins, in dorsal view; FP/HW > 0.75 (versus < 0.75).

### 30 Melanastera dimorpha sp. nov

(Figs 10J, 11A, 17D, E, 28G–L)

**Type material. Holotype m$: Brazil: MINAS GERAIS:** Vargem Bonita, Parque Nacional da Serra da Canastra, Cachoeira Casca d’Anta, plateau, S20.2976/2986, W46.5195/5289, 1160– 1250 m, 6.ix.2014, *Guatteria sellowiana* (D. Burckhardt & D.L. Queiroz) #144(6) (UFPR; dry).

**Paratypes. Brazil: MINAS GERAIS:** 5 m$, 7 f£, 5 immatures, same as holotype but (MMBC, NHMB, UFPR; dry, slide, 70% ethanol; NMB-PSYLL0007986, NMB- PSYLL0007987, NMB-PSYLL0007843 [LSMeldim-39], NMB-PSYLL0007844 [LSMeldim- 39], NMB-PSYLL0007845 [LSMeldim-39], NMB-PSYLL0007846 [LSMeldim-39], NMB-PSYLL0007847); 1 m$, 10 immatures, same but (D. Burckhardt & D.L. Queiroz) #144(2) (NHMB; 70% ethanol; NMB-PSYLL0007819).

**Description. *Adult.*** Coloration. Sexually dimorphic. Male dark brown to almost black.

Antenna whitish, scape and pedicel brown, apices of segments 3–8 and entire segments 9–10 dark brown. Forewing (Fig. 17D) relatively evenly dark brown, with white band in cell c+sc along R+M+Cu and along nodal line; in apical third of wing with sparse inconspicuous irregular light dots. Metatibia and metabasitarsus yellow, apical metatarsal segment brown. Female straw-coloured, head and thorax with small dark brown dots, mesoscutum with brown longitudinal bands. Antenna as in male. Forewing (Fig. 17E) ochreous with moderately dense brown dots, becoming denser towards apex; apex of pterostigma and apices of veins Rs, M_1+2_, M_3+4_, Cu_1a_ and Cu_1b_ dark brown. Coxae brown, meracanthus whitish; pro- and mesofemora with greyish brown transverse subapical band, metafemora with brownish dots; apical tarsal segments brown. Abdominal tergites partly, and sternites entirely dark brown. Younger specimens with less extended dark colour.

Structure. Microsculpture on vertex partly reduced in m$ (Fig. 10J), fully developed on f£ (Fig. 11A). Forewing oval, broadest distal of the middle, broadly, unevenly rounded apically; wing apex in male situated in cell r_2_ near apex of M_1+2_, in female at apex of M_1+2_; C+Sc weakly curved in distal third (male) or in the middle (female); pterostigma distinctly (in male) or slightly (in female) narrower in the middle than cell r_1_, irregularly convex; Rs in basal two thirds weakly, in apical third strongly curved to fore margin; M longer than M_1+2_ and M_3+4_; Cu_1a_ in basal two thirds moderately (in male) or weakly (in female) curved, ending proximal of M fork; surface spinules in male present in all cells except for most of cell c+sc, leaving narrow spinule-free stripes along the veins, forming hexagons of a single row of spinules in apical third of the wing; surface spinules in female present in all cells, covering the entire membrane, forming distinct hexagons of a double row of spinules. Hindwing with 8–11 ungrouped or grouped (3–4 + 3–4) costal setae. Metatibia bearing 5–7 grouped apical spurs, arranged as 2–3 + 3–4, separated 3 bristles.

Terminalia (Fig. 28G–L). Male. Proctiger, slender, subcylindrical, weakly produced posteriorly; densely covered with long setae in apical two thirds. Subgenital plate, in lateral view, irregularly ovoid; dorsal margin weakly curved, posterior margin slightly convex; with sparse, long setae. Paramere, in lateral view, relatively symmetrically, narrowly lanceolate; paramere apex, in dorsal view, blunt, directed inwards, lacking distinct sclerotised tooth; outer face with sparse, moderately long setae in apical two thirds; inner face with dense, moderately long setae; posterior margin with long setae. Proximal segment of aedeagus with apical part moderately subdivided. Distal segment of aedeagus weakly sinuate in basal three fifths; ventral process situated in apical third of segment, in lateral view, short, simple, knob-shaped, directed slightly obliquely ventro-apicad, almost in continuation of basal part of the segment, unevenly rounded apically, lacking lateral lobes; in ventral view, ventral process small, oval, with evenly rounded apex; apical dilation, in lateral view, long, slightly widening towards unevenly rounded apex, which is weakly directed ventrad, apical dilation dorsally bearing a large membranous lobe, dorsal margin convex, strongly angular basally; in dorsal view, apical dilation gradually widening towards evenly rounded apex; sclerotised end tube moderately long and slightly sinuate. – Female terminalia cuneate; densely covered with setae. Dorsal margin of proctiger, in lateral view, almost straight, apex subacute; in dorsal view, apex blunt; circumanal ring, in dorsal view, cruciform. Subgenital plate, in lateral view, irregularly narrowing to apex, pointed apically; in ventral view, apex narrowly truncate.

***Fifth instar immature.*** Coloration. Pale yellow; antenna pale yellow, scape and pedicel pale brown, gradually becoming darker towards apex, segments 9–10 entirely dark brown; cephalothoracic sclerite, wing pads, legs and caudal plate brown.

Structure. Eye with one short simple ocular seta dorsally. Antennal segments with following numbers of pointed sectasetae: 1(0), 2(1), 3(0), 4(2), 5(0), 6(2), 7(0), 8(1), 9(0),

10(0). Forewing pad with five marginal and one dorsal pointed sectasetae; hindwing pad with two marginal pointed sectasetae and one small dorsal pointed sectaseta. Metatibiotarsus relatively long, bearing eight pointed sectasetae in three rows on outer side; tarsal arolium broadly triangular apically, 1.5 times as long as claws. Abdomen with four lateral pointed sectasetae on each side anterior to caudal plate. Caudal plate with anterior margin remote from anterior margin of extra pore fields; with five pointed sectasetae on either side laterally, a single pointed sectasetae in the middle on either side of extra pore fields dorsally, and three pointed subapical sectasetae near circumanal ring on either side dorsally. Extra pore fields forming continuous outer and inner bands, consisting of relatively small oval patches; outer band long medially, end pointing outwards. Circumanal ring small.

**Host plant.** *Guatteria sellowiana* Schltdl. (Annonaceae).

**Distribution.** Brazil (MG).

**Derivation of name.** Named after the sexual dimorphism of the species.

**Comments.** Within the the *trematos*-group, *M. dimorpha* sp. nov. is characterised by its sexual dimorphism. It resembles *M. cabucu* sp. nov. in the knob-shaped ventral process of the distal aedeagal segment. The two species differ in details of the termialia: paramere slender in *M. dimorpha* and massive in *M. cabucu*; dorsal margin of the female proctiger straight in the former and sinuate in the latter.

### 31 Melanastera cabucu sp. nov

(Figs 11B, 17F, 28M–R)

**Type material. Holotype m$: Brazil: PARANÁ:** Morretes, Floresta Marumbi, trail to hotel, S25.4459, W48.8873, 170 m, 14.ix.2011, *Miconia formosa* (D. Burckhardt & D.L. Queiroz) #6(3) (UFPR; dry).

**Paratypes. Brazil: PARANÁ:** 3 m$, Morretes, Estação Marumbi, S25.4484/4492, W48.8901/8915, 230–240 m, 14.ix.2011, *Miconia formosa* (D. Burckhardt & D.L. Queiroz) #5(4) (NHMB; dry, slide, 70% ethanol; NMB-PSYLL0007640, NMB-PSYLL0007636,

NMB-PSYLL0008622 [LSMelcab-30], NMB-PSYLL0007637); 2 m$, 1 f£, same as holotype but (NHMB; slide; NMB-PSYLL0007638, NMB-PSYLL0007639, NMB-PSYLL0007641); 1

f£, Morretes, Embrapa campus, S25.4515, W48.8723, 30.xi.2004, *Miconia formosa* (D.L. Queiroz) (NHMB; dry; NMB-PSYLL0007642).

**Description. *Adult.*** Coloration. Straw-coloured. Head and thorax with small dark brown dots; mesoscutum with brown longitudinal stripes. Antennal segments 3–8 with dark brown apices, segments 9–10 entirely dark brown. Forewing (Fig. 17F) amber-coloured with sparse brown dots; base and apex of pterostigma and apices of veins Rs, M_1+2_, M_3+4_, Cu_1a_ and Cu_1b_ dark brown. Femora with brown dots; tarsi brown. Abdominal tergites partly and sternites entirely dark brown.

Structure. Forewing (Fig. 17F) oval, broadest distal of the middle, broadly and evenly rounded apically; wing apex situated in cell r_2_ near apex of M_1+2_; C+Sc eakly curved in distal third; pterostigma slightly narrower than r_1_ cell in the middle, weakly widening to apical two thirds; Rs almost straight in basal two thirds, obliquely curved to fore margin apically; M longer than M_1+2_ and M_3+4_; Cu_1a_ weakly irregularly curved, ending at level of M fork; surface spinules present in all cells, leaving narrow spinule-free stripes along the veins, forming hexagons of a double row of spinules, spinules absent from basal half of cell c+sc. Hindwing with 3–4 + 4 grouped costal setae. Metatibia bearing 8–10 grouped apical spurs, arranged as 3–4 + 4–6, anteriorly separated by 3 bristles.

Terminalia (Fig. 28M–R). Male. Proctiger tubular, in basal third weakly produced posteriorly; densely covered with long setae in apical two thirds. Subgenital plate, in lateral view, subglobular; dorsal margin strongly, irregularly curved, posterior margin curved; with moderately long setae. Paramere, in lateral view, asymmetrically lanceolate; apex, in lateral view, blunt, directed upwards; in dorsal view, apex blunt, directed upwards, slightly inwards and hardly posteriad, lacking distinct sclerotised tooth; outer face with sparse, moderately long setae mostly in apical half; inner face with dense, moderately long setae; posterior margin with long setae. Proximal segment of aedeagus with apical part strongly subdivided, bulbous. Distal segment of aedeagus with dorsal margin weakly curved in basal half; ventral process situated in apical third of segment, in both lateral and dorsal views, very short, simple, rounded, button-like, lacking lateral lobes; apical dilation, in lateral view, relatively large, subrectangular, base bearing a large membranous lobe at dorsal margin; in dorsal view, apical dilation elongate, with slightly truncate apex; sclerotised end tube short and weakly curved. – Female terminalia cuneate; densely covered with setae. Dorsal margin of proctiger, in lateral view, slightly convex distal to circumanal ring and distinctly concave in apical half, apex pointed, upturned; in dorsal view, apex blunt; circumanal ring, in dorsal view, distinctly cruciform. Subgenital plate, in lateral view, abruptly narrowing in apical half, pointed apically; in ventral view, apex blunt.

*Fifth instar immature.* Unknown.

**Host plant.** Adults were collected on *Miconia formosa* Cogn. (Melastomataceae), which is possibly a host.

**Distribution.** Brazil (PR).

**Derivation of name.** Named after *Miconia cabucu*, noun in apposition, one of the synonyms of the probable host *M. formosa*.

**Comments.** *Melanastera cabucu* sp. nov. resembles *M. dimorpha* sp. nov. in the knob-shaped ventral process of the distal aedeagal segment. The two species differ as outlined under *M. dimorpha*.

### 32 Melanastera itatiaia sp. nov

(Figs 11C, 17G, 28S–X)

**Type material. Holotype m$: Brazil: RIO DE JANEIRO:** Itatiaia, Parque Nacional do Itatiaia, Mirante do Último Adeus, S22.4589, W44.6071, 790 m, 15‒18.iv.2019, *Pleroma granulosum* (D. Burckhardt & D.L. Queiroz) #332(2) (UFPR; slide; [LSMelita-109]).

**Paratypes. Brazil: RIO DE JANEIRO:** 1 f£, same as holotype but (NHMB; slide; NMB- PSYLL0007936 [LSMelita-109]).

**Description. *Adult.*** Coloration. Yellow; head and thorax with dark brown dots. Antennal segments 3–9 apically and segment 10 entirely dark brown. Mesoscutum with four broad longitudinal stripes, margined by dark brown dots, and one row of dots in the middle.

Forewing (Fig. 17G) yellow with brown pattern consisting of distinct, irregularly spaced dots; apices of veins Rs, M_1+2_, M_3+4_, Cu_1a_ and Cu_1b_ brown.

Structure. Forewing (Fig. 17G) oblong-oval, widest in the middle, slightly irregularly rounded apically; wing apex situated in cell r_2_, near apex of M_1+2_; C+Sc curved in distal third; pterostigma relatively slightly wider than r_1_ cell in the middle, widening to apical quarter; Rs relatively straight in basal two thirds, weakly curved to fore margin apically; M longer than M_1+2_ and M_3+4_; Cu_1a_ irregularly curved, ending slightly proximal of M fork; surface spinules present in all cells, leaving very narrow spinule-free stripes along the veins, forming hexagons of a double row of spinules. Hindwing with 3 + 2 grouped costal setae. Metatibia bearing 8–9 grouped apical spurs, arranged as 3–4 + 4–5, anteriorly separated by 3 bristles.

Terminalia (Fig. 28S–X). Male. Proctiger with weakly convex posterior margin; densely covered with long setae in apical two thirds. Subgenital plate subglobular; dorsal margin strongly curved in distal third; posterior margin slightly concave; with long setae. Paramere, in lateral view, relatively evenly lanceolate, broadest medially; apex, in lateral view, blunt, directed upwards; in dorsal view, apex narrow, subacute, directed upwards and slightly inwards, lacking distinct sclerotised tooth; both outer and inner faces with dense, moderately long setae; posterior margin with longer setae. Proximal segment of aedeagus with apical part moderately subdivided. Distal segment of aedeagus in basal half with weakly sinuate dorsal margin; ventral process situated slightly distal of the middle of the segment, in lateral view, ventral process small, narrow, simple, lamellar, lacking lateral lobes, in dorsal view, ventral process lanceolate, broadest in apical third; apical dilation, in lateral view, relatively narrow, weakly curved, with narrowly rounded apex, dorsal part of apical dilation membranous and produced at base; sclerotised end tube moderately long and slightly curved. – Female terminalia cuneate; densely covered with setae. Dorsal margin of proctiger, in lateral view, weakly convex distal to circumanal ring, apex slightly upturned, pointed; in dorsal view, apex blunt; circumanal ring, in dorsal view, cruciform. Subgenital plate, in lateral view, distinctly, abruptly narrowing to apex in apical half, pointed apically; in ventral view, apex blunt or truncate.

*Fifth instar immature.* Unknown.

**Host plant.** Adults were collected on *Pleroma granulosum* (Desr.) D.Don (Melastomataceae) which is possibly a host.

**Distribution.** Brazil (RJ).

**Derivation of name.** Named after the Itatiaia National Park, where the types were collected.

**Comments.** Within the *trematos*-group, *M. itatiaia* sp. nov. resembles *M. pusilliflorae* sp. nov. in the forewing colour, shape and venation and the symmetrically lanceolate paramere. *Melanastera itatiaia* differs from *M. pusilliflorae* in the lighter body colour, FL/FW > 2.28 (versus FL/FW < 2.28), the shorter ventral process of the distal aedeagal segment and the apically pointed and slightly upturned female proctiger (versus subacute and straight).

### 33 Melanastera pusilliflorae sp. nov

(Figs 11D, 17H, 29A–F)

**Type material. Holotype m$: Brazil: MINAS GERAIS:** Viçosa, Universidade Federal Viçosa, UFV, campus Belvedere, S20.7566, W42.8600, 700 m, 8.vii.2012, *Miconia pusilliflora* (D. Burckhardt & D.L. Queiroz) #33(1) (UFPR; dry).

**Paratypes. Brazil: MINAS GERAIS:** 7 m$, 14 f£, 3 immatures, 3 skins, same as holotype but (MMBC, NHMB, UFPR; dry, slide, 70% ethanol; NMB-PSYLL0008056, NMB- PSYLL0008054, NMB-PSYLL0008070 [LSMelpus-2], NMB-PSYLL0008071 [LSMelpus- 2], NMB-PSYLL0008069 [LSMelpus-2], NMB-PSYLL0008068 [LSMelpus-2], NMB- PSYLL0008072, NMB-PSYLL0008073, NMB-PSYLL0008074, NMB-PSYLL0008047).—

**Paraná:** 4 immatures, Londrina, Parque Estadual Mata dos Godoy, S23.4428, W51.2423, 630 m, 12‒14.viii.2019, *Miconia pusilliflora* (D. Burckhardt & D.L. Queiroz) #343(15) (NHMB; slide, 70% ethanol; NMB-PSYLL0008048, NMB-PSYLL0008075).

**Description. *Adult.*** Coloration. Sexually dimorphic. Male dark brown. Antenna yellow, segments 1–2 partially brown, segments 4–9 with brown apices, segment 10 entirely brown. Forewing (Fig. 17H) with amber-coloured veins, irregular, sparse brown dots, absent from wing base; apices of pterostigma and veins Rs, M_1+2_, M_3+4_, Cu_1a_ and Cu_1b_ brown. Legs straw-coloured, femora with brown dots. Abdominal tergites and terminalia straw-coloured and brown. Female as male but head and thorax with brown or reddish brown pattern. Forewing with brown dots also present in basal part of wing. Abdominal tergites dark brown.

Structure. Forewing (Fig. 17H) oviform, broadest in the middle, broadly and evenly rounded apically; wing apex situated in cell r_2_ close to apex of M_1+2_; C+Sc weakly curved in distal third; pterostigma distinctly narrower in the middle than cell r_1_, widening to the middle; Rs relatively straight in basal two thirds, distinctly curved to fore margin apically; M longer than M_1+2_ and M_3+4_; Cu_1a_ irregularly curved, ending slightly proximal of M fork; surface spinules present in all cells, covering the entire membrane up to the veins, forming hexagons of a double row of spinules, spinules absent from base of cell c+sc. Hindwing with 5–6 grouped or ungrouped (3–4 + 3–4) costal setae. Metatibia bearing 7–9 grouped apical spurs, arranged as 3–4 + 4–5, anteriorly separated by ca. 3 bristles.

Terminalia (Fig. 29A–F). Male. Proctiger weakly produced posteriorly; densely covered with long setae in apical two thirds. Subgenital plate large, in lateral view, subglobular; dorsal and posterior margins almost straight; with densely spaced long setae. Paramere, in lateral view, evenly lanceolate, broadest medially; apex, in lateral view, blunt, directed upwards, in dorsal view, apex narrow, subacute, directed upwards and slightly inwards, lacking distinct sclerotised tooth; outer face with short setae in apical two thirds; inner face with dense, moderately long setae; posterior margin with long setae. Proximal segment of aedeagus with apical part moderately subdivided. Distal segment of aedeagus in basal half with weakly sinuate dorsal margin; ventral process situated slightly distal of the middle of the segment, in lateral view, ventral process moderately long, narrow, simple, lamellar, lacking lateral lobes, in dorsal view, ventral process lanceolate, broadest in the middle, slightly narrower than apical dilation; apical dilation, in lateral and dorsal views, relatively narrow, subparallel- sided, with broadly rounded apex, dorsal part of apical dilation membranous and expanded at base; sclerotised end tube moderately long and slightly curved. – Female terminalia cuneate; densely covered with setae. Dorsal margin of proctiger, in lateral view, slightly convex distal to circumanal ring, apex almost straight, blunt; in dorsal view, apex blunt; circumanal ring, in dorsal view, cruciform. Subgenital plate, in lateral view, abruptly narrowing to apex in apical half, pointed apically; in ventral view, apex truncate.

***Fifth instar immature.*** Coloration. Light; cephalothoracic sclerite, antenna, wing pads, legs and caudal plate dark brown.

Structure. Eye with one short simple ocular seta dorsally. Antennal segments with following numbers of pointed sectasetae: 1(0), 2(1), 3(0), 4(2), 5(0), 6(2), 7(1), 8(2), 9(0), 10(0). Forewing pad with 2–3 marginal pointed sectasetae; hindwing pad with one marginal pointed sectaseta. Metatibiotarsus long; tarsal arolium narrowly fan-shaped apically, 1.5 times as long as claws. Abdomen lacking larger setae dorsally, with one lateral pointed sectaseta on each side anterior to caudal plate. Caudal plate with anterior margin very close to anterior margin of extra pore fields; with four pointed sectasetae on either side laterally and two pointed sectasetae near circumanal ring on either side dorsally. Extra pore fields forming continuous outer and inner bands, consisting of relatively small oval patches; outer band long medially, end pointing outwards. Circumanal ring large.

**Host plant.** *Miconia pusilliflora* (DC.) Naudin (Melastomataceae).

**Distribution.** Brazil (MG, PR).

**Derivation of name.** Named after its host, *M. pusilliflora*.

**Comments.** *Melanastera pusilliflorae* sp. nov. resembles *M. itatiaia* sp. nov., from which it can be separated as indicated under the latter species.

### 34 Melanastera goldenbergi sp. nov

(Figs 11E, 17I, 29G–L)

**Type material. Holotype m$: Brazil: AMAZONAS:** Rio Preto da Eva, road from Embrapa, Fazenda Rio Urubu to Rio Preto da Eva, S2.4717, W59.5844, 120 m, 25.iv.2014*, Miconia manauara* (D. Burckhardt & D.L. Queiroz) #132(2) (UFPR; dry).

**Paratypes. Brazil: AMAZONAS:** 6 m$, 6 f£, 8 immatures, same as holotype but (MMBC, NHMB, UFPR; dry, slide, 70% ethanol; NMB-PSYLL0007892, NMB-PSYLL0008322 [LSMelman-11], NMB-PSYLL0007683, NMB-PSYLL0008263, NMB-PSYLL0007895 [LSMelman-11], NMB-PSYLL0007896 [LSMelman-11], NMB-PSYLL0007897); 2 m$, 8 f£, 12 immatures, same but Fazenda Rio Urubu, S2.4300/4744, W59.5617/5683, 50–100 m, 23– 24.iv.2014, *Miconia phanerostila* (D. Burckhardt & D.L. Queiroz) #131(4) (dry, slide, NHMB; 70% ethanol; NMB-PSYLL0007884, NMB-PSYLL0007898, NMB- PSYLL0007899, NMB-PSYLL0007900 [LSMelman-11], NMB-PSYLL0007901 [LSMelman-11], NMB-PSYLL0007891); 1 m$, Manaus, Adolpho Ducke Forest Reserve, S2.9645, W59.9202, 100 m, 17.xi.1995, fogging of *Eschweilera pseudodecolorans* (J.C.H. Guerrero, P.O.I. de Lima & G.P.L. Mesquita) #F2-32(1) (INPA; dry).

**Description. *Adult.*** Coloration. Brownish yellow; head and thorax with brown dots. Antenna yellow, scape and pedicel ochreous, segments 4–8 apically and segments 9–10 entirely brown. Mesoscutum with four broad ochreous longitudinal stripes. Forewing (Fig. 17I) with whitish veins and yellow membrane, in cell c+sc and r_1_ rather whitish, with only a few indistinct, irregular pale brown dots; base and apex of pterostigma and apices of veins R+M+Cu, Rs, M_1+2_, M_3+4_, Cu_1a_, Cu_1b_, and A medially with relatively large dark brown spots. Pro- and mesofemora with brown subapical band, pro- and mesotibiae dark yellow, protarsus brown.

Structure. Forewing (Fig. 17I) ovoid, widest slightly distal to the middle, broadly rounded apically; wing apex situated in the middle of margin of cell r_2_; C+Sc relatively evenly curved; pterostigma distinctly narrower than r_1_ cell in the middle, widening to apical two thirds; Rs hardly curved in basal two thirds, distinctly curved to fore margin apically; M longer than M_1+2_ and M_3+4_; Cu_1a_ relatively evenly curved, ending proximal of M fork; surface spinules present in all cells, leaving narrow spinule-free stripes along the veins, forming hexagons of a single row of spinules, spinules absent from basal half of cell c+sc. Hindwing with ungrouped (5–6) costal setae, sometimes with distinctly grouped (3 + 2–3) costal setae. Metatibia bearing 7–10 grouped apical spurs, arranged as 4 + 3–6, anteriorly separated by 1 bristle.

Terminalia (Fig. 29G–L). Male. Proctiger weakly produced posteriorly in basal half; densely covered with very long setae in apical two thirds. Subgenital plate, in lateral view, subtriangular; dorsal margin weakly concave; posterior margin irregularly convex; with sparse long setae. Paramere, in lateral view, narrowly irregularly lanceolate, with mostly subparallel margins in basal two thirds, irregularly narrowing to apex; apex sclerotised, in lateral view, narrow, subacute, directed slightly anteriad; in dorsal view, apex narrow, acute, directed inwards and slightly anteriad, bearing a small sclerotised tooth; outer face with sparse, moderately long setae laterally in apical two thirds; inner face with dense, relatively short setae in apical half and very long setae basally; posterior margin with very long setae. Proximal segment of aedeagus with apical part moderately subdivided. Distal segment of aedeagus almost straight in basal half with dorsal margin weakly sinuate; ventral process situated slightly distal of the middle of the segment, short, in lateral view, narrow, simple, lamellar, lacking lateral lobes, in dorsal view, ventral process ovoid, broadest medially, rounded apically; apical dilation, in both lateral and dorsal views, moderately broad, slightly widening towards broadly rounded apex, dorsal part of apical dilation membranous and produced at base; sclerotised end tube moderately long and almost straight. – Female terminalia cuneate; densely covered with long setae. Dorsal margin of proctiger, in lateral view, slightly concave distal to the circumanal ring, apical part straight, apex pointed; in dorsal view, apex blunt; circumanal ring, in dorsal view, cruciform. Subgenital plate, in lateral view, strongly narrowed in apical third, apex pointed; in ventral view, apex blunt.

***Fifth instar immature.*** Coloration. Pale yellow; cephalothoracic sclerite, antenna, wing pads, legs and caudal plate pale to bright brown.

Structure. Eye with one short simple ocular seta dorsally. Antennal segments with following numbers of pointed sectasetae: 1(0), 2(2: one large and one small), 3(0), 4(2), 5(0),

6(2), 7(1), 8(1), 9(0), 10(0). Forewing pad with five marginal pointed sectasetae; hindwing pad with two marginal pointed sectasetae; both pads lacking sectasetae dorsally.

Metatibiotarsus relatively long; tarsal arolium broadly triangular apically, > 1.5 times as long as claws. Abdomen lacking lateral pointed sectasetae anterior to caudal plate. Caudal plate with anterior margin remote from anterior margin of extra pore fields; with 3–4 (1 + 2 or 3) long and narrow pointed sectasetae on either side laterally, and three long and narrow pointed sectasetae subapically on either side of circumanal ring dorsally. Extra pore fields forming continuous outer and inner bands, consisting of large oval and rounded patches; outer band long medially, end pointing outwards. Circumanal ring large.

**Host plant.** *Miconia manauara* R. Goldenb., Caddah & Michelang., *M. phanerostila* Pilg. (Melastomataceae).

**Distribution.** Brazil (AM).

**Derivation of name.** Dedicated to Professor Renato Goldenberg, leading systematist of Melastomataceae, for his invaluable help in identifying many specimens of this taxonomically difficult plant family.

**Comments.** Within the *trematos*-group, *M. goldenbergi* sp. nov. resembles *M. clavata* sp. nov. and *M. dimorpha* sp. nov. in the narrowly lanceolate paramere. It differs from the last species in light body colour and the asymmetric (versus symmetric) paramere. *M. goldenbergi* differs from *M. clavata* in the much shorter ventral process of the distal aedeagal segment, and the straight (versus upturned) apex of the female proctiger.

### 35 Melanastera clavata sp. nov

(Figs 2B, 11F, 17J, 29M–R)

**Type material. Holotype m$: Brazil: MINAS GERAIS:** Vargem Bonita, Parque Nacional da Serra da Canastra, Cachoeira Casca d’Anta, plateau, S20.2976/2986, W46.5195/5289, 860 m, 5.ix.2014, *Miconia sellowiana* (D. Burckhardt & D.L. Queiroz) #144(1) (UFPR; dry).

**Paratypes. Brazil: MINAS GERAIS:** 5 m$, 5f£, 3 immatures, same as holotype but (MMBC, NHMB, UFPR; dry, slide, 70% ethanol; NMB-PSYLL0007812, NMB- PSYLL0007813, NMB-PSYLL0007681, NMB-PSYLL0007824 [LSMelcla-79], NMB-PSYLL0007825, NMB-PSYLL0007826 [LSMelcla-79]); 3 m$, 4f£, 3 immatures, same but Cachoeira Casca d’Anta, around park entrance, S24.8541/8573, W48.6982/7121, 850–860 m, 4–8.ix.2014, *Miconia pusilliflora* (D. Burckhardt & D.L. Queiroz) #141(20) (MMBC, NHMB; dry, slide, 70% ethanol; NMB-PSYLL00003133, NMB-PSYLL0007821, NMB- PSYLL0007822 [LSMelcla-79], NMB-PSYLL0007823 [LSMelcla-79], NMB- PSYLL00003130, NMB-PSYLL00003129).

**Description. *Adult.*** Coloration. Orange. Antenna yellow, apices of segments 4–9 and entire segment 10 brown. Thorax with indistinct light brown dots and patches; mesoscutum with four broad and one narrow median brown longitudinal stripes. Forewing (Fig. 17J) amber-coloured, with irregular, sparse brown dots mostly absent from wing base; apices of veins Rs, M_1+2_, M_3+4_, Cu_1a_ and Cu_1b_ inconspicuously brown.

Structure. Forewing (Fig. 17J) oblong-oval, broadest in the middle, narrowly, slightly unevenly rounded apically; wing apex situated in cell r_2_ near apex of M_1+2_; C+Sc relatively straight in basal two thirds, moderately curved in distal third; pterostigma narrower in the middle than r_1_ cell, widening to apical third; Rs mostly straight, strongly curved to fore margin in apical quarter; M longer than M_1+2_ and M_3+4_; Cu_1a_ irregularly convex, ending almost at M fork; surface spinules present in all cells, leaving narrow spinule-free stripes along the veins, forming hexagons mostly of a double row of spinules; spinules absent from base of cell c+sc. Hindwing with 4–5 + 3 grouped costal setae. Metatibia with 8–10 grouped apical spurs, arranged as 4 + 4–6, anteriorly separated by 2–3 bristles.

Terminalia (Fig. 29M–R). Male. Proctiger weakly produced posteriorly; densely covered with long setae in apical half. Subgenital plate subglobular; dorsal margin weakly curved; posterior margin slightly curved; with long setae, mostly posteriorly. Paramere, in lateral view, asymmetrically lanceolate; apex, in lateral view, blunt, directed slightly anteriad, in dorsal view, apex blunt, directed upwards and slightly inwards, lacking distinct sclerotised tooth; both outer and inner faces with dense, moderately long setae; posterior margin with longer setae. Proximal segment of aedeagus with apical part strongly subdivided. Distal segment of aedeagus with dorsal margin slightly sinuate in basal half; ventral process situated slightly distal to the middle of the segment, in both lateral and dorsal views, ventral process moderately long, simple, lacking lateral lobes, club-shaped, narrow basally, broadest subapically; apical dilation, in both lateral and dorsal views, moderately broad, oblong, subparallel-sided, with truncate apex; sclerotised end tube short and curved. – Female terminalia cuneate; densely covered with setae. Dorsal margin of proctiger, in lateral view, sinuate, apex pointed, slightly upturned; in dorsal view, apex blunt; circumanal ring, in dorsal view, distinctly cruciform. Subgenital plate, in lateral view, pointed apically; in ventral view, apex blunt.

***Fifth instar immature.*** Coloration. Yellow to orange; antenna pale brown, gradually becoming darker towards apex; cephalothoracic sclerite, wing pads, legs and caudal plate brown.

Structure. Body dorsum and venter of caudal plate densely covered with minute clavate and narrowly lanceolate setae. Eye with one short simple ocular seta dorsally. Antennal segments with following numbers of pointed sectasetae: 1(0), 2(1), 3(0), 4(2), 5(0), 6(2), 7(1), 8(1), 9(0), 10(0). Forewing pad with five marginal pointed sectasetae; hindwing pad with two marginal and 0–1 dorsal pointed sectasetae; both pads densely covered with many minute clavate and several narrowly lanceolate setae dorsally. Metatibiotarsus short; tarsal arolium broadly fan-shaped apically, 1.2 times as long as claws. Abdomen with two lateral pointed sectasetae laterally anterior to caudal plate. Caudal plate with anterior margin relatively distant from anterior margin of extra pore fields; with five (2 + 3) pointed sectasetae situated on two tubercles on either side laterally, and three pointed sectasetae subapically on either side of circumanal ring dorsally. Extra pore fields forming continuous outer and inner bands, consisting of small oval and round patches; outer band relatively short medially, end pointing outwards. Circumanal ring large.

**Host plants.** *Miconia pusilliflora* (DC.) Naudin, *M. sellowiana* Naudin (Melastomataceae).

**Distribution.** Brazil (MG).

**Derivation of name.** From the Latin adjective *clavatus* = with a club, referring to the shape of the ventral process of the distal segment of aedeagus and the presence of many minute clavate setae on dorsum of the immatures.

**Comments.** Within the *trematos*-group, *M. clavata* sp. nov. resembles *M. goldenbergi* sp. nov. in the narrowly, asymmetrically lanceolate paramere. The two species differ from each other as indicated under the description of *M. goldenbergi*.

### 36 Melanastera tibialis sp. nov

(Figs 11G, 18A, 29S–X)

**Type material. Holotype m$: Brazil: SÃO PAULO:** São José do Barreiro, Parque Nacional da Serra da Bocaina, Cachoeira das Posses, S22.7745, W44.6019, 1220 m, 7.iv.2019, *Tibouchina* sp. (D. Burckhardt & D.L. Queiroz) #319(2) (UFPR; dry).

**Paratypes. Brazil: SÃO PAULO:** 1 m$, 2 f£, 4 immatures, 5 skins, same as holotype but (NHMB; slide, 70% ethanol; NMB-PSYLL0008093, NMB-PSYLL0008153 [LSMeltib-104], NMB-PSYLL0008154 [LSMeltib-104], NMB-PSYLL0008155 [LSMeltib-104], NMB- PSYLL0008156, NMB-PSYLL0008157).

**Description. *Adult.*** Coloration. Yellow; head and thorax with dark brown dots. Antennal segments 3–8 apically and segments 9–10 entirely dark brown. Mesoscutum with four broad orange longitudinal stripes, margined by dark brown dots, and one row of brown dots in the middle. Forewing (Fig. 18A) yellow with scattered brown dots; apices of pterostigma, veins Rs, M_1+2_, M_3+4_, Cu_1a_ and Cu_1b_ brown. Legs yellow to brown, apical tarsal segments brown. Abdomen and terminalia orange.

Structure. Forewing (Fig. 18A) oblong-oval, widest slightly in the middle, relatively evenly rounded apically; wing apex situated in cell r_2_ near apex of M_1+2_ apex; C+Sc distinctly curved in distal third; pterostigma slightly wider than cell r_1_ in the middle, weakly expanding to apical third; Rs almost straight in basal two thirds, weakly curved to fore margin apically; M longer than M_1+2_ and M_3+4_; Cu_1a_ weakly irregularly curved, ending at M fork; surface spinules present in all cells, leaving narrow spinule-free stripes along the veins, forming hexagons of a double row of spinules, spinules absent from basal half of cell c+sc. Hindwing with 3–4 + 3 grouped costal setae. Metatibia bearing a posteriorly open crown of 8–11 ungrouped apical spurs.

Terminalia (Fig. 29S–X). Male. Proctiger, tubular, weakly produced posteriorly; densely covered with long setae in apical two thirds. Subgenital plate, in lateral view, subglobular; dorsal margin strongly curved; posterior margin slightly concave; with densely spaced long setae. Paramere, in lateral view, asymmetrically lanceolate; apex, in lateral view, blunt, directed upwards, in dorsal view, paramere relatively broad subapically, apex sclerotised and pointed, directed inwards; outer and inner faces with dense, moderately long setae; posterior margin with long setae. Proximal segment of aedeagus with apical part moderately subdivided. Distal segment of aedeagus with weakly convex ventral margin and weakly sinuate dorsal margin in basal half; ventral process situated slightly distal of the middle of the segment, simple, lacking lateral lobes, in lateral view, ventral process narrow, lamellar, in dorsal view, ventral process narrower than apical dilation, spatulate, constricted medially, broadest subapically, rounded apically); apical dilation broad, in lateral view, slightly widening towards broadly rounded apex, broadest in apical third, dorsal part of apical dilation membranous and expanded at base, in dorsal view, apical dilation broadest medially, narrowly rounded apically; sclerotised end tube short and weakly curved. – Female terminalia cuneate; densely covered with setae. Dorsal margin of proctiger, in lateral view, with a small convexity distal to circumanal ring, straight in apical third, apex blunt; in dorsal view, apex subacute; circumanal ring, in dorsal view, distinctly cruciform. Subgenital plate, in lateral view, abruptly narrowing to apex in apical half, apex pointed; in ventral view, apex blunt.

***Fifth instar immature.*** Coloration. Brownish yellow; antenna yellow to pale brown, gradually becoming darker towards apex, segments 6–10 dark brown.

Structure. Eye with one short simple ocular seta dorsally. Antennal segments with following numbers of pointed sectasetae: 1(0), 2(1), 3(0), 4(2), 5(0), 6(2), 7(1), 8(1), 9(0), 10(0). Each forewing pad with 6–7 pointed sectasetae marginally and ca. 20 pointed sectasetae or lanceolate setae dorsally; hindwing pad with 2–3 ponted sectasetae marginally and several pointed sectasetae and lanceolate setae dorsally. Metatibiotarsus long; tarsal arolium broadly fan-shaped apically, 1.5 times as long as claws. Abdominal tergites anterior to caudal plate with three rows of 4, 5–6 and 6 pointed sectasetae dorsally, respectively. Caudal plate dorsally with one row of 11 pointed sectasetae along anterior margin, a row of 10 pointed sectesetae submedially, two groups of 3 pointed sectasetae on each side in the middle and one group of 5 pointed sectasetae on each side subapically, anterior to circumanal ring. Caudal plate with anterior margin relatively remote from anterior margin of extra pore fields. Extra pore fields forming continuous outer and inner bands, consisting of large oval and rounded patches; outer band relatively short medially, end pointing backwards. Circumanal ring small.

**Host plant.** *Tibouchina* sp. (Melastomataceae).

**Distribution.** Brazil (SP).

**Derivation of name.** From the Latin noun *tibia =* shin, referring to the long metatibia.

**Comments.** *Melanastera tibialis* sp. nov. and *M. pleromatos* sp. nov. are morphologically similar and differ from the other members of the *trematos*-group in the ungrouped (versus grouped) metatibial spurs. *M. tibialis* differs from *M. pleromatos* as follows: MT > 0.4, MT/HW > 0.8 (versus MT < 0.4, MT/HW < 0.8), a slightly more (versus less) produced anterior margin of the paramere and the female proctiger bearing (versus lacking) a small swelling distal to the circumanal ring. The uncorrected p-distance between the two species

(*M. tibialis*: sample DB-DLQ#319(2) and *M. pleromatos*: sample DB-DLQ#187(3)) is 15.2% for *COI* and 16.2% for *cytb*.

### 37 Melanastera pleromatos sp. nov

(Figs 11H, 18B, 30A–F)

**Type material. Holotype m$: Brazil: RIO GRANDE DO SUL:** Cambará do Sul, Parque Nacional de Aparados da Serra, Itaimbezinho, park headquarters, S29.1583, W50.0800, 910 m, 25.i.2016, *Pleroma raddianum* (D. Burckhardt & D.L. Queiroz) #187(3) (UFPR; dry).

**Paratypes. Brazil: PARANÁ:** 2 m$, 1 f£, Antonina, Usina Parigot de Souza, S25.2557, W48.7787, 720 m, 19.vii.2017, *Pleroma raddianum* (I. Malenovský) #250 (MMBC; 70% ethanol); 5 m$, 7 f£, 1 immature, same but *Pleroma raddianum* (D. Burckhardt & D.L. Queiroz) #250(4) (NHMB; 70% ethanol; NMB-PSYLL0007500); 2 f£, same but (D. Burckhardt & D.L. Queiroz) #251 (NHMB; 70% ethanol; NMB-PSYLL0007501); 1 m$, 8 f£, Bocaiuva do Sul, BR-476 km 86, S25.1283/1900, W49.085/1283, 900–1080 m, 18.iv.2013, *Pleroma sellowianum*, remnants of Atlantic forest along plantations (D. Burckhardt & D.L. Queiroz) #104(6) (NHMB; 70% ethanol; NMB-PSYLL0007482); 5 m$, 6 f£, same but BR- 476 km72, S25.0783, W49.0916, 1140 m, 21.iv.2013, *Pleroma sellowianum* (D. Burckhardt & D.L. Queiroz) #108(1) (NHMB; 70% ethanol; NMB-PSYLL0007487); 3 m$, 1 f£, Cerro Azul, BR-476 km 69, S25.0685, W49.0877, 1080 m, 18–19.iv.2013 (D. Burckhardt & D.L. Queiroz) #106(6) (NHMB; slide, 70% ethanol; NMB-PSYLL0007484, NMB- PSYLL0007539); 1 f£, 1 immature, Colombo, Campus da Embrapa, S24.8526, W48.7358, 900 m, 12.ii.2014 (D.L. Queiroz) #591 (NHMB; 70% ethanol; NMB-PSYLL0007504); 1 m$, 1 f£, 1 immature, same but (D.L. Queiroz) #595 (NHMB; 70% ethanol; NMB-PSYLL0007505); 5 m$, 15 f£, 5 immatures, same but (D.L. Queiroz) #596 (NHMB; 70% ethanol; NMB-PSYLL0007506); 8 m$, 12 f£, 20 immatures, Curitiba, Parque Barigui, S25.4266, W49.3133, 910 m, 9.ii.2016, *Pleroma raddianum* (D. Burckhardt & D.L. Queiroz) #196(9) (NHMB; slide, 70% ethanol; NMB-PSYLL0007498, NMB-PSYLL0007535, NMB-PSYLL0007536); 1 m$, 4 f£, same but Pedreira Paulo, S25.3850, W49.2783, 950 m, 24.x.2012, *Pleroma raddianum* (D. Burckhardt & D.L. Queiroz) #47(3) (NHMB; 70% ethanol; NMB-PSYLL0007476); 1 m$, same but Parque Barreirinha, S25.3629, W49.2612, 1.xii.2012 (D. Burckhardt & D.L. Queiroz) #82 (NHMB; 70% ethanol; NMB- PSYLL0007486); 6 m$, 9 f£, same but Parque Barigui, S254269, W493134, 910 m, 4.xii.2012, *Pleroma raddianum* (D. Burckhardt & D.L. Queiroz) #85(6) (NHMB; 70% ethanol; NMB-PSYLL0007479); 4 m$, 5 f£, 2 immatures, same but Bosque Zaninelli, S254269, W493134, 950 m, 11.ii.2013, *Pleroma raddianum* (D. Burckhardt & D.L. Queiroz) #91(2) (NHMB; 70% ethanol; NMB-PSYLL0007480); 2 m$, 4 f£, same but Jardim Botânico, S25.4416, W49.2383, 910 m, 2.iv.2013, *Tibouchina* sp. (D. Burckhardt & D.L. Queiroz) #97(3) (NHMB; 70% ethanol; NMB-PSYLL0007481); 9 m$, 8 f£, Matinhos, Rodovia Elisio Pereira Alves Filho, S25.7100/7916, W48.5616/5766, 0–10 m, 29.xi.2012, *Pleroma raddianum* (D. Burckhardt & D.L. Queiroz) #81(2) (NHMB; 70% ethanol; NMB- PSYLL0007477); 1 f£, Morretes, Recanto Engenheiro Lacerda PR-410, S25.3333/3337, W48.9009/9015, 780–870 m, 13.ix.2011, *Pleroma raddianum* (D. Burckhardt & D.L. Queiroz) #2(1) (NHMB; 70% ethanol; NMB-PSYLL0007414); 13 m$, 23 f£, 3 immatures, 4 skins, same but Estação Marumbi, S25.4484/4492, W48.8901/8915, 230–240 m, 14.ix.2011, *Pleroma raddianum* (D. Burckhardt & D.L. Queiroz) #5(3) (MMBC, NHMB; dry, slide, 70% ethanol; NMB-PSYLL0007471, NMB-PSYLL0007472, NMB-PSYLL0007526 [LSMeltibo- 6], NMB-PSYLL0007529 [LSMeltibo-6], NMB-PSYLL0007530, NMB-PSYLL0007485 [voucher 5(4)]); 5 m$, 4 f£, same but Floresta Marumbi, trail to hotel, S25.4459, W48.8873, 170 m, 14.ix.2011, *Pleroma raddianum* (D. Burckhardt & D.L. Queiroz) #6(2) (NHMB; 70% ethanol; NMB-PSYLL0007473); 5 m$, 4 f£, 1 immature, same but BR-277, Cachoeira, S25.5748, W48.8279, 700 m, 28.xi.2012, *Pleroma raddianum* (D. Burckhardt & D.L. Queiroz) #80(2) (NHMB; 70% ethanol; NMB-PSYLL0007478); 1 adult, same but PR-410, Recanto Rio Cascata, S25.3340, W48.8984, 800 m, 26.vii.2017, *Pleroma raddianum*, Atlantic forest (L. Serbina & I. Malenovský) #DIC1 (NHMB; slide, NMB-PSYLL0007531 [voucher 6-1]); 1 f£, same but (L. Serbina & I. Malenovský) #DIC2+3 (MMBC; 70% ethanol); 1 f£, same but (L. Serbina & I. Malenovský) #DIC3 (MMBC; 70% ethanol); 8 m$, 26 f£, Piraquara, Parque Estadual do Marumbi, 25.1566/1600, W48.9745/9916, 890–1170 m, 23– 24.iv.2013, *Pleroma sellowianum* (D. Burckhardt & D.L. Queiroz) #109(1) (NHMB; 70% ethanol; NMB-PSYLL0007497); 12 m$, 4 f£, same but Marumbi, Estação Carvalho, S27.4727, W48.3781, 1000 m, 20.xi.2013, *Miconia sellowiana* (D.L. Queiroz) #599(4) (NHMB; 70% ethanol; NMB-PSYLL0007503, NMB-PSYLL0007532 [voucher 5-9]); 1 f£, Piraquara, Road to Marumbi, S25.4962, W48.9817, 990 m, 21.xi.2013 (D.L. Queiroz) #600 (NHMB; 70% ethanol; NMB-PSYLL0007507); 7 m$, 10 f£, 4 immatures, 1 skin, same but Marumbi, Road to Estação Carvalho, S25.4962, W48.9817, 990 m, 11.ii.2015 (D.L. Queiroz) #668 (NHMB; 70% ethanol; NMB-PSYLL0006044); 5 m$, 4 f£, same but S27.4727, W48.3781, 1000 m, 11.ii.2015 (D.L. Queiroz) #670 (NHMB; 70% ethanol; NMB-PSYLL0006050); 1 m$, 1 f£, same but Marumbi, Estação Carvalho, S27.4715, W48.3799, 1050 m, 11.ii.2015 (D.L. Queiroz) #671 (NHMB; 70% ethanol; NMB-PSYLL0006040 [male voucher]); 6 m$, 7 f£, 1 immature, Piraquara, BR-277 km50–55, S25.5733, W48.9900, 940 m, 28.xi.2012, *Pleroma raddianum* (D. Burckhardt & D.L. Queiroz) #79(4) (NHMB; 70% ethanol; NMB-PSYLL0007474); 1 m$, 8 f£, Tibagi, Parque Estadual Guartelá, S24.5623/5666, W50.2589, 920–950 m, 23–25.vi.2015, *Pleroma raddianum* (D. Burckhardt & D.L. Queiroz) #171(2) (NHMB; 70% ethanol; NMB-PSYLL0007489); 4 m$, 7 f£, Tibagi, Parque Estadual Guartelá, S24.5683, W50.2553, 940 m, 10–12.vii.2017, *Pleroma raddianum* (D. Burckhardt & D.L. Queiroz) #245(5) (NHMB; 70% ethanol; NMB-PSYLL0007499); 3 m$, 5 f£, 1 skin, Tibagi, Parque Estadual Guartelá, S24.5683, W50.2553, 940 m, 10.vii.2017, Cerrado vegetation (L. Serbina) #DIC4 (MMBC; 70% ethanol); 2 m$, 5 f£, Tunas do Paraná, BR-476 km60, S25.0083, W49.0800, 900 m, 18.iv.2013, *Miconia* sp. (D. Burckhardt & D.L. Queiroz) #105(2) (NHMB; slide, 70% ethanol; NMB-PSYLL0007483, NMB- PSYLL0007537); 9 m$, 12 f£, 6 immatures, 2 skins, Tunas do Paraná, Parque das Lauraceas, S25.0688, W49.0888, 890 m, 8.viii.2013 (D.L. Queiroz) #556 (NHMB; slide, 70% ethanol; NMB-PSYLL0007502, NMB-PSYLL0007525 [voucher 6-10]); 2 m$, 5 f£, Ventania, road BR-153, around 10 km from Ventania, 241,783, W50.2166, 870 m, 18.ix.2014, *Pleroma raddianum* (D. Burckhardt & D.L. Queiroz) #149(3) (NHMB; 70% ethanol; NMB- PSYLL0007488); 3 m$, 4 f£, 40 immatures, 1 skin, Morretes, PR-410, Recanto Rio Cascata, S25.3340, W48.8984, 800 m, 26.vii.2017, *Pleroma raddianum*, Atlantic forest (L. Serbina & I. Malenovský) #DIC2, 1 (MMBC; 70% ethanol).—**Rio Grande do Sul:** 6 m$, 16 f£, 10 immatures, 1 skin, Passo Fundo, S28.2288, W52.4061, 670 m, 7.ix.2012, *Pleroma sellowianum* (A.L. Marsaro Jr.) (NHMB; 70% ethanol; NMB-PSYLL0007510); 9 m$, 9 f£, 15 immatures, same but S28.2288, W52.4061, 670 m, 14.xi.2012 (A.L. Marsaro Jr.) (NHMB; 70% ethanol; NMB-PSYLL0007509); 18 m$, 27 f£, 5 immatures, Cambará do Sul, Parque Nacional de Aparados da Serra, Itaimbezinho, park headquarters, S29.1583, W50.0800, 910 m, 25.i.2016, *Pleroma raddianum* (D. Burckhardt & D.L. Queiroz) #187(3) (BMNH, MHNG, MMBC, NHMB, UFPR, USNM; dry, slide, 70% ethanol; NMB-PSYLL0007490–NMB- PSYLL0007496, NMB-PSYLL0007475, NMB-PSYLL0007560, NMB-PSYLL0007527 [voucher 6-3], NMB-PSYLL0007528 [voucher 6-4], NMB-PSYLL0007533 [voucher 6r-2], NMB-PSYLL0007534 [voucher 6r-1], NMB-PSYLL0007538).—**Santa Catarina:** 1 m$, Florianópolis, Costão do Santinho, S11.8339, W55.4996, 170 m, 16.ii.2015 (D.L. Queiroz) #672 (NHMB; 70% ethanol; NMB-PSYLL0007508).

**Description. *Adult.*** Coloration. Head and clypeus yellow with small dark dots on the vertex. Antenna whitish, apices of segments 4–9 and entire segment 10 dark brown. Thorax light orange brown, with scattered dark brown dots; mesopraescutum with two orange patches at the fore margin; mesoscutum with four broad and, in the middle, one narrow orange longitudinal stripes, margined by dark brown dots. Forewing (Fig. 18B) light amber-coloured with sparse, irregular brown dots; base and apex of pterostigma and apices of veins Rs, M_1+2_, M_3+4_, Cu_1a_ and Cu_1b_ brown. Legs yellow, pro- and mesofemora with two transverse brown bands. Abdomens and terminalia yellow.

Structure. Forewing (Fig. 18B) oblong-oval, widest in the middle, unevenly rounded apically; wing apex situated in cell r_2_ near apex of M_1+2_; C+Sc distinctly curved in distal third; pterostigma about as wide as r_1_ cell in the middle, weakly widening to the middle; Rs almost straight in basal two thirds, obliquely curved to fore margin apically; M longer than M_1+2_ and M_3+4_; Cu_1a_ relatively straight in basal two thirds, ending slightly proximal of M fork; surface spinules present in all cells, leaving narrow spinule-free stripes along the veins, forming hexagons of a double row of spinules, spinules absent from base of cell c+sc. Hindwing with 2 + 3–4 grouped costal setae. Metatibia short, bearing a posteriorly open crown of 8–9 ungrouped apical spurs.

Terminalia (Fig. 30A–F). Male. Proctiger tubular, weakly produced posteriorly; densely covered with short setae in apical two thirds. Subgenital plate irregularly ovoid; dorsal margin strongly curved in distal third; posterior margin irregularly convex; with densely spaced long setae. Paramere, in lateral view, irregularly lanceolate; apex, in lateral view, blunt, directed upwards and slightly anteriad, in dorsal view, paramere broad subapically, apex sclerotised and pointed, directed inwards; outer face with long setae mostly in apical half; inner face with dense moderately long setae; posterior margin with longer setae. Proximal segment of aedeagus with apical part moderately subdivided. Distal segment of aedeagus almost straight in basal half; ventral process situated slightly distad of the middle of the segment length, simple, lacking lateral lobes, in lateral view, ventral process narrow, lamellar, in dorsal view, ventral process narrower than apical dilation, slightly widening to apex, broadest apically; apical dilation broad, in lateral view, slightly widening towards broadly rounded apex, broadest in apical third, dorsal part of apical dilation membranous and expanded at base, in dorsal view, apical dilation broadest medially, rounded apically; sclerotised end tube short and weakly sinuate. – Female terminalia cuneate; densely covered with setae. Dorsal margin of proctiger, in lateral view, straight, apex pointed; in dorsal view, apex blunt; circumanal ring, in dorsal view, cruciform. Subgenital plate, in lateral view, irregularly narrowing to apex, apex pointed; in ventral view, apex truncate.

***Fifth instar immature.*** Coloration. Pale yellow; antenna gradually becoming darker towards apex, segments 9–10 brown; cephalothoracic sclerite, wing pads, legs and caudal plate yellow to pale brown. Structure. Eye with one short simple ocular seta dorsally. Antennal segments with following numbers of pointed sectasetae: 1(0), 2(2 – one large, one small), 3(0), 4(2), 5(0), 6(2), 7(1), 8(1), 9(0), 10(0). Forewing pad with five marginal and 1–2 dorsal pointed sectasetae; hindwing pad with two marginal and one dorsal pointed sectasetae. Metatibiotarsus short; tarsal arolium broadly triangular apically, 1.2 times as long as claws. Abdomen with four (2+2) pointed sectasetae laterally on either side anterior to caudal plate. Caudal plate with anterior margin relatively remote from anterior margin of extra pore fields; with five pointed sectasetae situated on weakly pronounced tubercles on either side laterally, and 3–4 pointed sectasetae subapically on either side of circumanal ring side dorsally. Extra pore fields forming continuous outer and inner bands, consisting of small oval patches; outer band relatively short medially, end pointing outwards. Circumanal ring small.

**Host plants.** *Pleroma raddianum* (DC.) Gardner, *P. sellowianum* (Cham) P.J.F.Guim. & Michelang. (Melastomataceae).

**Distribution.** Brazil (PR, RS, SC).

**Derivation of name.** Named after its host, *Pleroma*.

**Comments.** *Melanastera pleromatos* sp. nov. is morphologically close to *M. tibialis* sp. nov. The two species differ as indicated under *M. tibialis*.

## The *ingariko*-group

**Description. *Adult*.** Head, in lateral view, inclined at ≥ 45° from longitudinal axis of body (Fig. 7F). Vertex trapezoidal, covered with distinct imbricate microsculpture (Figs 11I, J, 12A, C–E) (rugosely imbricate in *M. acuminata* sp. nov.; Fig. 12B) and microscopical setae. Thorax moderately or strongly arched, covered with microscopical setae; metapostnotum usually with distinct longitudinal keel (indistinct in *M. caramambatai* sp. nov.). Forewing (Fig. 18C–I) uniformly coloured, with small brown dots; pterostigma distinctly expanding towards the middle or apical third, longer than 4.0 times as wide; vein R_1_ weakly convex or straight medially. Paramere, in lateral view, irregularly acuminate, falcate or lamellar. Distal segment of aedeagus bearing short ventral process, less than half as long as apical dilation, which is, in dorsal view, wider than the latter.

***Immature.*** Antenna 10-segmented.

**Comments.** The group comprises seven species from Brazil and one species from Panama, viz. *M. longitarsata* (Brown & Hodkinson). The Panamanian species differs from the other group members in the longer metatibia (MT/MF > 1.7 versus ≤ 1.7). Five species develop on Melastomataceae (*Clidemia*, *Miconia*) and two on Annonaceae (*Guatteria*, *Xylopia*).

### 38 Melanastera ingariko sp. nov

(Figs 11I, 18C, 30G–L)

**Type material. Holotype m$: Brazil: RORAIMA:** Uiramutã, Caramambatai, N5.1333/1466, W60.5850/5916, 970–1200 m, 7–14.iv.2015, *Miconia chrysophylla* (D. Burckhardt & D.L. Queiroz) #161(9) (UFPR; dry).

**Paratypes. Brazil: RORAIMA:** 9 m$, 21 f£, 8 immatures, 2 skins, same as holotype but (MMBC, NHMB, UFPR; dry, slide, 70% ethanol; NMB-PSYLL0007894, NMB- PSYLL0007977, NMB-PSYLL0007902 [LSMeling-14], NMB-PSYLL0007904 [LSMeling- 14], NMB-PSYLL0007905 [LSMeling-14], NMB-PSYLL0007906 [LSMeling-14], NMB- PSYLL0007907 [LSMeling-14], NMB-PSYLL0007909 [LSMeling-14], NMB- PSYLL0007910, NMB-PSYLL0007911, NMB-PSYLL0007912).

**Description. *Adult.*** Coloration. Straw-coloured; head and thorax with dark brown dots.

Antenna pale yellow, scape and pedicel brownish, apices of segments 3–8 and entire segments 9–10 brown. Mesopraescutum with two brown patches at fore margin; mesoscutum with four broad, brown longitudinal stripes, often margined by brown dots, and one or two median rows of dots. Forewing (Fig. 18C) pale ochreous with sparse irregular brown dots becoming darker and denser towards wing apex; apices of pterostigma and veins Rs, M_1+2_, M_3+4_, Cu_1a_ and Cu_1b_ dark brown. Metacoxa brown, femora with dark patches, pro- and mesotarsi brown. Abdomen yellow to brown, darker ventrally. Terminalia yellow to brown, female terminalia with darker apex. Abdominal tergites partly, sternites entirely dark brown. Male subgenital plate ventrally and female proctiger in distal third, dark brown.

Structure. Forewing (Fig. 18C) oval, widest in the middle, unevenly rounded apically; wing apex situated in cell r_2_ near M_1+2_ apex; C+Sc relatively evenly curved; pterostigma weakly widening to apical third; Rs almost straight in basal two thirds, weakly curved to fore margin apically; M longer than M_1+2_ and M_3+4_; Cu_1a_ irregularly curved, ending at M fork; surface spinules present in all cells, leaving narrow spinule-free stripes along the veins in apical part, forming hexagons of a single row of spinules, spinules absent from base of cell c+sc. Hindwing with 6 ungrouped costal setae. Metatibia bearing 7 grouped apical spurs, arranged as 3–4 + 3–4, anteriorly separated by 3–4 bristles.

Terminalia (Fig. 30G–L). Male. Proctiger relatively robust; densely covered with long setae in apical two thirds. Subgenital plate ovoid; dorsal margin strongly curved in apical third; posterior margin weakly convex; sparsely covered with long setae. Paramere, in lateral view, irregularly acuminate; apex narrow, subacute, weakly sclerotised, directed upwards and slightly inwards; in dorsal view, apex narrow, subacute, directed inwards, lacking a distinct sclerotised tooth; both outer and inner faces with dense, moderately long setae; posterior margin with long setae. Proximal segment of the aedeagus apically strongly subdivided. Distal segment of aedeagus straight in basal two thirds, strongly sclerotised dorsally; ventral process situated slightly distal of the middle of the segment, strongly sclerotised, short, simple, forming obtuse lobe directed apicad, in lateral view, in continuation of the basal part of the segment, in dorsal view, widened to apex which is irregularly truncate; apical dilation large, in lateral view, vaguely bean-shaped, constricted medially, broadly and unevenly rounded apically, in dorsal view, apical dilation oblong-obovate, widest in apical quarter, irregularly rounded apically; sclerotised end tube inserted at midlength of apical dilation, short and sinuate. – Female terminalia cuneate. Dorsal margin of proctiger, in lateral view, sinuate, apex upturned, pointed; in dorsal view, apex subacute; circumanal ring, in dorsal view, vaguely cruciform. Subgenital plate, in lateral view, irregularly narrowing to apex, pointed; in ventral view, apex blunt.

***Fifth instar immature.*** Coloration. Yellow; cephalothoracic sclerite, antenna, wing pads, legs and caudal plate brown.

Structure. Eye with one moderately long simple ocular seta dorsally. Antennal segments with following numbers of pointed sectasetae: 1(0), 2(1), 3(0), 4(2), 5(0), 6(2), 7(1), 8(1), 9(0), 10(0). Forewing pad with 5–6 marginal pointed sectasetae, lacking sectasetae dorsally; hindwing pad with two large and one small marginal pointed sectasetae. Metatibiotarsus short; tarsal arolium broadly triangular apically, 1.2–1.5 times longer than claws. Abdomen with three lateral pointed sectasetae anterior to caudal plate. Caudal plate with anterior margin remote from anterior margin of extra pore fields; with 5–6 (2 + 3–4) pointed sectasetae on either side laterally, and three pointed sectasetae subapically, on either side of circumanal ring dorsally. Extra pore field small, forming continuous outer bands, consisting of small oval patches; outer band short medially, end pointing outwards. Circumanal ring small.

**Host plant.** *Miconia chrysophylla* (Rich.) Urb. (Melastomataceae).

**Distribution.** Brazil (RR).

**Derivation of name.** Dedicated to the Ingarikó people, inhabiting the area where the species was collected. A noun in apposition.

**Comments.** *Melanastera ingariko* sp. nov. resembles *M. brevicauda* sp. nov. in the following characters: the acuminate paramere; the short ventral process of the distal aedeagal segment forming an obtuse lobe which is, in lateral view, in continuation of the basal part of the segment; the strongly sinuate dorsal margin of the female proctiger with an upturned apex. *Melanastera ingariko* differs from *M. brevicauda* in the following characters: forewing membrane with indistinct hexagons of a single row of surface spinules (versus distinct hexagons of a double row of spinules); the more massive male proctiger; the paramere which is narrower in apical half; the distal aedeagal segment with a wider apical dilation, and the sclerotised end tube inserted at midlength (versus subapically) of apical dilation; the shape of the female subgenital plate, in lateral view, with narrower, slightly upturned apex, in ventral view, with narrowly rounded apex, FP/SP < 2.0 (versus > 2.0).

### 39 Melanastera brasiliensis sp. nov

(Figs 11J, 18D, 30M–R)

**Type material. Holotype m$: Brazil: SÃO PAULO:** Santa Maria da Serra, Mina Velha, S22.6823, W48.3049, 450 m, 7.ii.2018, *Xylopia brasiliensis* (D. Burckhardt & D.L. Queiroz) #259(7) (UFPR; dry).

**Paratypes. Brazil: MINAS GERAIS:** 5 m$, 6 f£, Buenópolis, Parque Estadual da Serra do Cabral, headquarters: S17.8649, W44.1837, 590 m, 7.iv.2021, *Grona adscendens*, planted trees and cerrado vegetatio (D. Burckhardt & D.L. Queiroz) #383(2) (NHMB; 70% ethanol; NMB-PSYLL0008489); 1 m$, 6 f£, Diamantina, Parque Biribiri, S18.3712, W43.1099, 1080 m, 14.ix.2021, *Xylopia emarginata* (D.L. Queiroz) #1010(2) (NHMB; 70% ethanol; NMB- PSYLL0008559); 9 m$, 16 f£, same but Cachoeira Sentinela, S18.1863, W43.6143, 1100 m, 11.iv.2021, *Xylopia emarginata*, open cerrado vegetation near river on rocks (D. Burckhardt & D.L. Queiroz) #396(3) (NHMB; 70% ethanol; NMB-PSYLL0008490); 1 m$, 1 f£, São Gonçalo do Rio Preto, Parque Estadual do Rio Preto, Vau das Éguas, S18.0995, W43.3301, 740 m, 15.iv.2021, *Xylopia emarginata*, cerrado vegetation on rocky slope (D. Burckhardt & D.L. Queiroz) #407(2) (NHMB; 70% ethanol; NMB-PSYLL0008492); 4 m$, 8 f£, 1 immature same but Poço do Areia, S18.1310, W43.3379, 770 m, 16.iv.2021, *Xylopia emarginata*, scrub near river (D. Burckhardt & D.L. Queiroz) #409(1) (NHMB; 70% ethanol; NMB-PSYLL0008493); 1 f£, same but Forquilha, S18.1354, W43.3372, 780 m, 16.iv.2021, scrub near river (D. Burckhardt & D.L. Queiroz) #410(0) (NHMB; 70% ethanol; NMB- PSYLL0008494).—**São Paulo:** 7 m$, 7 f£, 13 immatures, 17 skins, same as holotype but (NHMB, UFPR; dry, slide; NMB-PSYLL0007766, NMB-PSYLL0007679, NMB- PSYLL0007680, NMB-PSYLL0007775[LSMelbra-66], NMB-PSYLL0007776, NMB- PSYLL0007777, NMB-PSYLL0007778 [LSMelbra-66], NMB-PSYLL0007779 [LSMelbra- 66]).

**Material not included in type series. Brazil**: **GOIÁS**: 15 m$, 19 f£, 7 immatures, 11 skins, Pirenópolis, Cachoeira do Abade, S15.8088, W48.9190, 955 m, 1.viii.2018 (D.L. Queiroz) #864(3) (NHMB; slide, 70% ethanol; NMB-PSYLL0007786, NMB-PSYLL0007773, NMB-PSYLL0007774 [LSMelbra-98]).—**Minas Gerais:** 4 m$, 4 f£, 8 immatures, Buenópolis, Parque Estadual da Serra do Cabral, Marcos Teixeira, S17.9035, W44.2308, 660 m, 19.ix.2019, *Xylopia emarginata* (D. Burckhardt & D.L. Queiroz) #369(1) (NHMB; 70% ethanol; NMB-PSYLL0007772); 2 m$, 4 f£, same but Passagem d’Anta, S17.9035, W44.2308, 1000 m, 19.ix.2019, *Xylopia emarginata* (D. Burckhardt & D.L. Queiroz) #374(2) (NHMB; 70% ethanol; NMB-PSYLL0007770); 1 m$, 1 f£, 5 immatures, same but Cuba, S17.9035, W44.2308, 1070 m, 22.ix.2019, *Xylopia emarginata* (D. Burckhardt & D.L. Queiroz) #376(2) (NHMB; 70% ethanol; NMB-PSYLL0007771); 6 m$, 9 f£, 3 immatures, 13 eggs, Coromandel, Fazenda Laje, S18.5610, W46.9020, 1060 m, 23.viii.2019, *Xylopia emarginata* (D. Burckhardt & D.L. Queiroz) #346(13) (NHMB; 70% ethanol; NMB- PSYLL0007755, NMB-PSYLL0005585); 10 m$, 20 f£, 8 immatures, Diamantina, Parque Estadual do Biribiri, Cachoeira Sentinela, S18.1886, W43.6229, 1090 m, 14‒15.ix.2019, *Xylopia emarginata* (D. Burckhardt & D.L. Queiroz) #357(2) (NHMB; 70% ethanol; NMB- PSYLL0007768); 18 m$, 17 f£, Diamantina, Parque Estadual do Biribiri, Poço do Estudante, S18.2040, W43.6216, 990 m, 15.ix.2019, *Xylopia emarginata* (D. Burckhardt & D.L. Queiroz) #359(1) (BMNH, MHNG, MMBC, NHMB, USNM; dry, 70% ethanol; NMB- PSYLL0007769, NMB-PSYLL0007971–NMB-PSYLL0007976); 1 m$, 4 f£, 1 immature,

São Gonçalo do Rio Preto, Parque Estadual do Rio Preto, Prainha, S18.1170, W43.2407, 760 m, 11‒12.ix.2019, *Xylopia emarginata* (D. Burckhardt & D.L. Queiroz) #351(5) (NHMB; 70% ethanol; NMB-PSYLL0007756); 1 f£, same but Vau das Éguas, S18.0990, W43.3210, 830 m, 12.ix.2019, *Xylopia emarginata* (D. Burckhardt & D.L. Queiroz) #355(1) (NHMB; 70% ethanol; NMB-PSYLL0007767).

**Description. *Adult.*** Coloration. Orange, head and thorax with dark brown dots. Antenna pale to brownish yellow, scape and pedicel brown, apices of segments 3–9 with and entire segment 10 dark brown. Mesopraescutum with two pale brown patches at the fore margin; mesoscutum with four broad pale brown longitudinal stripes, margined by dark brown dots, and one or two rows of median dots. Forewing (Fig. 18D) light amber-coloured, with irregular, dense dark brown dots; base and apex of pterostigma and apices of veins Rs, M_1+2_, M_3+4_, Cu_1a_ and Cu_1b_ dark brown. Femora with dark brown dots, tibiae and tarsi yellow.

Structure. Forewing (Fig. 18D) oval, broadest in the middle, narrowly rounded apically; wing apex situated in cell r_2_ near apex of apex M_1+2_; C+Sc distinctly curved in distal quarter; pterostigma as wide as or slightly narrower in the apical third than cell r_1_, weakly widening to apical third; Rs fairly straight in basal two thirds, weakly curved to fore margin apically; M longer than M_1+2_ and M_3+4_; Cu_1a_ weakly, relatively evenly curved, ending slightly proximal of M fork; surface spinules present in all cells, leaving narrow spinule-free stripes along the veins, forming hexagons of a single row of spinules, spinules sparse in basal third of cell c+sc. Hindwing with 3–4 + 2–3 grouped costal setae. Metatibia bearing 9 grouped apical spurs, arranged as 3–4 + 5–6, anteriorly separated by 3 bristles.

Terminalia (Fig. 30M–R). Male. Proctiger weakly produced postero-basally; densely covered with long setae in apical two thirds. Subgenital plate subglobular; dorsal margin almost straight; posterior margin irregularly convex; with dense, long setae mostly in apical half. Paramere, in lateral view, broadly acuminate; apex, in lateral view, blunt, directed upwards, in dorsal view, apex subacute, directed upwards and inwards, lacking distinct sclerotised tooth; both outer and inner faces with dense, moderately long setae in apical two thirds; posterior margin with longer setae. Proximal segment of aedeagus with apical part moderately subdivided. Distal segment of aedeagus with dorsal margin strongly sinuate in basal half; ventral process situated slightly distal of the middle of the segment, in lateral view, relatively short and broad, subrectangular, with apex slightly protruding ventrally; in dorsal view, ventral process relatively broad, broadest subapically, with a short, apically truncate median lobe arising from the base ventrally; apical dilation, in lateral view, broad, slightly widening towards broadly rounded apex, with a large membranous sack at dorsal margin basally; in dorsal view, apical dilation relatively narrow, oblong-obovate, apex evenly rounded; sclerotised end tube short and weakly curved. – Female terminalia cuneate; densely covered with setae. Dorsal margin of proctiger, in lateral view, weakly sinuous, apex blunt, in dorsal view, apex pointed; circumanal ring, in dorsal view, cruciform. Subgenital plate, in lateral view, abruptly narrowing in apical half, pointed apically; in ventral view, apex blunt.

***Fifth instar immature.*** Coloration. Light; antenna gradually becoming darker towards apex; cephalothoracic sclerite, wing pads, legs and caudal plate brown to dark brown.

Structure. Eye with one short, simple ocular seta dorsally. Antennal segments with following numbers of pointed sectasetae: 1(0), 2(1), 3(0), 4(2), 5(0), 6(2), 7(1), 8(1), 9(0), 10(0). Forewing pad with 4–5 (usually 5) marginal pointed sectasetae; hindwing pad with 1–2 (usually 2) marginal pointed sectasetae; both pads lacking sectasetae or larger lanceolate setae dorsally. Metatibiotarsus long; tarsal arolium broadly fan-shaped apically, 1.2 times as long as claws. Abdomen with 1–2 lateral pointed sectaseta on either side anterior to caudal plate. Caudal plate with anterior margin remote from anterior margin of extra pore fields; with four (2+2) pointed sectasetae and one smaller lanceolate seta on either side laterally, and three pointed sectasetae subapically, on either side of circumanal ring dorsally. Extra pore fields forming continuous outer and inner bands, consisting of moderate oval patches; outer band relatively long medially, end pointing outwards. Circumanal ring small.

**Host plant.** *Xylopia brasiliensis* Spreng., *X. emarginata* Mart. (Annonaceae).

**Distribution.** Brazil (GO, MG, SP).

**Derivation of name.** Named after one of its hosts, *X. brasiliensis*.

**Comments.** Within the *ingariko*-group, *M. brasiliensis* sp. nov. differs from the other species by its much broader paramere.

The sample from Goiás from *Xylopia sericea* (DLQ#864(3)) and some samples from Minas Gerais from *X. emarginata* (DB-DLQ#346(13), #351(5), #355(1), #357(2), #359(1), #369(1), #374(2), #376(2)) are excluded from the type series for differences in the DNA barcoding sequence (e.g. the uncorrected p-distance between the sample DB-DLQ#259(7) including the holotype and DLQ#864(3) is 1.74% for *COI*, and 3.19% for *cytb*) and/or in the host species. However, as there are no significant morphological differences and low sequence divergence between these samples, they are all referred to the same species.

### 40 Melanastera caramambatai sp. nov

(Figs 1C, 12A, 18E, 30S–X)

**Type material. Holotype m$: Brazil: RORAIMA:** Uiramutã, Caramambatai, N5.1333/1466, W60.5850/5916, 970–1200 m, 7–14.iv.2015, *Miconia* aff. *neourceolata* (D. Burckhardt & D.L. Queiroz) #161(19) (UFPR; dry).

**Paratypes. Brazil: RORAIMA:** 3 m$, 8 f£, 23 immatures, same as holotype but (NHMB; dry, slide, 70% ethanol; NMB-PSYLL0007988, NMB-PSYLL0007937 [LSMelcar-24], NMB-PSYLL0007938, NMB-PSYLL0007939 [LSMelcar-24], NMB-PSYLL0007940, NMB-PSYLL0007941).

**Material not included in type series. Brazil: RORAIMA:** 5 immature, 3 skins, Uiramutã, Caramambatai, N5.1333/1466, W60.5850/5916, 970–1200 m, 7–14.iv.2015, cf. *Virola* (D. Burckhardt & D.L. Queiroz) #161(16) (NHMB; slide, 70% ethanol; NMB-PSYLL0008203, NMB-PSYLL0008098).

**Description. *Adult.*** Coloration. Straw-coloured. Antenna brown, scape and pedicel yellow, segment 3 yellow at base. Mesoscutum with four indistinct broad and one narrow medial brown longitudinal stripes. Forewing (Fig. 18E) yellow with indistinct sparse brown dots mostly in apical two thirds of wing; apices of pterostigma and veins Rs, M_1+2_, M_3+4_, Cu_1a_ and Cu_1b_ brown. Legs yellow.

Structure. Metapostnotum lacking a distinct longitudinal keel. Forewing (Fig. 18E) obovoid, widest in apical third, broadly and evenly rounded apically; wing apex situated in cell r_2_ close to apex of M_1+2_; C+Sc hardly curved; pterostigma in the middle much narrower than cell r_1_, hardly narrowing apical quarter; Rs almost straight in basal two thirds, weakly curved to fore margin apically; M longer than M_1+2_ and M_3+4_; Cu_1a_ irregularly curved, ending proximal of M fork; surface spinules present in all cells, leaving very narrow or no spinule- free stripes along the veins, forming small hexagons of a single row of spinules. Hindwing with 6–7 ungrouped costal setae. Metatibia bearing 6–9 grouped apical spurs, arranged as 3–4

+ 3–5, anteriorly separated by 4–5 bristles.

Terminalia (Fig. 30S–X). Male. Proctiger long, hardly produced posteriorly; densely covered with moderately long setae in apical two thirds. Subgenital plate subtriangular; dorsal margin strongly curved; posterior margin weakly curved; with moderately long setae in ventral half. Paramere, in lateral view, acuminate, weakly curved backwards; apex, in lateral view, blunt, directed upwards, in dorsal view, apex blunt, directed inwards, upwards and slightly anteriad, lacking distinct sclerotised tooth; outer face with dense, long setae in apical half; inner face with dense, short to moderately long setae; posterior margin with long setae. Proximal segment of aedeagus with apical part moderately subdivided. Distal segment of aedeagus almost straight in basal half and with dorsal margin weakly sinuate; ventral process situated slightly distal of the middle of the segment, short, in lateral view, narrow, simple,

tubular, lacking lateral lobes, in dorsal view, ventral process relatively short and distinctly broader than apical dilation, with convex lateral margins, slightly produced medio-apically; apical dilation, in lateral view, moderately broad, slightly narrowing to evenly rounded apex, dorsal part of apical dilation membranous, hardly produced at base, in dorsal view, apical dilation oblong-obovate; sclerotised end tube short and sinuate. – Female terminalia cuneate; relatively densely covered with setae. Dorsal margin of proctiger, in lateral view, straight, apex pointed; in dorsal view, apex blunt; circumanal ring, in dorsal view, vaguely cruciform. Subgenital plate, in lateral view, irregularly narrowing to pointed apex; in dorsal view, apex blunt.

***Fifth instar immature.*** Coloration. Entirely pale yellow; antenna gradually becoming darker towards apex, caudal plate pale brown.

Structure. Body densely covered with minute narrowly lanceolate setae. Eye with one moderately long simple ocular seta dorsally. Antennal segments with following numbers of pointed sectasetae: 1(0), 2(1), 3(0), 4(2), 5(0), 6(2), 7(1), 8(2), 9(0), 10(0). Forewing pad with four marginal and one dorsal pointed sectasetae; hindwing pad with two marginal and one dorsal pointed sectasetae. Tarsal arolium narrowly fan-shaped apically, slightly longer or about as long as claws. Abdominal tergites anterior to caudal plate with three rows of 4, 2 and 5 pointed sectasetae dorsally, respectively. Caudal plate dorsally with one row of five pointed sectasetae along anterior margin, two rows each of four pointed sectesetae submedially, and with two groups of six pointed sectasetae on either side subapically, anterior to circumanal ring. Extra pore fields forming continuous outer and inner bands, consisting of large oval patches; outer band relatively short medially, end pointing backwards. Circumanal ring moderately elongate.

**Host plant and biology.** *Miconia* aff. *neourceolata* Michelang. (Melastomataceae).

Immatures develop on the shoots and axils of young leaves; they are conspicuous by the white wax they secrete (Fig. 1C). Some younger instars [DB-DLQ#161(16)] collected on ?*Virola* sp. (Myristicaceae) may not be conspecific with the specimens collected on *Miconia*.

**Distribution.** Brazil (RR).

**Derivation of name.** Named after the type locality, Caramambatai, near Mt Roraima in north Brazil.

**Comments.** *Melanastera caramambatai* sp. nov. is characterised within the *ingariko*- group by the brown (versus yellow with dark segmental tips) antennal flagellum, the obovate (versus oval) forewing shape and the absence (versus presence) of a distinct longitudinal keel on the metaposnotum.

The sample DB-DLQ#161(16) is not included in the type series. It is only provisionally referred to this species as it contains only young instars collected on ?*Virola* sp. (Myristicaceae).

### 41 Melanastera acuminata sp. nov

(Figs 12B, 18F, 31A–F)

**Type material. Holotype m$: Brazil: AMAZONAS:** Rio Preto da Eva, Embrapa, Fazenda Rio Urubu, S2.4300/4733, W59.5616/5683, 50–100 m, 23–24.iv.2014, *Guatteria* sp. (D.

Burckhardt & D.L. Queiroz) #131(2) (UFPR; dry).

**Paratypes. Brazil: AMAZONAS:** 2 m$, 3 f£, 6 immatures, 1 skin, same as holotype but (NHMB; dry, slide, 70% ethanol; NMB-PSYLL0007731, NMB-PSYLL0007732 [LSMelacu- 9], NMB-PSYLL0007744–NMB-PSYLL0007749).

**Description. *Adult.*** Coloration. Straw-coloured; head and thorax densely covered with dark brown dots and patches. Antenna yellow, segments 4–8 apically and, segments 9–10 entirely brown. Mesopraescutum with two dark brown patches at the fore margin; mesoscutum with two white narrow longitudinal bands lined with dark on either side, and one moderately wide median longitudinal band consisting of black brown and white lines.

Forewing (Fig. 18F) yellow with dense, mostly confluent dots; apices of pterostigma and veins Rs, M_1+2_, M_3+4_, Cu_1a_, Cu_1b_, partly indistinctly, dark brown. Femora with dark dots. Abdomen and terminalia brown.

Structure. Forewing (Fig. 18F) oblong-oval, widest in the middle, narrowly and evenly rounded apically; wing apex situated in cell r_2_ near apex of M_1+2_; C+Sc weakly curved; pterostigma in the middle about as wide as cell r_1_, strongly widening to apical third; Rs weakly, irregularly curved in basal two thirds, weakly curved to fore margin apically; M longer than M_1+2_ and M_3+4_; Cu_1a_ irregularly curved, ending at M fork; surface spinules present in all cells, leaving narrow spinule-free stripes along the veins in apical part, forming distinct hexagons of a double row of spinules, spinules sparse or absent from base of cell c+sc.

Hindwing with 3–4 + 2–3 indistinctly grouped costal setae. Metatibia bearing 6–7 grouped apical spurs, arranged as 3 + 3–4, anteriorly separated by 4 bristles.

Terminalia (Fig. 31A–F). Male. Proctiger weakly produced posteriorly; densely covered with moderately long setae in apical two thirds. Subgenital plate subglobular; dorsal margin straight; posterior margin irregularly convex; with long setae mostly posteriorly. Paramere, in lateral view, acuminate, falcate; apex, in lateral view, narrowly rounded, directed anteriad, in dorsal view, apex narrow, directed inwards and anteriad, with a small tooth; outer face with sparse, moderately long setae in basal half and few short setae in apical half; inner face with dense, moderately long setae; posterior margin with long setae. Proximal segment of aedeagus with apical part elongate and moderately subdivided. Distal segment of aedeagus almost straight in basal half, with dorsal margin hardly sinuate; ventral process situated in the middle of segment, in lateral view, narrow, simple, lamellar, lacking lateral lobes, in dorsal view, ventral process broadly obovoid, distinctly broader than apical dilation, broadest in apical third, unevenly rounded apically; apical dilation, in both lateral and dorsal views, broad, distinctly widening towards evenly rounded apex, dorsal part of apical dilation membranous and hardly expanded at base; sclerotised end tube short and almost straight. – Female terminalia cuneate; with dense setae. Dorsal margin of proctiger, in lateral view, weakly convex distal to circumanal ring, weakly concave in apical third, apex relatively straight, pointed; in dorsal view, apex subacute; circumanal ring, in dorsal view, distinctly cruciform. Subgenital plate, in lateral view, abruptly narrowing in apical third, pointed apically; in ventral view, apex blunt.

***Fifth instar immature.*** Coloration. Light; cephalothoracic sclerite, antenna, wing pads, legs and caudal plate pale brown.

Structure. Eye with one short, simple ocular seta dorsally. Antennal segments with following numbers of pointed sectasetae: 1(0), 2(1), 3(0), 4(2), 5(0), 6(2), 7(1), 8(1), 9(0), 10(0). Forewing pad with five marginal pointed sectasetae; hindwing pad with two marginal pointed sectasetae; both pads lacking sectasetae dorsally. Tarsal arolium broadly triangular apically, 1.1–1.4 times as long as claws. Abdomen with one lateral pointed sectasetae on each side anterior to caudal plate. Caudal plate with anterior margin relatively distant from anterior margin of extra pore fields; with four (1+4) pointed sectasetae on either side laterally, and three pointed sectasetae subapically, on either side of circumanal ring dorsally. Extra pore fields forming continuous outer and inner bands, consisting of small oval patches; outer band relatively short medially. Circumanal ring relatively large.

**Host plant.** *Guatteria* sp. (Annonaceae).

**Distribution.** Brazil (AM).

**Derivation of name.** From Latin *acuminatus* = pointed, referring to the pointed paramere.

**Comments.** *Melanastera acuminata* sp. nov. is separated from the other members of the *ingariko*-group by the the acuminate, falcate paramere. In the shape of the paramere, *M. acuminata* is similar to *M. mazzardoae* sp. nov. and *M. variegata* sp. nov., but it differs from these species in the forewing pattern with dense brownish dots (versus extensively yellow or brown, with colourless patches but without dots).

### 42 Melanastera melanocephala sp. nov

(Figs 1F, 7F, 12C, 18G, 31G–L, 40E, F)

**Type material. Holotype m$: Brazil: RORAIMA:** Uiramutã, Caramambatai, N5.1333/1466, W60.5850/5916, 970–1200 m, 7–14.iv.2015, *Miconia albicans* (D. Burckhardt & D.L. Queiroz) #161(6) (UFPR; dry).

**Paratypes. Brazil: GOIÁS:** 1 m$, 2 f£, Rio Verde, ca. 40 km E Jatai, BR-060, S17.8316, W51.3716, 820 m, 31.x.2012 (D. Burckhardt & D.L. Queiroz) #52 (NHMB; slide; NMB- PSYLL0008563); 1 f£, Mineiros, ca. 15 km NW of Mineiros, BR-364, S17.8310, W51.3709, 860 m, 1.xi.2012 (D. Burckhardt & D.L. Queiroz) #53 (NHMB; 70% ethanol; NMB- PSYLL0007429); 3 m$, 6 f£, 11 immatures, 1 skin, Pirenópolis, Cachoeira do Abade, S15.80877, W48.91899, 960 m, 1.viii.2018, *Miconia albicans* (D.L. Queiroz) #864(4) (NHMB; dry, 70% ethanol; NMB-PSYLL0007454, NMB-PSYLL0008266, NMB-PSYLL0008267); 1 m$, 1 f£, 14 immatures, 1 skin, same but Cachoeira do Lázaro e Santa Maria, S15.84558, W48.95808, 920 m, 1.viii.2018, *Miconia* sp. (D.L. Queiroz) #866(4) (MMBC, NHMB; slide, 70% ethanol; NMB-PSYLL0007455, NMB-PSYLL0007466, NMB-PSYLL0007467).—**Mato Grosso:** 2 m$, Alto Garças, ca. 120 km SE of Rondonópolis, BR- 364, S16.9000, W53.6550, 720 m, 1.xi.2012 (D. Burckhardt & D.L. Queiroz) #54 (NHMB; 70% ethanol; NMB-PSYLL0007430); 1 f£, Chapada dos Guimarães, MT-351, Rio Coxipó- açú, S14.53463, W55.80082, 260 m, 24.viii.2018 (D.L. Queiroz) #880 (NHMB; 70% ethanol; NMB-PSYLL0007456); 1 m$, Cuiabá, Chapada dos Guimarães, Caverna Aroe Jari, S15.5683, W55.4817, 750 m, 2.xi.2012 (D. Burckhardt & D.L. Queiroz) #56 (NHMB; 70% ethanol; NMB-PSYLL0007431); 4 f£, same but Parque Mãe Bonifácia, S15.5766, W56.1016, 200 m, 3.xi.2012 (D. Burckhardt & D.L. Queiroz) #57 (NHMB; 70% ethanol; NMB- PSYLL0007432); 1 f£, Lucas do Rio Verde, S13.2490, W56.0275, 400 m, 12.vi.2016 (L.A. Pezzini) #794 (NHMB; 70% ethanol; NMB-PSYLL0005769); 1 f£, 1 immature, 1 skin, same but BR-262, S20.4755, W53.6191, 490 m, 22.x.2021, *Miconia albicans*, Cerrado vegetation (D. Burckhardt & D.L. Queiroz) #482(3) (NHMB; 70% ethanol; NMB-PSYLL0008569); 2 m$, 15 immatures, 9 skins, Tabaporã, Fazenda Crestani, S11.3133/3666, W55.9616/9759, 330–380 m, 6–8.xi.2012, *Miconia rubiginosa* (D. Burckhardt & D.L. Queiroz) #62(2) (NHMB; slide, 70% ethanol; NMB-PSYLL0007433, NMB-PSYLL0007464, NMB- PSYLL0007465).—**Mato Grosso do Sul:** 2 m$, 2 f£, Campo Grande, near BR-163, S20.8907/8948, W54.5713/6599, 440–450 m, 16.xi.2012, *Miconia albicans* (D. Burckhardt & D.L. Queiroz) #74(3) (NHMB; 70% ethanol; NMB-PSYLL0007435); 3 m$, 1 f£, Rio Verde, MS-427, South of Rio Verde, S19.0183, W54.8583, 470 m, 14.xi.2012, *Miconia albicans* (D. Burckhardt & D.L. Queiroz) #69(2) (NHMB; 70% ethanol; NMB-PSYLL0007411); 1 m$, Rio Verde do Mato Grosso, BR-163, S18.9281/9519, W54.8357/9339, 350–440, 13.xi.2012 (D. Burckhardt & D.L. Queiroz) #68 (NHMB; 70% ethanol; NMB-PSYLL0007434); 1 f£, Ribas do Rio Pardo, Ribas do Rio Pardo, S20.4559, W53.7426, 390 m, 22.x.2021, *Miconia albicans*, edge of cerrado (D. Burckhardt & D.L. Queiroz) #481(7) (NHMB; 70% ethanol; NMB-PSYLL0008568).—**Minas Gerais:** 1 f£, 4 immatures, 1 skin, Buenópolis, Parque Estadual da Serra do Cabral, Marcos Teixeira, S17.9035, W44.2308, 1000 m, 20.ix.2019, *Miconia albicans* (D. Burckhardt & D.L. Queiroz) #369(5) (NHMB; 70% ethanol; NMB- PSYLL0007450); 1 m$, 1 f£, 2 immatures, same but S17.9122, W44.2092, 830 m, 9.iv.2021, *Miconia* sp., dense cerrado vegetation #391(1) (NHMB; 70% ethanol; NMB- PSYLL0008564); 3 m$, 4 f£, 1 immature, Chapada Gaúcha, Parque Nacional Grande Sertão Veredas, road to Base Itaguari, S15.1339, W45.8312, 710 m, 3.v.2021, *Miconia albicans*, Cerrado vegetation (D. Burckhardt & D.L. Queiroz) #451(2) (NHMB; 70% ethanol; NMB- PSYLL0008566); 1 m$, 1 f£, same but Rio Preto, S15.1213, W45.7378, 710 m, 4.v.2021, *Miconia albicans*, Cerrado vegetation (D. Burckhardt & D.L. Queiroz) #452(3) (NHMB; 70% ethanol; NMB-PSYLL0008567); 1 m$, Coromandel, Fazenda Laje, S18.5610, W46.9020, 1060 m, 12.ii.2018 (D. Burckhardt & D.L. Queiroz) #263 (NHMB; 70% ethanol; NMB- PSYLL0007448); 3 m$, 4 f£, same but S18.5610, W46.9020, 1060 m, 23.viii.2019, *Miconia albicans* (D. Burckhardt & D.L. Queiroz) #346(4) (NHMB; 70% ethanol; NMB- PSYLL0007449); 4 m$, 10 f£, 3 immatures, 2 skins, same but S18.57477, W46.88815, 1050 m, 5.iii.2014, *Miconia albicans* (D.L. Queiroz) #605(2) (MMBC, NHMB; slide, 70% ethanol; NMB-PSYLL0007451, NMB-PSYLL0007468–NMB-PSYLL0007470); 1 f£, same but S18.56773, W46.90342, 1060 m, 5.iii.2014 (D.L. Queiroz) #606 (NHMB; 70% ethanol; NMB-PSYLL0007452); 1 m$, 3 immatures, Lavras, S21.2672, W44.9353, 900 m, 1–6.vi.2010, *Miconia albicans*, edge of Atlantic forest around coffee plantation mixed with pastures (D. Burckhardt) #1(12) (NHMB; dry, 70% ethanol; NMB-PSYLL0007416, NMB- PSYLL0008585); 1 m$, 1 f£, Ouro Preto, Trevo da Chapada, cachoeira do moinho, S20.45302, W43.55230, 1240 m, 18.xi.2018 (D.L. Queiroz) #914 (NHMB; 70% ethanol; NMB-PSYLL0007457); 2 m$, 6 f£, Patos de Minas, Fazenda Laje, S18.5428/5584, W46.8992/9164, 990–1060 m, 28.x.2012, *Miconia albicans* (D. Burckhardt & D.L. Queiroz) #49(3) (NHMB; slide, 70% ethanol; NMB-PSYLL0007412, NMB-PSYLL0007440 [LSMelmel-10], NMB-PSYLL0007460 [LSMelmel-10], NMB-PSYLL0007462 [LSMelmel-10], NMB-PSYLL0007463 [LSMelmel-10]); 4 f£, 1 immature, 1 skin, São Gonçalo do Rio Preto, Parque Estadual do Rio Preto, near guest house, S18.1263, W43.3570, 850 m, 16.iv.2021, *Miconia* sp., planted trees (D. Burckhardt & D.L. Queiroz) #411(4) (NHMB; 70% ethanol; NMB-PSYLL0008565); 4 m$, 1 f£, 1 immature, São Roque de Minas, Parque Nacional da Serra da Canastra, 15 km before Cachoeira Casca d’Anta, S20.3391, W46.4392, 850–860 m, 4–8.ix.2014, *Miconia albicans* (D. Burckhardt & D.L. Queiroz) #142(1) (NHMB; 70% ethanol; NMB-PSYLL0007437); 1 f£, Uberlândia, Clube Caça e Pesca Itororo, S18.9932, W48.3065, 830 m, 7.x.2017, *Miconia* sp. (D.L. Queiroz) #836(7) (NHMB; 70% ethanol; NMB-PSYLL0007453); 1 immature, 2 skins, same but 9.ii.2018, *Miconia albicans* (D. Burckhardt & D.L. Queiroz) #262(3) (NHMB; 70% ethanol; NMB-PSYLL0007447); 2 m$, 1 f£, same but Panga, S19.1837, W48.3961, 810 m, 8.ii.2018, *Miconia albicans* (D. Burckhardt & D.L. Queiroz) #261(4) (NHMB; 70% ethanol; NMB-PSYLL0007446); 4 m$, 10 f£, Vargem Bonita, Parque Nacional da Serra da Canastra, Cachoeira Casca d’Anta, around park entrance, S24.8541/8573, W48.6982/7121, 850–860 m, 4–8.ix.2014, *Miconia albicans* (D. Burckhardt & D.L. Queiroz) #141(15) (NHMB; dry, 70% ethanol; NMB- PSYLL00003110, NMB-PSYLL00003118, NMB-PSYLL00003120); 2 m$, 6 f£, same but near waterfall, S20.3090, W46.5231, 860 m, 5.ix.2014, *Miconia albicans* (D. Burckhardt & D.L. Queiroz) #143(2) (NHMB; dry, 70% ethanol; NMB-PSYLL0007436, NMB- PSYLL0007458, NMB-PSYLL0007459); 1 m$, 1 f£, same but plateau, S20.2976/2986, W46.5195/5289, 1160–1250 m, 6.ix.2014, *Miconia* sp. (D. Burckhardt & D.L. Queiroz) #144(3) (NHMB; 70% ethanol; NMB-PSYLL0007438); 4 m$, 3 f£, Viçosa, S20.7633, W42.8717, 1.i.2006, *Miconia* sp. (E.G. Fidelis) (NHMB; dry; NMB-PSYLL0003236–NMB-PSYLL0003242).—**Paraná:** 1 m$, Colombo, Embrapa campus, S25.3196/3353, W49.1562/1677, 920 m, 1–5.iv.2013 (D. Burckhardt & D.L. Queiroz) #96 (NHMB; slide; NMB-PSYLL0007461); 1 f£, Jaguariaíva, Parque Estadual do Cerrado, S24.1638/1846, W49.6534/6665, 660–780 m, 26–27.vi.2015, *Miconia albicans* (D. Burckhardt & D.L. Queiroz) #172(6) (NHMB; 70% ethanol; NMB-PSYLL0007444); 1 m$, same but S24.1685/1833, W49.6603/6670, 780–820 m, 15–16.ii.2016, *Miconia albicans* (D.Burckhardt & D.L. Queiroz) #197(6) (NHMB; 70% ethanol; NMB-PSYLL0007445).—**Roraima:** 1 adult, Boa Vista, 41 km E Boa Vista, Campo Experimental Água Boa, N2.6716, W60.8400, 80 m, 2.iv.2015, *Guatteria* sp. (D. Burckhardt & D.L. Queiroz) #154(4) (NHMB; 70% ethanol; NMB-PSYLL0007443); 2 m$, 2 skins, Pacaraima, along RR-174 from Pacaraima to ca. 20 km S of Pacaraima, N5.1333/1466, W60.5850/5916, 490–910 m, 15.iv.2015, *Miconia albicans* (D. Burckhardt & D.L. Queiroz) #162(5) (NHMB; 70% ethanol; NMB-PSYLL0007442); 3 m$, 3 f£, 13 immatures, 2 skins, same as holotype but (NHMB; dry, slide, 70% ethanol; NMB-PSYLL0007439, NMB-PSYLL0007441, NMB- PSYLL0008264, NMB-PSYLL0008265, NMB-PSYLL0007511).

**Description. *Adult.*** Coloration. Sexually dimorphic. Orange. Male with head, pronotum and anterior half of mesopraescutum brown with dark brown or almost black dots. Antenna yellow, apices of segments 4–9 and entire segment 10 brown. Mesoscutum with four broad, brown longitudinal bands. Forewing (Fig. 18G) whitish with sparse, irregular brown dots; base and apex of pterostigma, apices of R+M+Cu, Rs, M_1+2_, M_3+4_, Cu_1a_, Cu_1b_ and middle section of A each with a dark brown dot. Legs yellow, femora with brown patches.

Abdominal tergites partly and sternites entirely brown. Female as male but with yellow vertex, pronotum and anterior half of mesopraescutum. Younger specimens with less expanded dark colour.

Structure. Forewing (Fig. 18G) oval, widest in the middle, broadly, evenly rounded apically; wing apex situated at apex of M_1+2_; C+Sc weakly curved in apical third; pterostigma in the middle distinctly narrower than cell r_1_, weakly widening to the middle; Rs weakly, irregularly convex; M slightly longer than M_1+2_ and M_3+4_; Cu_1a_ irregularly curved, ending at M fork; surface spinules present in all cells, dense, forming hexagons of a double row of spinules, leaving very narrow spinule-free stripes along the veins. Hindwing with 3–4 + 2–5 grouped costal setae. Metatibia bearing 4–5 grouped apical spurs, arranged as 2 + 2–3, anteriorly separated by 5–7 bristles.

Terminalia (Fig. 31G–L). Male. Proctiger tubular; densely covered with moderately long setae in apical two thirds. Subgenital plate, in lateral view, subglobular; dorsal margin weakly curved; posterior margin weakly rounded; with sparse moderately long setae. Paramere, in lateral view, narrowly acuminate; apex blunt, directed upwards; in dorsal view, apex narrow, acute, directed upwards and slightly inwards, bearing two small sclerotised teeth; outer face with relatively short setae in apical two thirds; inner face with dense, moderately long setae; posterior margin with long setae. Proximal segment of aedeagus with apical part moderately subdivided. Distal segment of aedeagus almost straight in basal half with dorsal margin weakly sinuate; ventral process situated in apical third of segment, short, in lateral view, lamellar, with more or less distinct lateral lobes, in dorsal view, ventral process distinctly broader than apical dilation, widening to apical third, indistinctly three-lobed, median lobe rounded apically; apical dilation, in lateral view, moderately broad, subparallel-sided, dorsal part of apical dilation membranous and slightly produced at base, in dorsal view, apical dilation broadest in apical third, rounded apically; sclerotised end tube short and weakly sinuate. – Female terminalia cuneate; densely covered with mostly short setae. Dorsal margin of proctiger, in lateral view, concave in apical third, apex upturned, pointed; in dorsal view, apex subacute; circumanal ring, in dorsal view, vaguely cruciform. Subgenital plate, in lateral view, abruptly narrowing to apex in apical third, apex upturned, pointed; in ventral view, apex blunt.

***Fifth instar immature.*** Coloration. Yellow to reddish; cephalothoracic sclerite, antennae, wing pads, legs and caudal plate pale to dark brown.

Structure. Eye with one moderately long simple ocular seta dorsally. Antennal segments with following numbers of pointed sectasetae: 1(0), 2(1), 3(0), 4(2), 5(0), 6(2), 7(1), 8(1), 9(0), 10(0). Forewing pad with 7–8 marginal and 1–2 dorsal pointed sectasetae; hindwing pad with 2–3 marginal and one dorsal pointed sectasetae. Metatibiotarsus bearing 3–4 pointed sectasetae on outer side; tarsal arolium narrowly fan-shaped apically, slightly longer or about as long as claws. Abdomen with three lateral pointed sectasetae anterior to caudal plate.

Caudal plate with anterior margin relatively close to anterior margin of extra pore fields; with five pointed sectasetae on either side laterally, one pointed sectaseta medially on each inner side of extra pore fields, and four pointed sectasetae on either side subapically, anterior to circumanal ring. Extra pore fields forming continuous outer and inner bands, consisting of small oval and rounded patches; outer band long medially, end pointing outwards. Circumanal ring small.

**Host plants and biology.** *Miconia albicans* (Sw.) Steud., *M. rubiginosa* (Bonpl.) DC. (Melastomataceae). Immatures covered with waxy secretions develop often on inflorescences and infrutescences (Fig. 1F) but also on young leaves and shoots. The feeding damages the leaf epidermis leading to a crumpling of the leaf (Fig. 40F). The eggs, first white, later turning black, are laid on the young shoots (Fig. 40E).

**Distribution.** Brazil (GO, MG, MS, MT PR, RR).

**Derivation of name.** From the Ancient Greek μέλας = black and κεφαλή = head, referring to the cospicuously black head in the male.

**Comments.** *Melanastera melanocephala* sp. nov. differs from other species of the *ingariko*-group by the conspicuously dark head and pronotum of the male, strongly contrasting from the rest of the thorax, and the straight, slender paramere.

### 43 Melanastera brevicauda sp. nov

(Figs 12D, 18H, 31M–R)

**Type material. Holotype m$: Brazil: MINAS GERAIS:** Conselheiro Lafaiete, S20.5797, W43.7199, 1020 m, 20.viii.2013, *Toona* plantation (D.L. Queiroz) #560 (UFPR; dry).

**Paratypes. Brazil: MINAS GERAIS:** 5 m$, 5 f£, 3 immatures, same as holotype but (NHMB; dry, slide; NMB-PSYLL0007741, NMB-PSYLL0007794, NMB-PSYLL0007795 [LSMelbrev-19], NMB-PSYLL0007796 [LSMelbrev-19], NMB-PSYLL0007797, NMB- PSYLL0007798, NMB-PSYLL0007799, NMB-PSYLL0007792); 4 m$, 3f£, Paula Cândido, S20.8081, W42.9839, 780 m, 22.viii.2013, *Miconia trianae*, experimental plot coffee–*Inga* plantation (D.L. Queiroz) #564(1) (NHMB; dry; NMB-PSYLL0007682, NMB- PSYLL0007793).

**Description. *Adult.*** Coloration. Straw-coloured; head and thorax densely covered with dark brown dots. Antennal segments 4–7 apically and segments 8–10 entirely brown.

Mesopraescutum with two brown patches at fore margin; mesoscutum with four broad orange longitudinal stripes, margined with dark brown dots, and one or two median rows of dark dots. Forewing (Fig. 18H) straw-coloured with sparse, irregular brown dots; apices of pterostigma and veins Rs, M_1+2_, M_3+4_, Cu_1a_ and Cu_1b_ dark brown. Femora with dark brown dots; pro- and mesotarsi brown. Abdominal sternites partly brown.

Structure. Forewing (Fig. 18H) oblong-oval, broadest in the middle, unevenly rounded apically; wing apex situated in cell r_2_ near apex of M_1+2_; C+Sc weakly curved in apical quarter; pterostigma slightly narrower in the middle than r_1_ cell, distinctly widening to apical third; Rs weakly irregularly convex; M longer than M_1+2_ and M_3+4_; Cu_1a_ irregularly convex, ending slightly proximal of M fork; surface spinules present in all cells, leaving very narrow spinule-free stripes along the veins, forming conspicuous hexagons of a double row of spinules. Hindwing with 7–8 ungrouped or 4–5 + 3–4 grouped costal setae. Metatibia bearing 7–9 grouped apical spurs, arranged as 3–4 + 4–5, anteriorly separated by 3 bristles.

Terminalia (Fig. 31M–R). Male. Proctiger weakly produced in posterior half; densely covered with moderately long setae in apical two thirds. Subgenital plate irregularly ovoid; dorsal margin slightly sinuate; posterior margin regularly convex; with moderately long setae.

Paramere, in lateral view, lamellar in basal two thirds, acuminate in apical third; apex, in lateral view, relatively narrow, subacute, directed upwards, in dorsal view, apex truncate, directed inwards, lacking sclerotised tooth; both outer and inner faces with dense, moderately long setae; posterior margin with long setae. Proximal segment of aedeagus with apical part moderately subdivided. Distal segment of aedeagus weakly curved in basal half, with slightly sinuate dorsal margin; ventral process situated slightly distal of the middle of the segment, in lateral view, tubular: short, broad, simple, rounded apically, lacking lateral lobes; in dorsal view, ventral process broad, broadest slightly distal of the middle, broadly rounded laterally and truncate apically; apical dilation, in lateral view, narrow, subparallel-sided, with rounded apex; dorsal part of apical dilation bearing membranous sack at base, appearing angular; in dorsal view, apical dilation, obovoid, broadest in apical third, rounded apically; sclerotised end tube short and weakly sinuate. – Female terminalia cuneate; densely covered with setae. Dorsal margin of proctiger, in lateral view, with imperceptible swelling distal to circumanal ring, concave in apical third, apex acute, upturned; in dorsal view, apex subacute; circumanal ring, in dorsal view, distinctly cruciform. Subgenital plate, in lateral view, abruptly narrowing to subacute apex, which is not upturned; in ventral view, apex truncate, weakly concave in the middle.

***Fifth instar immature.*** Coloration. Light; cephalothoracic sclerite, antenna, wing pads, legs and caudal plate brown.

Structure. Eye with one moderately long simple ocular seta dorsally. Antennal segments with following numbers of pointed sectasetae: 1(0), 2(1), 3(0), 4(2), 5(0), 6(2), 7(1), 8(1), 9(0), 10(0). Forewing pad with 10 marginal and 6–7 dorsal pointed sectasetae; hindwing pad with three marginal and three dorsal pointed sectasetae. Tarsal arolium broadly triangular apically, 1.6 times longer than claws. Abdominal tergites anterior to caudal plate with three rows of 4, 2 and 6 pointed sectasetae dorsally, respectively. Caudal plate dorsally with one row of four pointed sectasetae along anterior margin, one row of four pointed sectasetae submedially, two groups of 2 + 3 sectasetae on lateral tubercles, two sectasetae medially between extra pore fields and two groups of five sectasetae on either side subapically, anterior to circumanal ring; with a few long simple setae on either side ventrally. Extra pore fields forming continuous outer and inner bands, consisting of small rounded patches; outer band very short, end pointing outwards. Circumanal ring large.

**Host plant.** Adults were collected on *Miconia trianae* Cogn. (Melastomataceae) which is a possible host. The sample DLQ#560, which includes immatures, lacks information on the host plant.

**Distribution.** Brazil (MG).

**Derivation of name.** From the Latin adjective *brevis* = short, and the noun *cauda* = tail. The noun in ablative case means “with a short tail” and refers to the short female terminalia, in particular the subgenital plate.

**Comments.** *Melanastera brevicauda* sp. nov. and *M. michali* sp. nov. differ from the other species of the *ingariko*-group in the apically trucate paramere (in dorsal view). *Melanastera brevicauda* differs from *M. michali* in the dark dots on the forewing, sparse in the former, dense and often confluent in the latter, in addition to details in the male and female terminalia.

### 44 Melanastera michali sp. nov

(Figs 2A, 4B, 12E, 18I, 31T–Y)

**Type material. Holotype m$: Brazil: RORAIMA:** 1 f£, Uiramutã, Caramambatai, N5.1333/1466, W60.5850/5916, 970–1200 m, 7–14.iv.2015, *Miconia* cf. *cephaloides* (D. Burckhardt & D.L. Queiroz) #161(1) (UFPR; dry).

**Paratypes. Brazil: RORAIMA:** 3 f£, same as holotype but (NHMB; dry, slide; NMB- PSYLL0007935, NMB-PSYLL0007967 [LSMelmich-37], NMB-PSYLL0008039); 1 m$, 1

f£, same but *Miconia* sp. (D. Burckhardt & D.L. Queiroz) #161(15) (NHMB; slide; NMB- PSYLL0007951 [LSMelmich-37], NMB-PSYLL0007952 [LSMelmich-37]).

**Description. *Adult.*** Coloration. Straw-coloured; head and thorax densely covered with dark brown dots. Antenna yellow, scape and pedicel brown, segments 4–8 apically and segments 9–10 entirely dark brown. Mesopraescutum with two brown patches at fore margin; mesoscutum with four broad orange longitudinal stripes margined with dark brown dots, and one or two median rows of dark dots. Forewing (Fig. 18I) light amber-coloured with irregular, very dense dark dots, often confluent, denser in males than in females; base and apex of pterostigma and apices of Rs, M_1+2_, M_3+4_, Cu_1a_ and Cu_1b_ dark brown. Femora with dark brown dots; pro- and mesotarsi brown. Abdominal sclerites and terminalia of male dark brown, of female straw-coloured with dark brown sternites.

Structure. Forewing (Fig. 18I) oval, widest in the middle, broadly rounded apically; wing apex situated in cell r_2_, near apex of M_1+2_; C+Sc weakly curved in distal quarter; pterostigma narrower than cell r_1_, weakly widening to apical third; Rs weakly, irregularly convex; M longer than M_1+2_ and M_3+4_; Cu_1a_ irregularly convex, ending at M fork; surface spinules present in all cells, forming hexagons of a double row of spinules, spinules leaving very narrow spinule-free stripes along the veins, spinules absent from base of cell c+sc. Hindwing with 6 ungrouped or 3–4 + 2–4 grouped costal setae. Metatibia bearing 9–10 grouped apical spurs, arranged as 3–4 + 5–6, anteriorly separated by 3 bristles.

Terminalia (Fig. 31T–Y). Male. Proctiger weakly produced posteriorly; densely covered with long setae in apical two thirds. Subgenital plate irregularly ovoid; dorsal margin curved; posterior margin slightly sinuate; with a few moderately long setae. Paramere, in lateral view, lamellar in basal two thirds, acuminate in apical third; apex, in lateral view, blunt, in dorsal view, apex truncate, lacking sclerotised tooth, directed upwards and slightly inwards; outer face with dense, moderately long setae in apical half and long setae posteriorly; inner face with long setae subbasally and posteriorly, dense moderately long setae medially and short setae apically. Proximal segment of aedeagus with apical part strongly subdivided. Distal segment of aedeagus weakly curved in basal half, with dorsal margin sinuate; ventral process situated slightly distal of the middle of the segment, in lateral view, narrow, simple, lamellar, lacking lateral lobes, in dorsal view, ventral process lanceolate, broadest at apical third, slightly broader than apical dilation; apical dilation, in lateral view, long and broad, slightly expanding towards rounded apex, in dorsal view, oblong, widest subapically, rounded apically; sclerotised end tube short and weakly sinuate. – Female terminalia cuneate; densely covered with setae. Dorsal margin of proctiger, in lateral view, with a conspicuous bump distal to circumanal ring, strongly concave in apical third, apex pointed and upturned; in dorsal view, apex blunt; circumanal ring, in dorsal view, distinctly cruciform. Subgenital plate, in lateral view, pointed apically; in ventral view, apex truncate.

***Fifth instar immature.*** Unknown (see comments).

**Host plant.** *Miconia* cf. *cephaloides* Michelang. (Melastomataceae) (see comments). The immatures are on the new flush and secrete a lot of flocculent wax (D. Burckhardt & D.L. Queiroz, pers. obs.).

**Distribution.** Brazil (RR).

**Derivation of name.** Dedicated to dearest husband of LŠS, Michal Štarha for his continuous help and support during this project.

**Comments.** Adults and immatures were collected on *Miconia* cf. *cephaloides*. The samples DB-DLQ#161(1) with adult *M. michali* sp. nov. from *Miconia* and #161(16) with adult *M. stanopekari* sp. nov. from ?*Virola*, both with immatures, were inadvertently mixed in the laboratory (D. Burckhardt, pers. comm.). As the immatures are mostly young instars, they cannot be attributed at present to any of the two species.

*Melanastera michali* sp. nov. resembles *M. brevicauda* sp. nov.; see comments under the latter species.

## The *duckei*-group

**Description. *Adult*.** Head, in lateral view, inclined at ≥ 45° from longitudinal axis of body. Vertex (Figs 12D–L; 13A, B) trapezoidal, covered with imbricate microsculpture and microscopical setae. Thorax weakly to strongly arched, covered with microscopical setae; metapostnotum with distinct longitudinal keel. Forewing (Figs 18J, 19A–H) uniformly coloured, with small brown dots (absent in *M. lucens*); pterostigma distinctly expanding towards the middle or apical third, longer than 4.0 times as wide; vein R_1_ weakly convex or straight medially. Paramere, in lateral view, irregularly acuminate or lanceolate. Ventral process of the distal aedeagal segment, in lateral view, straight or hardly curved and with short apico-median lobe, more than half as long as apical dilation; in dorsal view, at least slightly wider than apical dilation, apico-median lobe mostly visible.

***Immature.*** Antenna 10-segmented.

**Comments.** The group includes nine species from Brazil. One species from Costa Rica, viz. *M. lucens* (Burckhardt, Hanson & Madrigal), resembles the *duckei*-group in the structure of the distal segment of the aedeagus and may be closely related to it, but it differs from the Brazilian species in the bright orange body and the absence of dark dots on the forewings (the latter character it shares with *olgae*-group). Confirmed or probable hosts of five Brazilian species are Melastomataceae (*Miconia*), of two species Annonaceae (*Guatteria*, *Xylopia*), and one species develops on Myristicaceae (cf. *Virola*). For one species the host is unknown.

### 45 Melanastera duckei sp. nov

(Figs 12F, 18J, 32A–F)

**Type material. Holotype m$: Brazil: AMAZONAS:** Manaus, Adolpho Ducke Forest Reserve, S2.9645, W59.9202, 100 m, 22.vi.1996, fogging of *Eschweilera pseudodecolorans* (J.C.H. Guerrero, P.O.I. de Lima & G.P.L. Mesquita) F3-129(2) (INPA; dry).

**Paratypes. Brazil: AMAZONAS:** 1 m$, 1 f£, same as holotype but 18.vi.1996, fogging of *Eschweilera rodriguesiana* F3-30(2) (INPA; slide; [LSMelduc-44]); 1 m$, same but 18.vi.1996, fogging of *Eschweilera rodriguesiana* F3-30(4) (BMNH; slide; [LSMelduc-44]); 1 f£, same but 22.vii.1995, fogging of *Ecclinusa guianensis* F2-29(9) (BMNH; slide; [LSMelduc-44]); same but 26.vi.1996, fogging of *Pouteria glomerata* F3-122A(10) (INPA; dry); 1 m$, 1 f£, 1 adult, Manaus, Bairro Tarumã-Açu, BR-174 km 1, S2.9466, W60.0333, 100 m, 2.v.2014 (D. Burckhardt & D.L. Queiroz) #134 (NHMB; slide; NMB-PSYLL0007865 [LSMelduc-44], NMB-PSYLL0007866 [LSMelduc-44], NMB-PSYLL0007867).

**Description. *Adult.*** Coloration. Straw-coloured; head and thorax with dark brown dots.

Antenna yellow; segments 3–9 apically and segment 10 entirely brown. Mesopraescutum with brown patches at fore margin; mesoscutum with four broad, pale brown longitudinal stripes, margined by brown dots, and one or two rows of median dots. Forewing (Fig. 18J) yellow with sparse irregular brown dots; apices of pterostigma and Rs, M_1+2_, M_3+4_, Cu_1a_ and Cu_1b_ dark brown. Legs yellow, pro- and mesofemora with light brown patches.

Structure. Forewing (Fig. 18J) oviform, widest in the middle, broadly rounded apically; wing apex situated in cell r_2_, in distance of apex of M_1+2_; C+Sc curved in apical quarter; pterostigma distinctly narrower than r_1_ cell in the middle, slightly widening to apical third; Rs weakly irregularly convex; M longer than M_1+2_ and M_3+4_; Cu_1a_ weakly convex, ending proximal to M fork; surface spinules present in all cells covering the entire surface up to the veins, forming hexagons of a double row of spinules, spinules sparse to almost absent from base of cell c+sc. Hindwing with 4 + 2–3 grouped costal setae. Metatibia bearing 7–8 grouped apical spurs, arranged as 3–4 + 3–5, anteriorly separated by 3 bristles.

Terminalia (Fig. 32A–F). Male. Proctiger weakly produced posteriorly; densely covered with long setae in apical two thirds. Subgenital plate, in lateral view, subtriangular; dorsal margin almost straigh; posterior margin weakly convex; with sparse, long setae. Paramere, in lateral view, irregularly acuminate; apex, in lateral view, blunt, directed anteriad, in dorsal view, apex narrow, subacute, directed distinctly anteriad and upwards, with a small sclerotised tooth; outer face with sparse, short setae in apical two thirds; inner face with dense, short setae; with long setae posteriorly. Proximal segment of aedeagus with apical part strongly subdivided. Distal segment of aedeagus thick in basal half, with dorsal margin strongly sinuate; ventral process situated slightly proximal of the middle of segment, in lateral view, process large with lateral lobes, in dorsal view, slightly obovate, widest apically, apex with three small lobes of the same length, the median lobe slightly narrower than the lateral ones; apical dilation long and narrow, in both lateral and dorsal views, subparallel-sided, with rounded apex, with membranous lobe basally at dorsal margin; sclerotised end tube moderately long and weakly sinuate. – Female terminalia cuneate; densely covered with setae. Dorsal margin of proctiger, in lateral view, with shallow bulge distal to circumanal ring, weakly concave in apical third, apex subacute; in dorsal view, apex blunt; circumanal ring, in dorsal view, distinctly cruciform. Subgenital plate, in lateral view, pointed apically; in ventral view, apex blunt.

*Fifth instar immature.* Unknown.

**Host plant.** Unknown. Adults were collected by fogging *Ecclinusa guianensis*, *Pouteria glomerata* (Sapotaceae), *Eschweilera pseudodecolorans* and *E. rodriguesiana* (Lecythidaceae), which are unlikely hosts.

**Distribution.** Brazil (AM).

**Derivation of name.** Named after the eminent entomologist and botanist Adolfo Ducke, who investigated the fauna and flora of Amazonia.

**Comments.** *Melanastera duckei* sp. nov. is characterised within the *duckei*-group by its acuminate paramere with the apical portion which is bent forward.

### 46 Melanastera burchellii sp. nov

(Figs 1D, 12G, 19A, 32G–L)

**Type material. Holotype m$: Brazil: GOIÁS:** Pirenópolis, Parque dos Pirineus, S25.3299, W49.1612, 1300 m, 2.viii.2018, *Miconia burchellii* (D.L. Queiroz) #867(2) (UFPR; dry).

**Paratypes. Brazil: GOIÁS:** 1 m$, 3 f£, 14 immatures, 2 skins, same as holotype but (MMBC, NHMB; slide, 70% ethanol; NMB-PSYLL0007791, NMB-PSYLL0007807 [LSMelbur-101], NMB-PSYLL0007808 [LSMelbur-101], NMB-PSYLL0007810, NMB-PSYLL0007811); 1 f£, 2 immatures, 1 skin, Pirenópolis, Cachoeira do Abade, S15.8088, W48.9190, 960 m, 1.viii.2018, *Miconia burchellii* (D.L. Queiroz) #864(6) (NHMB; slide, 70% ethanol; NMB-PSYLL0007789, NMB-PSYLL0007809); 1 f£, 9 immatures, 1 skin, Pirenópolis, Cachoeira do Lázaro e Santa Maria, S15.8456, W48.9581, 920 m, 2.viii.2018, *Miconia burchellii* (D.L. Queiroz) #866(2) (NHMB; 70% ethanol; NMB-PSYLL0007790).

**Description. *Adult.*** Coloration. Yellow. Head and thorax with dark brown dots. Antennal segments 4–8 apically and segments 9–10 entirely brown. Mesopraescutum with two orange patches at the fore margin; mesoscutum with four orange longitudinal stripes. Forewing (Fig. 19A) yellow with sparse, irregular, pale brown dots, dots absent from antero-basal part of wing; apices of pterostigma and Rs, M_1+2_, M_3+4_, Cu_1a_ and Cu_1b_ brown. Pro- and mesofemora with brown patches. Structure. Forewing (Fig. 19A) oblong-oval, broadest in apical third, evenly rounded apically; wing apex situated in cell r_2_ near apex of M_1+2_; C+Sc distinctly convex in apical third; pterostigma slightly narrower in the middle than cell r_1_, weakly widening to the middle; Rs weakly irregularly convex; M longer than M_1+2_ and M_3+4_; Cu_1a_ irregularly convex, ending slightly proximal to (male) or at (female) M fork; surface spinules present in all cells, leaving narrow spinule-free stripes along the veins, dense, forming hexagons of a single row of spinules, spinules absent from basal half of cell c+sc. Hindwing with 3–4 + 3 grouped costal setae. Metatibia bearing 7 grouped apical spurs, arranged as 3–4 + 3–4, anteriorly separated by 2 bristles.

Terminalia (Fig. 32G–L). Male. Proctiger robust, slightly convex posteriorly; densely covered with moderately long setae in apical two thirds. Subgenital plate subglobular; dorsal margin weakly curved; posterior margin convex; with dense, moderately long setae, mostly posteriorly. Paramere, in lateral view, acuminate; apex, in lateral view, blunt, directed upwards, in dorsal view, apex directed inwards and slightly upwards, lacking distinct sclerotised tooth; both outer and inner faces with dense, moderately long setae in apical two thirds; posterior margin with longer setae. Proximal segment of aedeagus with apical part moderately subdivided. Distal segment of aedeagus in basal half with ventral margin convex and dorsal margin sinuate; ventral process large, situated slightly proximal of the middle of segment, in lateral view, tubular, curved, with distinct lateral lobes and apical process directed ventrad; in dorsal view, ventral process much broader than apical dilation, with broad wing-like lateral lobes on each side and a relatively small button-like median lobe directed slightly ventrad; apical dilation, in lateral view, elongate, subparallel-sided, rounded apically, in dorsal view, apical dilation, narrow, slightly widening towards rounded apex; with small membranous lobe at dorsal margin basally; sclerotised end tube short and nearly straight. – Female terminalia cuneate; densely covered with setae. Dorsal margin of proctiger, in lateral view, weakly concave, apex pointed, slightly upturned; in dorsal view, apex blunt; circumanal ring, in dorsal view, vaguely cruciform. Subgenital plate, in lateral view, abruptly narrowing to apex in apical half, pointed apically; in ventral view, apex subacute.

***Fifth instar immature.*** Coloration. Pale yellow; antenna yellow, gradually becoming darker towards apex; legs yellow with brown distal tarsal segment; cephalothoracic sclerite, wing pads, and caudal plate brownish yellow to pale brown.

Structure. Eye with one ocular pointed sectaseta dorsally. Antennal segments with following numbers of pointed sectasetae: 1(0), 2(1), 3(0), 4(2), 5(0), 6(2), 7(1), 8(2), 9(0),

10(0). Forewing pad with 9–10 marginal and 12–15 pointed sectasetae and lanceolate setae scattered dorsally; hindwing pad with 2 marginal and 3–4 dorsal pointed sectasetae. Tarsal arolium broadly fan-shaped apically, 1.5 times as long as claws. Abdominal tergites anterior to caudal plate with three rows of 4, 2 and 6 pointed sectasetae dorsally, respectively. Caudal plate dorsally with one row of 5–6 pointed sectasetae along anterior margin, two rows each of four pointed sectesetae submedially, and with two groups of four pointed sectasetae on either side subapically, anterior to circumanal ring. Extra pore fields forming continuous outer and inner bands, consisting of small oval and round patches; outer band short medially, end pointing backwards. Circumanal ring small.

**Host plant and biology.** *Miconia burchellii* Triana (Melastomataceae). Immatures develop at the base of infrutescences. Their presence can be often visible only by their waxy secretions (Fig. 1D, arrow).

**Distribution.** Brazil (GO).

**Derivation of name.** Named after its host, *M. burchellii*.

**Comments.** *Melanastera burchellii* sp. nov. differs from the other species of the *duckei*- group in the absence of dark dots in the forewing cells c+sc and r_1_.

### 47 Melanastera barretoi sp. nov

(Figs 12H, 19B, 32M–R)

**Type material. Holotype m$: Brazil: MATO GROSSO:** Sorriso, Fazenda Mazzardo II, S12.4217, W55.8432, 390 m, 15.viii.2013 (T. Mazzardo) (UFPR; dry).

**Paratypes. Brazil: MATO GROSSO:** 3 m$, 2 f£, same as holotype but (NHMB, UFPR; dry, slide; NMB-PSYLL0007788, NMB-PSYLL0007803, NMB-PSYLL0007804, NMB- PSYLL0007805 [LSMelbar-M3]); 4 m$, 2 f£, same but S12.4233, W55.5957, 390 m, 26.ii.2015, *Guatteria schomburgkiana* (D.L. Queiroz) #678(3) (NHMB; dry, slide, 70% ethanol; NMB-PSYLL0007787, NMB-PSYLL0007802, NMB-PSYLL0007842, NMB- PSYLL0008283 [LSMelbar-M3]).

**Description. *Adult.*** Coloration. Yellow, head and thorax with dark brown dots. Antenna with brown scape and pedicel, and apices of segments 4–8 and entire segments 9–10 dark brown. Mesopraescutum with pale brown patches at fore margin; mesoscutum with four broad pale brown and one median narrow dark brown longitudinal stripes. Forewing (Fig. 19B) yellow with scattered, irregular brown dots; base and apex of pterostigma and apices of Rs, M_1+2_, M_3+4_, Cu_1a_ and Cu_1b_ dark brown. Femora with dark brown dots, apical tarsal segments brown. Abdominal sternites dark brown, female terminalia brown.

Structure. Forewing (Fig. 19B) oblong-oval, widest in the middle, broadly rounded apically; wing apex situated in cell r_2_, near apex of M_1+2_; C+Sc weakly curved in distal third; pterostigma slightly narrower than r_1_ cell;widening to apical third; Rs weakly convex; M longer than M_1+2_ and M_3+4_; Cu_1a_ weakly convex; distinctly distal to M fork; surface spinules present in all cells, leaving narrow spinule-free stripes along the veins, forming hexagons of a double row of spinules, in male with a single row in basal part of the wing, spinules absent from basal part of cell c+sc. Hindwing with 3–4 + 2–4 grouped costal setae. Metatibia bearing 4–5 apical spurs, arranged as 2 + 2–3, anteriorly separated by 3–4 bristles.

Terminalia (Fig. 32M–R). Male. Proctiger weakly produced posteriorly; densely covered with moderately long setae in apical two thirds. Subgenital plate subglobular; dorsal margin almost straight; posterior margin convex; with a few moderately long setae. Paramere, in lateral view, slightly irregularly acuminate; apex, in lateral view, blunt, in dorsal view, paramere gradually narrowing to blunt apex which is directed upwards and slightly inwards, lacking distinct sclerotised tooth; both outer and inner faces with dense, moderately long setae; posterior margin with long setae. Proximal segment of aedeagus with apical part moderately subdivided. Distal segment of aedeagus weakly curved in basal half; ventral process situated slightly distal of the middle of the segment, in lateral view, ventral process long, tubular, with apical part slightly curved ventrad and two large lateral lobes, in dorsal view, ventral process much broader than apical dilation, irregularly lanceolate; apical dilation, in lateral view, narrow, with slightly convex dorsal margin and rounded apex; sclerotised end tube moderately long and almost straight. – Female terminalia cuneate; densely covered with setae. Dorsal margin of proctiger, in lateral view, weakly convex distal to circumanal ring, apex straight and subacute; in dorsal view, apex subacute; circumanal ring, in dorsal view, distinctly cruciform. Subgenital plate, in lateral view, pointed apically; in ventral view, apex truncate.

*Fifth instar immature.* Unknown.

**Host plant.** Adults were collected on *Guatteria schomburgkiana* Mart. (Annonaceae) which is a possible host.

**Distribution.** Brazil (MT).

**Derivation of name.** Dedicated to Professor Marliton Rocha Barreto, for his help and support in the field.

**Comments.** Within the *duckei*-group, *Melanastera barretoi* sp. nov. shares the slender, acuminate, straight paramere with *M. burchellii* sp. nov. and *M. sericeae* sp. nov. It differs from the former species in the presence (versus absence) of dark dots in forewing cells c+sc and r_1_. From the latter species, it differs as indicated in the key. *Melanastera barretoi* sp. nov. and *M. sericeae* sp. nov. also differ in the DNA barcoding sequence: the uncorrected p-distance between them is 8.62% for *COI* and 8.57% for *cytb*.

### 48 Melanastera sericeae sp. nov

(Figs 12I, 19C, 32S–X)

**Type material. Holotype m$: Brazil: MINAS GERAIS:** Vargem Bonita, Parque Nacional da Serra da Canastra, Cachoeira Casca d’Anta, around park entrance, S24.8541/8573, W48.6982/7121, 860 m, 4–8.ix.2014, *Xylopia sericea* (D. Burckhardt & D.L. Queiroz) #141(10) (UFPR; dry).

**Paratypes. Brazil: MINAS GERAIS:** 18 m$, 22 f£, same as holotype but (MMBC, NHMB, UFPR; dry, slide, 70% ethanol; NMB-PSYLL0008594–NMB-PSYLL0008597, NMB- PSYLL0008086 [LSMelser-43], NMB-PSYLL0008085 [LSMelser-43], NMB-PSYLL00003113–NMB-PSYLL00003116); 7 m$, 2 f£, same but near waterfall, S20.3090, W46.5231, 860 m, 5.ix.2014 (D. Burckhardt & D.L. Queiroz) #143 (NHMB; slide, 70% ethanol; NMB-PSYLL0008094 [LSMelser-43], NMB-PSYLL0008083 [LSMelser-43], NMB- PSYLL0008084 [LSMelser-43], NMB-PSYLL0008082 [LSMelser-43]).—**Paraná:** 1 f£, Ibaiti, Fazendinha Nova Esperança, BR-153, S23.9416, W50.2400, 635 m, 18.ix.2014 (D. Burckhardt & D.L. Queiroz) #148 (NHMB; slide; NMB-PSYLL0008082 [LSMelser-43]).

**Description. *Adult.*** Coloration. Orange; head and thorax with expanded white patches and dark brown dots; orange colour on head reduced. Antenna yellow, apices of segments 4, 6 and 8 and entire segments 9–10 dark brown. Mesopraescutum with pale brown patches at fore margin; mesoscutum with four broad pale brown longitudinal stripes and one row of brown dots in the middle. Forewing (Fig. 19C) amber-coloured with sparse, irregular, brown dots; base and apex of pterostigma and apices of Rs, M_1+2_, M_3+4_, Cu_1a_ and Cu_1b_ dark brown. Legs yellow, femora with dark patches, pro- and mesotarsi light brown. Male tergites and terminalia yellow, sternites dark brown; female abdomen dark brown and terminalia brownish yellow.

Structure. Forewing (Fig. 19C) obovate, widest in the middle, broadly, slightly unevenly rounded apically; wing apex situated in cell r_2_, near M_1+2_ apex; C+Sc weakly curved in distal half of its section between base and costal break; pterostigma long and narrow, slightly narrower than r_1_ cell in the middle, slightly convex in apical two thirds; Rs weakly convex in apical third; M longer than M_1+2_ and M_3+4_; Cu_1a_ irregularly convex, ending at M fork; surface spinules present in all cells, forming hexagons of a double row of spinules, spinules leaving very narrow spinule-free stripes along the veins, surface spinules absent from basal part of cell c+sc. Hindwing with 6–7 + 1–2 grouped costal setae. Metatibia bearing 4–6 grouped apical spurs, arranged as 2–3 + 2–4, separated by 3 bristles.

Terminalia (Fig. 32S–X). Male. Proctiger moderately produced posteriorly; densely covered with long setae in apical two thirds. Subgenital plate irregularly ovoid; dorsal margin almost straight; posterior margin regularly convex; with few moderately long setae. Paramere, in lateral view, narrowly acuminate; apex blunt, directed upwards, slightly anteriad and inwards, lacking distinct sclerotised tooth; outer face with sparse, moderately long setae in apical two thirds; inner face with dense, moderately long setae; posterior margin with long setae. Proximal segment of aedeagus with apical part moderately subdivided. Distal segment of aedeagus weakly sinuate in basal half; ventral process situated slightly distal to the middle of segment, in lateral view, broadly tubular widening to apex with lateral lobes, in dorsal view, ventral process subcircular, weakly produced apico-medially, much broader than apical dilation; apical dilation, in lateral view, elongate, subparallel-sided, with rounded apex, in dorsal view, apical dilation narrow, weakly widening towards apex; sclerotised end tube short and almost straight. – Female terminalia cuneate; densely covered with setae. Dorsal margin of proctiger, in lateral view, weakly concave, apex straight, blunt; in dorsal view, apex subacute; circumanal ring, in dorsal view, distinctly cruciform. Subgenital plate, in lateral view, pointed apically; in ventral view, apex blunt.

*Fifth instar immature.* Unknown.

**Host plant.** Adults were collected on *Xylopia sericea* A.St.-Hil. (Annonaceae) which is a possible host.

**Distribution.** Brazil (MG, PR).

**Derivation of name.** Named after its probable host *Xylopia sericea*.

**Comments.** Within the *duckei*-group, *Melanastera sericeae* sp. nov. resembles *M. barretoi*

sp. nov. and *M. burchellii* sp. nov. For details see comments under *M. barretoi*.

### 49 Melanastera tristis sp. nov

(Figs 12J, 19D, 33A–G)

**Type material. Holotype m$: Brazil: PARANÁ:** Morretes, PR-410, near Rio São João, S25.3757, W48.8692, 130 m, 26.vii.2017, *Miconia tristis* subsp. *australis*, roadside vegetation, Atlantic forest (L. Serbina & I. Malenovský) #DIC4 (UFPR; dry).

**Paratypes. Brazil: PARANÁ:** 7 f£, Morretes, Embrapa, S25.4494/4515, W48.8779/8792, 40 m, 13.ix.2011, *Miconia tristis* (D. Burckhardt & D.L. Queiroz) #4(2) (NHMB; slide, 70% ethanol; NMB-PSYLL0008100, NMB-PSYLL0008130–NMB-PSYLL0008132); 6 m$, 10 f£, 15 immatures, same as holotype but (MMBC, NHMB, UFPR; dry, slide, 70% ethanol; NMB- PSYLL0008604, NMB-PSYLL0008152, NMB-PSYLL0008129 [LSMelmel-27]).

**Material not included in type series. Brazil: MINAS GERAIS:** 1 m$, 2 f£, Catas Altas da Noruega, Beira da estrada, S20.58059, W43.72178, 720 m, 20.viii.2013, degraded Atlantic forest (D.L. Queiroz) #559 (NHMB; slide; NMB-PSYLL0008141, NMB-PSYLL0008142 [LSMeltris-58], NMB-PSYLL0008143 [LSMeltris-58]).—**RIO DE JANEIRO:** 1 m$, 2 f£, 1 immature, Itatiaia, Parque Nacional do Itatiaia, Travessia Ruy Braga, S224322, W446251, 1220 m, 18.iv.2019, *Miconia* cf. *paniculata* (D. Burckhardt & D.L. Queiroz) #336(3) (NHMB; slide, 70% ethanol; NMB-PSYLL0008103, NMB-PSYLL0008134 [LSMeltris-104]); 2 m$, 7 f£, 8 immatures, 1 skin, same but Travessia Ruy Braga, S224322, W446251, 1220 m, 18.iv.2019, *Miconia* cf. *paniculata* (D. Burckhardt & D.L. Queiroz) #336(7) (NHMB; dry, slide, 70% ethanol; NMB-PSYLL0008102, NMB-PSYLL0008140 [LSMelmel-102], NMB-PSYLL0008605).—**São Paulo:** 1 m$, 5 f£, 2 immatures, 2 skins, São José do Barreiro, Parque Nacional da Serra da Bocaina, Cachoeira das Posses, S22.7745, W44.6019, 1220 m, 7.iv.2019, *Miconia* sp. (D. Burckhardt & D.L. Queiroz) #319(3) (NHMB; slide, 70% ethanol; NMB-PSYLL0008101, NMB-PSYLL0008133 [LSMeltris-104]).

**Description. *Adult.*** Coloration. Orange; head and thorax with dark brown dots. Antenna with brown scape and pedicel, and dark brown apical portions of segments 3–9 and entire segment 10. Mesopraescutum with two brown patches at the fore margin; mesoscutum with four broad and one narrow median brown longitudinal stripes margined by dark brown dots. Forewing (Fig. 19D) amber-coloured with irregular brown dots, denser females; apices of veins Rs, M_1+2_, M_3+4_, Cu_1a_ and Cu_1b_ dark brown. Femora with dark patches.

Structure. Forewing (Fig. 19D) oblong-oval, widest in the middle, unevenly rounded apically; wing apex situated in cell r_2_ near apex of M_1+2_; C+Sc distinctly curved in distal third; pterostigma narrower in the middle than r_1_ cell, weakly widening to apical third; Rs weakly irregularly convex; M longer than M_1+2_ and M_3+4_; Cu_1a_ irregularly convex, ending at M fork;

surface spinules present in all cells, leaving very narrow spinule-free stripes along the veins, forming hexagons of a single and/or a double row of spinules, spinules sparse or absent from base of cell c+sc. Hindwing with 3–4 + 2–3 grouped or 7 ungrouped costal setae. Metatibia bearing 7–8 grouped apical spurs, arranged as 3–4 + 4, anteriorly separated by 4–5 bristles.

Terminalia (Fig. 33A–G). Male. Proctiger strongly produced posteriorly in basal third; densely covered with long setae in apical two thirds. Subgenital plate big, subglobular; dorsal margin strongly curved; posterior margin irregularly curved; with sparsely spaced long setae. Paramere, in lateral view, irregularly lanceolate, broadest in the middle; apex, in lateral view, relatively blunt, directed upwards, in dorsal view, apex subacute, directed dorsad and slightly inwards, lacking distinct sclerotised tooth; outer face with dense moderately long setae mostly in apical half; inner face with dense moderately long setae; posterior margin with long setae. Proximal segment of aedeagus with apical part well sclerotised, not distinctly subdivided.

Distal segment of aedeagus weakly curved in basal half; ventral process situated slightly distal of the middle of segment, in lateral view, with short stem and lateral lobes, in dorsal view, ventral process approximately subcircular, broader than apical dilation; apical dilation, in lateral view, narrow, curved, unevenly rounded apically, in dorsal view, apical dilation narrowly obovate; sclerotised end tube short and slightly sinuate. – Female terminalia cuneate; densely covered with setae. Dorsal margin of proctiger, in lateral view, sinuate, apex pointed; in dorsal view, apex blunt; circumanal ring, in dorsal view, vaguely cruciform.

Subgenital plate, in lateral view, pointed apically; in ventral view, apex blunt.

***Fifth instar immature.*** Coloration. Pale to bright yellow; antennal scape and pedicel and segments 6–7 brown, segments 3–5 pale yellow, segments 8–10 dark brown; cephalothoracic sclerite, wing pads, legs and caudal plate dark brown.

Structure. Body densely covered with minute clavate and narrowly lanceolate setae. Eye with one short simple ocular seta dorsally. Antennal segments with following numbers of pointed sectasetae: 1(0), 2(2 – one large, one small), 3(0), 4(2), 5(0), 6(2), 7(1), 8(1), 9(0), 10(0). Forewing pad with five marginal pointed sectasetae; hindwing pad with two marginal pointed sectasetae; both pads densely covered with minute clavate and narrowly lanceolate setae over their entire surface. Tarsal arolium fan-shaped apically, slightly longer than claws. Abdomen with three pointed sectasetae laterally on each side anterior to caudal plate. Caudal plate with anterior margin relatively distant from anterior margin of extra pore fields; with five (2 + 3) pointed sectasetae on well-pronounced tubercles on either side laterally, and three pointed sectasetae subapically on each side of circumanal ring dorsally. Extra pore fields forming continuous outer and inner bands, consisting of moderately large and small oval patches; outer band relatively short medially, end pointing outwards. Circumanal ring large.

**Host plants.** *Miconia* cf. *paniculata* (Mart. & Schrank ex DC.) Naudin, *M. tristis* subsp.

*australis* Wurdack (Melastomataceae).

**Distribution.** Brazil (MG, PR, RJ, SP).

**Derivation of name.** Named after one of its hosts, *M. tristis*.

**Comments.** Within the *duckei*-group, *Melanastera tristis* sp. nov. differs from the other species by the posteriorly strongly produced male proctiger.

The samples DLQ#559 from Minas Gerais and DB-DLQ#336(7) from Rio de Janeiro differ from the type series (#DIC4) in the DNA barcoding sequences: the uncorrected p-distances between #DIC4 and DLQ#559 are 7.45% for *COI* and 10.2% for *cytb*; between #DIC4 and DB+DLQ#336(7) 3.07% for *COI*. Although we could not find any morphological differences between the samples, we did not include the material from MG, RJ and SP in the type series as the high genetic distance would suggest that *M. tristis* may represent cryptic species.

### 50 Melanastera discoloris sp. nov

(Figs 4A, 12K, 19E, 33H–M)

**Type material. Holotype m$: Brazil: PARANÁ:** Londrina, Parque Estadual Mata dos Godoy, S23.4428, W51.2423, 630 m, 12‒14.viii.2019, *Miconia discolor* (D. Burckhardt & D.L. Queiroz) #343(3) (UFPR; dry).

**Paratypes. Brazil: PARANÁ:** 9 m$, 15 f£, 3 immatures, 1 skin, same as holotype but (NHMB; dry, slide, 70% ethanol; NMB-PSYLL0007828, NMB-PSYLL0007989–NMB- PSYLL0007992, NMB-PSYLL0007849, NMB-PSYLL0007850 [LSMeldis-116], NMB- PSYLL0007851 [LSMeldis-116], NMB-PSYLL0007852 [LSMeldis-116], NMB-PSYLL0007853 [LSMeldis-116]); 3 m$, 6 f£, same but *Miconia buddlejoides* (D. Burckhardt & D.L. Queiroz) #343(2) (MMBC, NHMB; 70% ethanol; NMB-PSYLL0007827).

**Description. *Adult.*** Coloration. Straw-coloured, head and thorax with dark brown dots.

Antennal segments 4, 6 and 8 apically and segments 9–10 entirely dark brown. Mesopraescutum with two orange patches at the fore margin; mesoscutum with four broad orange longitudinal stripes margined by brown dots, and one median row of dots. Forewing (Fig. 19E) straw-coloured with dark irregularly spaced brown dots, few or absent from cell c+sc; base and apex of pterostigma, and apices of veins Rs, M_1+2_, M_3+4_, Cu_1a_ and Cu_1b_ dark brown. Femora with dark brown patches. Female proctiger brown apically.

Structure. Forewing (Fig. 19E) obovate, widest in apical third, broadly rounded apically; wing apex situated in cell r_2_ near apex of M_1+2_; C+Sc curved in distal third; pterostigma slightly narrower than r_1_ cell in the middle, weakly widening to middle; Rs weakly convex in basal two thirds, strongly curved to fore margin apically; M longer than M_1+2_ and M_3+4_; Cu_1a_ irregularly convex, ending at M fork; surface spinules present in all cells, leaving very narrow spinule-free stripes along the veins, forming hexagons of a double row of spinules, spinules sparse at base of cell c+sc. Hindwing with 4 + 2–3 grouped costal setae. Metatibia bearing 6– 7 grouped apical spurs, arranged as 3 + 3–4, anteriorly separated by 5 bristles.

Terminalia (Fig. 33H–M). Male. Proctiger weakly produced posteriorly; densely covered with moderately long setae in apical two thirds. Subgenital plate subglobular; dorsal margin strongly curved; posterior margin convex; with moderately long setae. Paramere, in lateral view, irregularly lanceolate, posterior margin slightly indented in the middle; apex, in lateral view, blunt, directed upwards; in dorsal view, paramere apex subacute, directed inwards, lacking sclerotised tooth; outer and inner faces with dense, moderately long setae; posterior margin with long setae. Proximal segment of aedeagus with apical part moderately subdivided. Distal segment of aedeagus with slightly convex ventral margin and slightly sinuate dorsal margin in basal half; ventral process situated slightly proximad of the middle of segment, in lateral view, ventral process clavate with lateral lobes, in dorsal view, ventral process much broader than apical dilation, with a subtriangular lobe on either side, apex blunt, lying in the same plane as lateral lobes; apical dilation, in lateral view, broad, slightly widening towards broadly rounded apex, broadest near apex, dorsal part of apical dilation forming membranous lobe at base, in dorsal view, apical dilation narrow and long,

subparallel-sided, blunt apically; sclerotised end tube short and weakly curved. – Female terminalia cuneate; densely covered with setae. Dorsal margin of proctiger, in lateral view, weakly sinuate, apex pointed, upturned; in dorsal view, apex blunt; circumanal ring, in dorsal view, distinctly cruciform. Subgenital plate, in lateral view, abruptly narrowing to apex in apical half, apex pointed, strongly upturned; in ventral view, apex blunt or truncate.

***Fifth instar immature.*** Coloration. Brownish yellow; cephalothoracic sclerite, antenna, wing-pads, legs and caudal plate brown.

Structure. Eye with one small simple ocular seta dorsally. Forewing pad with five marginal pointed sectasetae; hindwing pad with two marginal pointed sectasetae; both pads lacking sectasetae dorsally. Tarsal arolium broadly triangular apically, 1.4 times as long as claws Abdomen with one pointed sectaseta on either side laterally anterior to caudal plate. Caudal plate with anterior margin relatively remote from anterior margin of extra pore fields; with five (2 + 3) pointed sectasetae on small tubercles on either side laterally, and two pointed sectasetae subapically, on either side of circumanal ring dorsally. Extra pore fields forming continuous outer and inner bands, consisting of large and small oval and elongate patches; outer band long medially, end pointing outwards. Circumanal ring large.

**Host plants.** *Miconia discolor* DC. (Melastomataceae); adults were also collected on *M. buddlejoides* Triana, which is a possible host.

**Distribution.** Brazil (PR).

**Derivation of name.** Named after one of its hosts, *M. discolor*.

**Comments.** *Melanastera discoloris* sp. nov. resembles *M. stanopekari* sp. nov. in the posteriorly indented paramere. It differs from the latter species in the sparser brown dots on the forewing and in the details of the male and female terminalia.

### 51 Melanastera truncata sp. nov

(Figs 12L, 19F, 33N–S)

**Type material. Holotype m$: Brazil: PARANÁ:** Curitiba, Parque Tanguá, S25.3816, W49.2850, 930 m, 6.ii.2013, *Miconia cinerascens* (D. Burckhardt & D.L. Queiroz) #90(7) (UFPR; slide; [LSMeltru-26]).

**Paratypes. Brazil: PARANÁ:** 1 m$, 2 f£, same as holotype but (NHMB; slide; NMB- PSYLL0008144 [LSMeltru-26], NMB-PSYLL0008145 [LSMeltru-26], NMB- PSYLL0008147 [LSMeltru-26]).

**Description. *Adult.*** Coloration. Yellow; head and thorax with dense, dark brown dots. Antennal segments 1–2 partly brown, segments 4–9 brownish apically, segment 10 entirely brown. Mesopraescutum with brown patches at fore margin; mesoscutum with four longitudinal brown stripes. Forewing (Fig. 19F) yellow with sparse, irregular brown dots; apex and base of pterostigma and apices of Rs, M_1+2_, M_3+4_, Cu_1a_ and Cu_1b_ dark brown.

Femora with brown patches. Female terminalia with brown apex.

Structure. Forewing (Fig. 19F) oval, broadest in the middle, broadly and evenly rounded apically; wing apex situated in cell r_2_ near apex of M_1+2_; C+Sc weakly curved in distal third; pterostigma slightly narrower in the middle than cell r_1_, distinctly widening to the middle; Rs weakly convex; M longer than M_1+2_ and M_3+4_; Cu_1a_ irregularly curved, ending at M fork; surface spinules present in all cells, leaving narrow spinule-free stripes along the veins, forming hexagons of a double row of spinules in apical third of the wing. Hindwing with 4 + 4 grouped or 7 ungrouped costal setae. Metatibia bearing 8–9 grouped apical spurs, arranged as 4 + 4–5, anteriorly separated by 3 bristles.

Terminalia (Fig. 33N–S). Male. Proctiger weakly produced posteriorly; densely covered with moderately long setae in apical two thirds. Subgenital plate irregularly ovoid; dorsal margin strongly curved; posterior margin weakly convex; with moderately long setae.

Paramere, in lateral view, lanceolate; apex, in lateral view, narrow, subacute, directed upwards and slightly inwards, in dorsal view, apex triangular, subacute, directed upwards and slightly inwards, lacking distinct sclerotised tooth; outer face with dense, moderately long setae in apical two thirds; inner face with dense, moderately long setae; posterior margin with long setae. Proximal segment of aedeagus with apical part moderately subdivided. Distal segment of aedeagus almost straight in basal half, with weakly sinuate dorsal margin; ventral process situated slightly distad of the middle of segment, in lateral view, spatulate, truncate apically; in dorsal view, ventral process much broader than apical dilation, subcircular; apical dilation, in lateral view, relatively narrow, subparallel-sided, rounded apically; in dorsal view, apical dilation widening towards apex; sclerotised end tube moderately long and almost straight. – Female terminalia cuneate; densely covered with setae. Dorsal margin of proctiger, in lateral view, sinuate, apex upturned, pointed; in dorsal view, apex blunt; circumanal ring, in dorsal view, distinctly cruciform. Subgenital plate, in lateral view, abruptly narrowed in apical half, pointed apically; in ventral view, apex blunt or truncate.

*Fifth instar immature.* Unknown.

**Host plant.** Adults were collected on *Miconia cinerascens* Miq. (Melastomataceae) which is a possible host.

**Distribution.** Brazil (PR).

**Derivation of name.** From the Latin adjective *truncatus* = truncate, referring to the shape of the ventral process of the distal segment of aedeagus.

**Comments.** Within the *duckei*-group, *Melanastera truncata* sp. nov. is characterised by the spatulate, apically truncate ventral process of the distal aedeagal segment.

### 52 Melanastera atlantica sp. nov

(Figs 1E, 13A, 19G, 33T–Y)

**Type material. Holotype m$: Brazil: RIO DE JANEIRO:** Itatiaia, Parque Nacional do Itatiaia, Cachoeira Maromba, S22.4297, W44.6199, 1100 m, 17‒18.iv.2019, *Miconia* cf. *petropolitana* (D. Burckhardt & D.L. Queiroz) #335(5) (UFPR; dry).

**Paratypes. Brazil: PARANÁ:** 2 f£, 2 immatures, Antonina, Usina Parigot de Souza, S25.2438, W48.7511, 30 m, 17‒20.vii.2017, *Guatteria australis* (D. Burckhardt & D.L. Queiroz) #248(9) (NHMB; slide, 70% ethanol; NMB-PSYLL0007736, NMB- PSYLL0007780, NMB-PSYLL0007781, NMB-PSYLL0007737 [LSMelatl-35]).—**RIO DE Janeiro:** 8 m$, 18 f£, same as holotype but (MMBC, NHMB, UFPR; dry, slide, 70% ethanol; NMB-PSYLL0007737, NMB-PSYLL0007675–NMB-PSYLL0007677, NMB-PSYLL0007678 [LSMelatl-35]); 3 m$, 3 f£, 1 immature, 2 skins, same but Travessia Ruy Braga, S22.4322, W44.6251, 1200 m, 18.iv.2019, *Miconia cinnamomifolia* (D. Burckhardt & D.L. Queiroz) #336(8) (NHMB; 70% ethanol; NMB-PSYLL0007738, NMB- PSYLL0007785, NMB-PSYLL0008117 [LSMelatl-35]).

**Description. *Adult.*** Coloration. Sexually dimorphic. Male orange. Antennal segments 3–9 apically, segment 10 entirely dark brown. Forewing (Fig. 19G) yellow, sometimes with sparse, irregular, brown dots; base and apex of pterostigma and apices of vens Rs, M_1+2_, M_3+4_, Cu_1a_ and Cu_1b_ dark brown. Pro- and mesofemora with brown transverse subapical band.

Female as male but with dark brown dots on head and thorax; dark dots very dense on head and pronotum. Mesopraescutum with orange to brown patches at fore margin; mesoscutum with four broad and one narrow median orange longitudinal stripes, margined by dark brown dots. Femora with dark brown patches. Abdominal sternites dark brown; terminalia brown.

Structure. Forewing (Fig. 19G) oval, broadest in the middle, broadly rounded apically; wing apex situated at apex of M_1+2_ or nearby in cell r_2_; C+Sc weakly curved in distal third; pterostigma about as wide as r_1_ cell in the middle, widening to apical third; Rs irregularly convex; M longer than M_1+2_ and M_3+4_; Cu_1a_ irregularly convex, ending at M fork; surface spinules present in all cells, leaving narrow spinule-free stripes along the veins, forming hexagons of a double row of spinules, spinules absent or sparse in basal half of cell c+sc. Hindwing with 4 + 3–4 grouped costal setae. Metatibia bearing 7–9 grouped apical spurs, arranged as 3–4 + 4–5, anteriorly separated by 2 bristles.

Terminalia (Fig. 33T–Y). Male. Proctiger weakly produced posteriorly; densely covered with long setae. Subgenital plate subglobular; distal margin strongly curved; posterior margin convex; with dense, long setae mostly posteriorly. Paramere, in lateral view, lanceolate; apex, in lateral view, blunt, directed upwards, in dorsal view, apex blunt, directed upwards and inwards, lacking distinct sclerotised tooth; both outer and inner faces with dense, moderately long setae; posterior margin with longer setae. Proximal segment of aedeagus with apical part moderately subdivided. Distal segment of aedeagus in basal half almost straight, dorsal margin weakly sinuate; ventral process situated slightly distal of the middle of segment, in lateral view, short and broad, tubular, rounded apically; in dorsal view, ventral process relatively narrow, only slightly broader than apical dilation, widening to apical third, with a short median lobe; apical dilation, in lateral view, relatively broad, hardly widening to broadly rounded apex, with an angular membranous lobe at base dorsally; in dorsal view, apical dilation oblong-oval, with rounded apex; sclerotised end tube moderately long and weakly curved. – Female terminalia cuneate; densely covered with setae. Dorsal margin of proctiger, in lateral view, slightly concave in apical third, apex pointed; in dorsal view, apex blunt; circumanal ring, in dorsal view, distinctly cruciform. Subgenital plate, in lateral view, pointed apically; in ventral view, apex blunt.

***Fifth instar immature.*** Coloration. Pale yellow; cephalothoracic sclerite, antenna, wing pads, legs and caudal plate brown.

Structure. Eye with one short simple ocular seta dorsally. Antennal segments with following numbers of pointed sectasetae 1(0), 2(1), 3(0), 4(2), 5(0), 6(2), 7(1), 8(1), 9(0), 10(0). Forewing pad with 4–5 marginal and 0–1 dorsal pointed sectasetae; hindwing pad with 2 marginal pointed sectasetae. Tarsal arolium broadly fan-shaped apically, slightly longer than claws. Abdomen with one lateral pointed sectaseta on either side anterior to caudal plate. Caudal plate with anterior margin close to anterior margin of extra pore fields; with three pointed sectasetae on either side laterally, and three pointed sectasetae subapically, on either side of circumanal ring dorsally. Extra pore fields forming continuous outer and inner bands, consisting of moderate oval patches; outer band long medially, end pointing outwards. Circumanal ring small.

**Host plant.** *Miconia cinnamomifolia* (DC.) Naudin (Melastomataceae); adults were also collected on *Miconia* cf. *petropolitana* Cogn. which is a possible host. According to the field notes, the sample DB-DLQ#248(9) was collected on *Guatteria australis* A.St.-Hil. (Annonaceae) which is most probably erroneous (D. Burckhardt, unpublished data).

Immatures develop in leaf axils and produce flocculent waxy secretions. Nearby the adults lay eggs on the stems (Fig. 1E).

**Distribution.** Brazil (PR, RJ).

**Derivation of name.** Named after the habitat of the species, the Atlantic Forest.

**Comments.** Within the *duckei*-group, *Melanastera atlantica* sp. nov. shares with *M. truncata* sp. nov. the lanceolate paramere. The two species differ in the details of the male and female terminalia.

### 53 Melanastera stanopekari sp. nov

(Figs 13B, 19H, 34A–F)

**Type material. Holotype m$: Brazil: RORAIMA:** Uiramutã, Caramambatai, N5.1333/1466, W60.5850/5916, 970–1200 m, 7–14.iv.2015, cf. *Virola* (D. Burckhardt & D.L. Queiroz) #161(16) (UFPR; slide; [LSMelsta-51]).

**Paratypes. Brazil: RORAIMA:** 1 f£, same as holotype but (NHMB; slide; NMB- PSYLL0008603 [LSMelsta-51]).

**Description. *Adult.*** Coloration. Yellowish to orange; head and thorax with dark brown dots; head lighter than thorax. Antenna pale brown, scape and pedicel brown, apices of segments 4–8 and entire segments 9–10 dark brown. Mesopraescutum with brown patches at fore margin; mesoscutum with four broad pale brown longitudinal stripes, margined by dark brown dots, and one or two median rows of dark brown dots. Forewing (Fig. 19H) yellow with irregular, sparse brown dots; apices of pterostigma and Rs, M_1+2_, M_3+4_, Cu_1a_ and Cu_1b_

dark brown. Femora with brown patches; apical tarsal segments brown. Abdominal sternites dark brown.

Structure. Forewing (Fig. 19H) oval, widest in the middle, broadly rounded apically; wing apex situated in cell r_2_, near apex of M_1+2_; C+Sc weakly curved in distal third; pterostigma slightly narrower than cell r_1_, widening to apical third; Rs relatively straight in basal two thirds, weakly curved to fore margin apically; M longer than M_1+2_ and M_3+4_; Cu_1a_ irregularly convex, ending at M fork; surface spinules present in all cells, forming hexagons mostly of a double row of spinules, spinules leaving narrow spinule-free stripes along the veins, spinules absent from basal part of cell c+sc. Hindwing with (3–4) + (2–3) grouped costal setae.

Metatibia bearing 9–10 grouped apical spurs, arranged as (4–5) + (5–6), anteriorly separated by 3 bristles.

Terminalia (Fig. 34A–F). Male. Proctiger produced posteriorly; densely covered with long setae in apical two thirds. Subgenital plate subglobular; dorsal margin curved; posterior margin weakly convex; with long setae. Paramere, in lateral view, irregularly lanceolate, incised in basal quarter at posterior margin; apex subacute, in dorsal view, apex directed upwards and slightly inwards, lacking distinct sclerotised tooth; outer face with dense, moderately long setae in apical half and long setae posteriorly; inner face with long setae subbasally and posteriorly, dense moderately long setae medially and short setae apically.

Proximal segment of aedeagus with apical part strongly subdivided. Distal segment of aedeagus weakly curved in basal half, with dorsal margin weakly sinuate; ventral process situated slightly distal of the middle of segment, in lateral view, ventral process tubular with lateral lobes, in dorsal view, ventral process much broader than apical dilation, bearing a subtriangular lobe on either side, apex rounded, lying in the same plane as lateral lobes; apical dilation long and narrow, subparallel-sided, with rounded apex; sclerotised end tube short and nearly straight. – Female terminalia cuneate; densely covered with setae. Dorsal margin of proctiger, in lateral view, sinuate, apex pointed, slightly upturned; in dorsal view, apex blunt; circumanal ring, in dorsal view, distinctly cruciform. Subgenital plate, in lateral view, pointed apically; in ventral view, apex blunt.

*Fifth instar immature.* Unknown.

**Host plant.** Unidentified Myristicaceae (cf. *Virola* sp.).

**Distribution.** Brazil (RR).

**Derivation of name.** Dedicated to Professor Stano Pekár, group leader at the Department of Botany and Zoology, Faculty of Science, Masaryk University, Brno, for his support and help during the project.

**Comments.** Adults and immatures were collected on cf. *Virola* sp. The samples DB- DLQ#161(16) with adult *M. stanopekari* sp. nov. from ?*Virola* and #161(1) with adult *M. michali* sp. nov. from *Miconia*, both with immatures, were inadvertedly mixed in the lab (D. Burckhardt, pers. comm.). As the immatures are mostly young instars, they cannot be attributed at present to any of the two species.

*Melanastera stanopekari* sp. nov. resembles *M. discoloris* sp. nov. For details see comments under the latter species.

## The *smithi*-group

**Description. *Adult*.** Head, in lateral view, inclined at ≥ 45° from longitudinal axis of body (Fig. 7G, H). Vertex (Fig. 13C–J) trapezoidal, covered with distinct imbricate microsculpture and microscopical setae. Thorax weakly to strongly arched, covered with microscopical setae; metapostnotum with distinct longitudinal keel. Forewing (Figs 19I, J, 20A–F) uniformly coloured, with small brown dots; pterostigma expanding towards the middle or apical third, longer than 4.0 times as wide; vein R_1_ weakly convex or straight medially; M longer than M_1+2_ and M_3+4_. Paramere, in lateral view, irregularly acuminate or lanceolate. Ventral process of the distal aedeagal segment, in lateral view, curved and with long apico-median lobe; in dorsal view, wider than apical dilation.

***Immature.*** Antenna 10-segmented.

**Comments.** The group comprises eight species from Brazil, including *Melanastera smithi* (Burckhardt, Morais & Picanço). All species develop on Melastomataceae (*Miconia*, *Pleroma*).

### 54 Melanastera sellowianae sp. nov

(Figs 3E, 7G, 13C, 19I, 34G–L)

*Diclidophlebia* sp., Barreto *et al*. (2020): 381.

**Type material. Holotype m$: Brazil: PARANÁ:** 1 m$, Tibagi, Parque Estadual Guartelá, S24.5623/5666, W50.2589, 920–950 m, 23–25.vi.2015, *Miconia sellowiana* (D. Burckhardt & D.L. Queiroz) #171(1) (UFPR; dry).

**Paratypes. Brazil: PARANÁ:** 1 m$, 2 f£, Cerro Azul, BR-476 km 69, S25.0685, W49.0877, 1080 m, 18–19.iv.2013 (D. Burckhardt & D.L. Queiroz) #106(6) (NHMB; 70% ethanol; NMB-PSYLL0007600); 1 m$, Colombo, Campus da Embrapa, S25.0399, W49.0925, 950 m, 12.vii.2013 (D.L. Queiroz) #546 (NHMB; 70% ethanol; NMB-PSYLL0007631); 14 m$, 22 f£, same but Jardim Botânico, S25.4417, W49.2394, 910 m, 24.v.2018 (D.L. Queiroz) (NHMB; 70% ethanol; NMB-PSYLL0007635); 35 m$, 54 f£, 1 immature, 1 skin, same but S25.4416/4433, W49.2366/2383, 930 m, 19.vii.2012, *Miconia sellowiana* (D. Burckhardt & D.L. Queiroz) #44(4) (NHMB; 70% ethanol; NMB- PSYLL0007601); 5 m$, 7 f£, same but S25.4417, W49.2394, 920 m, 12.iii.2013 (D.L. Queiroz) #462 (NHMB; 70% ethanol; NMB-PSYLL0007622); 1 m$, 1 f£, same but S21.7248, W48.3569, 920 m, 28.viii.2014 (D.L. Queiroz) #649 (NHMB; 70% ethanol; NMB-PSYLL0007634); 4 m$0, 40 f£, Curitiba, UFPR, Centro Politécnico, S25.4450, W49.2316, 890 m, 1.vii.2015, *Miconia* sp. (D. Burckhardt & D.L. Queiroz) #173(10) (NHMB; 70% ethanol; NMB-PSYLL0007611); 13 m$, 15 f£, same but S25.4450, W49.2450, 900 m, 15.vi.2016, *Miconia sellowiana* (D. Burckhardt & D.L. Queiroz) #200(11) (NHMB; 70% ethanol; NMB-PSYLL0005914); 9 m$, 16 f£, 1 skin, same but S25.44662, W49.23206, 840 m, 23.vi.2017, *Miconia sellowiana* (D. Burckhardt & D.L. Queiroz) #231(9) (NHMB; 70% ethanol; NMB-PSYLL0007615); 7 m$, 10 f£, same but S25.4466, W49.2323, 840 m, 23.vi.2017, *Miconia sellowiana* (D. Burckhardt & D.L. Queiroz) #244(3) (NHMB; 70% ethanol; NMB-PSYLL0007620); 1 m$, 1 f£, 4 immatures, same but S25.4478, W49.2316, 920 m, 14.vii.2017, *Miconia sellowiana* (M.R. Barreto) #P1 (NHMB; 70% ethanol; NMB- PSYLL0008554; 10 m$, 16 f£, 3 immatures, same but S25.4478, W49.2316, 920 m, 14.vii.2017, *Miconia sellowiana* (I. Malenovský) #P2 (NHMB; 70% ethanol; NMB- PSYLL0008555); 2 m$, 5 f£, 1 skin, same but S25.4478, W49.2316, 920 m, 14.vii.2017, *Miconia sellowiana* (I. Malenovský) #P3 (NHMB; 70% ethanol; NMB-PSYLL0008556); 3 m$, 5 f£, same but S25.4478, W49.2316, 920 m, 14.vii.2017, *Miconia sellowiana* (I. Malenovský) #P4 (NHMB; 70% ethanol; NMB-PSYLL0008557); 16 m$, 40 f£, same but S25.4478, W49.2316, 920 m, 27.vii.2017, *Miconia sellowiana* (I. Malenovský) (MMBC; 70% ethanol); 2 m$, 2 f£, same but Parque Tanguá, S25.3770, W49.2839, 870 m, 25.vi.2017, *Miconia sellowiana* (D. Burckhardt & D.L. Queiroz) #234(5) (NHMB; 70% ethanol; NMB- PSYLL0007616); 21 m$, 54 f£, Jaguariaíva, Parque Estadual do Cerrado, S24.1638/1846, W49.6534/6665, 660–780 m, 26–27.vi.2015, *Miconia sellowiana* (D. Burckhardt & D.L. Queiroz) #172(5) (NHMB; 70% ethanol; NMB-PSYLL0007610); 2 m$, 9 f£, same but S24.1685/1833, W49.6603/6670, 780–820 m, 15–16.ii.2016, *Miconia sellowiana* (D. urckhardt & D.L. Queiroz) #197(4) (NHMB; 70% ethanol; NMB-PSYLL0007613); 6 f£, same but S24.1829, W49.6617, 807 m, 10.vii.2013, Cerrado vegetation (D.L. Queiroz) #530 (NHMB; 70% ethanol; NMB-PSYLL0007628); 1 m$, 1 f£, same but S24.6562, W49.9988, 870 m, 10.vii.2013, Cerrado vegetation (D.L. Queiroz) #533 (NHMB; 70% ethanol; NMB- PSYLL0007627); 12 m$, 17 f£, Mallet, ARIE Serra do Tigre, S25.9465, W50.8069, 1010 m, 13‒14.vi.2017, *Miconia sellowiana* (D. Burckhardt & D.L. Queiroz) #228(5) (NHMB; 70% ethanol; NMB-PSYLL0007614); 2 f£, Palmeira, Recanto dos Papagaios, S25.4600, W49.7667, 960 m, 7.ii.2016 (D. Burckhardt & D.L. Queiroz) #193 (NHMB; 70% ethanol; NMB-PSYLL0007612); 4 m$, 11 f£, Piraquara, Marumbi, Estação Carvalho, S25.4950, W48.9838, 1000 m, 20.xi.2013 (D.L. Queiroz) #599(4) (NHMB; 70% ethanol; NMB-PSYLL0007633); 5 m$, 14 f£, Ponta Grossa, Parque Estadual de Vila Velha, S25.2238, W49.9927, 960 m, 12.vii.2017 (M.R. Barreto) (NHMB; 70% ethanol; NMB- PSYLL0005478); 1 m$, 4 f£, same but S25.22384/24645, W49.99272/50.03994, 750‒870 m, 12‒14.vii.2017, *Miconia sellowiana* (D. Burckhardt & D.L. Queiroz) #246(21) (NHMB; 70% ethanol; NMB-PSYLL0007617); 9 m$, 16 f£, 1 immature, same but S25.22384/24645, W49.99272/50.03994, 750‒870 m, 12‒14.vii.2017, *Miconia sellowiana* (D. Burckhardt & D.L. Queiroz) #246(9) (NHMB; 70% ethanol; NMB-PSYLL0007619); 1 m$, 1 f£, same but S25.2392, W49.9844, 950 m, 12.vii.2013, Cerrado vegetation (D.L. Queiroz) #538 (NHMB; 70% ethanol; NMB-PSYLL0007629); 2 m$, 4 f£, same but S25.3183, W49.1517, 970 m, 12.vii.2013, Cerrado vegetation (D.L. Queiroz) #540 (NHMB; 70% ethanol; NMB- PSYLL0007630); 5 m$, 14 f£, 1 skin, same but S25.2238/2465, W49.99272/50.0399, 750‒870 m, 14.vii.2017, *Miconia sellowiana*, *Araucaria* forest, transitional forest, *Baccharis* scrub (L. Serbina) #DIC4 (MMBC; 70% ethanol); 5 m$, 6 f£, Tibagi, Parque Guartelá, S24.5683, W50.2653, 930 m, 10.vii.2017 (M.R. Barreto) (NHMB; 70% ethanol; NMB-PSYLL0005463); 99 m$, 132 f£, 2 immatures, 2 skins, same but S24.5623/5666, W50.2589, 920–950 m, 23–25.vi.2015, *Miconia sellowiana* (D. Burckhardt & D.L. Queiroz) #171(1) (BMNH, MHNG, MMBC, NHMB, UFPR, USNM; dry, slide, 70% ethanol; NMB- PSYLL0007608, NMB-PSYLL0007643–NMB-PSYLL0007648, NMB-PSYLL0008286, NMB-PSYLL0008287, NMB-PSYLL0007655, NMB-PSYLL0007656, NMB- PSYLL0007664, NMB-PSYLL0007658 [LSMelsel-4], NMB-PSYLL0007659 [LSMelsel-4], NMB-PSYLL0007660 [LSMelsel-4], NMB-PSYLL0007661 [LSMelsel-4], NMB- PSYLL0007599, NMB-PSYLL0007662, NMB-PSYLL0007663, NMB-PSYLL0007609); 11 m$, 44 f£, 1 immature, same but S24.5683, W50.2553, 940 m, 10–12.vii.2017, *Miconia sellowiana* (D. Burckhardt & D.L. Queiroz) #245(2) (NHMB; slide, 70% ethanol; NMB- PSYLL0007657, NMB-PSYLL0007665, NMB-PSYLL0007618); 3 m$, 11 f£, 1 immature, same but S24.5683, W50.2553, 938 m, 10.vii.2017, *Miconia sellowiana*, Cerrado vegetation (L. Serbina) #DIC4 (MMBC; 70% ethanol); 1 m$, 6 f£, same but S24.5683, W50.2553, 938 m, 11.vii.2017, *Miconia sellowiana*, Cerrado vegetation (L. Serbina) #DIC4 (MMBC; 70% ethanol); 2 m$, 4 f£, 1 adult, same but S24.5683, W50.2553, 750‒870 m, 12.vii.2017, *Araucaria* forest, transitional forest, *Baccharis* scrub (L. Serbina) #DIC4 (MMBC; 70% ethanol); 15 m$, 26 f£, same but S24.5638, W50.2559, 930 m, 8.xii.2013, Cerrado vegetation (D.L. Queiroz) #522 (NHMB; 70% ethanol; NMB-PSYLL0007623); 5 m$, 5 f£, 2 immatures, same but S24.5640, W50.2521, 910 m, 9.vii.2013, Cerrado vegetation (D.L. Queiroz) #524 (NHMB; 70% ethanol; NMB-PSYLL0007624); 7 m$, 11 f£, same but S24.1655, W49.6663, 990 m, 9.vii.2013, Cerrado vegetation (D.L. Queiroz) #527 (NHMB; 70% ethanol; NMB- PSYLL0007625); 11 m$, 10 f£, same but S24.1646, W49.6602, 1040 m, 9.vii.2013, Cerrado vegetation (D.L. Queiroz) #528 (NHMB; 70% ethanol; NMB-PSYLL0007626); 1 m$, 1 f£, Tunas do Paraná, Parque das Lauraceas, S24.8774, W48.7485, 860 m, 7.viii.2013, Atlantic forest (D.L. Queiroz) #555 (NHMB; 70% ethanol; NMB-PSYLL0007632).—**São Paulo:** 4 m$, 4 f£, São José do Barreiro, Parque Nacional da Serra da Bocaina, Cachoeira Santo Isidro, S22.7472, W44.6167, 1220 m, 7.iv.2019, *Miconia sellowiana* (D. Burckhardt & D.L. Queiroz) #320(1) (NHMB; 70% ethanol; NMB-PSYLL0007621).

**Description. *Adult.*** Coloration. Yellowish-orange; head and thorax with brown dots. Male generally paler than female. Antenna yellow, apices of segments 4–9 and entire segment 10 brown. Mesopraescutum with yellow to orange patches at fore margin; mesoscutum with four broad yellow to orange longitudinal stripes. Forewing (Fig. 19I) yellow, with sparse, irregular brown dots; apices of pterostigma and Rs, M_1+2_, M_3+4_, Cu_1a_ and Cu_1b_ dark brown. Legs yellow.

Structure. Forewing (Fig. 19I) oblong-oval, with subparallel costal and anal margins, broadest in the middle, evenly rounded apically; wing apex situated at apex of M_1+2_; C+Sc distinctly curved in distal third; pterostigma distinctly narrower in the middle than r_1_ cell, weakly widening to apical third; Rs weakly, irregularly convex; Cu_1a_ irregularly, weakly curved, ending at M fork; surface spinules large, present in all cells, leaving no spinule-free stripes along the veins, forming hexagons of a double row of spinules. Hindwing with 6–7 ungrouped costal setae. Metatibia bearing 6–10 grouped apical spurs, arranged as (3–5) + (3– 5), anteriorly separated by 2 or more bristles.

Terminalia (Fig. 34G–L). Male. Proctiger weakly produced posteriorly; densely covered with short setae in apical two thirds. Subgenital plate small, irregularly ovoid; dorsal margin strongly curved in proximal third; posterior margin relatively straight; with long setae.

Paramere, in lateral view, irregularly lamellar, acuminate in apical fifth; apex, in lateral view, blunt, directed slightly anteriad, in dorsal view, apex blunt, directed inwards, lacking distinct sclerotised tooth; outer face with short setae in apical half; inner face densely covered with moderately long setae; posterior margin with very long setae. Proximal segment of aedeagus with apical part weakly subdivided. Distal segment of aedeagus strongly curved in basal half; ventral process situated in the middle, with short stem, bearing three large lobes, the median lobe hemispherical, directed ventrad, the two lateral lobes wing-like, oriented laterad, perpendicularly to the median lobe, in dorsal view, twice larger than diameter of the apical dilation; apical dilation narrow, evenly rounded apically; sclerotised end tube short and weakly sinuate. – Female terminalia cuneate; relatively densely covered with setae. Dorsal margin of proctiger, in lateral view, weakly sinuate, apex pointed; in dorsal view, apex blunt; circumanal ring, in dorsal view, vaguely cruciform. Subgenital plate, in lateral view, pointed apically; in ventral view, apex blunt.

***Fifth instar immature.*** Coloration. Light; cephalothoracic sclerite, antenna, wing pads, legs and caudal plate brown.

Structure. Eye with one tiny ocular seta dorsally. Antennal segments with following numbers of pointed sectasetae: 1(0), 2(1), 3(0), 4(2), 5(0), 6(2), 7(0), 8(1), 9(0), 10(0). Forewing pad with seven marginal pointed sectasetae and several narrowly truncate apically lanceolate setae dorsally; hindwing pad with two marginal pointed sectasetae and several (3– 7) narrowly truncate apically lanceolate setae dorsally. Tarsal arolium broadly triangular apically, slightly longer than claws, with relatively short petiole. Abdomen anterior to caudal plate with three transverse rows of sparse (4, i.e. 2 on either side of each segment) narrowly truncate apically lanceolate setae on dorsum and 2 + 2 pointed sectasetae on either side laterally; margin of abdomen including caudal plate covered with dense microtrichia and sparse minute clavate setae. Caudal plate distinctly sinuous subapically; with three rows of four (i.e. two on either side) sparsely spaced pointed sectasetae or narrowly truncate lanceolate setae, 3 + 3 pointed sectasetae laterally on each marginal hump, and 3–4 pointed sectasetae on either side of circumanal ring dorsally. Extra pore fields absent. Circumanal ring large.

**Host plant.** *Miconia sellowiana* Naudin (Melastomataceae).

**Distribution.** Reported from Brazil (PR) (Barreto *et al*. 2020; as *Diclidophlebia* sp.), Brazil (PR, SP).

**Derivation of name.** Named after its host, *Miconia sellowiana*.

**Comments.** Within the *smithi*-group, *Melanastera sellowianae* sp. nov. resembles *M. simillima* sp. nov. in the small body size (FL < 1.5 mm), the short antenna (AL < 0.5 mm, AL/HW < 1.0); elongate forewing (FL/FW > 2.55) with only weakly widening pterostigma, and the lack of extra pore fields on the caudal plate in the immature. *Melanastera sellowianae* differs from *M. simillima* as follows: ML/M_1+2_ < 2.00 (versus > 2.00); male subgenital plate, in lateral view, relatively straight (versus slightly concave) posteriorly; anterior margin of paramere, in lateral view, concave (versus straight) subapically; apical dilation of the distal aedeagal segment short (versus long) relative to ventral process; FP/SP > 1.8 (versus < 1.8); fifth instar immatures: AL < 0.33 (versus > 0.33); BL/AL > 3.0 (versus < 3.0); forewing pad with less (versus more) than eight marginal pointed sectasetae; lanceolate setae on dorsum of wing pads and abdomen narrowly truncate (versus pointed) apically.

### 55 Melanastera simillima sp. nov

(Figs 13D, 19J, 34M–R, 37A–C)

**Type material. Holotype m$: Brazil: PARANÁ:** 1 m$, Morretes, Recanto Engenheiro Lacerda PR-410, S25.3333/3337, W48.9009/9015, 780–870 m, 13.ix.2011, *Pleroma raddianum* (D. Burckhardt & D.L. Queiroz) #2(1) (UFPR; dry).

**Paratypes. Brazil: PARANÁ:** 10 m$, 19 f£, 2 adults without abdomen, 9 immatures, Antonina, Usina Parigot de Souza, S25.2557, W48.7787, 720 m, 19.vii.2017, *Pleroma raddianum* (L. Serbina & I. Malenovský) (MMBC; slide, 70% ethanol); 10 m$, 16 f£, same as holotype but (NHMB, UFPR; dry, slide, 70% ethanol; NMB-PSYLL0008598–NMB- PSYLL0008602, NMB-PSYLL0008087, NMB-PSYLL0008107 [LSMelsim-32], NMB- PSYLL0008109 [LSMelsim-32], NMB-PSYLL0008110 [LSMelsim-32], NMB-PSYLL0008108 [LSMelsim-32]); 2 f£, same but Recanto Ferradura PR-410, S25.4528, W48.8794, 40 m, 13.ix.2011, *Pleroma reitzii* (D. Burckhardt & D.L. Queiroz) #3(1) (NHMB; dry, slide; NMB-PSYLL0008088, NMB-PSYLL0008111); 2 f£, same but PR-410, Recanto Rio Cascata, S25.3340, W48.8984, 800 m, 26.vii.2017, Atlantic forest, *Pleroma raddianum* (L. Serbina & I. Malenovský) #DIC1 (MMBC; 70% ethanol); 1 m$, 3 f£, same but PR-410, Recanto Rio Cascata, S25.3340, W48.8984, 790 m, 26.vii.2017, Atlantic forest, roadside vegetation with dominant *Pleroma raddianum* (L. Serbina & I. Malenovský) #DIC2+3 (MMBC; 70% ethanol); 2 m$, 7 f£, Piraquara, Parque Estadual do Marumbi, S25.1566/1583, W48.9750/9933, 890–1170 m, 23–24.iv.2013, *Miconia pusilliflora* (D. Burckhardt & D.L. Queiroz) #109(10) (NHMB; slide, 70% ethanol; NMB-PSYLL0008090, NMB- PSYLL0008112, NMB-PSYLL0008106, NMB-PSYLL0008104 [LSMelsim-32], NMB- PSYLL0008105 [LSMelsim-32]); 4 m$, 5 f£, same but, S25.1566/1583, W48.9750/9933, 890–1170 m, 23–24.iv.2013, *Miconia pusilliflora* (D. Burckhardt & D.L. Queiroz) #109(6) (NHMB; 70% ethanol; NMB-PSYLL0008089); 1 m$, 1 f£, same but Marumbi, Estação Carvalho, S27.4727, W48.3781, 1002 m, 11.ii.2015 (D.L. Queiroz) #670 (NHMB; 70% ethanol; NMB-PSYLL0008091).—**Santa Catarina:** 1 f£, Florianópolis, Costão do Santinho, S11.8339, W55.4996, 170 m, 16.ii.2015 (D.L. Queiroz) #672 (NHMB; 70% ethanol; NMB-PSYLL0008092).

**Description. *Adult.*** Coloration. As in *M. sellowianae* sp. nov. Structure. Forewing (Fig. 19J) oblong-oval, costal and anal margins subparallel, widest in the middle, evenly rounded apically; wing apex situated in cell r_2_, near apex of M_1+2_; C+Sc moderately curved in distal third; pterostigma distinctly narrower in the middle than r_1_ cell, weakly widening to apical third; Rs weakly convex; Cu_1a_ weakly irregularly curved, ending proximal of M fork; surface spinules large, dense, present in all cells, leaving no or very narrow spinule-free stripes along the veins, forming hexagons of a double row of spinules. Hindwing with 5–6 ungrouped costal setae. Metatibia bearing 6–10 grouped apical spurs, arranged as (3–5) + (3–5), separated by 2 or more bristles.

Terminalia (Fig. 34M–R). Male. Proctiger moderately produced posteriorly; densely covered with short setae in apical two thirds. Subgenital plate small, irregularly ovoid; dorsal margin curved; posterior margin concave; with moderately long setae. Paramere, in lateral view, irregularly acuminate; apex, in lateral view, blunt, directed inwards, in dorsal view, apex lacking distinct sclerotised tooth; outer face with short setae in apical half; inner face with dense moderately long setae; posterior margin with very long setae. Proximal segment of aedeagus with apical part weakly subdivided. Distal segment of aedeagus strongly curved in basal half; ventral process situated in the middle, with a long stem bearing three small lobes: the median directed ventrad, the two lateral lobes oriented laterad, perpendicularly to the median lobe, in dorsal view, about the same size as apical dilation; apical dilation slightly widening towards rounded apex; sclerotised end tube of ductus ejaculatorius short and weakly sinuate. – Female terminalia cuneate; densely covered with setae. Dorsal margin of proctiger, in lateral view, weakly sinuate, apex pointed; in dorsal view, apex blunt; circumanal ring, in dorsal view, vaguely cruciform. Subgenital plate, in lateral view, pointed apically; in ventral view, apex blunt.

***Fifth instar immature.*** Coloration. Pale yellow; antenna pale yellow to pale brown, three apical segments dark brown; cephalothoracic sclerite, wing pads, legs and caudal plate pale brown.

Structure. Eye with one short ocular seta dorsally. Antennal segments with following numbers of pointed sectasetae: 1(0), 2(1), 3(0), 4(2), 5(0), 6(2), 7(0), 8(1), 9(0), 10(0). Forewing pad with 9–10 marginal pointed sectasetae and several (6–8) pointed lanceolate setae dorsally; hindwing pad with two marginal and 3–4 lanceolate setae dorsally. Tarsal arolium (Fig. 37C) broadly triangular apically, about as long as claws, with relatively short petiole. Abdomen anterior to caudal plate with three rows of four sparsely spaced pointed sectasetae dorsally and 2 + 2 pointed sectasetae on either side laterally; margin of abdomen including caudal plate covered with dense microtrichia and sparse minute clavate setae.

Caudal plate (Fig. 37A, B) distinctly sinuous subapically; with three rows of 4–6 sparsely spaced pointed sectasetae or lanceolate setae dorsally, 2 + 2 pointed sectasetae on each marginal hump laterally, and four pointed sectasetae near circumanal ring on either side dorsally. Extra pore fields absent. Circumanal ring large.

**Host plant.** *Pleroma raddianum* (Cham.) Cogn. (Melastomataceae); adults were also collected on *Miconia pusilliflora* (DC.) Naudin and *Pleroma reitzii* (Brade) P.J.F.Guim. & Michelang. which are possible hosts.

**Distribution.** Brazil (PR, SC).

**Derivation of name.** From the Latin adjective *simillimus* = most similar, referring to the close resemblance to *M*. *sellowianae*.

**Comments.** Details on the morphological similarities and diagnostic characters for *Melanastera simillima* sp. nov. and *M. sellowianae* sp. nov. are given under the latter species.

### 56 Melanastera trianae sp. nov

(Figs 13E, 20A, 34S–X)

**Type material. Holotype m$: Brazil: MINAS GERAIS:** Paula Cândido, S20.80806, W42.98389, 780 m, 22.viii.2013, *Miconia trianae*, coffee plantation with *Inga* (D.L. Queiroz) #564 (UFPR; dry).

**Paratypes. Brazil: MINAS GERAIS:** 9 m$, 18 f£, 9 immatures, 8 skins, same as holotype but (MMBC, NHMB, UFPR; dry, slide, 70% ethanol; NMB-PSYLL0008201, NMB- PSYLL0008202, NMB-PSYLL0008151, NMB-PSYLL0008096, NMB-PSYLL0008320 [LSMeltria-20], NMB-PSYLL0008125, NMB-PSYLL0008124, NMB-PSYLL0008126, NMB-PSYLL0008123, NMB-PSYLL0008128, NMB-PSYLL0008127); 1 f£, 9 immatures, 8 skins, same but (D.L. Queiroz) #565 (MMBC, NHMB, UFPR; dry; NMB-PSYLL0008097).

**Description. *Adult.*** Coloration. Yellow; head and thorax with brown dots. Antenna light brown, segment 3 yellow, apices of segments 4–8 and entire segments 9–10 brown. Mesopraescutum with two orange patches at the fore margin; mesoscutum with four broad orange longitudinal stripes, often margined by brown dots, and medial row of dots. Forewing (Fig. 20A) yellow with irregular, moderately dense brown dots; base and apex of pterostigma and apices of Rs, M_1+2_, M_3+4_, Cu_1a_ and Cu_1b_ dark brown. Femora with dark brown patches.

Structure. Forewing (Fig. 20A) oblong-oval, broadest in the middle, broadly, evenly rounded apically; wing apex situated at apex of M_1+2_; C+Sc almost straight; pterostigma slightly narrower in the middle than r_1_ cell, weakly widening to apical third; Rs almost straight, weakly curved to fore margin apically; Cu_1a_ irregularly curved, ending at M fork; surface spinules present in all cells, leaving narrow spinule-free stripes along the veins, forming hexagons of a single row of spinules, spinules absent from basal half of c+sc. Hindwing with 6–7 ungrouped or 4 + 3 grouped costal setae. Metatibia bearing 9–11 grouped apical spurs, arranged as (4–5) + (5–6) exteriorly, anteriorly separated by 1–2 bristles.

Terminalia (Fig. 34S–X). Male. Proctiger weakly curved posteriorly; densely covered with moderately long setae in apical two thirds. Subgenital plate subglobular; dorsal margin strongly curved; posterior margin almost straight; with moderately long setae. Paramere, in lateral view, irregularly lanceolate; apex, in lateral view, directed slightly anteriad, blunt apically, in dorsal view, apex truncate, directed upwards, slightly inwards and anteriad, lacking distinct sclerotised tooth; outer and inner faces with dense, moderately long setae; posterior margin also with moderately long setae. Proximal segment of aedeagus with apical part broad, strongly subdivided. Distal segment of aedeagus thick in basal half, with sinuate dorsal margin; ventral process situated in the middle, in lateral view, ovoid, with large apical extension directed ventrad, in dorsal view, much broader than apical dilation, with regularly curved margins, broadest medially; apical dilation narrow, in lateral view, subparallel-sided, hardly widening towards rounded apex, dorsal margin weakly indented subapically, in dorsal view, apical dilation broadest medially, with narrowly rounded apex; sclerotised end tube moderately long and almost straight. – Female terminalia cuneate; densely covered with setae. Dorsal margin of proctiger, in lateral view, weakly sinuate, apex pointed; in dorsal view, apex blunt; circumanal ring, in dorsal view, distinctly cruciform. Subgenital plate, in lateral view, abruptly narrowing in the middle, apex pointed; in ventral view, apex blunt.

***Fifth instar immature.*** Coloration. Yellow; cephalothoracic sclerite, antenna, wing pads, legs and caudal plate brown.

Structure. Eye with one moderately long, simple ocular seta dorsally. Antennal segments with following numbers of pointed sectasetae: 1(0), 2(1), 3(0), 4(2), 5(0), 6(2), 7(1), 8(1), 9(0), 10(0). Forewing pad with 5–6 marginal and 2 dorsal pointed sectasetae; hindwing pad with 2–3 marginal and one dorsal pointed sectasetae. Metatibiotarsus bearing five pointed lanceolate setae on outer side; tarsal arolium broadly fan-shaped apically, slightly longer than claws. Abdomen with 1–2 lateral pointed sectasetae anterior to caudal plate. Caudal plate with anterior margin close to anterior margin of extra pore fields; with four pointed sectasetae on either side laterally, one pointed sectaseta submedially on either inner side of extra pore fields, and three pointed sectasetae on either side subapically, anterior to circumanal ring.

Extra pore fields forming continuous outer and inner bands, consisting of small and large oval patches; outer band long medially, end pointing outwards. Circumanal ring small.

**Host plant.** *Miconia trianae* Cogn. (Melastomataceae).

**Distribution.** Brazil (MG).

**Derivation of name.** Named after its host, *M. trianae*.

**Comments.** *Melanastera trianae* sp. nov. differs from the other species of the *smithi*-group in the almost straight (versus weakly curved in apical third) vein C+Sc.

### 57 Melanastera cinerascentis sp. nov

(Figs 13F, 20B, 35A–F)

**Type material. Holotype m$: Brazil: PARANÁ:** Antonina, Usina Parigot de Souza, S25.2438, W48.7511, 30 m, 17‒20.vii.2017, *Miconia cinerascens* var. *robusta* (D. Burckhardt & D.L. Queiroz) #248(5) (UFPR; dry).

**Paratypes. Brazil: PARANÁ:** 2 m$, 2 f£, 18 immatures, 4 skins, same as holotype but (NHMB; slide, 70% ethanol; NMB-PSYLL0007848, NMB-PSYLL0007857 [LSMelcin-26], NMB-PSYLL0007854 [LSMelcin-26], NMB-PSYLL0007855 [LSMelcin-26], NMB- PSYLL0007856 [LSMelcin-26], NMB-PSYLL0007861 [LSMelcin-26], NMB- PSYLL0007862 [LSMelcin-26], NMB-PSYLL0007863 [LSMelcin-26], NMB- PSYLL0008204).

**Material not included in type series. Brazil: MINAS GERAIS:** 1 immature, 2 skins, Lavras, near Lavras, Quedas do Rio Bonito, Parque Ecológico, BR-354 km9, S21.3294, W44.9690, 1000 m, 19.vi.2010, *Miconia pepericarpa*, riverine forest (D. Burckhardt) #10(1) (NHMB; slide, 70% ethanol; NMB-PSYLL0007830, NMB-PSYLL0007864 [LSMelcin- 110]).

**Description. *Adult.*** Coloration. Orange; head and abdomen with dark brown dots. Antenna yellow, scape and pedicel pale brown, segments 4–8 with brown apices, segments 9– 10 entirely brown. Mesopraescutum with pale brown patches at fore margin; mesoscutum with four broad orange longitudinal stripes margined by brown dots and several median rows of brown dots. Forewing (Fig. 20B) yellow with moderately dense, irregular brown dots; apices of pterostigma and Rs, M_1+2_, M_3+4_, Cu_1a_ and Cu_1b_ dark brown. Femora with dark patches.

Structure. Forewing (Fig. 20B) oval, widest in the middle, broadly, slightly unevenly rounded; wing apex situated in cell r_2_ near apex of M_1+2_; C+Sc weakly curved in distal third; pterostigma distinctly narrower than r_1_ cell in the middle, weakly widening to the middle; Rs weakly convex in basal two thirds, moderately strongly curved to fore margin apically; Cu_1a_ unevenly convex, ending at M fork; surface spinules present in all cells, forming hexagons of a single row of spinules, spinules leaving narrow spinule-free stripes along the veins, absent from basal half of cell c+sc. Hindwing with 7–10 ungrouped costal setae. Metatibia bearing 9–11 grouped apical spurs, arranged as 4 + (5–7), anteriorly separated by 4 bristles.

Terminalia (Fig. 35A–F). Male. Proctiger weakly produced posteriorly; densely covered with long setae in apical two thirds. Subgenital plate irregularly ovoid; dorsal margin curved in apical third; posterior margin weakly convex; with moderately long setae. Paramere, in lateral view, irregularly lanceolate; apex, in lateral view, blunt, directed slightly anteriad, in dorsal view, apex blunt, directed upwards, slightly inwards and slightly anteriad, lacking distinct sclerotised tooth; both outer and inner faces with dense, moderately long setae; posterior margin with slightly longer setae. Proximal segment of aedeagus with apical part strongly subdivided. Distal segment of aedeagus thick basally, with dorsal margin weakly sinuate in basal half; ventral process situated slightly proximal to the middle of segment, in lateral view, ventral process relatively robust and long, expanded laterally and with apical lobe directed ventrad; in dorsal view, ventral process lanceolate, distinctly broader than apical dilation, apex incised dorsally and produced into a broadly tubular lobe directed ventrad; apical dilation tubular, long, in lateral view, slightly widening towards rounded apex, in dorsal view, apical dilation widest medially; sclerotised end tube moderately long and almost straight. – Female terminalia cuneate; densely covered with setae. Dorsal margin of proctiger, in lateral view, weakly, irregularly concave, apex pointed, upturned; in dorsal view, apex blunt; circumanal ring, in dorsal view, distinctly cruciform. Subgenital plate, in lateral view, pointed apically; in ventral view, apex blunt.

***Fifth instar immature.*** Coloration. Brown; antenna gradually becoming darker towards apex; cephalothoracic sclerite, wing pads, legs and caudal plate dark brown.

Structure. Eye with one short simple ocular seta dorsally. Antennal segments with following numbers of pointed sectasetae: 1(0), 2(1-2), 3(0-1), 4(2), 5(0), 6(2), 7(1), 8(1), 9(0), 10(0). Forewing pad with 4–5 marginal and one dorsal pointed sectasetae; hindwing pad with two marginal and 0–1 dorsal pointed sectasetae. Tarsal arolium broadly fan-shaped apically, slightly longer than claws or about as long Abdomen with 2–3 pointed sectasetae laterally anterior to caudal plate. Caudal plate with anterior margin close to anterior margin of extra pore fields; with 4–5 pointed sectasetae on either side laterally, and three pointed sectasetae subapically, on either side of circumanal ring dorsally. Extra pore fields forming continuous outer and inner bands, consisting of small oval patches; outer band long medially, end pointing outwards. Circumanal ring small.

**Host plants.** *Miconia cinerascens* var. *robusta* Wurdack, *M. pepericarpa* Mart. ex DC. (Melastomataceae).

**Distribution.** Brazil (MG, PR).

**Derivation of name.** Named after its host, *M. cinerascens*.

**Comments.** *Melanastera cinerascentis* sp. nov. shares with *M. cannabinae* sp. nov, *M. minutiflorae* sp. nov., *M. prasinae* sp. nov., *M. smithi* (Burckhardt, Morais & Picanço) and *M. trianae* sp. nov. the lanceolate paramere and the thick, short basal portion of the distal aedeagal segment. The species can be identified based on the morphological characters provided in the keys.

As the sample DB#10(1) contains only a single immature and skins which were collected on *M. pepericarpa* rather than *M. cinerascens*, it is not included in the type series. The uncorrected p-distance between DB-DLQ#248(5) (type material) and DB#10(1) is 2.41% for *COI* and 3.97% for *cytb* (poor quality). As the immatures from the two samples are morphologically similar, they are referred here to the same species.

### 58 Melanastera minutiflorae sp. nov

(Figs 13G, 20C, 35G–L)

**Type material. Holotype m$: Brazil: CEARÁ:** Ubajara, ICMBIO headquarters, S3.8389, W40.9393, 860 m, 2‒7.vii.2016, *Miconia minutiflora* (D. Burckhardt & D.L. Queiroz) #212(4) (UFPR; dry).

**Paratypes. Brazil: CEARÁ:** 8 m$, 12 f£, 13 immatures, same as holotype but (NHMB, MHNG, MMBC, UFPR, USNM; dry, slide, 70% ethanol; NMB-PSYLL0005917, NMB- PSYLL0007649, NMB-PSYLL0007650, NMB-PSYLL0007725, NMB-PSYLL0007726 [LSMelmin-28], NMB-PSYLL0007727 [LSMelmin-28], NMB-PSYLL0007724, NMB-

PSYLL0007719); 13 m$, 28 f£, 67 immatures, 21 skins, Ubajara, Parque Nacional de Ubajara, S3.8368/8371, W40.8980/9096, 780‒820 m, 1‒7.vii.2016, *Miconia crenata* (D. Burckhardt & D.L. Queiroz) #211(9) (MMBC, NHMB; dry, 70% ethanol; NMB- PSYLL0005915, NMB-PSYLL0007743, NMB-PSYLL0007729, NMB-PSYLL0007728, NMB-PSYLL0007740, NMB-PSYLL0007651, NMB-PSYLL0007652, NMB-PSYLL0007653); 1 m$, 2 f£, 4 immatures, same but Planalto, S3.8416, W40.9100, 840 m, 5‒ 6.vii.2016, *Miconia pusilliflora* (D. Burckhardt & D.L. Queiroz) #218(5) (NHMB; 70%

ethanol; NMB-PSYLL0005918); 14 m$, 8 immatures, 1 skin, Tianguá, Parque Nacional de Ubajara, Torres, S3.7983, W40.9050, 880 m, 6.vii.2016, *Miconia pusilliflora* (D. Burckhardt & D.L. Queiroz) #220(7) (NHMB; 70% ethanol; NMB-PSYLL0005921).

**Description. *Adult.*** Coloration. Orange; head and thorax with brown dots. Antenna brownish yellow, segments 3–9 with brown apices, segment 10 entirely brown.

Mesopraescutum with orange to pale brown patches at fore margin; mesoscutum with four broad and one narrow medial orange to pale brown longitudinal stripes. Forewing (Fig. 20C) yellow with irregular, moderately dense, brown dots; apices of pterostigma and Rs, M_1+2_, M_3+4_, Cu_1a_ and Cu_1b_ dark brown. Femora with dark patches. Abdomen sclerites partly dark brown.

Structure. Forewing (Fig. 20C) oval, widest in the middle, broadly rounded apically; wing apex situated in cell r_2_, near the apex of M_1+2_; C+Sc slightly convex in distal third; pterostigma slightly narrower than cell r_1_ in the middle, widening to apical third; Rs relatively straight in basal two thirds, obliquely curved to fore margin apically; Cu_1a_ irregularly convex, ending slightly proximal of M fork or at the level of M fork; surface spinules present in all cells, forming hexagons of a single row (male) or a double row of spinules (female), spinules leaving narrow spinule-free stripes along the veins, absent from basal third of cell c+sc. Hindwing with 4 + (2–4) grouped costal setae. Metatibia bearing 8–10 grouped apical spurs, arranged as (4–5) + (4–5), anteriorly separated by 3 or more bristles.

Terminalia (Fig. 35G–L). Male. Proctiger weakly produced posteriorly; densely covered with long setae in apical two thirds. Subgenital plate irregularly ovoid; dorsal margin curved; posterior margin relatively straight; with long setae. Paramere, in lateral view, asymmetrically lanceolate; apex, in lateral view, narrow, directed anteriad, in dorsal view, apex blunt, directed upwards, slightly inwards and slightly anteriad, lacking distinct sclerotised tooth; both outer and inner faces with dense moderately long setae; posterior margin with moderately long setae. Proximal segment of aedeagus with apical part broad, strongly subdivided. Distal segment of aedeagus thick in basal half, with sinuate dorsal margin; ventral process situated in the middle, in lateral view, ovoid, with small tubular apical extension directed ventrad, in dorsal view, ovoid part much broader than apical dilation; apical dilation narrow, in lateral view, subparallel-sided, weakly widening towards rounded apex, dorsal margin almost straight subapically, in dorsal view, apical dilation subparallel-sided, rounded apically; sclerotised end tube moderately long and almost straight. – Female terminalia cuneate; densely covered with setae. Dorsal margin of proctiger, in lateral view, hardly convex distal to circumanal ring, concave in apical third, apex pointed, slightly upturned; in dorsal view, apex blunt; circumanal ring, in dorsal view, vaguely cruciform. Subgenital plate, in lateral view, abruptly narrowing in apical third, apex pointed; in ventral view, apex blunt.

***Fifth instar immature.*** Coloration. Pale yellow; antenna pale yellow to pale brown, segment 10 with dark brown apex; cephalothoracic sclerite, wing pads, legs and caudal plate pale brown.

Structure. Eye with one short simple ocular seta dorsally. Antennal segment with following numbers of pointed sectasetae: 1(0), 2(2 – one large, one small), 3(0), 4(2), 5(0), 6(2), 7(1), 8(1), 9(0), 10(0). Forewing pad with 5–6 marginal pointed sectasetae; hindwing pad with two marginal pointed sectasetae; both pads lacking sectasetae dorsally. Tarsal arolium narrowly fan-shaped apically, slightly longer than claws. Abdomen with four (2+2) pointed sectasetae on either side laterally anterior to caudal plate. Caudal plate with anterior margin relatively close to anterior margin of extra pore fields; with four (2 + 2) pointed sectasetae on either side laterally, and three pointed sectasetae subapically on either side of circumanal ring dorsally.

Extra pore fields forming continuous outer and inner bands, consisting of small oval patches; outer band long medially, end pointing outwards. Circumanal ring small.

**Host plants.** *Miconia crenata* (Vahl.) Michelang., *Miconia minutiflora* (Bonpl.) DC., *M. pusilliflora* (DC.) Naudin (Melastomataceae).

**Distribution.** Brazil (CE).

**Derivation of name.** Named after one of its hosts, *Miconia minutiflora*.

**Comments.** *Melanastera minutiflorae* sp. nov. differs from the other species of the *smithi*- group as indicated in the keys (see also comments under *M. cinerascentis* sp. nov.).

### 59 Melanastera cannabinae sp. nov

(Figs 13H, 20D, 35M–R)

**Type material. Holotype m$: Brazil: AMAZONAS:** Iranduba, Olaria Montemar, S3.1400, W60.3583, 40 m, 15.iv.2014, *Miconia cannabina* (D. Burckhardt & D.L. Queiroz) #126(3) (UFPR; dry).

**Paratypes. Brazil: AMAZONAS:** 14 m$, 21 f£, 17 immatures, same as holotype but (NHMB; dry, slide, 70% ethanol; NMB-PSYLL0007801, NMB-PSYLL0007984, NMB- PSYLL0007985, NMB-PSYLL0007831 [LSMelcan-3], NMB-PSYLL0007832 [LSMelcan- 3], NMB-PSYLL0007833, NMB-PSYLL0007834, NMB-PSYLL0007835, NMB- PSYLL0007836 [LSMelcan-3], NMB-PSYLL0007837).

**Material not included in type series. Trinidad and Tobago: TRINIDAD:** 1 m$, 2 adults, North Coast Road, 6.iii.1988 (D. Hollis) #M40 (BMNH; dry, slide; [LSMeltrin-sp2]); 2 m$, 2 f£, 12 km SW Point Fortin, 9.iii.1989 (D. Hollis) #M46 (BMNH; dry).

**Description. *Adult.*** Coloration. Yellow; Head and thorax with several scattered dark brown dots. Antenna yellow, apices of segents 4, 6–7 and entire segments 8–10 brown. Mesoscutum with four longitudinal orange bands. Forewing (Fig. 20D) yellow with sparse, irregular brown dots slightly denser distal of nodal line; base and apex of pterostigma and apices of Rs, M_1+2_, M_3+4_, Cu_1a_ and Cu_1b_ brown, sometimes indistinct. Femora with light brown patches.

Structure. Forewing (Fig. 20D) oval, broadest in the middle, broadly rounded apically; wing apex situated in cell r_2_ near apex of M_1+2_; C+Sc slightly convex in distal third; pterostigma narrower in the middle than r_1_ cell, widening to apical third; Rs almost straight in basal two thirds, weakly curved to fore margin apically; Cu_1a_ weakly convex, ending at M fork; surface spinules present in all cells, leaving narrow spinule-free stripes along the veins, forming hexagons mostly of a single (male) or a double (female) row of spinules, spinules absent from base of cell c+sc. Hindwing often with distinctly grouped costal setae (as 4–5 + 3–4). Metatibia bearing 9–10 grouped apical spurs, arranged as 4 + (5–6), anteriorly separated by 2 or more bristles.

Terminalia (Fig. 35M–R). Male. Proctiger distinctly produced posteriorly in basal third; densely covered with long setae in apical two thirds. Subgenital plate subglobular; dorsal margin weakly curved; posterior margin weakly convex; with moderately long setae.

Paramere, in lateral view, irregularly lanceolate; apex, in lateral view, directed obliquely anteriad, blunt apically, in dorsal view, apex blunt, directed upwards, slightly inwards and anteriad, lacking distinct sclerotised tooth; outer and inner faces with dense, moderately long setae; posterior margin also with moderately long setae. Proximal segment of aedeagus with apical part broad, strongly subdivided. Distal segment of aedeagus thick in basal half, with sinuate dorsal margin; ventral process situated in the middle, in lateral view, ovoid, with large tubular apical lobe directed ventrad, in dorsal view, the ovoid part much broader than apical dilation, broadest medially; apical dilation narrow, in lateral view, subparallel-sided, hardly widening to rounded apex, dorsal margin weakly sinuate subapically, in dorsal view, apical dilation broadest medially, gradually narrowing in apical half to rounded apex; sclerotised end tube moderately long, almost straight. – Female terminalia cuneate; densely covered with setae. Dorsal margin of proctiger, in lateral view, hardly convex distal to circumanal ring, weakly concave in apical third, apex pointed, straight; in dorsal view, apex blunt; circumanal ring, in dorsal view, distinctly cruciform. Subgenital plate, in lateral view, abruptly narrowing in apical third, apex pointed; in ventral view, apex blunt.

***Fifth instar immature.*** Coloration. Entirely pale to bright yellow.

Structure. Eye with one short ocular seta dorsally. Antennal segment with following numbers of pointed sectasetae: 1(0), 2(1), 3(0), 4(2), 5(0), 6(2), 7(1), 8(1), 9(0), 10(0). Forewing pad with five marginal pointed sectasetae; hindwing pad with two marginal sectasetae. Tarsal arolium broadly triangular apically, slightly longer than claws. Abdomen with one lateral pointed sectasetae on either side anterior to caudal plate. Caudal plate with anterior margin close to anterior margin of extra pore fields; with four pointed sectasetae on either side laterally, and three pointed sectasetae subapically, on either side of circumanal ring dorsally. Extra pore fields forming continuous outer and inner bands, consisting of small oval patches; outer band long medially, end pointing outwards. Circumanal ring small.

**Host plant.** *Miconia cannabina* Markgr. (Melastomataceae). **Distribution.** Brazil (AM), Trinidad and Tobago (Trinidad). **Derivation of name.** Named after its host, *Miconia cannabina*.

**Comments.** *Melanastera cannabinae* sp. nov. differs from the other species of the *smithi*- group as indicated in the keys (see also comments under *M. cinerascentis* sp. nov.).

The uncorrected p-distance between the samples DB-DLQ#126(3) from Brazil and DH#M40 from Trinidad is 1.69% for *COI*. As the specimens from the two samples do not display morphological differences, they are considered conspecific, but the specimens from Trinidad are not included in the type series.

### 60 Melanastera prasinae sp. nov

(Figs 13I, 20E, 35S–X)

**Type material. Holotype m$: Brazil: RORAIMA:** 1 m$, Pacaraima, along RR-174 from Pacaraima to ca. 20 km S of Pacaraima, N4.4033/4733, W61.1483/1633, 490–910 m, 15.iv.2015, *Miconia prasina* (D. Burckhardt & D.L. Queiroz) #162(2) (UFPR; dry).

**Paratypes. Brazil: AMAZONAS:** 1 m$, 1 f£, 1 immature, Iranduba, Embrapa, Campo Experimental, Caldeirão, S3.2516/2566, W60.2216/2233, 30–50 m, 14–17.iv.2014, *Miconia splendens*/*umbrosa* (D. Burckhardt & D.L. Queiroz) #125(9) (NHMB; slide, 70% ethanol; NMB-PSYLL0008042, NMB-PSYLL0008063 [LSMelpra-76], NMB-PSYLL0008062 [LSMelpra-76]).—**MATO GROSSO:** 1 m$, 1 f£, Sorriso, S12.5926, W55.7947, 340 m, 27.ii.2015 (L.A. Pezzini) #483 (NHMB; 70% ethanol; NMB-PSYLL0005701); 1 m$, 1 f£,

Tabaporã, S11.5816, W55.7666, 370 m, 8.xi.2012 (D. Burckhardt & D.L. Queiroz) #64 (NHMB; 70% ethanol; NMB-PSYLL0008041).—**Minas Gerais:** 4 m$, 1 f£, 2 immatures, 2

skins, Vargem Bonita, Parque Nacional da Serra da Canastra, Cachoeira Casca d’Anta, near waterfall, S20.3090, W46.5231, 860 m, 5.ix.2014, *Miconia calvescens* (D. Burckhardt & D.L. Queiroz) #143(8) (NHMB; slide, 70% ethanol; NMB-PSYLL0008044, NMB- PSYLL0008064); 4 m$, 3 f£, 1 immature, same but Cachoeira Casca d’Anta, around park entrance, S24.8541/8573, W48.6982/7121, 850–860 m, 4–8.ix.2014, *Miconia calvescens* (D. Burckhardt & D.L. Queiroz) #141(19) (NHMB; dry, slide, 70% ethanol; NMB-

PSYLL0003117, NMB-PSYLL0008057 [LSMelpra-76A], NMB-PSYLL0008058 [LSMelpra-76A], NMB-PSYLL00003132, NMB-PSYLL00003131).—**Roraima:** 15 m$, 7

f£, 4 immatures, 1 skin, Amajari, Tepequém, along river, N3.7700, W61.7400, 600 m, 4.iv.2015, *Miconia alata* (D. Burckhardt & D.L. Queiroz) #156(2) (NHMB; slide, 70% ethanol; NMB-PSYLL0008045, NMB-PSYLL0008060 [LSMelpra-65], NMB-

PSYLL0008059 [LSMelpra-65], NMB-PSYLL0008061); 8 m$, 7 f£, 3 immatures, 2 skins,

same as holotype but (MMBC, NHMB, UFPR; dry, slide, 70% ethanol; NMB- PSYLL0008064, NMB-PSYLL0008043, NMB-PSYLL0008027, NMB-PSYLL0008065 [LSMelpra-33], NMB-PSYLL0008066 [LSMelpra-33]).

**Description. *Adult.*** Coloration. Sraw-coloured; head and thorax with brown dots. Antennal segments 3–8 apically and segments 9–10 entirely brown. Mesopraescutum with orange to pale brown patches at fore margin; mesoscutum with four broad and one narrow median orange to pale brown longitudinal stripes. Forewing (Fig. 20E) yellow with irregular, sparse, brown dots; apices of pterostigma and of Rs, M_1+2_, M_3+4_, Cu_1a_ and Cu_1b_ dark brown. Femora with dark brown patches. Abdominal sclerites partly dark brown.

Structure. Forewing (Fig. 20E) oval, widest in the middle, broadly, slightly unevenly rounded; wing apex situated in cell r_2_, near apex of M_1+2_; C+Sc moderately curved in distal third; pterostigma slightly narrower than r_1_ cell, widening to apical third; Rs almost straight in basal two thirds, obliquely curved to fore margin apically; Cu_1a_ irregularly convex, ending distal of M fork; surface spinules present in all cells, forming hexagons mostly of a single row of spinules, spinules leaving narrow spinule-free stripes along the veins, surface spinules absent from basal portion of cell c+sc. Hindwing with (4–5) + (2–3) grouped costal setae. Metatibia bearing 9–11 grouped apical spurs, arranged as (4–5) + (5–6), anteriorly separated by 2 or more bristles.

Terminalia (Fig. 35S–X). Male. Proctiger distinctly produced posteriorly in basal third; densely covered with long setae in apical two thirds. Subgenital plate subglobular; dorsal margin curved; posterior margin weakly convex; with moderately long setae. Paramere, in lateral view, irregularly lanceolate; apex, in lateral view, directed obliquely anteriad, blunt apically, in dorsal view, apex slightly truncate, directed upwards, slightly inwards and anteriad, lacking distinct sclerotised tooth; outer and inner faces with dense, long setae; posterior margin with long setae. Proximal segment of aedeagus with apical part broad, strongly subdivided. Distal segment of aedeagus thick in basal half, with weakly sinuate dorsal margin; ventral process situated slightly proximal of the middle, in lateral view, ovoid, with large tubular apical lobe directed ventrad, in dorsal view, the ovoid part much broader than apical dilation, broadest medially; apical dilation narrow, in lateral view, subparallel- sided, hardly widening to rounded apex, dorsal margin weakly indented subapically, in dorsal view, apical dilation narrow, broadest medially, with blunt apex; sclerotised end tube moderately long and almost straight. – Female terminalia cuneate; densely covered with setae. Dorsal margin of proctiger, in lateral view, hardly convex distal to circumanal ring, weakly concave in apical third, apex pointed, slightly upturned; in dorsal view, apex blunt; circumanal ring, in dorsal view, distinctly cruciform. Subgenital plate, in lateral view, abruptly narrowing to apex in apical third, pointed apically; in ventral view, apex blunt.

***Fifth instar immature.*** Coloration. Pale yellow; cephalothoracic sclerite, antenna, wing pads, legs and caudal plate pale brown.

Structure. Eye with one short simple ocular seta dorsally. Antennal segments with following numbers of pointed sectasetae: 1(0), 2(2 – one large, one small), 3(0), 4(2), 5(0),

6(2), 7(1), 8(1), 9(0), 10(0). Forewing pad with 4–5 marginal pointed sectasetae; hindwing pad with two marginal pointed sectasetae; both pads lacking sectasetae dorsally. Tarsal arolium narrowly fan-shaped apically, 1.2–1.5 times longer than claws. Abdomen with 2–3 pointed sectasetae laterally on each side anterior to caudal plate. Caudal plate with anterior margin relatively close to anterior margin of extra pore fields; with four (2 + 2) pointed sectasetae one weakly pronounced tubercles on either side laterally and three pointed sectasetae subapically on each side of circumanal ring dorsally. Extra pore fields forming continuous outer and inner bands, consisting of small oval patches; outer band long medially, end pointing outwards. Circumanal ring small.

**Host plants.** *Miconia alata* (Aubl.) DC., *M. calvescens* DC., *M. prasina* (Sw.) DC.,

*Miconia splendens* (Sw.) Griseb./*M. umbrosa* Cogn. (Melastomataceae).

**Distribution.** Brazil (AM, MG, MT, RR).

**Derivation of name.** Named after *Miconia prasina*, one of its hosts.

**Comments.** *Melanastera prasinae* sp. nov. differs from the other species of the *smithi*- group as indicated in the keys (see also comments under *M. cinerascentis* sp. nov.). The specimens from the samples DB-DLQ#162(2) collected on *M. prasina* and DB- DLQ#156(2) collected on *M. alata*, both from Roraima, differ in the DNA barcoding sequences (uncorrected p-distance 1.95% for *COI* and 1.56% for *cytb*). Similarly, the uncorrected p-distance between DB-DLQ#162(2) and the samples DB-DLQ#143(8) and DB- DLQ#141(19) collected on *M. calvescens* in Minas Gerais is 1.73% for *COI* and 1.56% for *cytb*. The uncorrected p-distance between DB-DLQ#162(2) and the sample DB-DLQ#125(9) collected on *M. splendens*/*M. umbrosa* in Amazonas is 2.85% for *COI* and 1.82% for *cytb*. As no morphological differences were found between the specimens, they are all considered conspecific.

### 61 *Melanastera smithi* (Burckhardt, Morais & Picanço, 2006)

(Figs 7H, 13H, 20F, 36A–F, 38B)

*Diclidophlebia*, Picanço *et al*. (2005): 3.

*Diclidophlebia smithi* Burckhardt, Morais & Picanço, 2006: 242.

*Melanastera smithi*, Burckhardt *et al*. (2023): 412.

**Material examined. Holotype m$: Brazil: MINAS GERAIS:** Viçosa, S20.7633, W42.8717, 650 m, 3.viii.2004, *Miconia calvescens* (E.G. Fidelis M.) (NHMB; dry; NMB- PSYLL0008290).

**Paratypes: Brazil: MINAS GERAIS:** 5 m$, 6 f£, 7 immatures, same as holotype but (NHMB; dry, slide, 70% ethanol; NMB-PSYLL0008291–NMB-PSYLL0008297, NMB- NMB-PSYLL0008288, NMB-PSYLL0008289, NMB-PSYLL0008213, NMB- PSYLL0008214, NMB-PSYLL0008160 NMB-PSYLL0008319 [LSMelsmit], NMB-

PSYLL0008215); 1 immature, same but 1.i.2006, *Miconia calvescens* (E.G. Fidelis M.) (NHMB; 70% ethanol; NMB-PSYLL0008318).

**Amended description. *Adult.*** Coloration. Reddish to yellowish brown; head and thorax with brown dots. Antenna yellow, apices of segments 4, 6, 8–9 and entire segment 10 brown. Mesoscutum with four broad and one narrow, orange or pale brown median longitudinal stripes. Forewing (Fig. 20F) amber-coloured with irregular brown dots, very sparse at base, relatively dense apically; apices of pterostigma and of Rs, M_1+2_, M_3+4_, Cu_1a_ dark brown, sometimes faint or almost wanting. Femora with dark brown patches. Abdominal sclerites with brown margins.

Structure. Forewing (Fig. 20F) oval, widest in the middle, broadly, evenly rounded apically; wing apex situated in cell r_2_ near apex of M_1+2_; C+Sc weakly curved in distal third; pterostigma distinctly narrower than r_1_ cell in the middle, widening to the middle; Rs relatively straight in basal two thirds, obliquely curved to fore margin apically; Cu_1a_ irregularly convex, ending slightly proximal to M fork; surface spinules present in all cells, leaving very narrow spinule-free stripes along the veins, forming hexagons of a double row of spinules, spinules absent from or sparse in basal third of cell c+sc. Hindwing with 6–10 ungrouped or (3–6) + (3–4) grouped costal setae. Metatibia bearing 7–8 grouped apical spurs, arranged as (3–4) + (4–5), anteriorly separated by 2 or more bristles.

Terminalia (Fig. 36A–F). Male. Proctiger produced posteriorly in the middle; densely covered with long setae in apical two thirds. Subgenital plate subglobular; dorsal margin curved; posterior margin slightly convex; with long setae. Paramere, in lateral view, irregularly lanceolate; apex, in lateral view, narrowly blunt, directed upwards and slightly anteriad, in dorsal view, apex subacute, directed upwards, slightly inwards and anteriad, lacking distinct sclerotised tooth; outer and inner faces with dense, moderately long setae; posterior margin with longer setae. Proximal segment of aedeagus with apical part broad, strongly subdivided. Distal segment of aedeagus thick and with weakly sinuate dorsal margin in basal half; ventral process situated slightly proximal to the middle, in lateral view, ovoid, with large tubular apical lobe directed ventrad, in dorsal view, ovoid part much broader than apical dilation, broadest medially; apical dilation narrow, in lateral view, slightly widening to truncate apex, with a small membranous extension at base dorsally; in dorsal view, apical dilation narrow, broadest in apical quarter, with narrowly rounded apex; sclerotised end tube moderately long, weakly curved. – Female terminalia cuneate; densely covered with setae.

Dorsal margin of proctiger, in lateral view, slightly convex distal to circumanal ring, weakly concave in apical third, apex pointed, slightly upturned; in dorsal view, apex blunt; circumanal ring, in dorsal view, distinctly cruciform. Subgenital plate, in lateral view, abruptly narrowing to apex in apical third, pointed apically; in ventral view, apex blunt.

***Fifth instar immature*** (Fig. 38B). Coloration. Yellow; cephalothoracic sclerite, antenna, wing pads, legs and caudal plate brown. Structure. Body densely covered with minute lanceolate setae. Eye with one short simple ocular seta dorsally. Antennal segments with following numbers of pointed sectasetae: 1(0), 2(2 – one large, one small), 3(0), 4(2), 5(0), 6(2), 7(1), 8(1), 9(0), 10(0). Forewing pad with five marginal pointed sectasetae and densely arranged minute lanceolate setae; hindwing pad with two marginal pointed sectasetae. Tarsal arolium narrowly fan-shaped apically, about as long as claws. Abdomen with two pointed sectasetae laterally on each side anterior to caudal plate. Caudal plate with anterior margin close to anterior margin of extra pore fields; with three pointed sectasetae on either side laterally, and 3–4 (1 or 2 + 2) pointed sectasetae near circumanal ring on weak tubercles on either side dorsally. Extra pore fields forming continuous outer and inner bands, consisting of small and large oval and rounded patches; outer band long medially, end pointing outwards. Circumanal ring small.

**Host plant.** *Miconia calvescens* DC. (Melastomataceae).

**Distribution.** Brazil (MG) (Picanço *et al*. 2005, as *Diclidophlebia* sp.; Burckhardt *et al*.

2006b; de Morais *et al*. 2008, 2010, 2013, as *Diclidophlebia smithi*).

**Comments.** *Melanastera smithi* differs from the other species of the *smithi*-group as indicated in the keys (see also comments under *M. cinerascentis* sp. nov.).

In view of the control of *Miconia calvenscens* on several Pacific islands, de Morais *et al*. (2008, 2010, 2013) studied several aspects of the biology and life history of *M. smithi*.

## Ungrouped species

### 62 Melanastera granulosi sp. nov

(Figs 13K, 20G, 36G)

**Type material. Holotype f£: Brazil: RIO DE JANEIRO:** Itatiaia, Parque Nacional do Itatiaia, Cachoeira Maromba, S22.4297, W44.6199, 1100 m, 17‒18.iv.2019, *Pleroma granulosum* (D. Burckhardt & D.L. Queiroz) #335(3) (UFPR; slide; [LSMelgra-106]).

**Paratypes. Brazil: RIO DE JANEIRO:** 5 immatures, 2 skins, same as holotype but (NHMB; slide, 70% ethanol; NMB-PSYLL0007893, NMB-PSYLL0007924 [LSMelgra-106], NMB- PSYLL0007925).

**Description. *Adult.*** Coloration. Body pale yellow. Eyes dark. Antenna pale yellow, segment 8 with dark brown apex, segments 9–10 entirely dark brown. Mesopraescutum with

two orange patches along the fore margin; mesoscutum with four broad orange longitudinal stripes. Forewing (Fig. 20G) pale yellow with whitish stripes along veins, lacking dark dots. Protarsus dark brown.

Structure. Head, in lateral view, inclined 45° from longitudinal body axis. Vertex (Fig. 13K) trapezoidal, with imbricate microsculpture and microscopic setae. Thorax moderately arched, with microscopic setae. Forewing (Fig. 20G) oblong-oval, with subparallel costal and anal margins, broadest slightly distal of the middle, broad rounded apically; wing apex in the middle of margin of cell r_2_; C+Sc weakly curved in apical third; pterostigma slightly narrower in apical third than r_1_ cell, curved in apical third; Rs straight, weakly curved to costal margin in apical quarter; M much longer than M_1+2_ and M_3+4_; Cu_1a_ straight in basal two thirds, curved to anal margin in apical third, ending at M fork; cell cu_1_ narrow; surface spinules present in all cells, absent from basal half of cell c+sc, leaving broad spinule-free stripes along the veins, forming hexagons of a single row of spinules. Metatibia bearing 9 grouped apical spurs, arranged as 3 + 6, anteriorly separated by 1 bristle.

Terminalia. Male. Unknown. – Female terminalia (Fig. 36G) cuneate; densely covered with setae. Dorsal margin of proctiger, in lateral view, straight posterior to circumanal ring, apex blunt, in dorsal view, apex blunt; circumanal ring, in dorsal view, vaguely cruciform. Subgenital plate with long apical process; in lateral view, abruptly narrowing in apical half, pointed apically; in ventral view, apex blunt.

***Fifth instar immature.*** Coloration. Pale yellow.

Structure. Eye with one long simple ocular seta dorsally. Antennal segments with following numbers of pointed sectasetae: 1(0), 2(1), 3(0), 4(2), 5(0), 6(2), 7(1), 8(1), 9(0), 10(0). Each forewing pad with 9–11 marginal sectasetae and ca. 30 large sectasetae or smaller lanceolate setae dorsally. Metatibiotarsus long; tarsal arolium broadly fan-shaped apically, 1.2 times as long as claws. Abdominal tergites anterior to caudal plate with three rows of 5–6, 7– 8 and 15 pointed sectasetae dorsally, respectively. Caudal plate dorsally with one row of 14– 18 pointed sectasetae along anterior margin, a row of 13–15 pointed sectesetae submedially, two groups of 2–3 pointed sectasetae on each side in the middle and one group of 4–6 pointed sectasetae on each side subapically, anterior to circumanal ring. Caudal plate large; anterior margin remote from anterior margin of extra pore fields. Extra pore fields forming continuous outer and inner bands, consisting of large and small oval and round patches; outer band relatively long medially, end pointing outwards. Circumanal ring small.

**Host plant.** *Pleroma granulosum* (Desr.) D.Don (Melastomataceae).

**Distribution.** Brazil (RJ).

**Derivation of name.** Named after its host, *P. granulosum*.

**Comments.** *Melanastera granulosi* sp. nov. resembles *M. macaireae* sp. nov. and *M. tijuca* sp. nov. in the narrow, yellow forewing with light stripes along the veins, from which it differs in the longer metatibia (MT 0.6 versus 0.4; MT/HW 1.1 versus < 0.9) and the female terminalia (FP 0.6 versus 0.4).

### 63 Melanastera sebiferae sp. nov

(Figs 13L, 20H, 36H)

**Type material. Holotype f£: Brazil: AMAZONAS:** Manaus, Sede Embrapa, Campo Experimental, AM-010, km 29, S2.8950, W59.9733, 100 m, 27–30.iv.2014, *Virola sebifera* (D. Burckhardt & D.L. Queiroz) 133(3) (UFPR; slide; [LSMelseb-63B]).

**Paratypes. Brazil: AMAZONAS:** 2 immatures, same as holotype but (NHMB; slide, 70% ethanol; NMB-PSYLL0008067, NMB-PSYLL0008081 [LSMelseb-63B]).

**Description. *Adult.*** Coloration. Brownish yellow; head and thorax with dark brown dots. Antenna pale yellow, segments 3–9 with dark brown apices, segment 10 entirely dark brown.

Mesopraescutum with pale brown patches at fore margin; mesoscutum with four broad pale brown longitudinal stripes, margined by dark brown dots, and one or two median rows of dots. Forewing (Fig. 20H) yellow, with irregular, moderately dense, dark brown dots; base and apex of pterostigma and apices of Rs, M_1+2_, M_3+4_, Cu_1a_ and Cu_1b_ dark brown. Femora with brown patches.

Structure. Head, in lateral view, inclined 45° from longitudinal axis of body. Vertex (Fig. 13L) trapezoidal, slightly shorter than half as wide, with imbricate microsculpture and microscopic setae. Thorax moderately arched, with microscopic setosity. Forewing (Fig. 20H) obovate, widest in the middle, broadly and unevenly rounded apically; wing apex situated in cell r_2_ near apex of M_1+2_; C+Sc weakly curved in distal third; pterostigma about as wide as cell r_1_ in the middle, widening to apical third; Rs weakly convex in basal two thirds, obliquely curved to fore margin apically; M longer than M_1+2_ and M_3+4_; Cu_1a_ irregularly convex, ending slightly proximal to M fork; surface spinules present in all cells, leaving very narrow spinule- free stripes along the veins, forming hexagons of a single row of spinules, spinules absent from base of cell c+sc. Hindwing with 4 + 3 grouped costal setae. Metatibia bearing 8–9 grouped apical spurs, arranged as 4 + (4–5), anteriorly separated by several bristles.

Terminalia. Male. Unknown. – Female terminalia (Fig. 36H) cuneate; covered with setae.

Dorsal margin of proctiger, in lateral view, with a shallow hump distal to circumanal ring, concave in apical half, apex pointed, upturned; in dorsal view, apex subacute; circumanal ring, in dorsal view, cruciform. Subgenital plate, in lateral view, abruptly narrowing in apical third, apex pointed; in ventral view, apex blunt.

***Fifth instar immature.*** Coloration. Pale yellow; antenna pale brown with dark brown apical segment; cephalothoracic sclerite, wing pads, legs and caudal plate pale brown. Structure. Eye with one tiny, simple ocular seta dorsally. Antennal segments with following numbers of pointed sectasetae: 1(0), 2(1), 3(0), 4(2), 5(0), 6(2), 7(1), 8(1), 9(0), 10(0). Forewing pad with 9–11 large marginal and two smaller dorsal pointed sectasetae; hindwing pad with three large marginal and one smaller dorsal pointed sectasetae. Tarsal arolium broadly fan-shaped apically, 1.5 times as long as claws. Abdomen with two lateral pointed sectasetae anterior to circumanal ring. Caudal plate with anterior margin remote from anterior margin of extra pore fields; with two pointed sectasetae (one lateral and one sublateral) at anterior margin on either side, six pointed sectasetae (situated on tubercles in groups of 3 + 3) on either side laterally and more posteriorly, and five pointed subapical sectasetae near circumanal ring on either side dorsally. Extra pore fields forming continuous outer and inner bands, consisting of small oval and round patches; outer band moderately long medially, end pointing outwards. Circumanal ring small.

**Host plant.** *Virola sebifera* Aubl. (Myristicaceae).

**Distribution.** Brazil (AM).

**Derivation of name.** Named after its host, *Virola sebifera*.

**Comments.** *Melanastera sebiferae* sp. nov. differs from the other three Brazilian congeners developing on *Virola* spp. as follows: from *M. falcata* sp. nov. in the relatively shorter and wider forewing (FL/FW < 2.1 versus > 2.5); from *M. virolae* sp. nov. in the dorsal margin of the female proctiger which is distinctly concave (versus straight) in apical third; and from *M. stanopekari* sp. nov. in the surface spinules of the forewing forming hexagons of a single (versus double) row of spinules.

## Discussion and conclusions

**Taxonomy.** The Paurocephalini are one of two tribes of Liviinae, which together with the Euphyllurinae and Neophyllurinae form the relatively small family Liviidae. While the Paurocephalini is predominantly native to the tropics, its sister taxon, the Liviini, is most species-rich in the north temperate regions (Burckhardt *et al*. 2023). Here, 60 new species of the Paurocephalini are described from Brazil (one with additional material from Trinidad), bringing the number of extant species of the tribe to 130. The new species belong to the genera *Klyveria* (1 sp.) and *Melanastera* (59 spp.), bringing the numbers of known extant species to three (both worldwide and in Brazil) for *Klyveria* and 69/60 (worldwide/Brazil) for *Melanastera*. While the morphological differences between the *Klyveria* species are clear, this is not always the case with *Melanastera*, as the morphological differences between the individual species are often small. The diagnostically most important structures are the distal segment of the aedeagus (in lateral and dorsal view) and the paramere. Often, the forewing (shape, venation, surface spinules and colour pattern) and the female terminalia in adults, as well as the chaetotaxy, the tarsal arolium and the shape of the additional pore fields on the caudal plate in the last instar immatures provide additional characters for species delimitation.

**DNA barcoding.** Some recent taxonomic studies have successfully used molecular methods to diagnose psyllid species (Martoni *et al*. 2018; Percy 2018; Cho *et al*. 2020; Pramatarova *et al*. 2024). For the present study, two mitochondrial genes, *COI* and *cytb*, were sequenced for most Brazilian Paurocephalini species. The 3% maximum intraspecific genetic divergence criterion for *COI* (Martoni *et al*. 2018) characterised most *Melanastera* species.

*Cytb* often showed a relatively higher genetic variation, and we propose the uncorrected p-distance threshold of 5% below which samples of *Melanastera* can be considered conspecific. Although the genetic divergence criterion generally helped to delimit species, we found a high degree of intraspecific variation in some species of *Melanastera*. In *M. falcata* sp. nov., the *COI* and *cytb* sequences differed by up to 7.4% and 10%, respectively. As this species is morphologically well diagnosable and homogeneous, we interpret the molecular differences as intraspecific variation. A similar situation was found in *M. obscura* sp. nov. where *cytb* sequences diverged by almost 9% between samples, and in *M. tristis* sp. nov. with genetic differences reaching 7.45% for *COI* and 10.2% for *cytb*, but the morphology is homogeneous in both species. Extremely high sequence diversity was found within a single sample of *M. amazonica* sp. nov. (14% for *COI* and 13.3% for *cytb*) and between samples from different host plants, *Annona foetida* and *Guatteria* spp. Again, we interpret these differences as intraspecific, as all samples are morphologically homogeneous, although the presence of cryptic species within these taxa cannot be completely ruled out.

In summary, the circumscriptions of species in Paurocephalini by DNA barcoding and by morphology were generally congruent. DNA barcoding thus provides a useful identification tool in this taxonomically difficult tribe, despite the relatively high genetic variation in some species.

**Host plant patterns.** Psyllids are generally monophagous or narrowly oligophagous, and related psyllid species tend to develop on related plant taxa (Hollis 2004; Hodkinson 2009; Burckhardt *et al*. 2014; Ouvrard *et al*. 2015). The three extant species of *Klyveria* are narrowly oligophagous on *Luehea* species (Malvaceae, Malvales, malvids, rosids) and, thus, fit the general pattern of psyllids. Confirmed or probable host plants (not all are identified to species) are known for 55 of the 60 Brazilian *Melanastera* species (Table 6). The largest number of species, i.e. 28, develop (or are likely to develop) on Melastomataceae (Myrtales, rosids), followed by 18 species on Annonaceae (Magnoliales, magnoliids), four species each on Asteraceae (Asterales, campanulids, asterids) and Myristicaceae (Magnoliales, magnoliids) and one species on Cannabaceae (Rosales, fabids, rosids). These host orders belong to unrelated groups of angiosperms (APG IV 2016; Stevens 2017; Zuntini *et al*. 2024). The hosts of the eight extant Neotropical species of *Melanastera* described so far are Melastomataceae (7 spp.) and Cannabaceae (1 sp.). The two Old World species of *Melanastera* develop on Malvaceae (Burckhardt *et al*. 2006a; He *et al*. 2024). Annonaceae and Myristicaceae have not yet been reported as hosts for Paurocephalini (Burckhardt & Mifsud 2003; Burckhardt *et al*. 2023). The wide range of unrelated host taxa within the same genus is atypical for Psylloidea (Ouvrard *et al*. 2015), but is shared with other genera of Paurocephalini such as *Diclidophlebia* Crawford, *Haplaphalara* Uichanco, *Lilaoshia* Burckhardt, Serbina & Malenovský and *Paurocephala* Crawford; in the sister tribe Liviini, however, the host range is homogeneous within each genus (Burckhardt *et al*. 2023, 2024b).

**TABLE 6.**
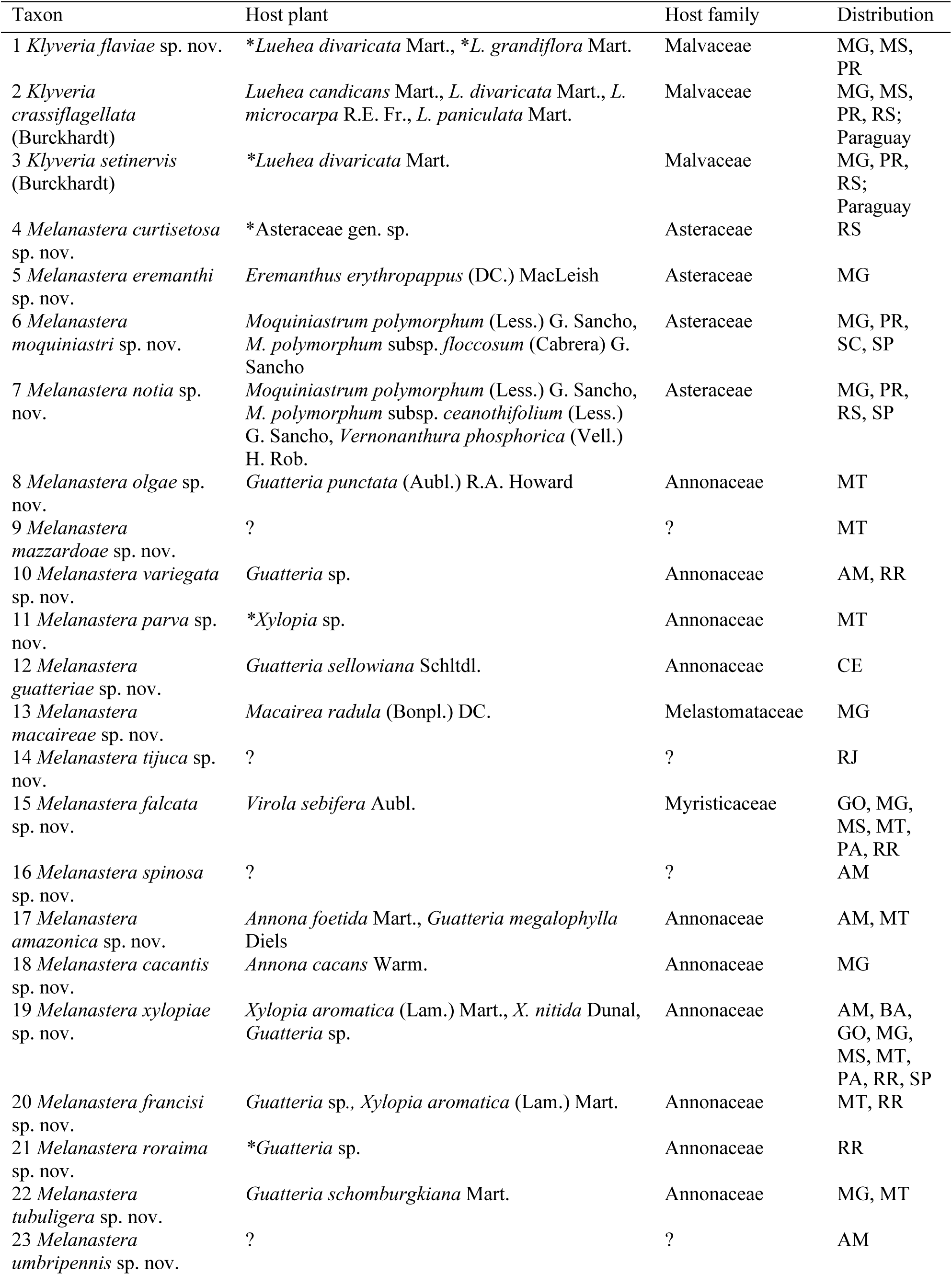
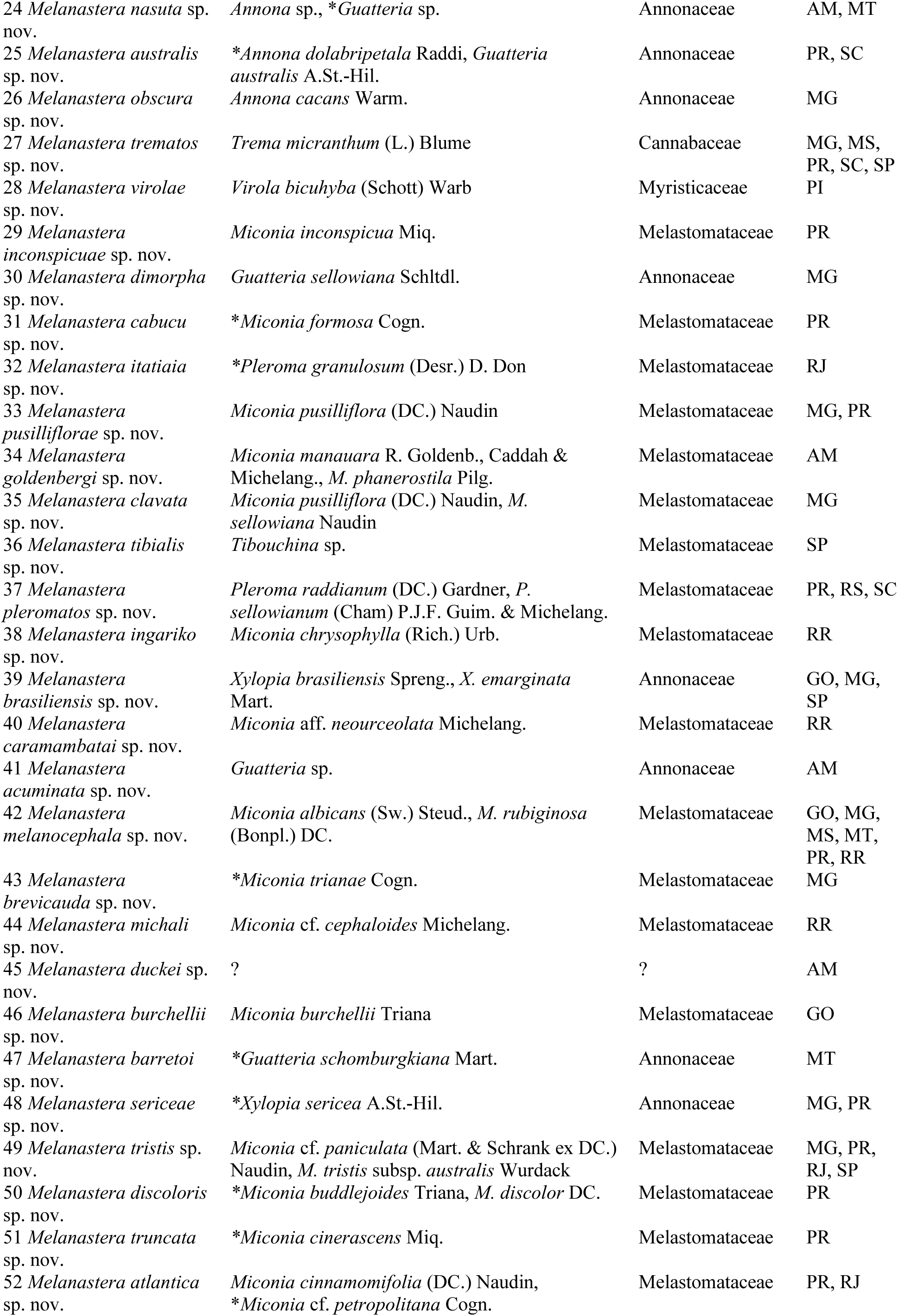
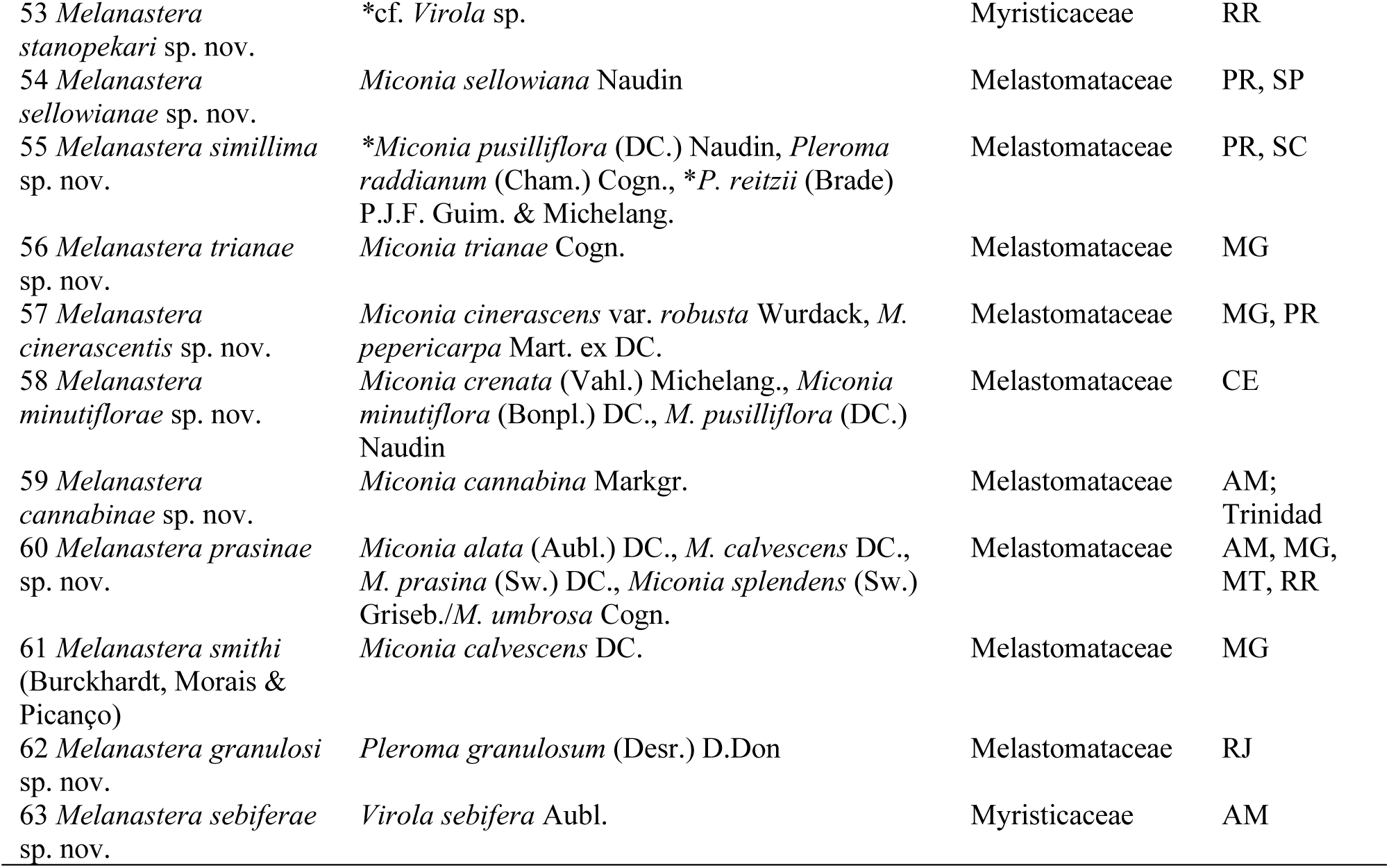
Brazilian Paurocephalinae with host plants (* indicates probable hosts not confirmed by the presence of immatures) and distribution (for abbreviations of Brazilian states see Material and methods).

Of the 55 Brazilian *Melanastera* species with known hosts, 36 species are monophagous, 13 are narrowly oligophagous and six are widely oligophagous. Among Melastomataceae feeders, 22 species are monophagous or narrowly oligophagous on *Miconia*, three on *Pleroma* spp. and one each on *Macairea* and *Tibouchina*, while one species is widely oligophagous on *Miconia* and *Pleroma*. Similar ratios are found among the Annonaceae feeders: eight species are monophagous or narrowly oligophagous on *Guatteria* and three each on *Annona* and *Xylopia* spp., while two species are widely oligophagous on *Annona* and *Guatteria* spp. and two on *Guatteria* and *Xylopia* spp. Three of the four Asteraceae-feeding species are monophagous on *Eremanthus*, *Moquiniastrum* and one unidentified species, while one is oligophagous on *Moquiniastrum* and *Vernonanthura* spp. The four species associated with *Virola* (Myristicaceae) and the one with *Trema* (Cannabaceae) are monophagous.

The term “superhost” has been coined for plants that harbour a rich fauna of gall- inducing insects (Santos *et al*. 2013). The term has also been used for some psyllid hosts, although they do not harbour only gall-inducing species. Examples include *Copaifera langsdorffii* (Fabaceae), which hosts 11 psyllid species from three genera, some of which induce galls, often with four or more species on the same tree (Burckhardt & Queiroz 2020), or *Mimosa scabrella* (Fabaceae) with five associated psyllid species (Burckhardt 2021). Among the *Melanastera* hosts, there are no species that could be considered as “superhost”. Although *Miconia pusilliflora* (Melastomataceae), the species with the highest number of associated Paurocephalini species in Brazil, hosts four species, these are mostly allopatric. In this context, *Miconia calvescens*, a noxious weed on some Pacific islands, is worth mentioning as it hosts three, also allopatric, species, *M. lucens* (Burckhardt, Hanson & Madrigal) in Costa Rica, and *M. prasina* sp. nov. and *M. smithi* (Burckhardt, Morais & Picanço) in Brazil.

**Biology.** Nothing is known about the biology of *Klyveria* spp. but there are studies on the biology of *Melanastera sellowianae* sp. nov. in Paraná (Barreto *et al*. 2020) and *M. smithi* in Minas Gerais (de Morais *et al*. 2008, 2013). Both species are polyvoltine with overlapping generations and population peaks during the cooler and drier months from April to July. This also seems to be the case for other *Melanastera* spp. (D. Burckhardt & D.L. Queiroz, pers. obs.). Adults and immatures of *M. smithi* have been found to attack buds, inflorescences and infrutescences (de Morais *et al*. 2008). Our field observations suggest that this is similar in other *Melanastera* species associated with Melastomataceae (Fig. 1C–F). Morais *et al*. (2013) showed that feeding by psyllids causes a collapse of the leaf epidermis. The presence of *M. melanocephala* sp. nov. on *Miconia albicans* is often indicated by severely deformed leaves (Fig. 40F). These deformations are probably due to physical damage to the leaf rather than physiologically induced galls. This is in contrast to *Melanastera* on Asteraceae, where immatures develop on the leaf blade (Fig. 39C–F) or in probably physiologically induced galls (Fig. 40A, B). The life history of *Melanastera* on Annonaceae, Cannabaceae and Myristicaceae is probably similar to that of species on Melastomataceae.

De Morais *et al*. (2008, 2013) observed syrphid larvae in small numbers predating on *M. smithi*, but never found parasitoids. Our observations (D. Burckhardt & D.L. Queiroz, pers. obs.) suggest that this is also the case for the other *Melanastera* spp. We never found mummies, which are often numerous in other psyllid taxa. In *M. moquiniastri* sp. nov. we observed that immatures produce large amounts of honeydew and are attended by two ant species, *Camponotus* sp. and *Crematogaster crinosa* (I. Malenovský, pers. obs.). Whether these ants only collect the honeydew or also prey on the psyllids remains unknown.

**Biogeography.** Species of *Klyveria* and *Melanastera* have been found in 15 Brazilian states (Table 6), and 60 species are endemic to the country. Two Brazilian species of *Klyveria* are also found in Paraguay and one of *Melanastera* in Trinidad. Of the 60 endemic species, 37 are restricted to a single state and 11 to two states. The most widespread species is *Melanastera xylopiae* sp. nov. which occurs in nine states, followed by *M. falcata* sp. nov. and *M. melanocephala* sp. nov. (6 states), and *M. trematos* sp. nov. (5 states).

The states with the highest number of Paurocephalini species are Minas Gerais (25 spp.) and Paraná (20 spp.). This may be due to the much greater collecting efforts compared to other states, but it also documents the biological richness of the two biomes, the Atlantic Forest and the Cerrado. Three states (Amazonas 12 spp., Mato Grosso 12 spp. and Roraima 11 spp.) with low collection effort have relatively many species, probably reflecting true species richness. The other states have fewer than 10 known species and are under-sampled. It is interesting to note that the four species associated with Asteraceae are restricted to southern and southeastern Brazil. A similar distribution is observed for *Klyveria*.

**Diversity.** The 60 new species of Paurocephalini described here increase the number of extant known species of this tribe by 1.7 times. With 67 extant species in the Americas, *Melanastera* becomes one of the most species-rich psyllid genera in the Neotropics after *Calophya* Löw (Calophyidae) (70 spp.) but before *Mitrapsylla* Crawford (Psyllidae) (51 spp.) (Ouvrard 2023). With 69 species, *Melanastera* is now also the most species-rich genus of Liviinae worldwide, followed by *Paurocephala* (46 spp.) (Burckhardt *et al*. 2023). In addition to the extant species, the Paurocephalini includes four species (two of which belong to *Melanastera*, one to *Klyveria* and one to *Diclidophlebia*) that have been described from Dominican amber (Burckhardt *et al*. 2024a).

In addition to the *Melanastera* species recognised here, other species are represented in the material at hand that are not formally described due to insufficient material. This, together with the fact that many regions in Brazil and other South American countries have not been explored or are insufficiently sampled, suggests that at least another 60 new species of Paurocephalinae can be expected in the Neotropical region. Priority for fieldwork should be given to areas at risk of destruction or degradation by human activities, such as the Amazon rainforest, the Atlantic Forest and the Cerrado.

## Acknowledgements

We are very grateful to R. Goldenberg (Universidade Federal do Paraná, Curitiba, PR), M.L. Brotto and J.T.W. Motta (Museu Botânico Municipal, Curitiba, PR) for identifying our plant specimens, to L. Beenken (ETH, Zürich) for identifying a sample of fungi, and M. Escárraga and L. Pedraza (UFPR, Curitiba) for identifying a sample with ants. We are indebted to P. Brown (BMNH), S.E. Halbert (FSCA) and J.A. Rafael (INPA) for the loan of material. We thank L. Dušátková and K. Křížová (Department of Botany and Zoology, Faculty of Science, Masaryk University, Brno) for their kind assistance in the molecular laboratory, E. Kvinikadze (Department of Botany and Zoology, Faculty of Science, Masaryk University, Brno) for help with preparation of slides, and D. Mathys (Swiss Nanoscience Institute, Nano Imaging, University of Basel) for his great help in taking SEM photos. The careful revision of an earlier manuscript version with useful suggestions by G. Taylor, Y.-C. Liao and D. Ouvrard is much appreciated. We gratefully acknowledge receiving the following collecting permits in Brazil: CNPq, IBAMA / SISBIO (11832; 37053). LŠS was partly funded by a grant from the Swiss National Science Foundation (SNSF; P2BSP3_168733).

---

**FIGURE 7.**
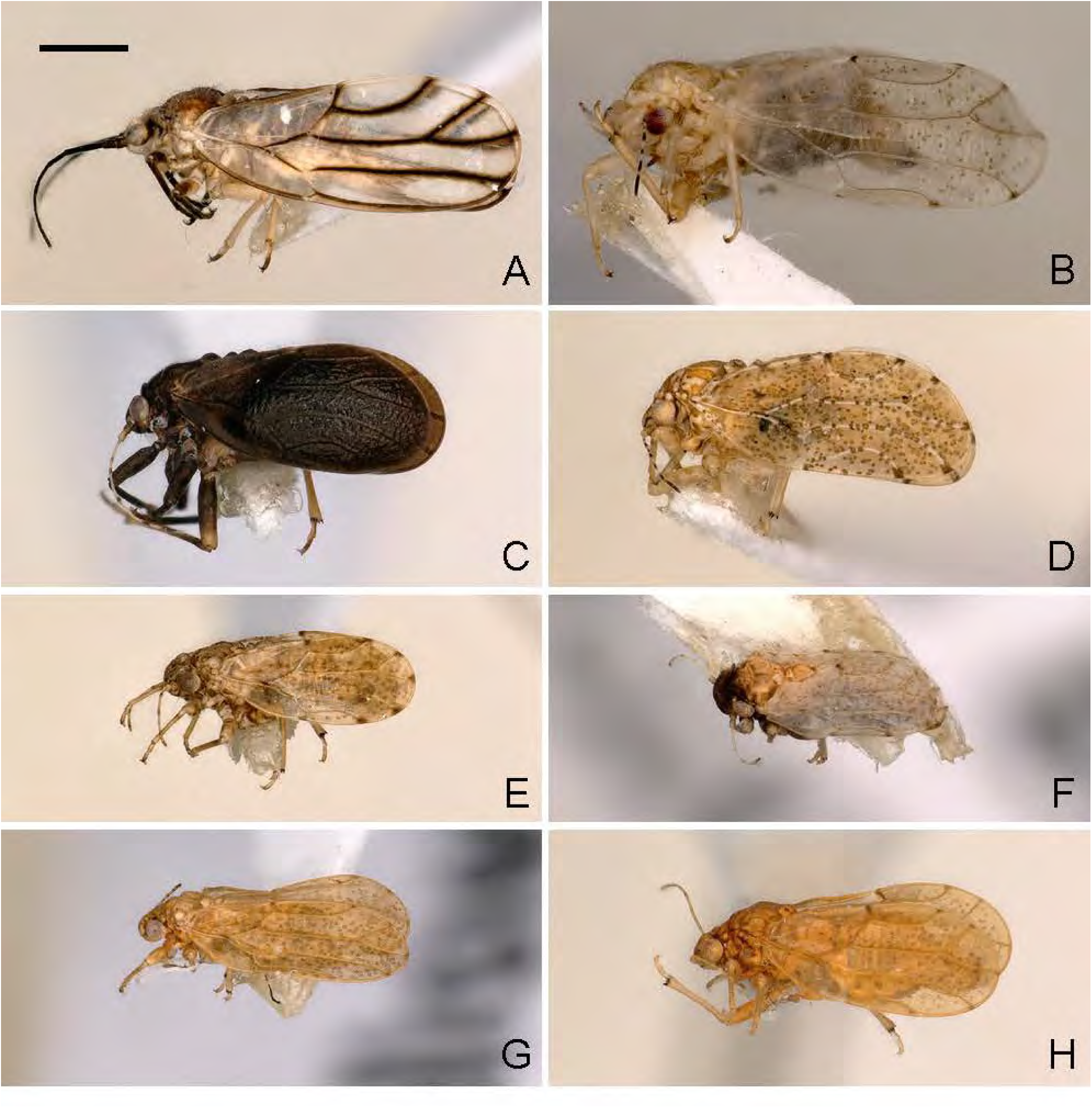
Habitus of adult Paurocephalini. A, *Klyveria crassiflagellata* (Burckhardt); B, *Melanastera moquiniastri* sp. nov.; C, *M. olgae* sp. nov.; D, *M. xylopiae* sp. nov.; E, *M. virolae* sp. nov.; F, *M. melanocephala* sp. nov.; G, *M. sellowianae* sp. nov.; H, *M. smithi* (Burckhardt, Morais & Picanço). Scale bar: 0.5 mm.

**FIGURE 8.**
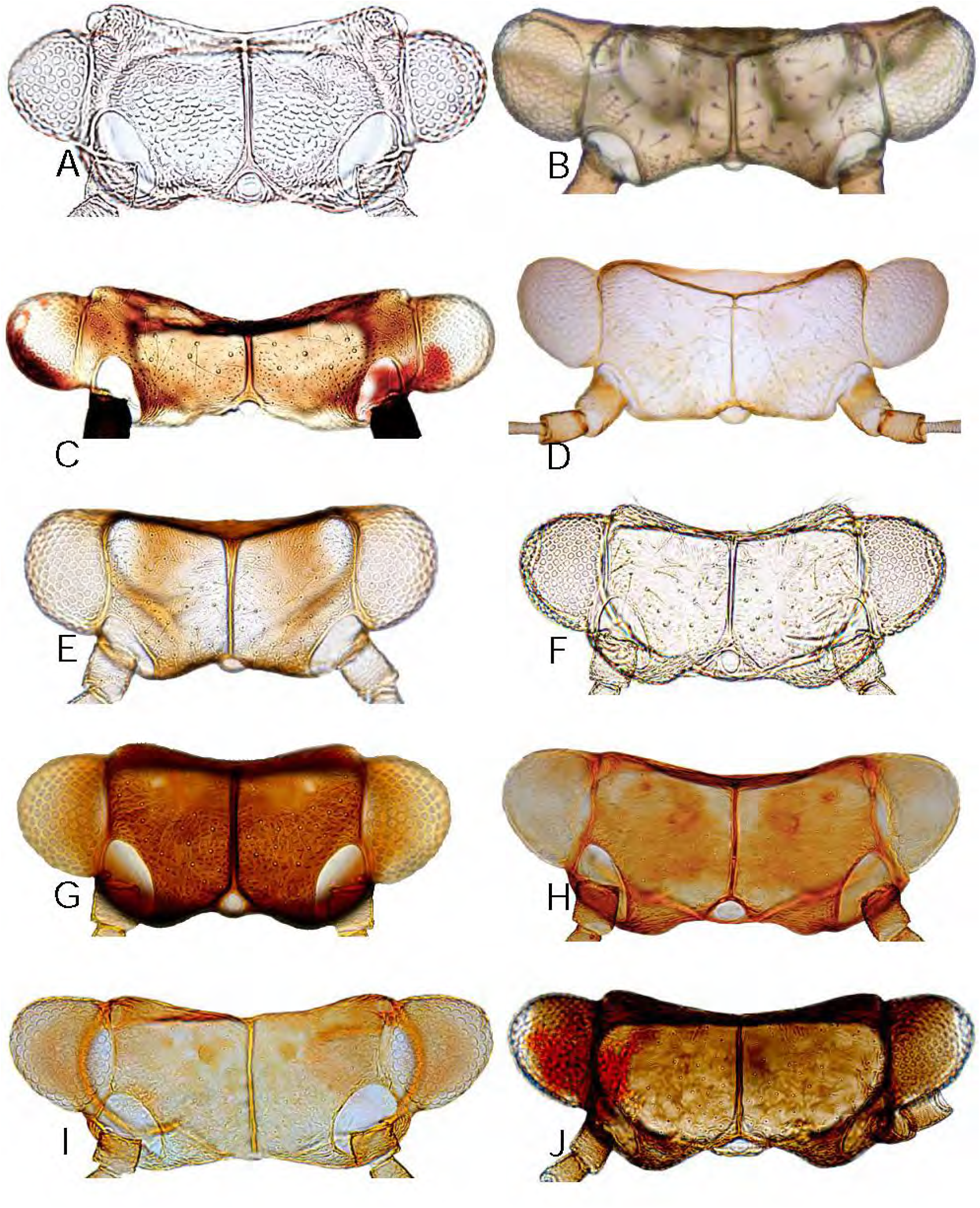
*Klyveria* and *Melanastera* spp., head, dorsal view. A, *K. flaviae* sp. nov.; B, *K. crassiflagellata* (Burckhardt); C, *K. setinervis* (Burckhardt); D, *M. curtisetosa* sp. nov.; E, *M. eremanthi* sp. nov.; F, *M. moquiniastri* sp. nov.; G, *M. notia* sp. nov.; H, *M. olgae* sp. nov.; I, *M. mazzardoae* sp. nov.; J, *M. variegata* sp. nov.

**FIGURE 9.**
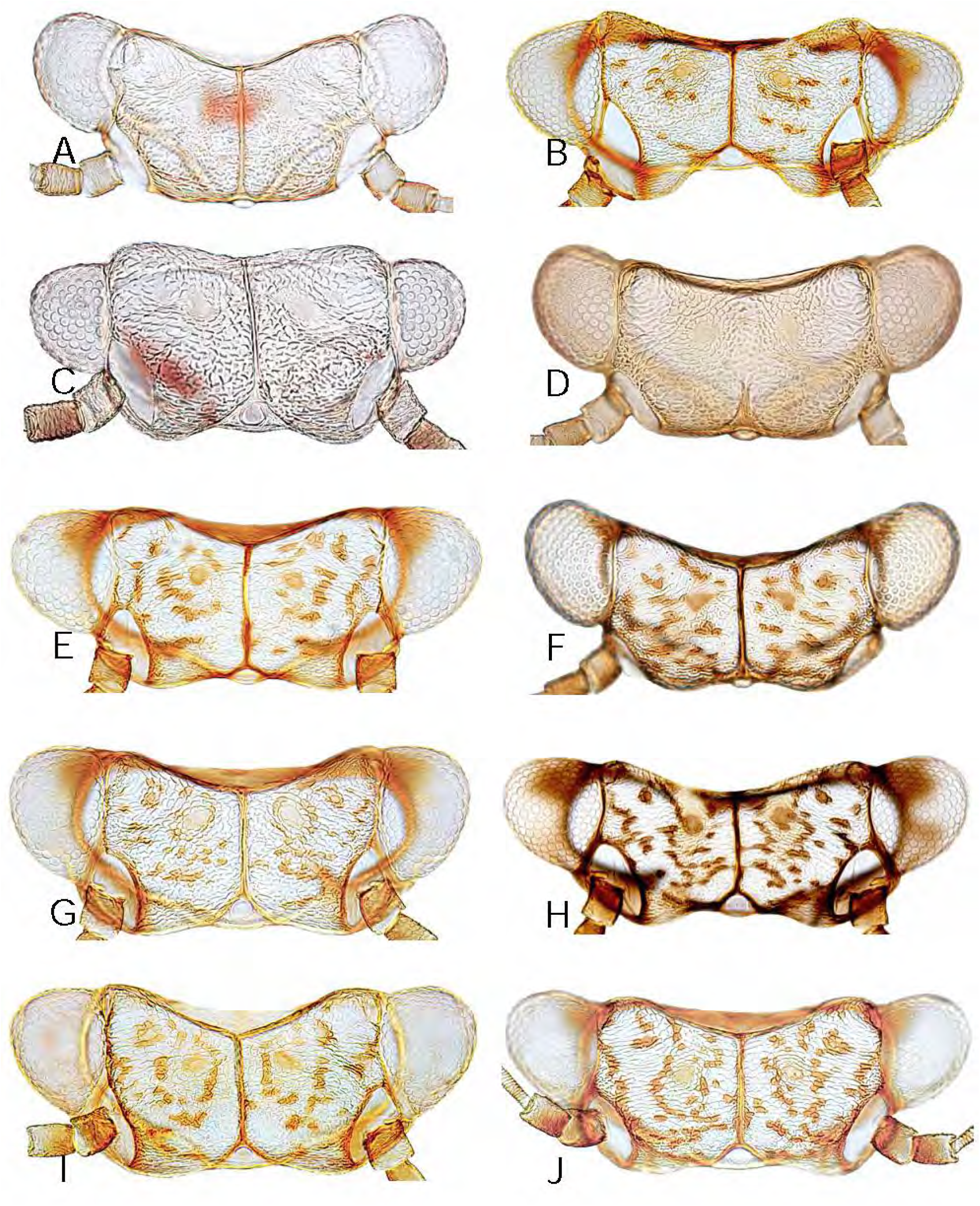
*Melanastera* spp., head, dorsal view. A, *M. parva* sp. nov.; B, *M. guatteriae* sp. nov.; C, *M. macaireae* sp. nov.; D, *M. tijuca* sp. nov.; E, *M. falcata* sp. nov.; F, *M. spinosa* sp. nov.; G, *M. amazonica* sp. nov.; H, *M. cacans* sp. nov.; I, *M. xylopiae* sp. nov.; J, *M. francisi* sp. nov.

**FIGURE 10.**
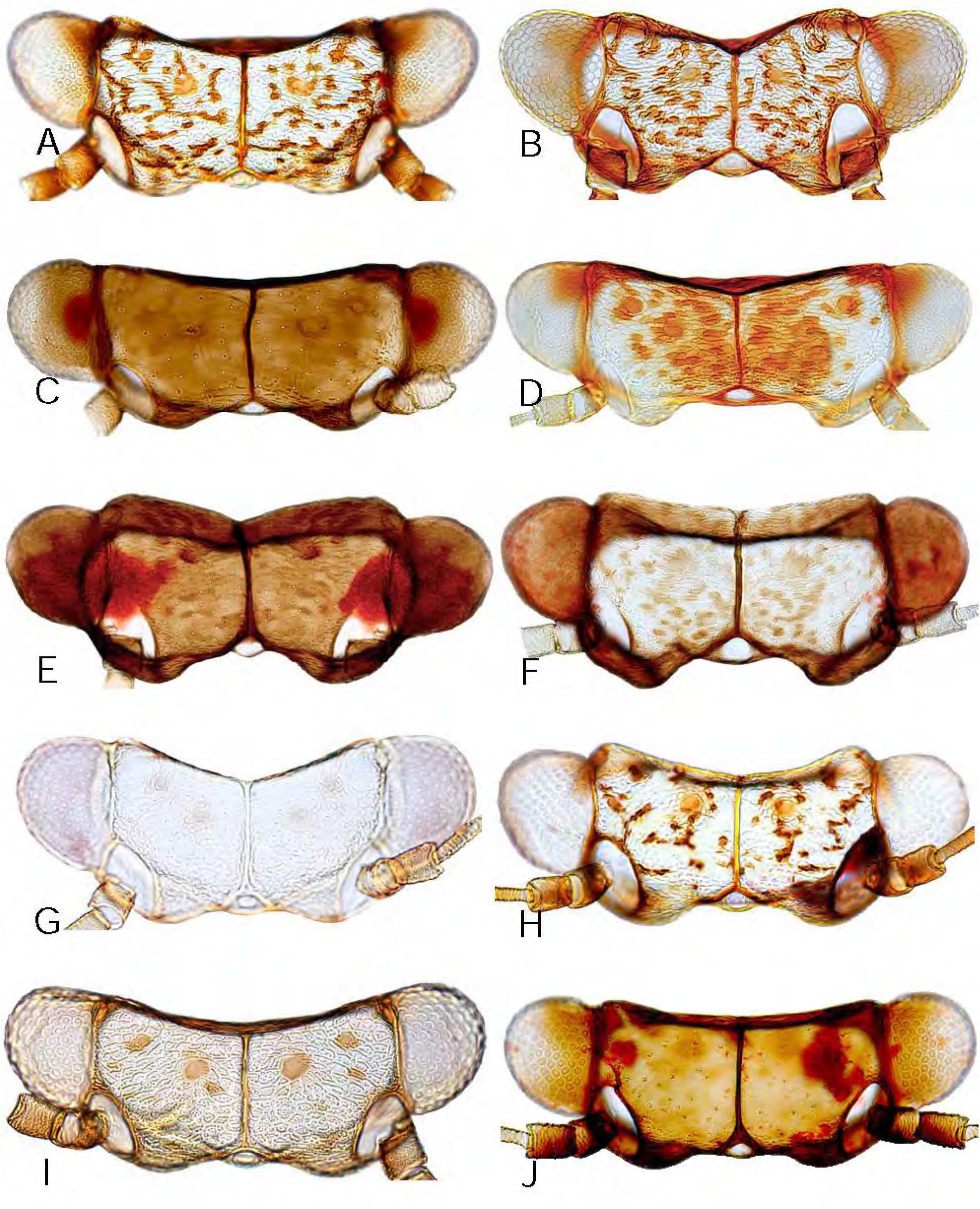
*Melanastera* spp., head, dorsal view. A, *M. roraima* sp. nov.; B, *M. tubuligera* sp. nov.; C, *M. umbripennis* sp. nov.; D, *M. nasuta* sp. nov.; E, *M. australis* sp. nov.; F, *M. obscura* sp. nov.; G, *M. trematos* sp. nov.; H, *M. virolae* sp. nov.; I, *M. inconspicuae* sp. nov.; J, *M. dimorpha* sp. nov., m$.

**FIGURE 11.**
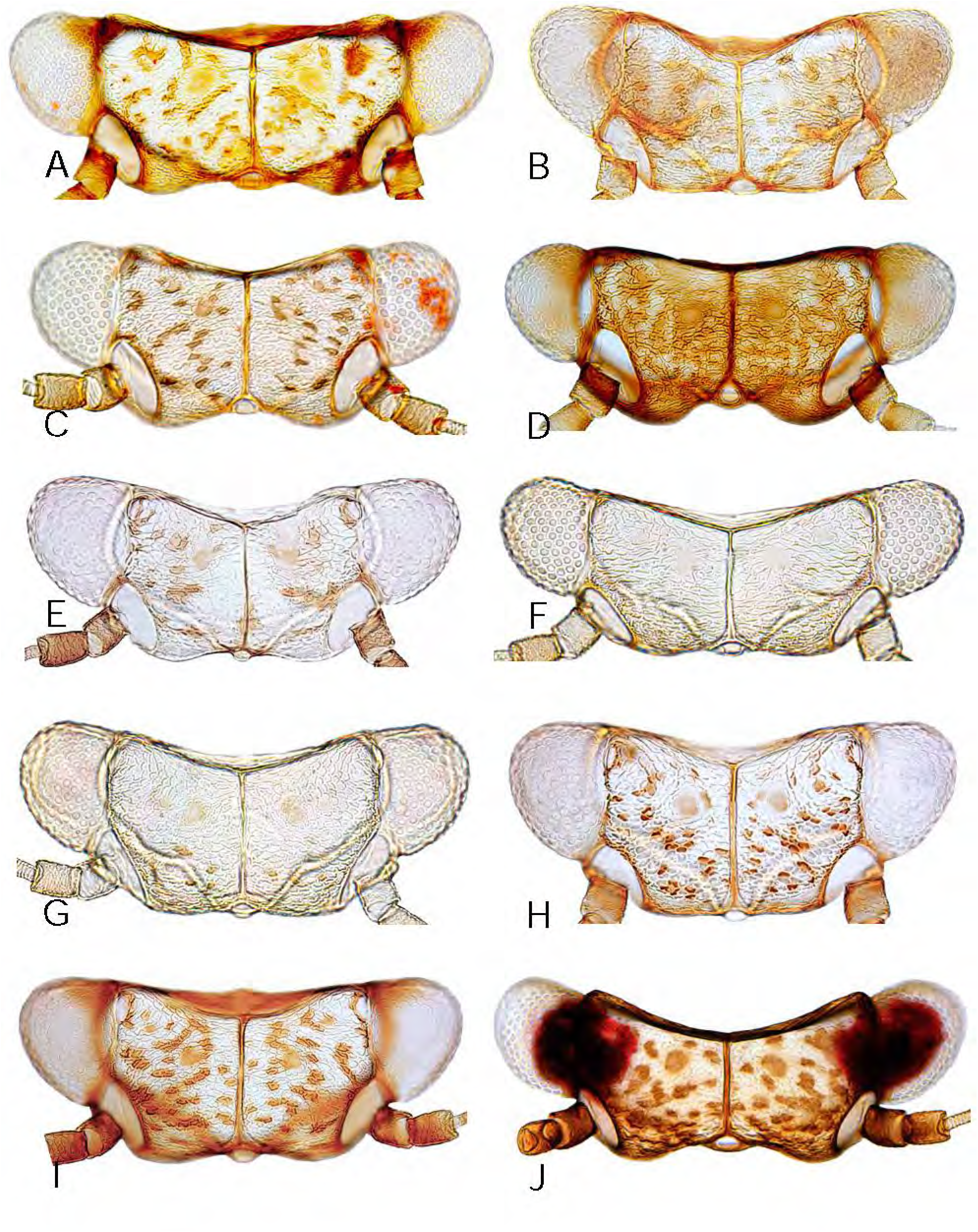
*Melanastera* spp., head, dorsal view. A, *M. dimorpha* sp. nov., f£; B, *M. cabucu* sp. nov.; C, *M. itatiaia* sp. nov.; D, *M. pusilliflorae* sp. nov.; E, *M. goldenbergi* sp. nov.; F, *M. clavata* sp. nov.; G, *M. tibialis* sp. nov.; H, *M. pleromatos* sp. nov.; I, *M. ingariko* sp. nov.; J, *M. brasiliensis* sp. nov.

**FIGURE 12.**
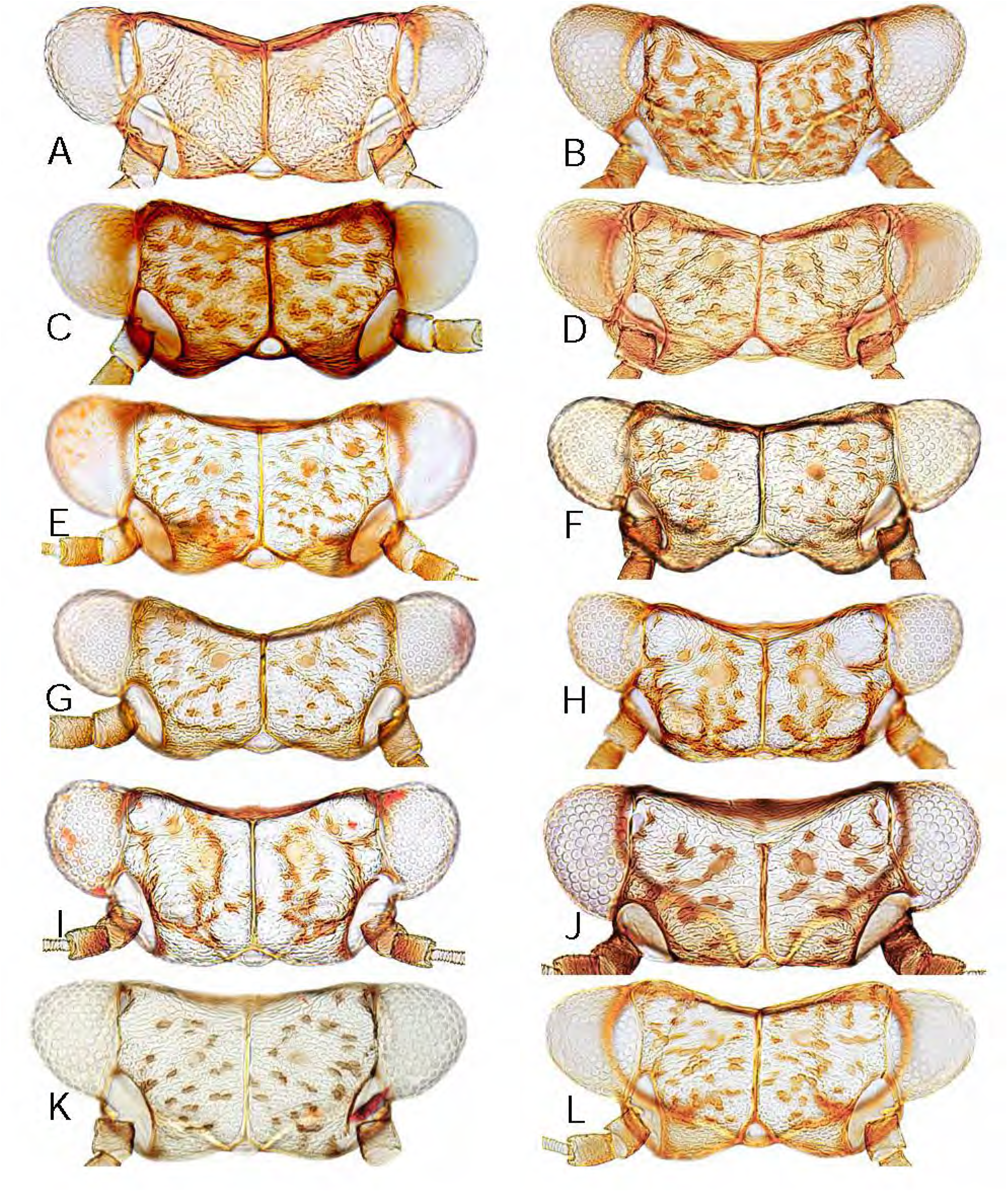
*Melanastera* spp., head, dorsal view. A, *M. caramambatai* sp. nov.; B, *M. acuminata* sp. nov.; C, *M. melanocephala* sp. nov.; D, *M. brevicauda* sp. nov.; E, *M. michali* sp. nov.; F, *M. duckei* sp. nov.; G, *M. burchellii* sp. nov.; H, *M. barretoi* sp. nov.; I, *M. sericeae* sp. nov.; J, *M. tristis* sp. nov.; K, *M. discoloris* sp. nov.; L, *M. truncata* sp. nov.

**FIGURE 13.**
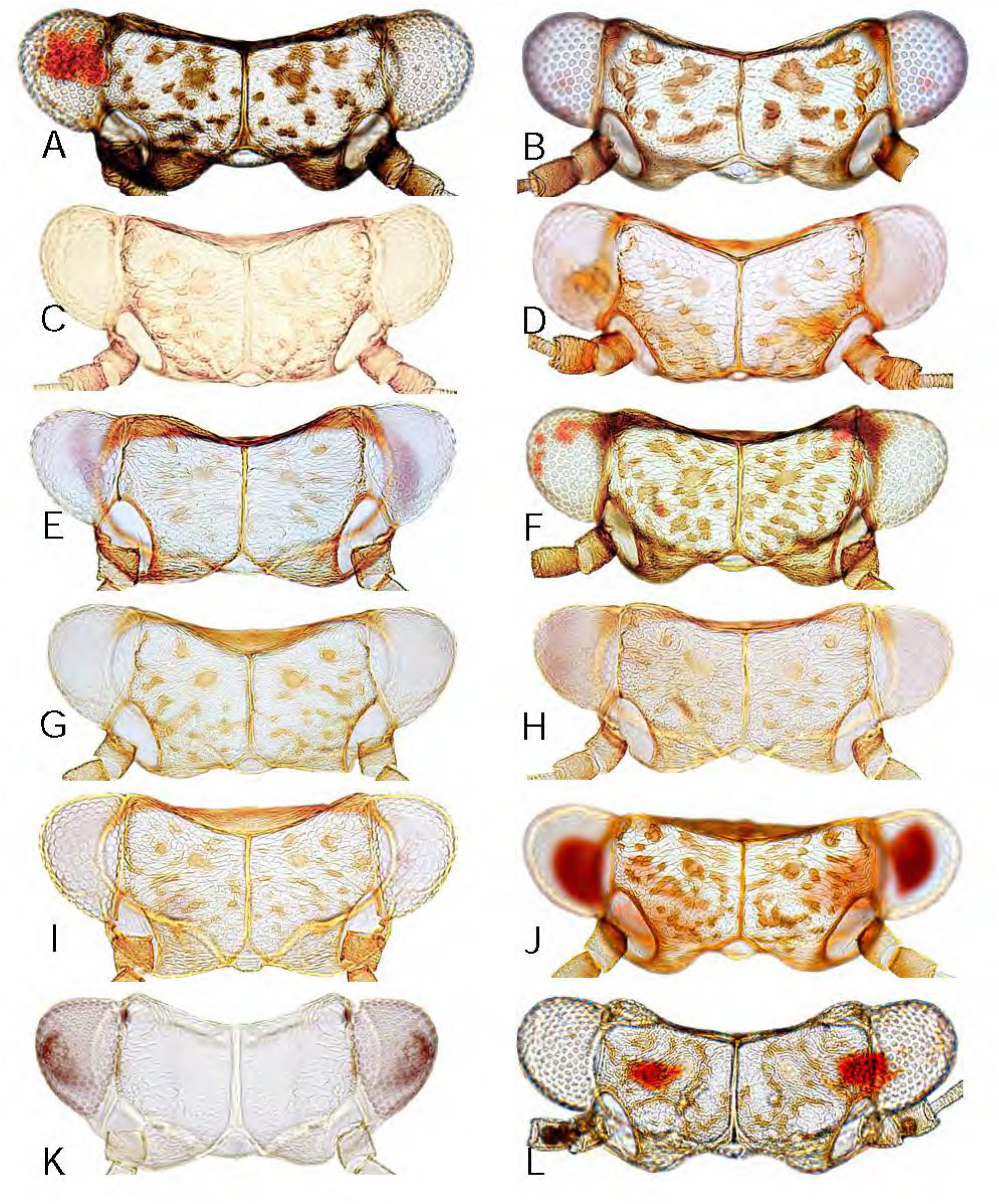
*Melanastera* spp., head, dorsal view. A, *M. atlantica* sp. nov.; B, *M. stanopekari* sp. nov; C, *M. sellowianae* sp. nov.; D, *M. simillima* sp. nov.; E, *M. trianae* sp. nov.; F, *M. cinerascentis* sp. nov.; G, *M. minutiflorae* sp. nov.; H, *M. cannabinae* sp. nov.; I, *M. prasinae* sp. nov.; J, *M. smithi* (Burckhardt, Morais & Picanço); K, *M. granulosi* sp. nov.; L, *M. sebiferae* sp. nov.

**FIGURE 14.**
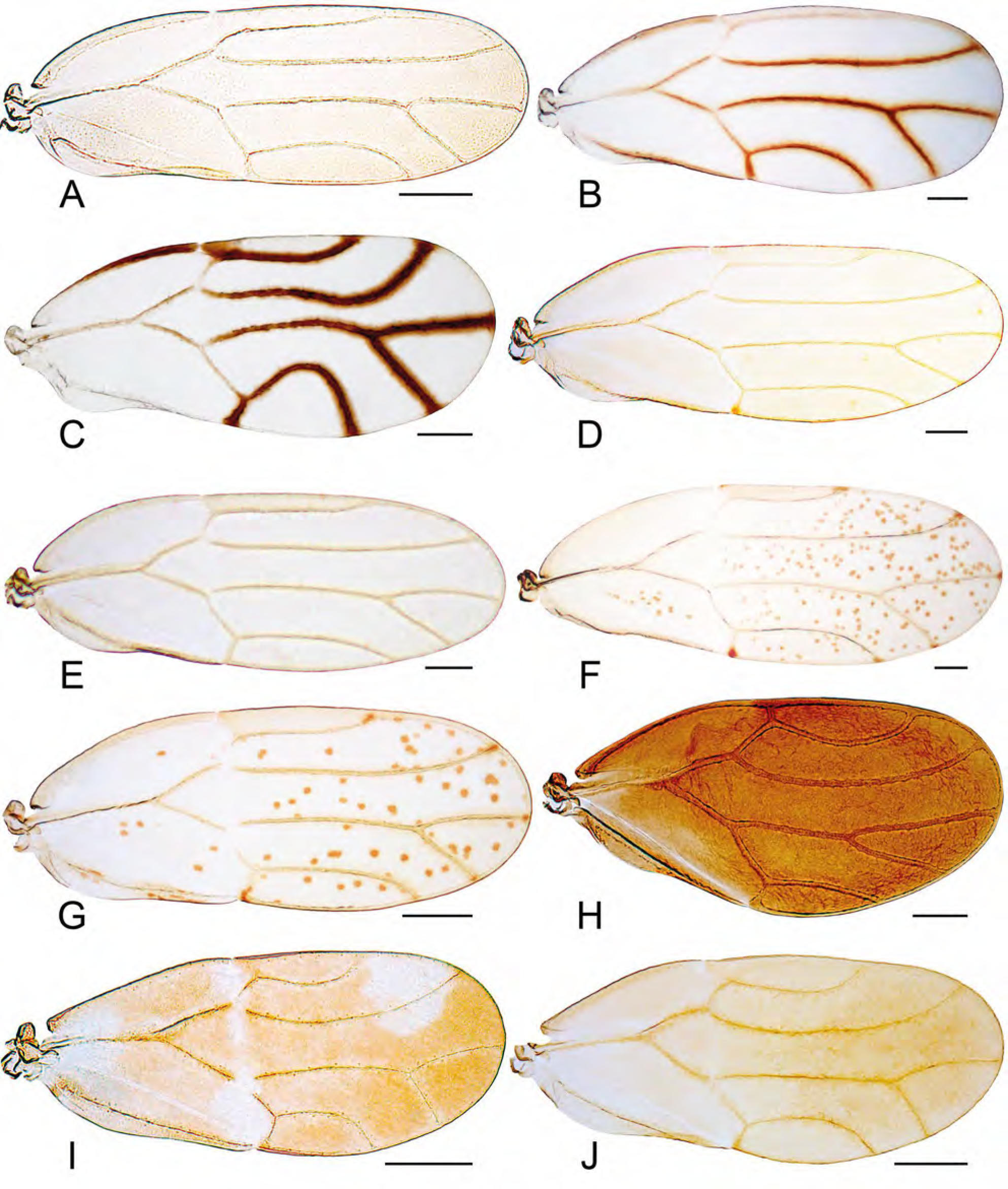
*Klyveria* and *Melanastera* spp., forewing. A, *K. flaviae* sp. nov.; B, *K. crassiflagellata* (Burckhardt); C, *K. setinervis* (Burckhardt); D, *M. curtisetosa* sp. nov.; E, *M. eremanthi* sp. nov.; F, *M. moquiniastri* sp. nov.; G, *M. notia* sp. nov.; H, *M. olgae* sp. nov.; I, *M. mazzardoae* sp. nov., m$; J, *M. mazzardoae* sp. nov., f£. Scale bars: 0.2 mm.

**FIGURE 15.**
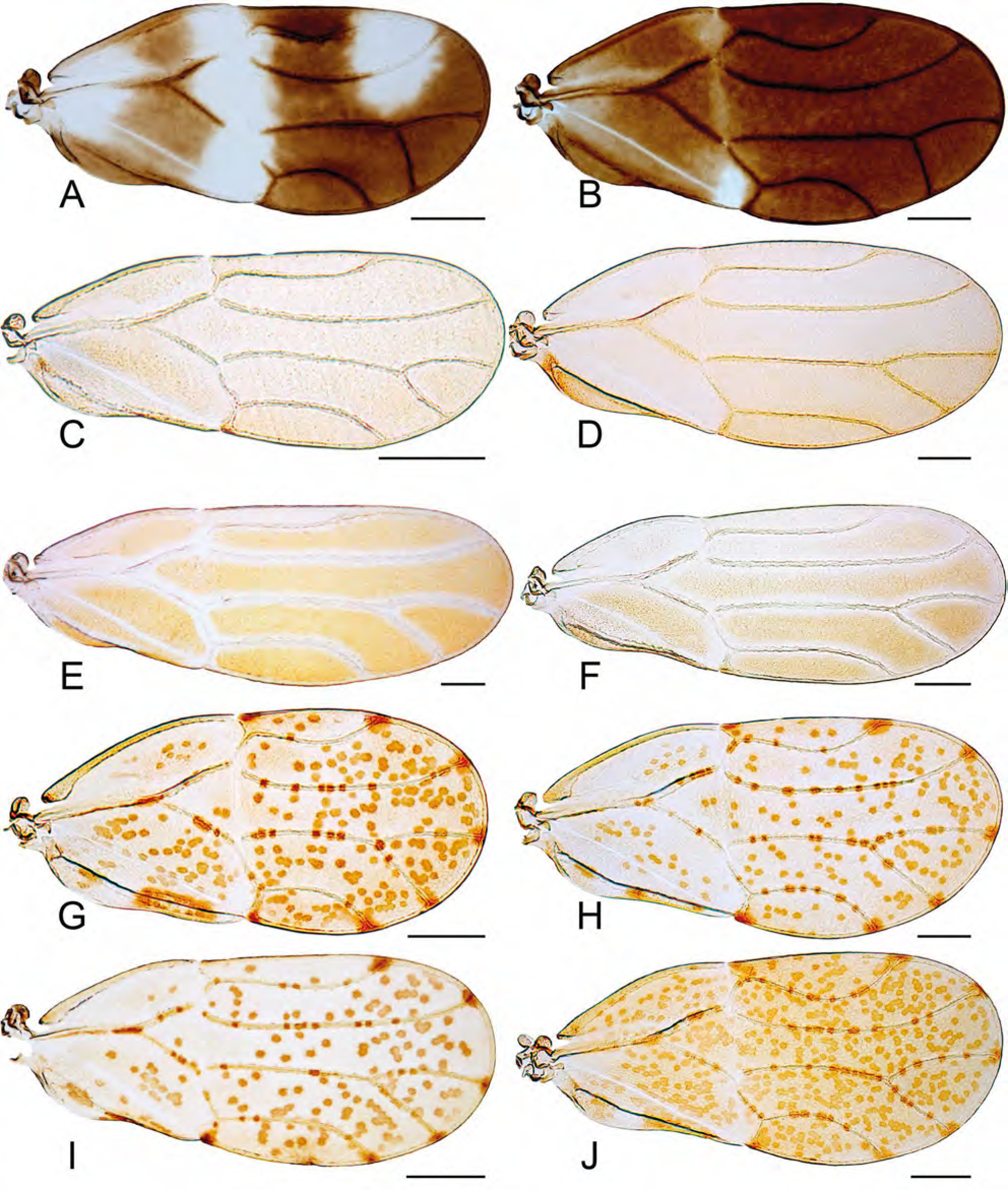
*Melanastera* spp., forewing. A, *M. variegata* sp. nov., m$; B, *M. variegata* sp. nov., f£; C, *M. parva* sp. nov.; D, *M. guatteriae* sp. nov.; E, *M. macaireae* sp. nov.; F, *M. tijuca* sp. nov.; G, *M. falcata* sp. nov., m$; H, *M. falcata* sp. nov., f£; I, *M. spinosa* sp. nov.; J, *M. amazonica* sp. nov. Scale bars: 0.2 mm.

**FIGURE 16.**
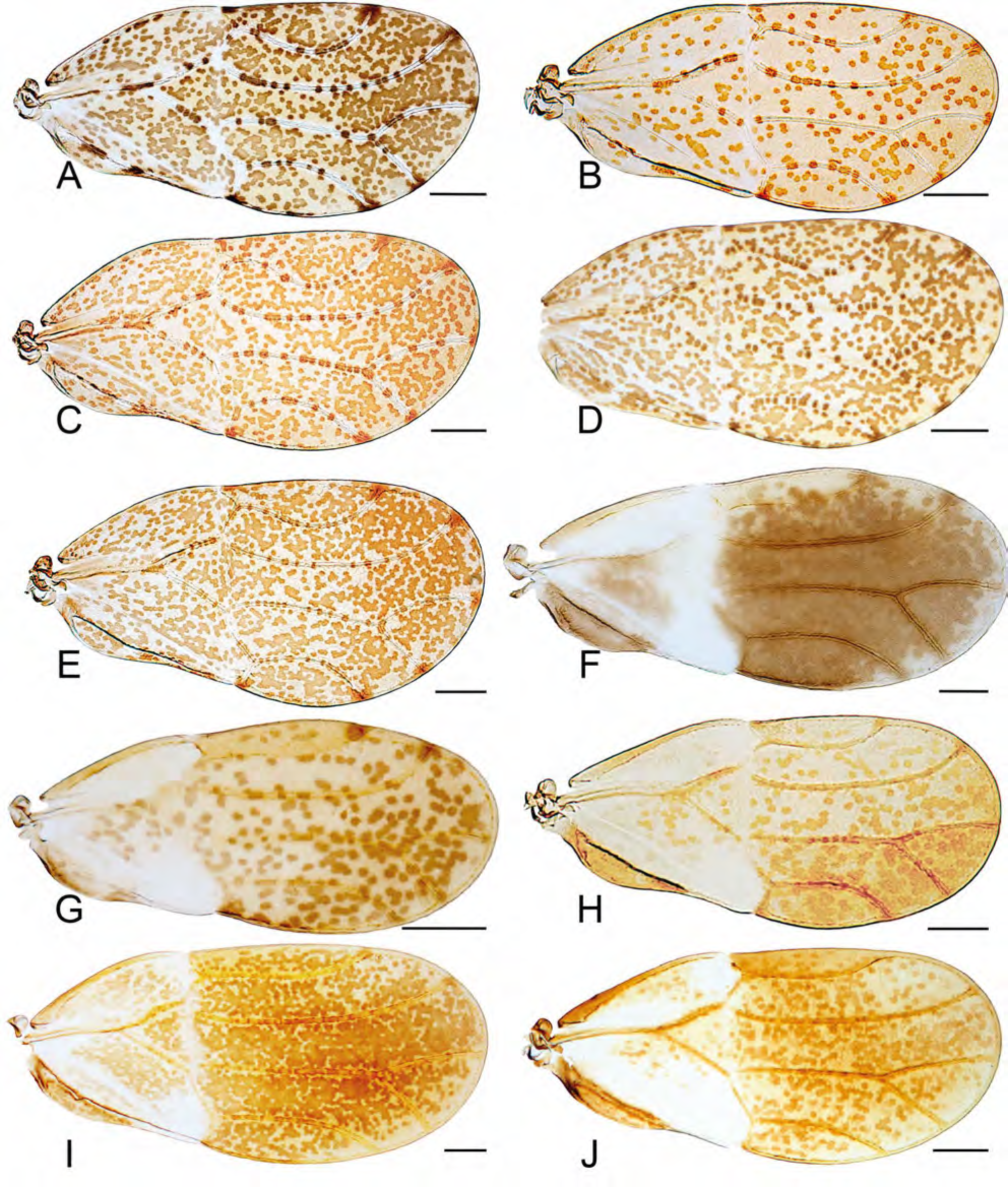
*Melanastera* spp., forewing. A, *M. cacans* sp. nov.; B, *M. xylopiae* sp. nov.; C, *M. francisi* sp. nov.; D, *M. roraima* sp. nov.; E, *M. tubuligera* sp. nov.; F, *M. umbripennis* sp. nov., m$; G, *M. umbripennis* sp. nov., f£; H, *M. nasuta* sp. nov.; I, *M. australis* sp. nov.; J, *M. obscura* sp. nov. Scale bars: 0.2 mm.

**FIGURE 17.**
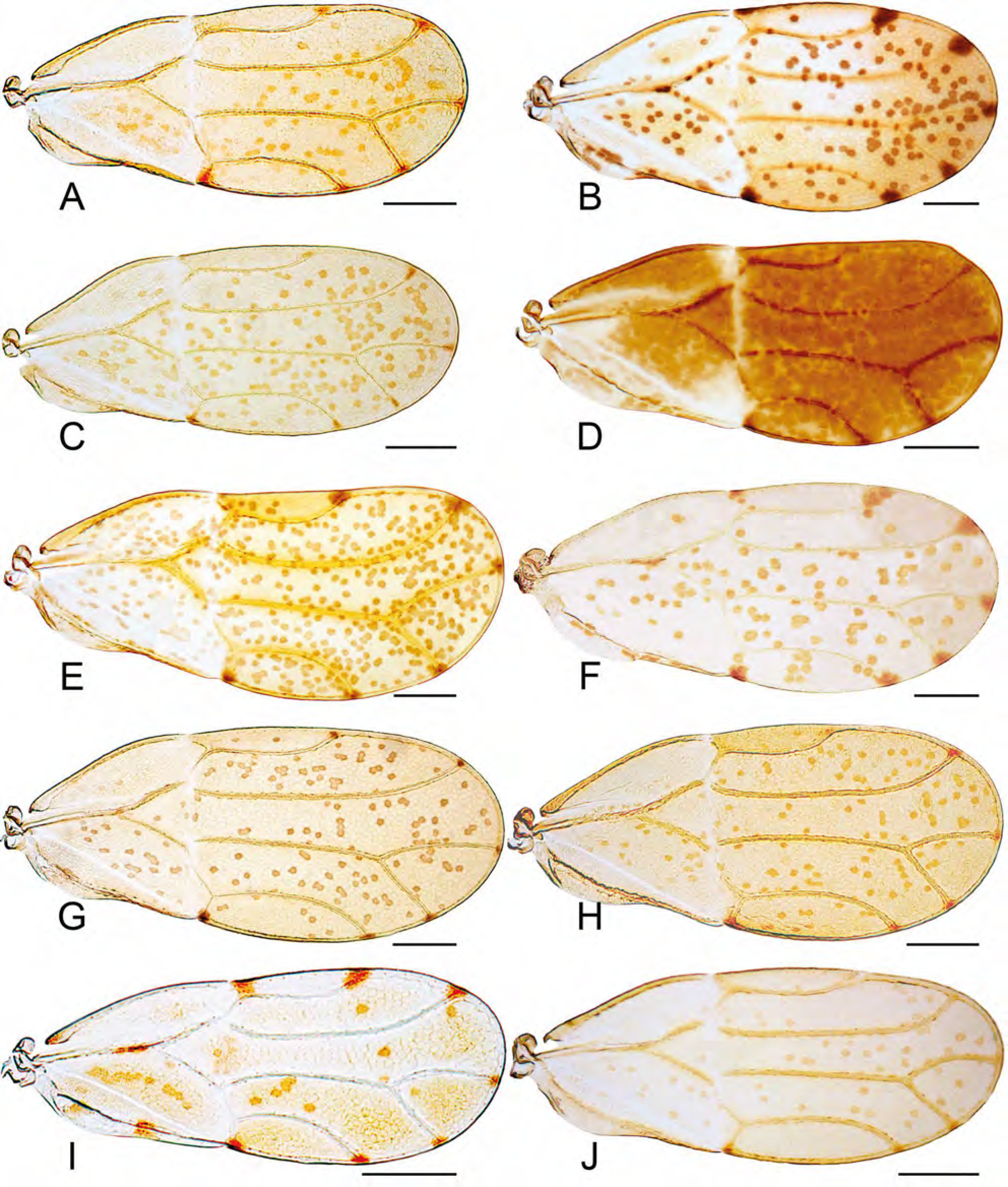
*Melanastera* spp., forewing. A, *M. trematos* sp. nov.; B, *M. virolae* sp. nov.; C, *M. inconspicuae* sp. nov.; D, *M. dimorpha* sp. nov., m$; E, *M. dimorpha* sp. nov., f£; F, *M. cabucu* sp. nov.; G, *M. itatiaia* sp. nov.; H, *M. pusilliflorae* sp. nov.; I, *M. goldenbergi* sp. nov.; J, *M. clavata* sp. nov. Scale bars: 0.2 mm.

**FIGURE 18.**
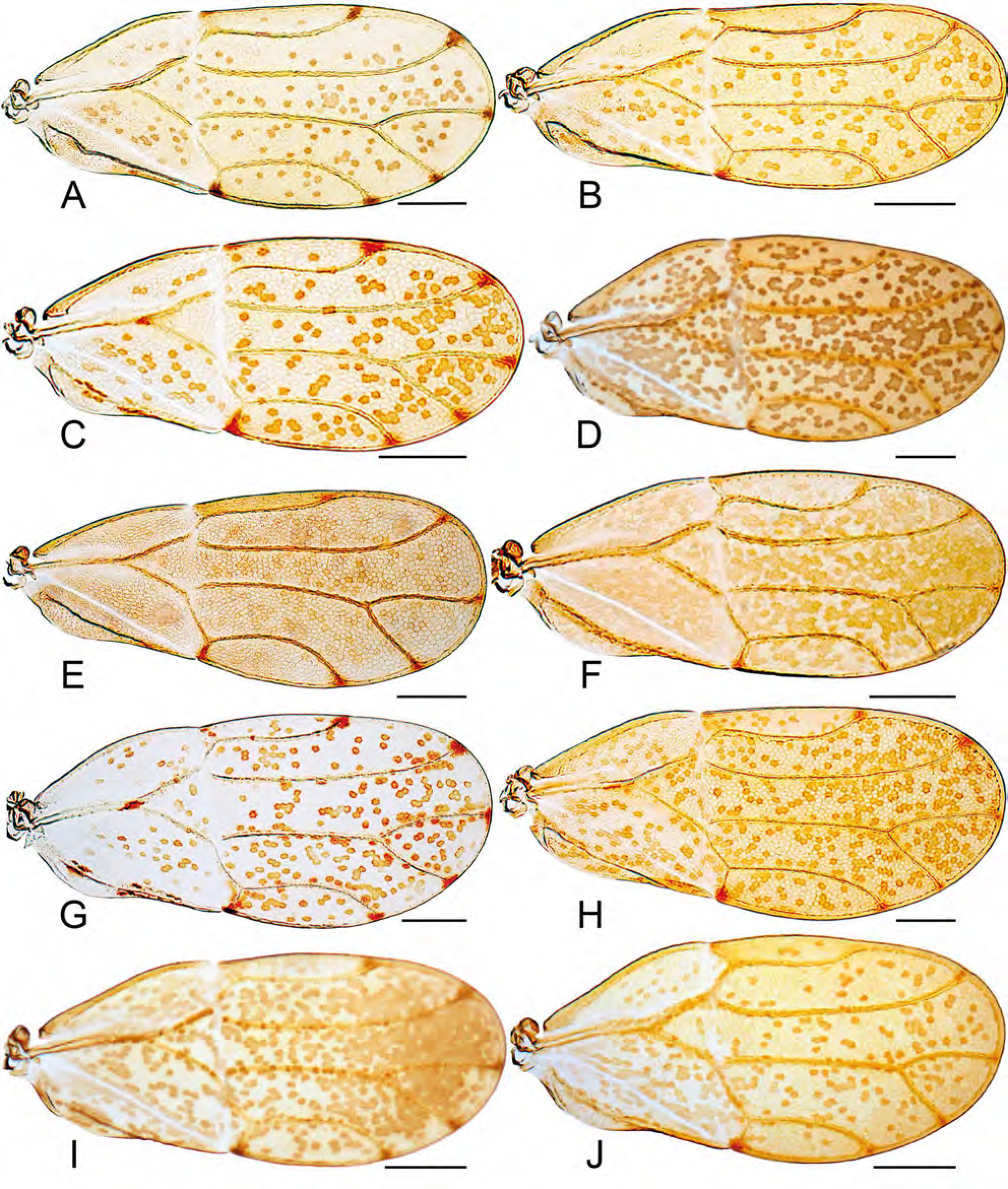
*Melanastera* spp., forewing. A, *M. tibialis* sp. nov.; B, *M. pleromatos* sp. nov.; C, *M. ingariko* sp. nov.; D, *M. brasiliensis* sp. nov.; E, *M. caramambatai* sp. nov.; F, *M. acuminata* sp. nov.; G, *M. melanocephala* sp. nov.; H, *M. brevicauda* sp. nov.; I, *M. michali* sp. nov.; J, *M. duckei* sp. nov. Scale bars: 0.2 mm.

**FIGURE 19.**
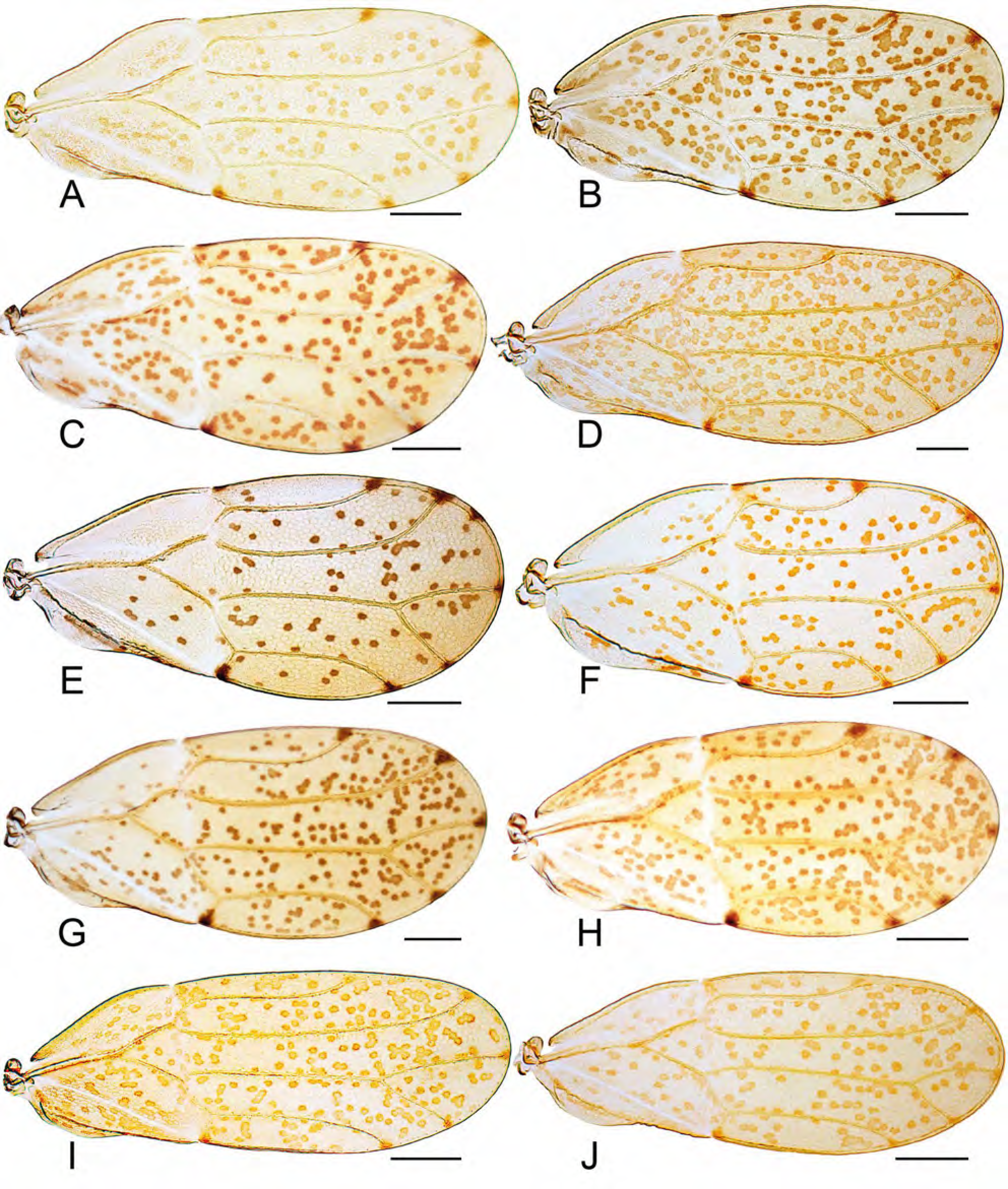
*Melanastera* spp., forewing. A, *M. burchellii* sp. nov.; B, *M. barretoi* sp. nov.; C, *M. sericeae* sp. nov.; D, *M. tristis* sp. nov.; E, *M. discoloris* sp. nov.; F, *M. truncata* sp. nov.; G, *M. atlantica* sp. nov.; H, *M. stanopekari* sp. nov.; I, *M. sellowianae* sp. nov.; J, *M. simillima* sp. nov. Scale bars: 0.2 mm.

**FIGURE 20.**
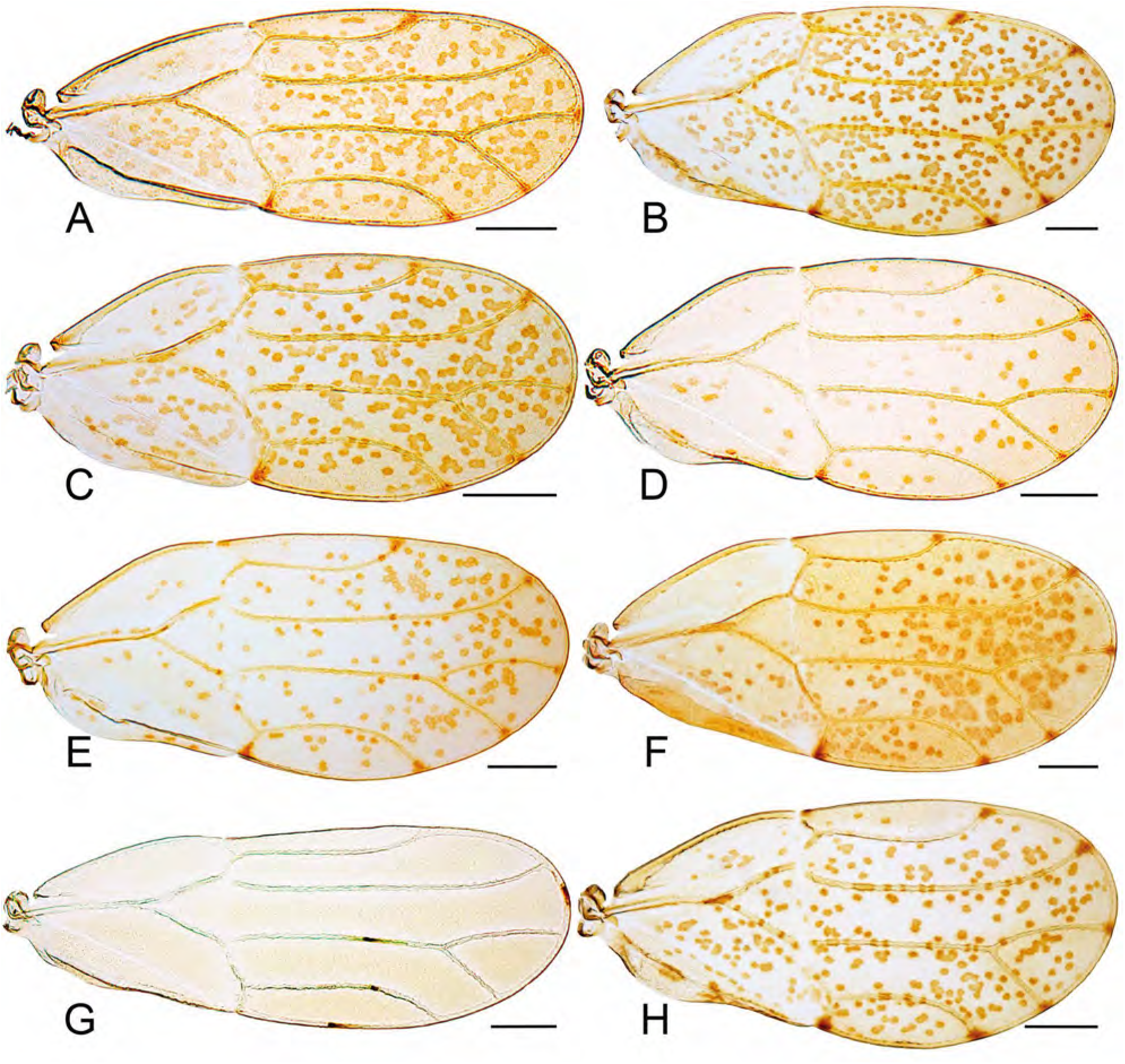
*Melanastera* spp., forewing. A, *M. trianae* sp. nov.; B, *M. cinerascentis* sp. nov.; C, *M. minutiflorae* sp. nov.; D, *M. cannabinae* sp. nov.; E, *M. prasinae* sp. nov.; F, *M. smithi* (Burckhardt, Morais & Picanço); G, *M. granulosi* sp. nov.; H, *M. sebiferae* sp. nov. Scale bars: 0.2 mm.

**FIGURE 21.**
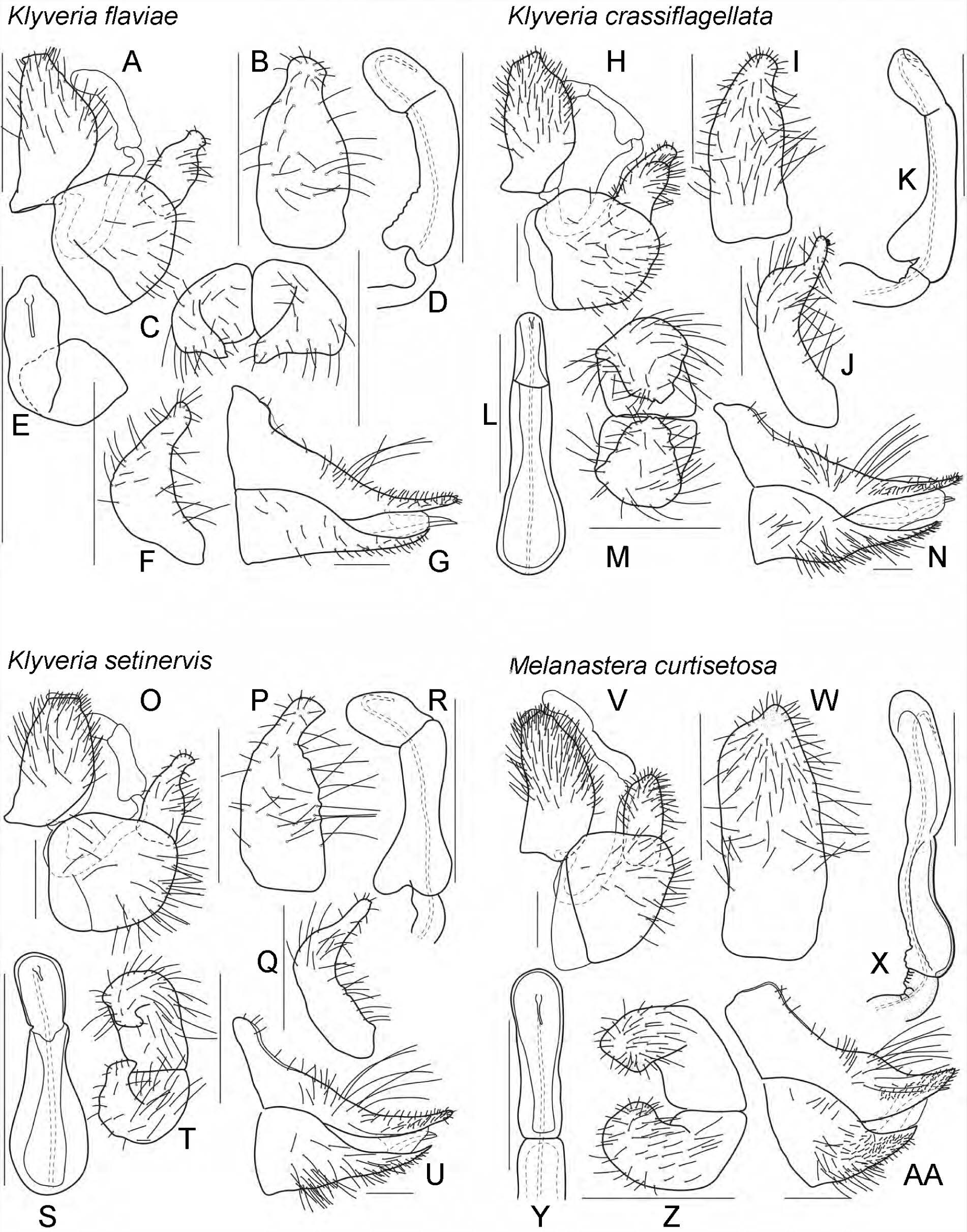
*Klyveria flaviae* sp. nov. A, m$ terminalia, lateral view; B, paramere, inner face, lateral view; C, paramere, dorsal view; D, distal segment of aedeagus, lateral view; E, distal segment of aedeagus, dorsal view; F, paramere, posterior view; G, f£ terminalia, lateral view. – *K. crassiflagellata* (Burckhardt). H, m$ terminalia, lateral view; I, paramere, inner face, lateral view; J, paramere, posterior view; K, distal segment of aedeagus, lateral view; L, distal segment of aedeagus, dorsal view; M, paramere, dorsal view; N, f£ terminalia, lateral view. – *K. setinervis* (Burckhardt). O, m$ terminalia, lateral view; P, paramere, inner face, lateral view; Q, paramere, posterior view; R, distal segment of aedeagus, lateral view; S, distal segment of aedeagus, dorsal view; T, paramere, dorsal view; U, f£ terminalia, lateral view. – 1. *M. curtisetosa* sp. nov. V, m$ terminalia, lateral view; W, paramere, inner face, lateral view; X, distal segment of aedeagus, lateral view; Y, distal segment of aedeagus, dorsal view; Z, paramere, dorsal view; AA, f£ terminalia, lateral view. Scale bars: 0.1 mm.

**FIGURE 22.**
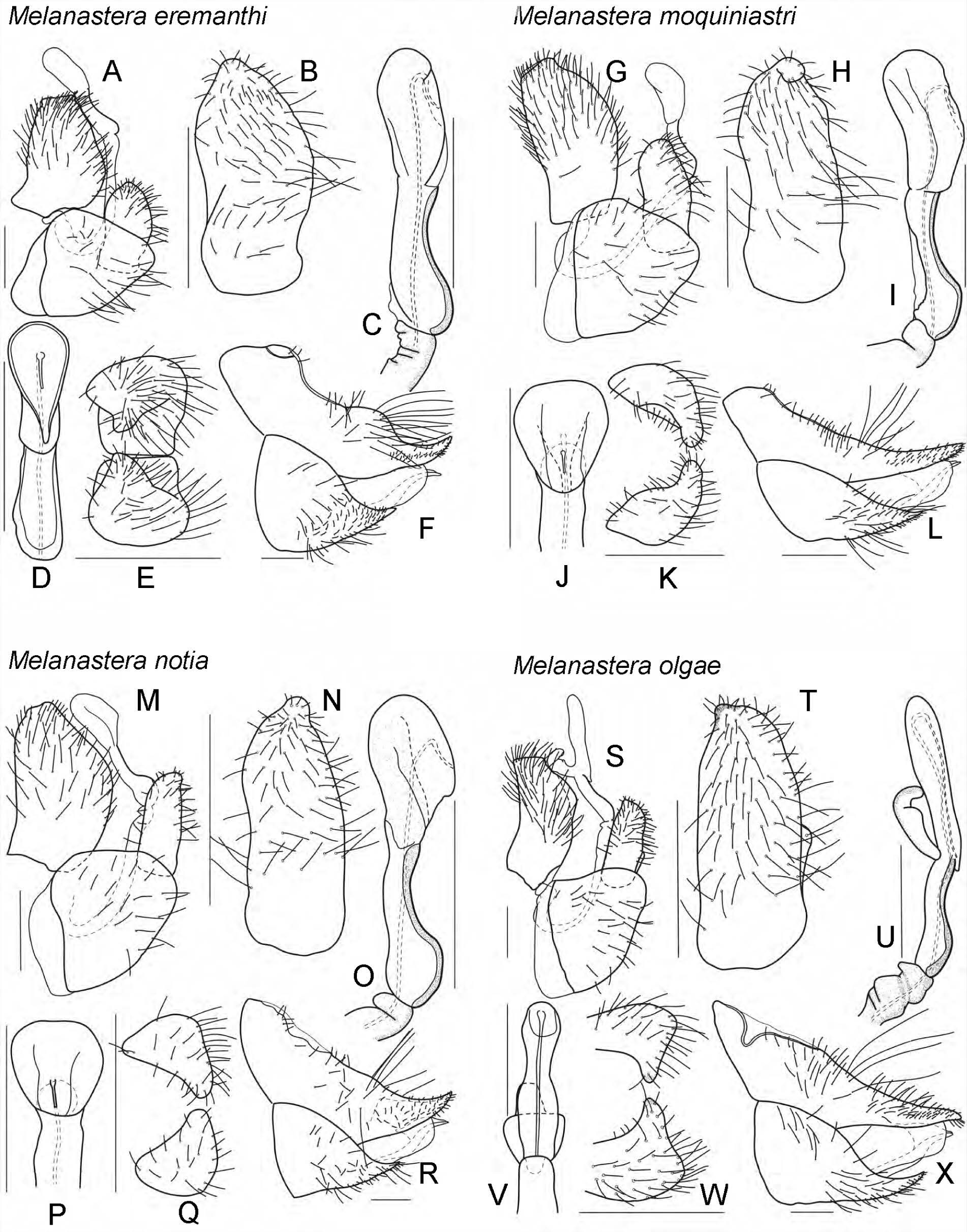
*Melanastera eremanthi* sp. nov. A, m$ terminalia, lateral view; B, paramere, inner face, lateral view; C, distal segment of aedeagus, lateral view; D, distal segment of aedeagus, dorsal view; E, paramere, dorsal view; F, f£ terminalia, lateral view. – *M. moquiniastri* sp. nov. G, m$ terminalia, lateral view; H, paramere, inner face, lateral view; I, distal segment of aedeagus, lateral view; J, distal segment of aedeagus, dorsal view; K, paramere, dorsal view; L, f£ terminalia, lateral view. – *M. notia* sp. nov. M, m$ terminalia, lateral view; N, paramere, inner face, lateral view; O, distal segment of aedeagus, lateral view; P, distal segment of aedeagus, dorsal view; Q, paramere, dorsal view; R, f£ terminalia, lateral view. – *M. olgae* sp. nov. S, m$ terminalia, lateral view; T, paramere, inner face, lateral view; U, distal segment of aedeagus, lateral view; V, distal segment of aedeagus, dorsal view; W, paramere, dorsal view; X, f£ terminalia, lateral view. Scale bars: 0.1 mm.

**FIGURE 23.**
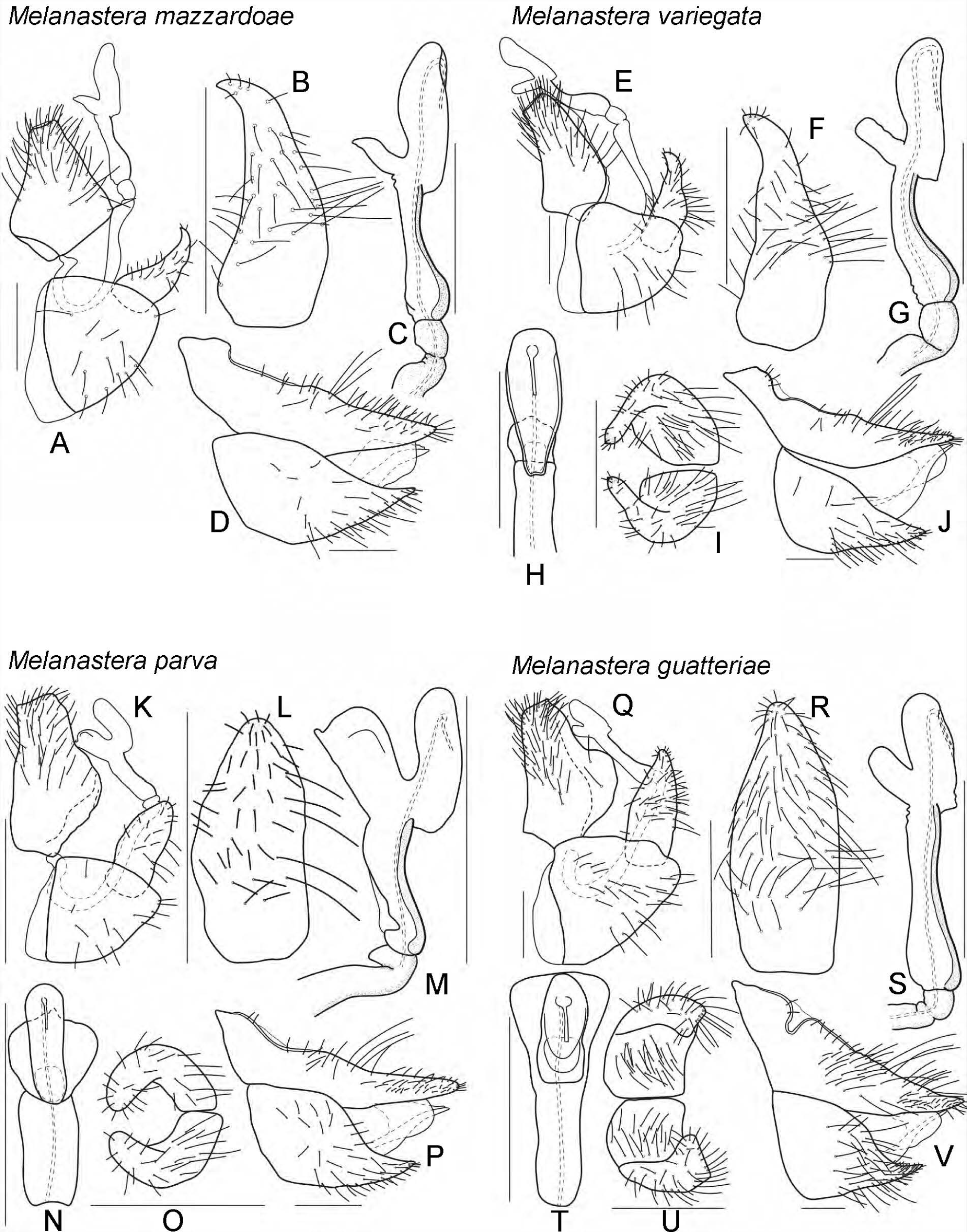
*Melanastera mazzardoae* sp. nov. A, m$ terminalia, lateral view; B, paramere, inner face, lateral view; C, distal segment of aedeagus, lateral view; D, f£ terminalia, lateral view. – *M. variegata* sp. nov. E, m$ terminalia, lateral view; F, paramere, inner face, lateral view; G, distal segment of aedeagus, lateral view; H, distal segment of aedeagus, dorsal view; I, paramere, dorsal view; J, f£ terminalia, lateral view. – *M. parva* sp. nov. K, m$ terminalia, lateral view; L, paramere, inner face, lateral view; M, distal segment of aedeagus, lateral view; N, distal segment of aedeagus, dorsal view; O, paramere, dorsal view; P, f£ terminalia, lateral view. – *M. guatteriae* sp. nov. Q, m$ terminalia, lateral view; R, paramere, inner face, lateral view; S, distal segment of aedeagus, lateral view; T, distal segment of aedeagus, dorsal view; U, paramere, dorsal view; V, f£ terminalia, lateral view. Scale bars: 0.1 mm.

**FIGURE 24.**
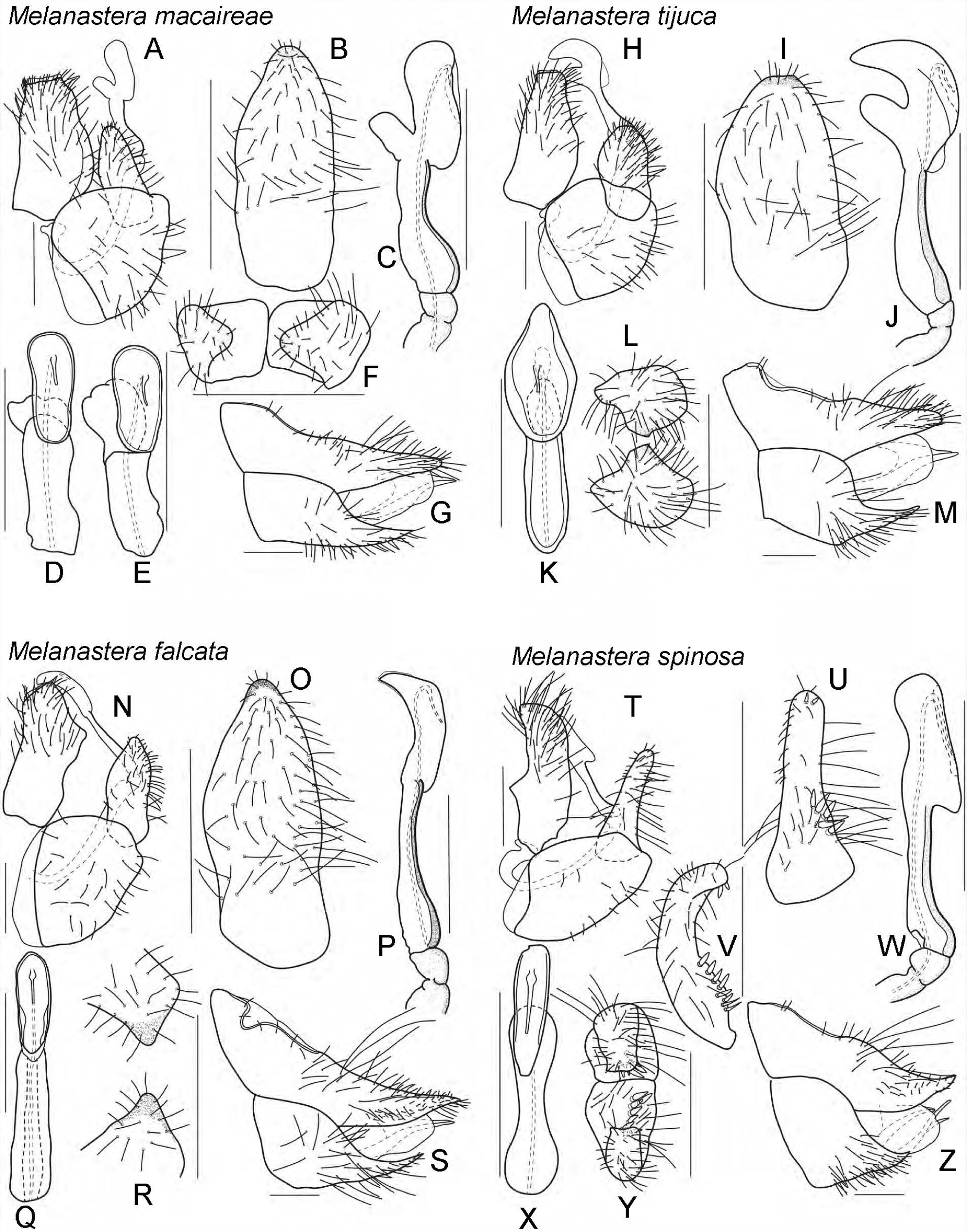
*Melanastera macaireae* sp. nov. A, m$ terminalia, lateral view; B, paramere, inner face, lateral view; C, distal segment of aedeagus, lateral view; D–E, distal segment of aedeagus, dorsal view; F, paramere, dorsal view; G, f£ terminalia, lateral view. – *M. tijuca* sp. nov. H, m$ terminalia, lateral view; I, paramere, inner face, lateral view; J, distal segment of aedeagus, lateral view; K, distal segment of aedeagus, dorsal view; L, paramere, dorsal view; M, f£ terminalia, lateral view. – *M. falcata* sp. nov. N, m$ terminalia, lateral view; O, paramere, inner face, lateral view; P, distal segment of aedeagus, lateral view; Q, distal segment of aedeagus, dorsal view; R, paramere, dorsal view; S, f£ terminalia, lateral view. – *M. spinosa* sp. nov. T, m$ terminalia, lateral view; U, paramere, inner face, lateral view; V, paramere, posterior view; W, distal segment of aedeagus, lateral view; X, distal segment of aedeagus, dorsal view; Y, paramere, dorsal view; Z, f£ terminalia, lateral view. Scale bars: 0.1 mm.

**FIGURE 25.**
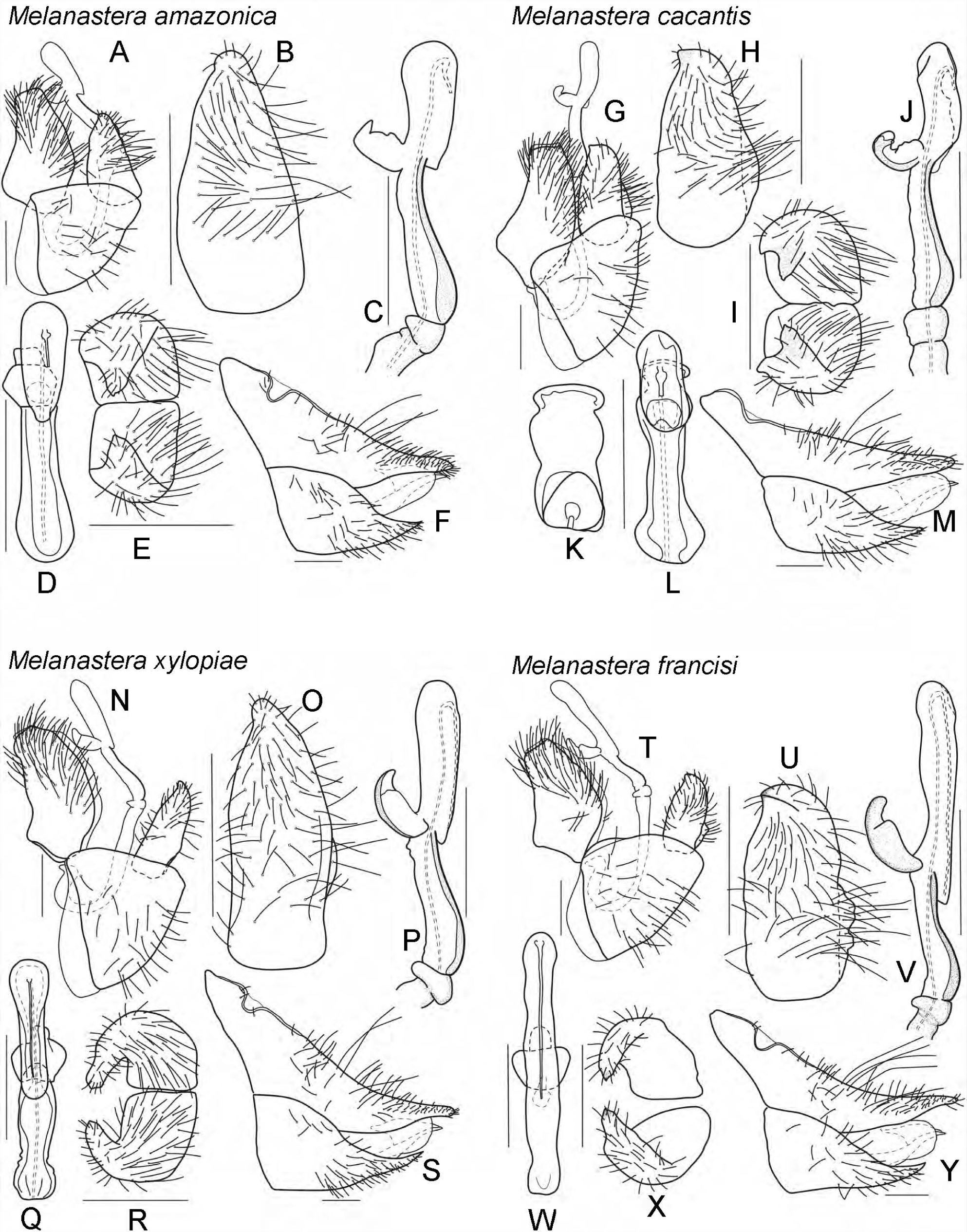
*Melanastera amazonica* sp. nov. A, m$ terminalia, lateral view; B, paramere, inner face, lateral view; C, distal segment of aedeagus, lateral view; D, distal segment of aedeagus, dorsal view; E, paramere, dorsal view; F, f£ terminalia, lateral view. – *M. cacans* sp. nov. G, m$ terminalia, lateral view; H, paramere, inner face, lateral view; I, paramere, dorsal view; J, distal segment of aedeagus, lateral view; K–L, distal segment of aedeagus, dorsal view; M, f£ terminalia, lateral view. – *M. xylopiae* sp. nov. N, m$ terminalia, lateral view; O, paramere, inner face, lateral view; P, distal segment of aedeagus, lateral view; Q, distal segment of aedeagus, dorsal view; R, paramere, dorsal view; S, f£ terminalia, lateral view. – *M. francisi* sp. nov. T, m$ terminalia, lateral view; U, paramere, inner face, lateral view; V, distal segment of aedeagus, lateral view; W, distal segment of aedeagus, dorsal view; X, paramere, dorsal view; Y, f£ terminalia, lateral view. Scale bars: 0.1 mm.

**FIGURE 26.**
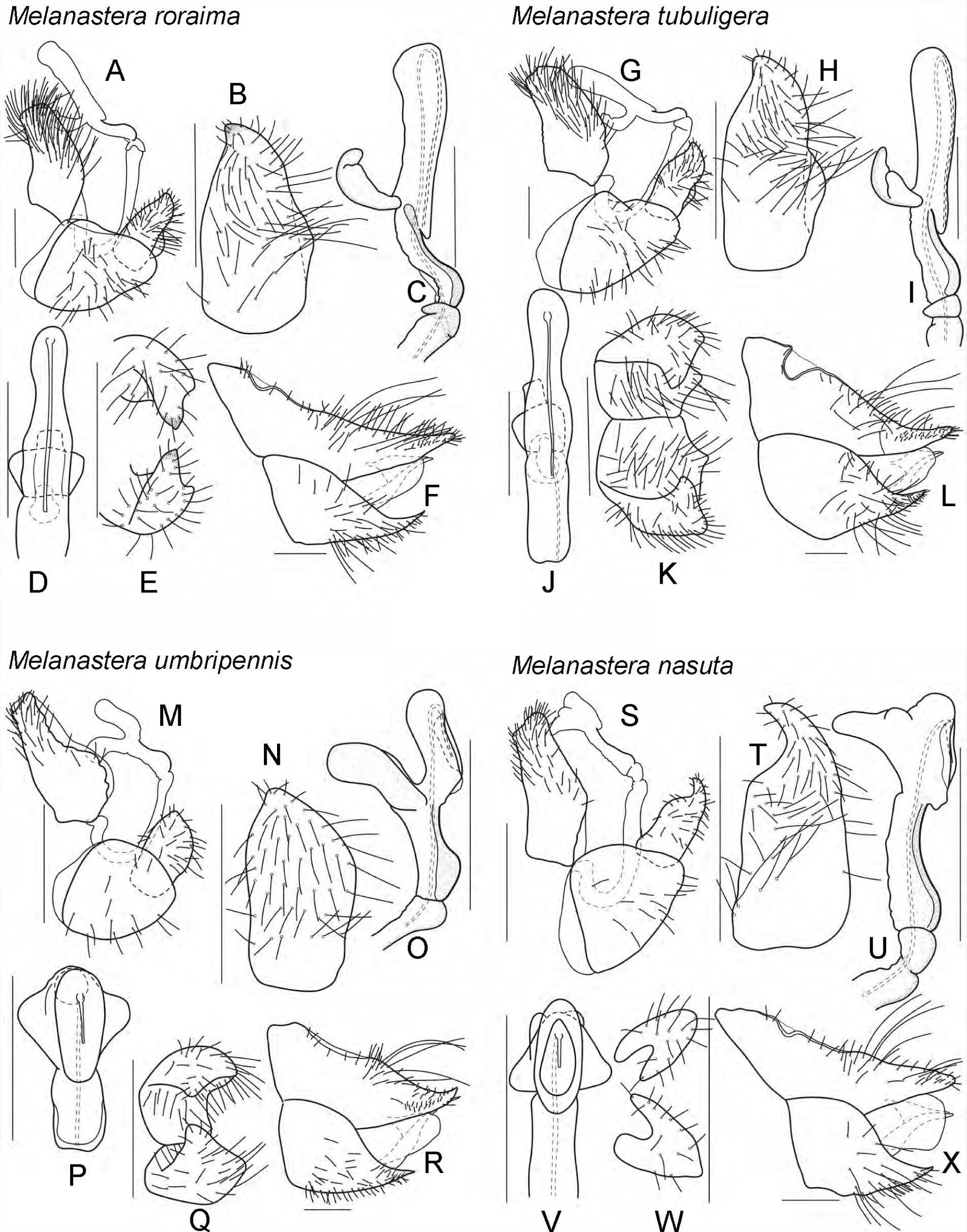
*Melanastera roraima* sp. nov. A, m$ terminalia, lateral view; B, paramere, inner face, lateral view; C, distal segment of aedeagus, lateral view; D, distal segment of aedeagus, dorsal view; E, paramere, dorsal view; F, f£ terminalia, lateral view. – *M. tubuligera* sp. nov. G, m$ terminalia, lateral view; H, paramere, inner face, lateral view; I, distal segment of aedeagus, lateral view; J, distal segment of aedeagus, dorsal view; K, paramere, dorsal view; L, f£ terminalia, lateral view. – *M. umbripennis* sp. nov. M, m$ terminalia, lateral view; N, paramere, inner face, lateral view; O, distal segment of aedeagus, lateral view; P, distal segment of aedeagus, dorsal view; Q, paramere, dorsal view; R, f£ terminalia, lateral view. – *M. nasuta* sp. nov. S, m$ terminalia, lateral view; T, paramere, inner face, lateral view; U, distal segment of aedeagus, lateral view; V, distal segment of aedeagus, dorsal view; W, paramere, dorsal view; X, f£ terminalia, lateral view. Scale bars: 0.1 mm.

**FIGURE 27.**
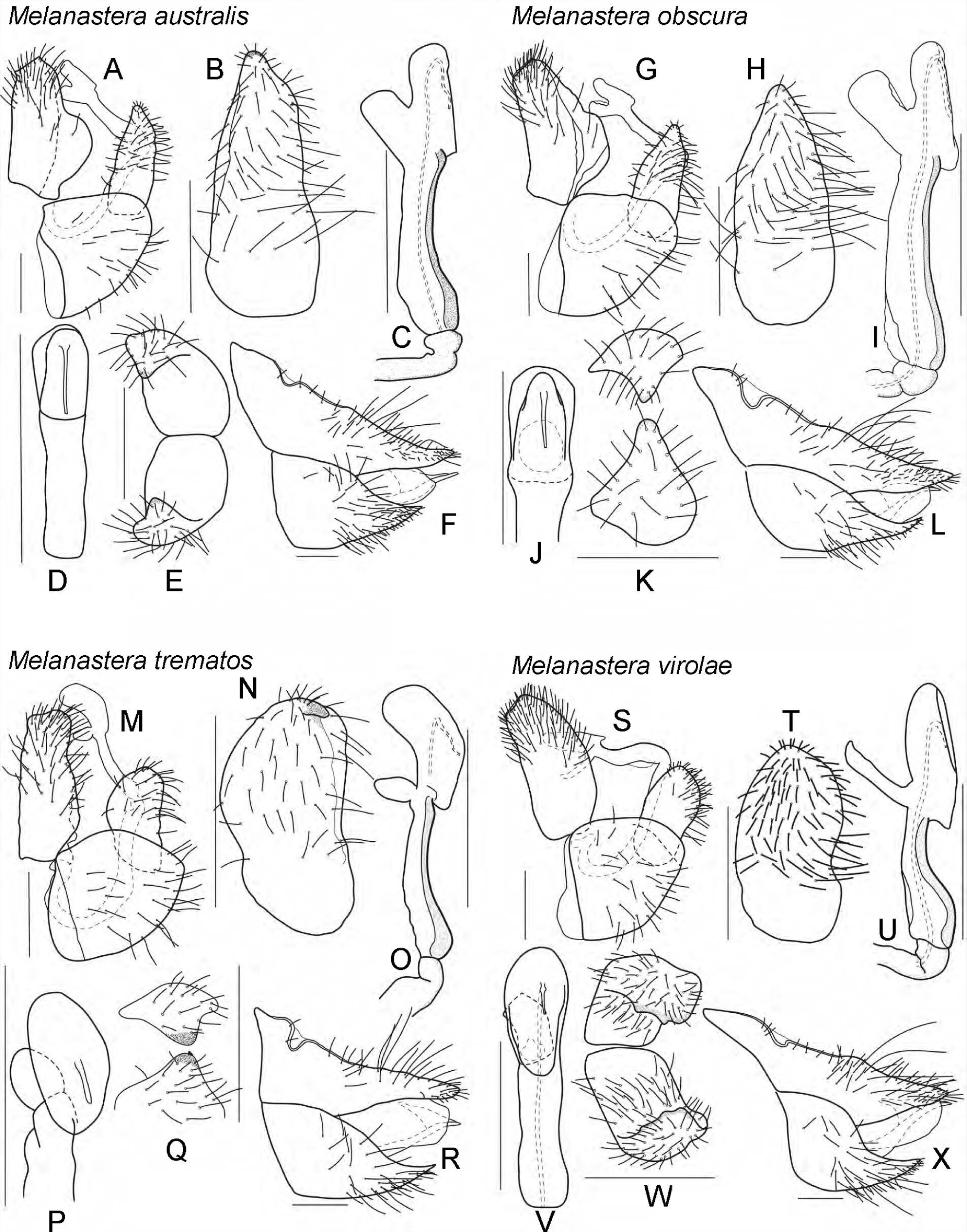
*Melanastera australis* sp. nov. A, m$ terminalia, lateral view; B, paramere, inner face, lateral view; C, distal segment of aedeagus, lateral view; D, distal segment of aedeagus, dorsal view; E, paramere, dorsal view; F, f£ terminalia, lateral view. – *M. obscura* sp. nov. G, m$ terminalia, lateral view; H, paramere, inner face, lateral view; I, distal segment of aedeagus, lateral view; J, distal segment of aedeagus, dorsal view; K, paramere, dorsal view; L, f£ terminalia, lateral view. – *M. trematos* sp. nov. M, m$ terminalia, lateral view; N, paramere, inner face, lateral view; O, distal segment of aedeagus, lateral view; P, distal segment of aedeagus, dorsal view; Q, paramere, dorsal view; R, f£ terminalia, lateral view. – *M. virolae* sp. nov. S, m$ terminalia, lateral view; T, paramere, inner face, lateral view; U, distal segment of aedeagus, lateral view; V, distal segment of aedeagus, dorsal view; W, paramere, dorsal view; X, f£ terminalia, lateral view. Scale bars: 0.1 mm.

**FIGURE 28.**
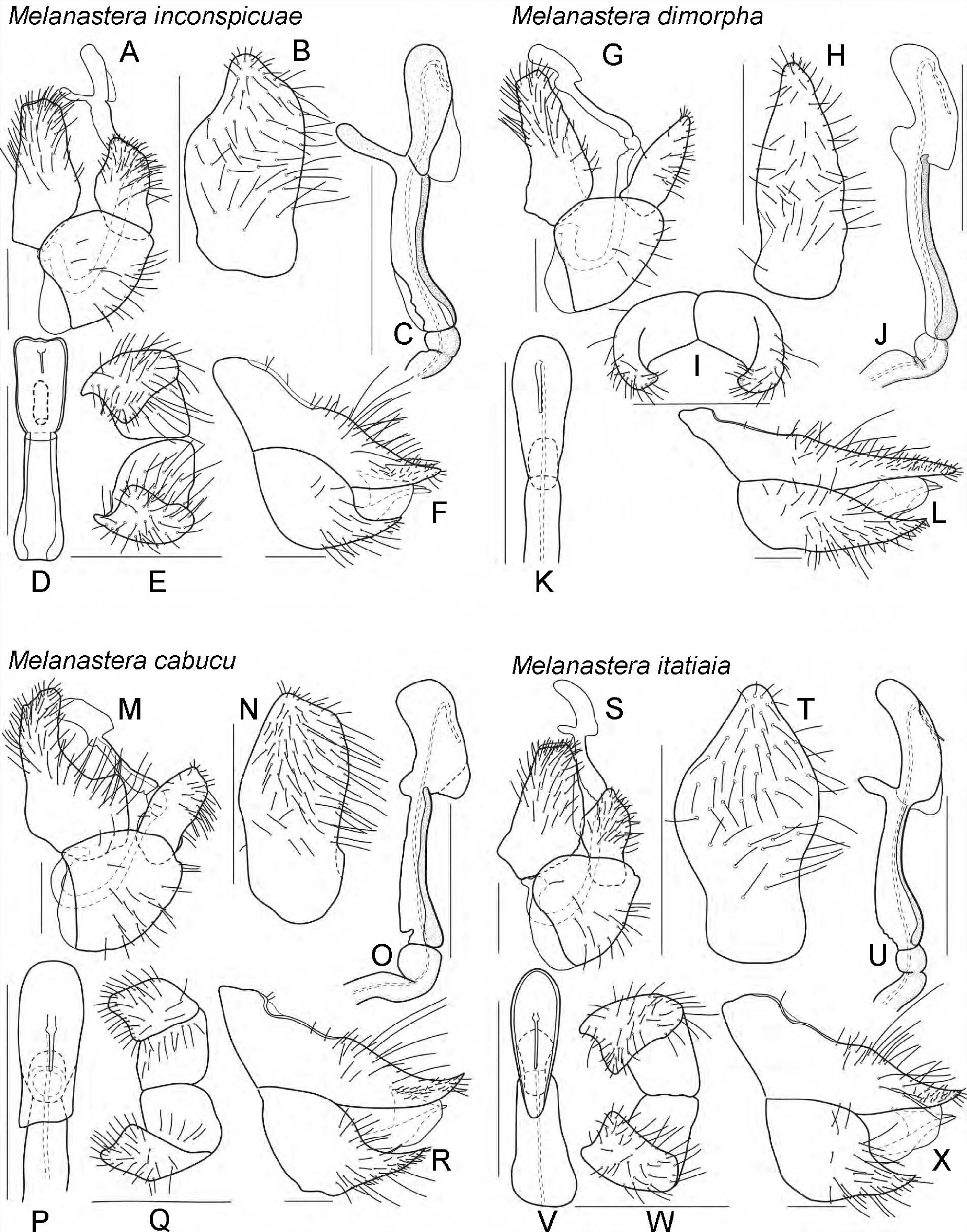
*Melanastera inconspicuae* sp. nov. A, m$ terminalia, lateral view; B, paramere, inner face, lateral view; C, distal segment of aedeagus, lateral view; D, distal segment of aedeagus, dorsal view; E, paramere, dorsal view; F, f£ terminalia, lateral view. – *M. dimorpha* sp. nov. G, m$ terminalia, lateral view; H, paramere, inner face, lateral view; I, paramere, dorsal view; J, distal segment of aedeagus, lateral view; K, distal segment of aedeagus, dorsal view; L, f£ terminalia, lateral view. – *M. cabucu* sp. nov. M, m$ terminalia, lateral view; N, paramere, inner face, lateral view; O, distal segment of aedeagus, lateral view; P, distal segment of aedeagus, dorsal view; Q, paramere, dorsal view; R, f£ terminalia, lateral view. – *M. itatiaia* sp. nov. S, m$ terminalia, lateral view; T, paramere, inner face, lateral view; U, distal segment of aedeagus, lateral view; V, distal segment of aedeagus, dorsal view; W, paramere, dorsal view; X, f£ terminalia, lateral view. Scale bars: 0.1 mm.

**FIGURE 29.**
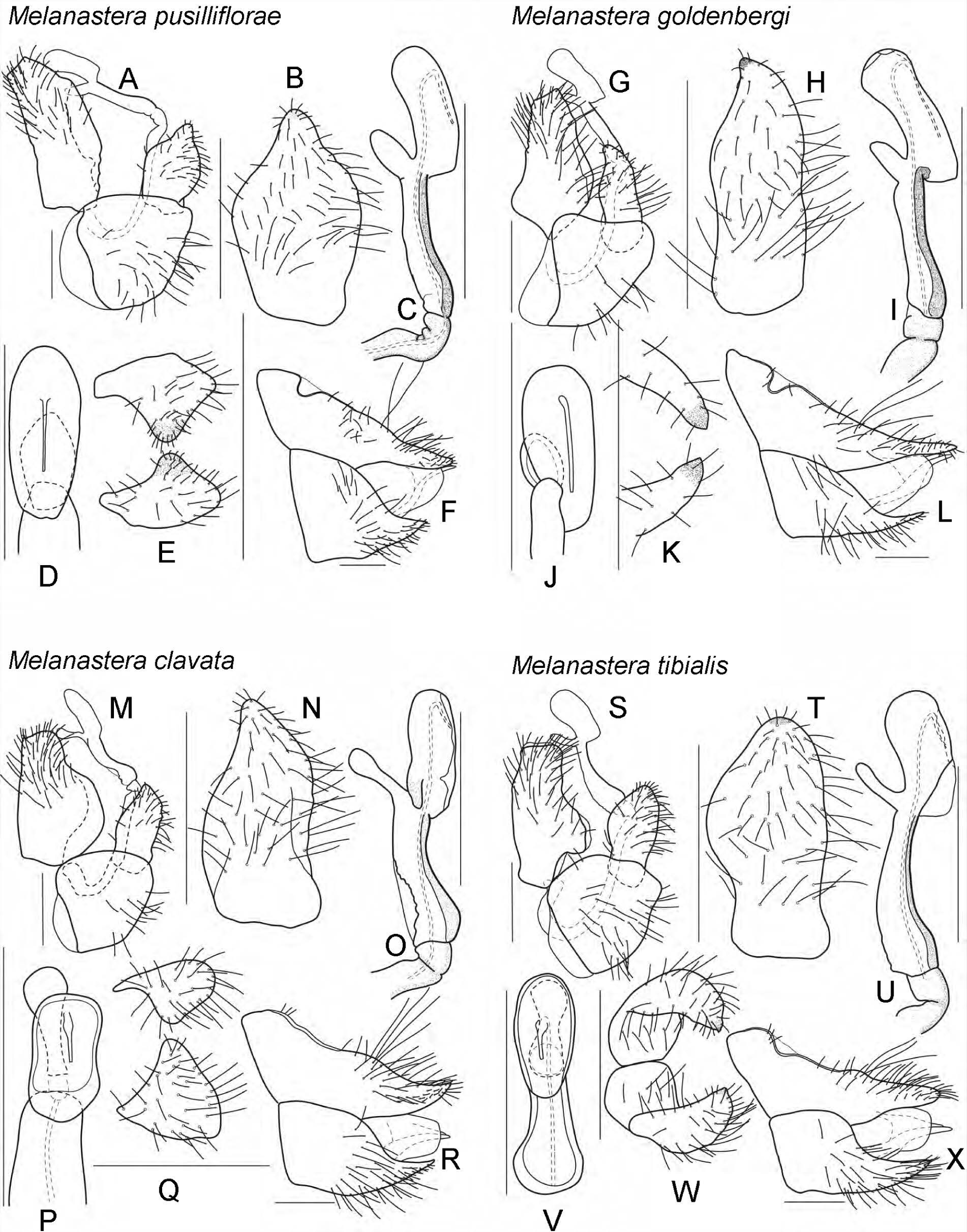
*Melanastera pusilliflorae* sp. nov. A, m$ terminalia, lateral view; B, paramere, inner face, lateral view; C, distal segment of aedeagus, lateral view; D, distal segment of aedeagus, dorsal view; E, paramere, dorsal view; F, f£ terminalia, lateral view. – *M. goldenbergi* sp. nov. G, m$ terminalia, lateral view; H, paramere, inner face, lateral view; I, distal segment of aedeagus, lateral view; J, distal segment of aedeagus, dorsal view; K, paramere, dorsal view; L, f£ terminalia, lateral view. – *M. clavata* sp. nov. M, m$ terminalia, lateral view; N, paramere, inner face, lateral view; O, distal segment of aedeagus, lateral view; P, distal segment of aedeagus, dorsal view; Q, paramere, dorsal view; R, f£ terminalia, lateral view. – *M. tibialis* sp. nov. S, m$ terminalia, lateral view; T, paramere, inner face, lateral view; U, distal segment of aedeagus, lateral view; V, distal segment of aedeagus, dorsal view; W, paramere, dorsal view; X, f£ terminalia, lateral view. Scale bars: 0.1 mm.

**FIGURE 30.**
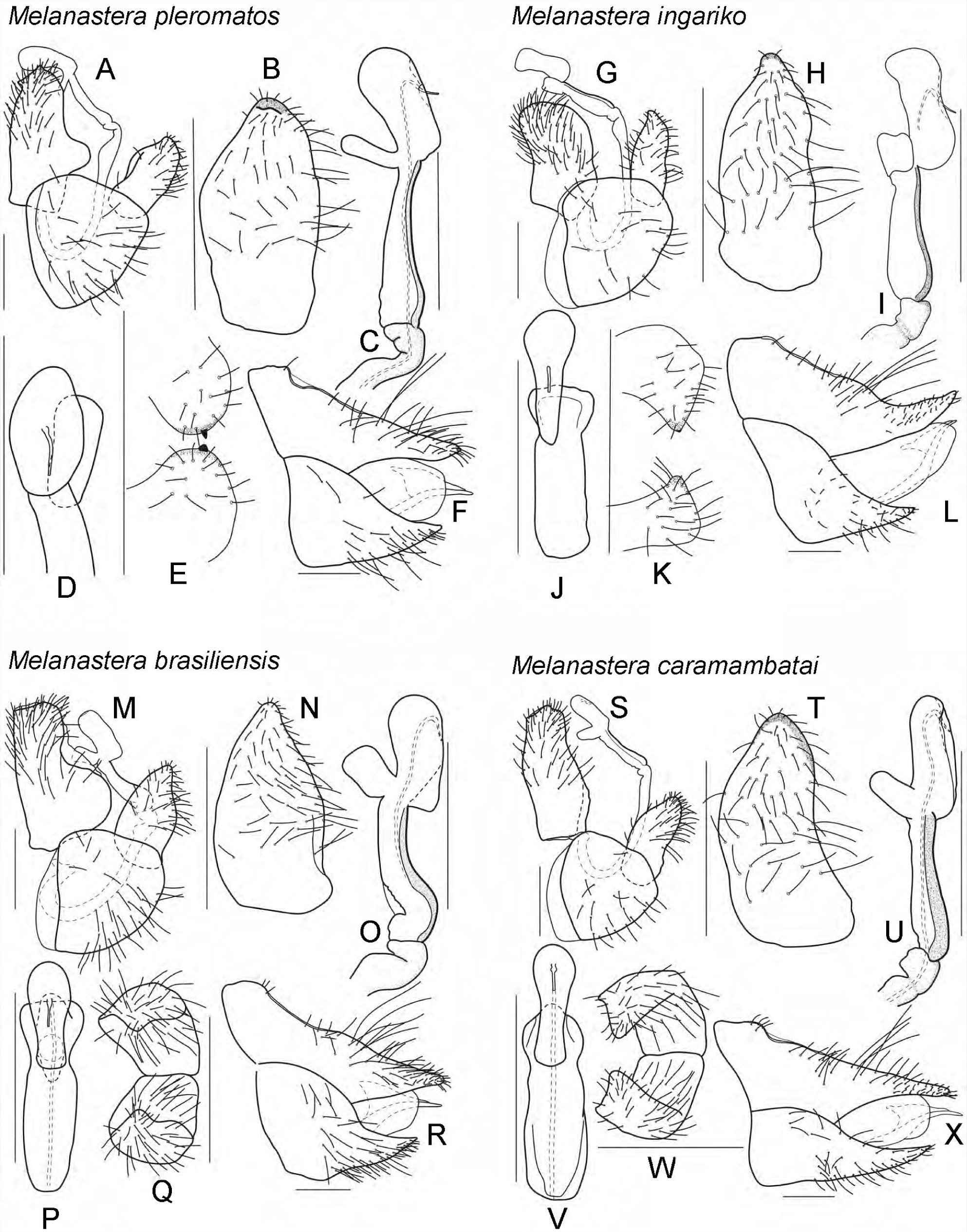
*Melanastera pleromatos* sp. nov. A, m$ terminalia, lateral view; B, paramere, inner face, lateral view; C, distal segment of aedeagus, lateral view; D, distal segment of aedeagus, dorsal view; E, paramere, dorsal view; F, f£ terminalia, lateral view. – *M. ingariko* sp. nov. G, m$ terminalia, lateral view; H, paramere, inner face, lateral view; I, distal segment of aedeagus, lateral view; J, distal segment of aedeagus, dorsal view; K, paramere, dorsal view; L, f£ terminalia, lateral view. – *M. brasiliensis* sp. nov. M, m$ terminalia, lateral view; N, paramere, inner face, lateral view; O, distal segment of aedeagus, lateral view; P, distal segment of aedeagus, dorsal view; Q, paramere, dorsal view; R, f£ terminalia, lateral view. – *M. caramambatai* sp. nov. S, m$ terminalia, lateral view; T, paramere, inner face, lateral view; U, distal segment of aedeagus, lateral view; V, distal segment of aedeagus, dorsal view; W, paramere, dorsal view; X, f£ terminalia, lateral view. Scale bars: 0.1 mm.

**FIGURE 31.**
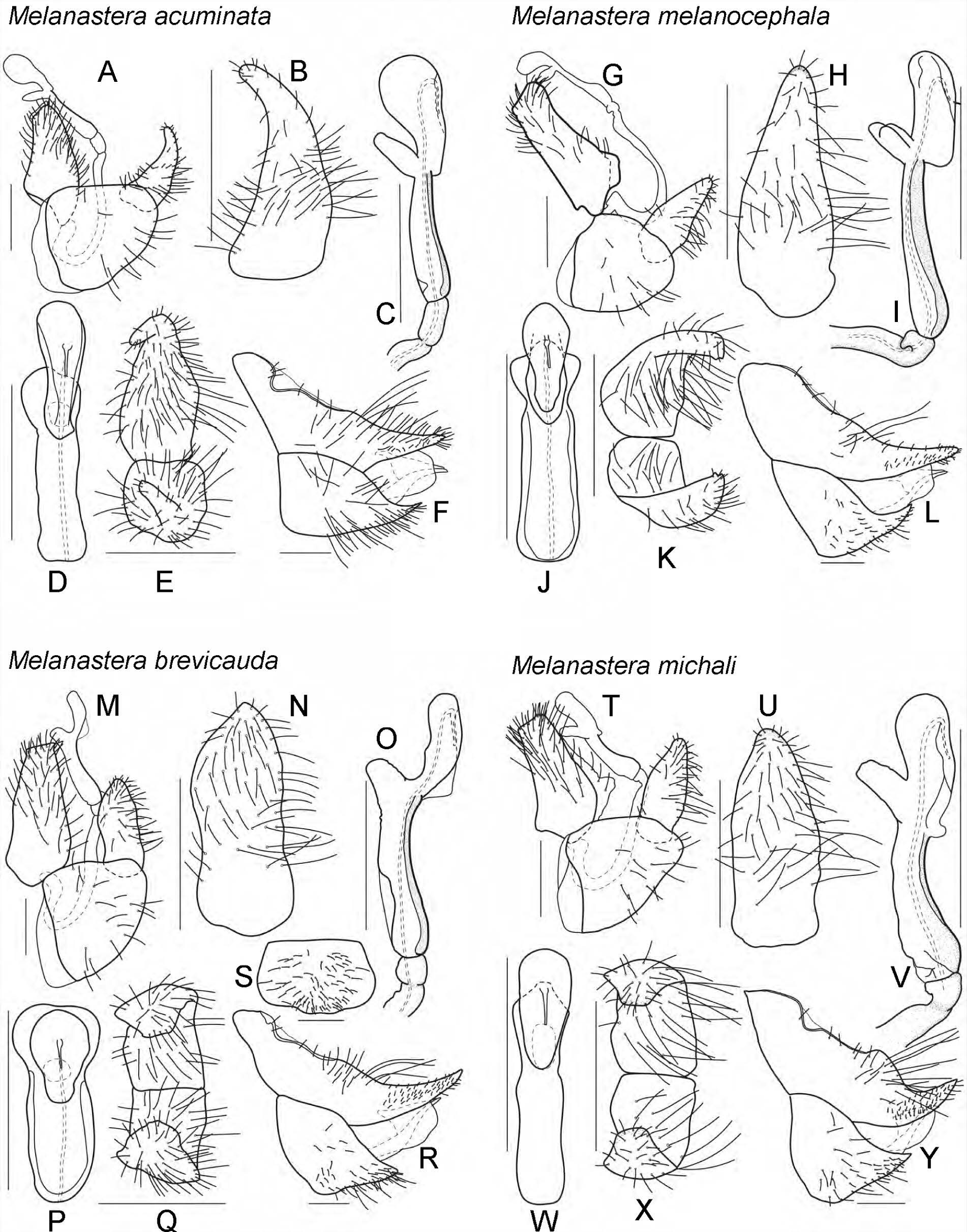
*Melanastera acuminata* sp. nov. A, m$ terminalia, lateral view; B, paramere, inner face, lateral view; C, distal segment of aedeagus, lateral view; D, distal segment of aedeagus, dorsal view; E, paramere, dorsal view; F, f£ terminalia, lateral view. – *M. melanocephala* sp. nov. G, m$ terminalia, lateral view; H, paramere, inner face, lateral view; I, distal segment of aedeagus, lateral view; J, distal segment of aedeagus, dorsal view; K, paramere, dorsal view; L, f£ terminalia, lateral view. – *M. brevicauda* sp. nov. M, m$ terminalia, lateral view; N, paramere, inner face, lateral view; O, distal segment of aedeagus, lateral view; P, distal segment of aedeagus, dorsal view; Q, paramere, dorsal view; R, f£ terminalia, lateral view; S, f£ subgenital plate, ventral view. – *M. michali* sp. nov. T, m$ terminalia, lateral view; U, paramere, inner face, lateral view; V, distal segment of aedeagus, lateral view; W, distal segment of aedeagus, dorsal view; X, paramere, dorsal view; Y, f£ terminalia, lateral view. Scale bars: 0.1 mm.

**FIGURE 32.**
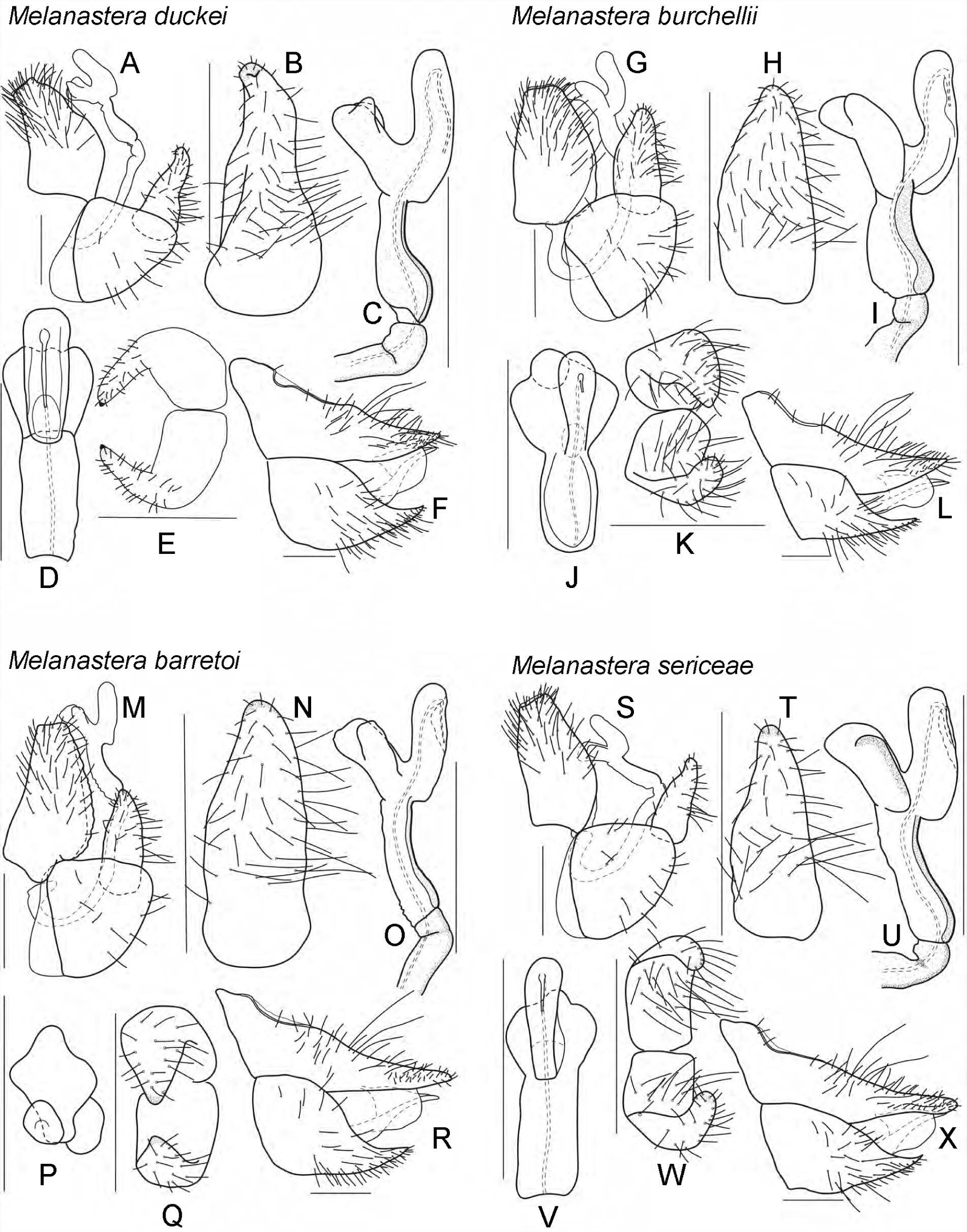
*Melanastera duckei* sp. nov. A, m$ terminalia, lateral view; B, paramere, inner face, lateral view; C, distal segment of aedeagus, lateral view; D, distal segment of aedeagus, dorsal view; E, paramere, dorsal view; F, f£ terminalia, lateral view. – *M. burchellii* sp. nov. G, m$ terminalia, lateral view; H, paramere, inner face, lateral view; I, distal segment of aedeagus, lateral view; J, distal segment of aedeagus, dorsal view; K, paramere, dorsal view; L, f£ terminalia, lateral view. – *M. barretoi* sp. nov. M, m$ terminalia, lateral view; N, paramere, inner face, lateral view; O, distal segment of aedeagus, lateral view; P, distal segment of aedeagus, dorsal view; Q, paramere, dorsal view; R, f£ terminalia, lateral view. – *M. sericeae* sp. nov. S, m$ terminalia, lateral view; T, paramere, inner face, lateral view; U, distal segment of aedeagus, lateral view; V, distal segment of aedeagus, dorsal view; W, paramere, dorsal view; X, f£ terminalia, lateral view. Scale bars: 0.1 mm.

**FIGURE 33.**
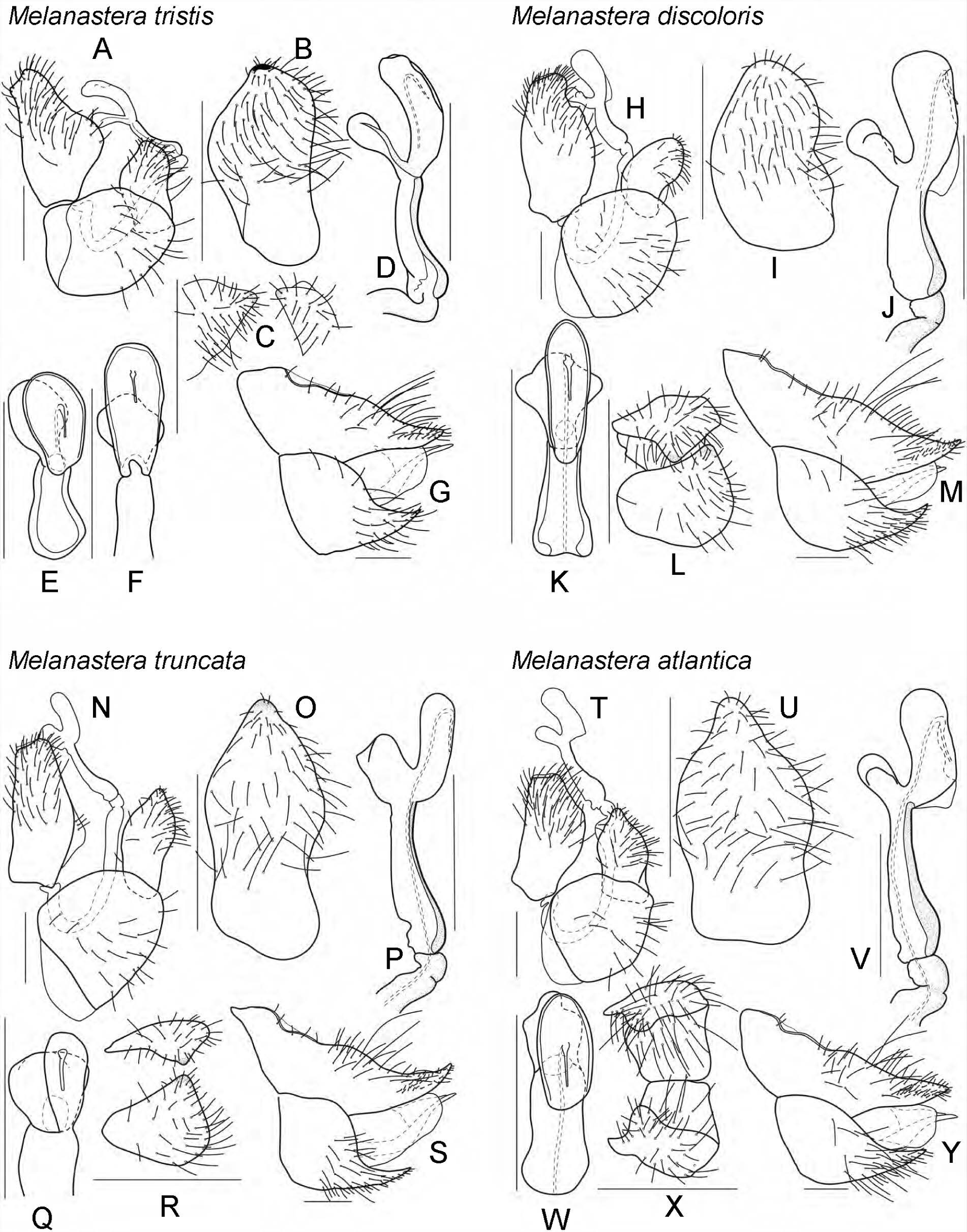
*Melanastera tristis* sp. nov. A, m$ terminalia, lateral view; B, paramere, inner face, lateral view; C, paramere, dorsal view; D, distal segment of aedeagus, lateral view; E–F, distal segment of aedeagus, dorsal view; G, f£ terminalia, lateral view. – *M. discoloris* sp. nov. H, m$ terminalia, lateral view; I, paramere, inner face, lateral view; J, distal segment of aedeagus, lateral view; K, distal segment of aedeagus, dorsal view; L, paramere, dorsal view; M, f£ terminalia, lateral view. – *M. truncata* sp. nov. N, m$ terminalia, lateral view; O, paramere, inner face, lateral view; P, distal segment of aedeagus, lateral view; Q, distal segment of aedeagus, dorsal view; R, paramere, dorsal view; S, f£ terminalia, lateral view. – *M. atlantica* sp. nov. T, m$ terminalia, lateral view; U, paramere, inner face, lateral view; V, distal segment of aedeagus, lateral view; W, distal segment of aedeagus, dorsal view; X, paramere, dorsal view; Y, f£ terminalia, lateral view. Scale bars: 0.1 mm.

**FIGURE 34.**
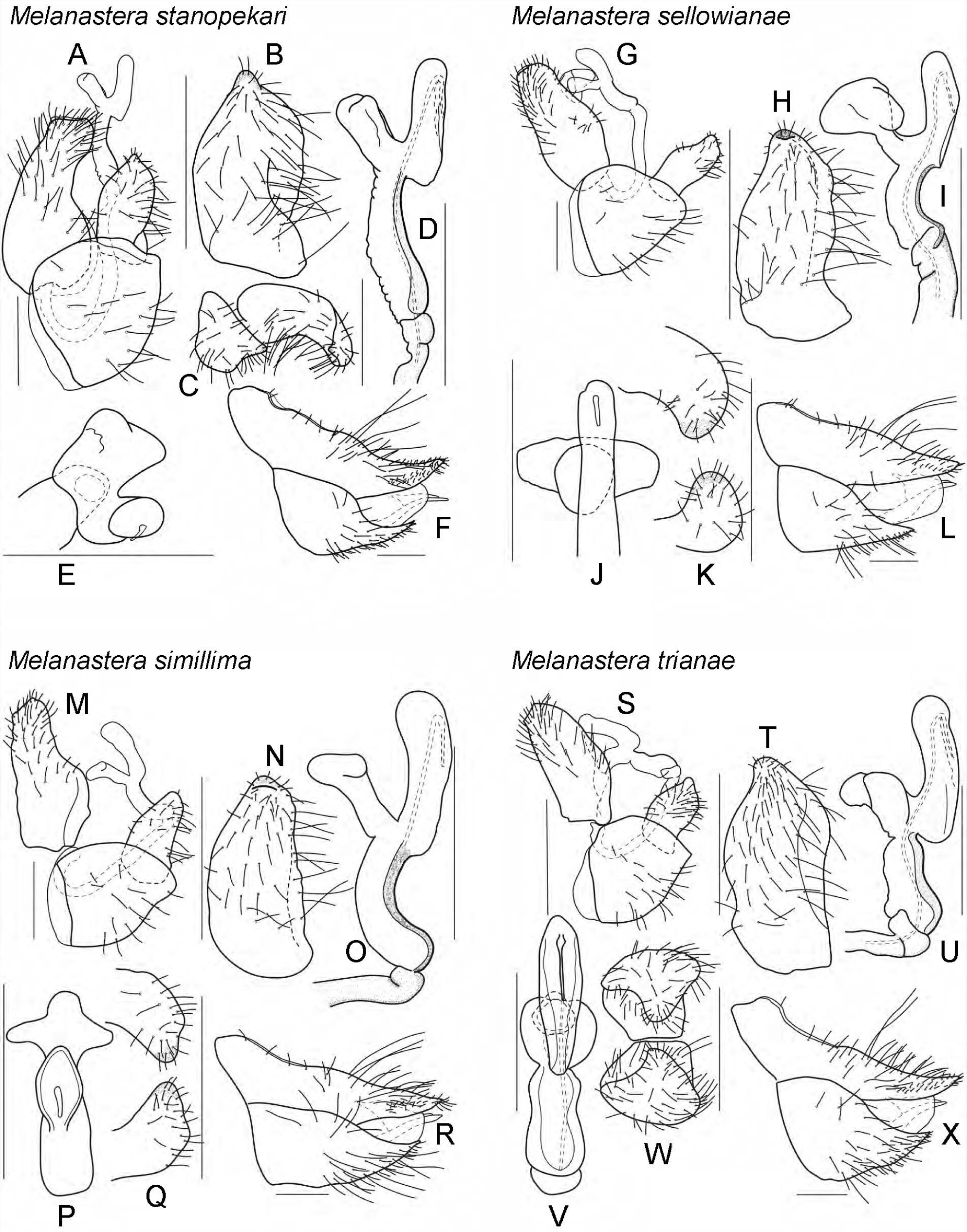
*Melanastera stanopekari* sp. nov. A, m$ terminalia, lateral view; B, paramere, inner face, lateral view; C, paramere, dorsal view; D, distal segment of aedeagus, lateral view; E, distal segment of aedeagus, dorsal view; F, f£ terminalia, lateral view. – *M. sellowianae* sp. nov. G, m$ terminalia, lateral view; H, paramere, inner face, lateral view; I, distal segment of aedeagus, lateral view; J, distal segment of aedeagus, dorsal view; K, paramere, dorsal view; L, f£ terminalia, lateral view. – *M. simillima* sp. nov. M, m$ terminalia, lateral view; N, paramere, inner face, lateral view; O, distal segment of aedeagus, lateral view; P, distal segment of aedeagus, dorsal view; Q, paramere, dorsal view; R, f£ terminalia, lateral view. – *M. trianae* sp. nov. S, m$ terminalia, lateral view; T, paramere, inner face, lateral view; U, distal segment of aedeagus, lateral view; V, distal segment of aedeagus, dorsal view; W, paramere, dorsal view; X, f£ terminalia, lateral view.

**FIGURE 35.**
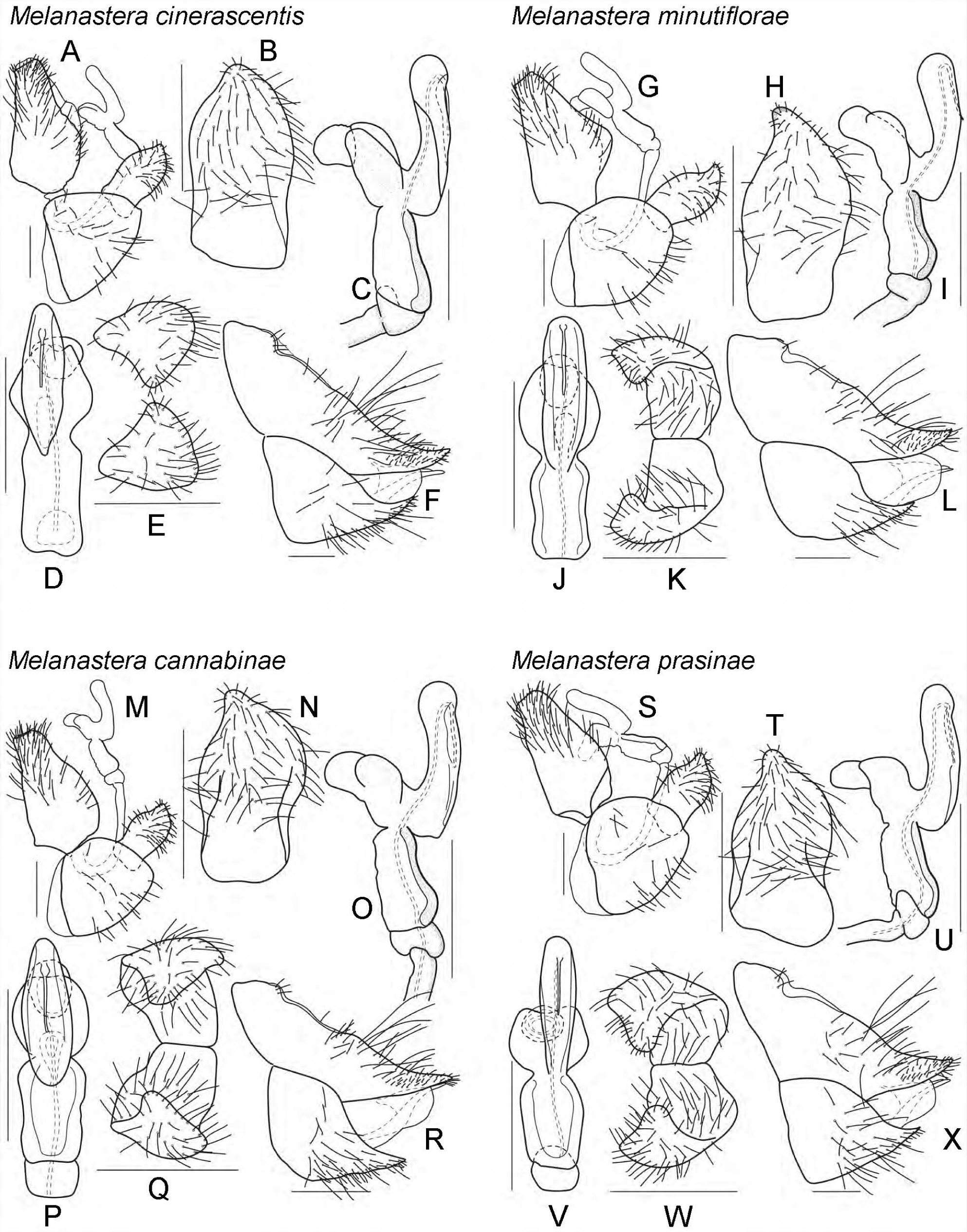
*Melanastera cinerascentis* sp. nov. A, m$ terminalia, lateral view; B, paramere, inner face, lateral view; C, distal segment of aedeagus, lateral view; D, distal segment of aedeagus, dorsal view; E, paramere, dorsal view; F, f£ terminalia, lateral view. – *M. minutiflorae* sp. nov. G, m$ terminalia, lateral view; H, paramere, inner face, lateral view; I, distal segment of aedeagus, lateral view; J, distal segment of aedeagus, dorsal view; K, paramere, dorsal view; L, f£ terminalia, lateral view. – *M. cannabinae* sp. nov. M, m$ terminalia, lateral view; N, paramere, inner face, lateral view; O, distal segment of aedeagus, lateral view; P, distal segment of aedeagus, dorsal view; Q, paramere, dorsal view; R, f£ terminalia, lateral view. – *M. prasinae* sp. nov. S, m$ terminalia, lateral view; T, paramere, inner face, lateral view; U, distal segment of aedeagus, lateral view; V, distal segment of aedeagus, dorsal view; W, paramere, dorsal view; X, f£ terminalia, lateral view. Scale bars: 0.1 mm.

**FIGURE 36.**
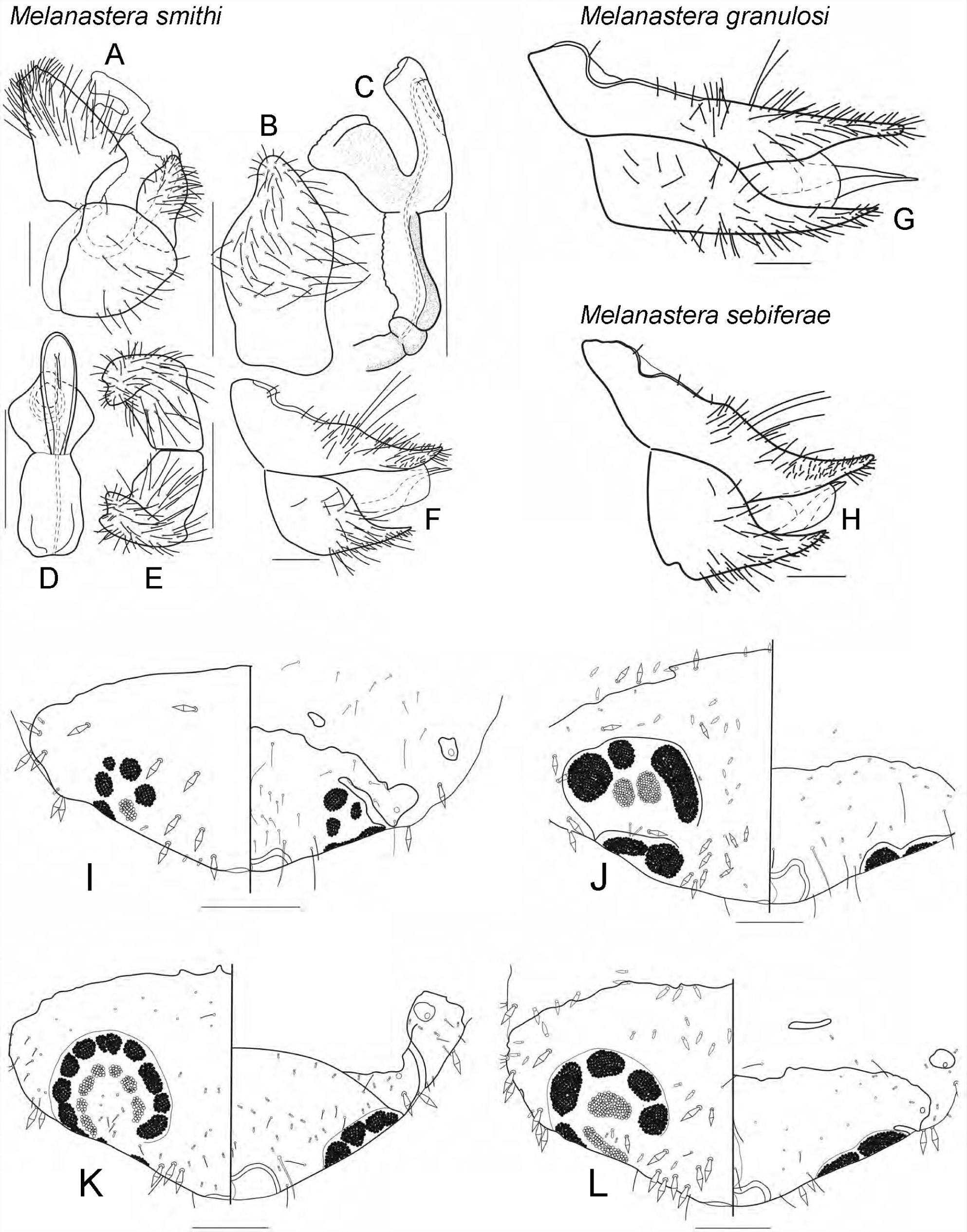
*Melanastera smithi* (Burckhardt, Morais & Picanço). A, m$ terminalia, lateral view; B, paramere, inner face, lateral view; C, distal segment of aedeagus, lateral view; D, distal segment of aedeagus, dorsal view; E, paramere, dorsal view; F, f£ terminalia, lateral view. G, – *M. granulosi* sp. nov., f£ terminalia, lateral view. – H, *M. sebiferae* sp. nov., f£ terminalia, lateral view. – Caudal plate, dorsal (left) and ventral (right) views, of fifth instar immature of *Klyveria* and *Melanastera* spp. I, *K. classiflagellata* (Burckhardt); J, *M. moquiniastri* sp. nov.; K, *M. eremanthi* sp. nov.; L, *M. notia* sp. nov. Scale bars: 0.1 mm.

**FIGURE 37.**
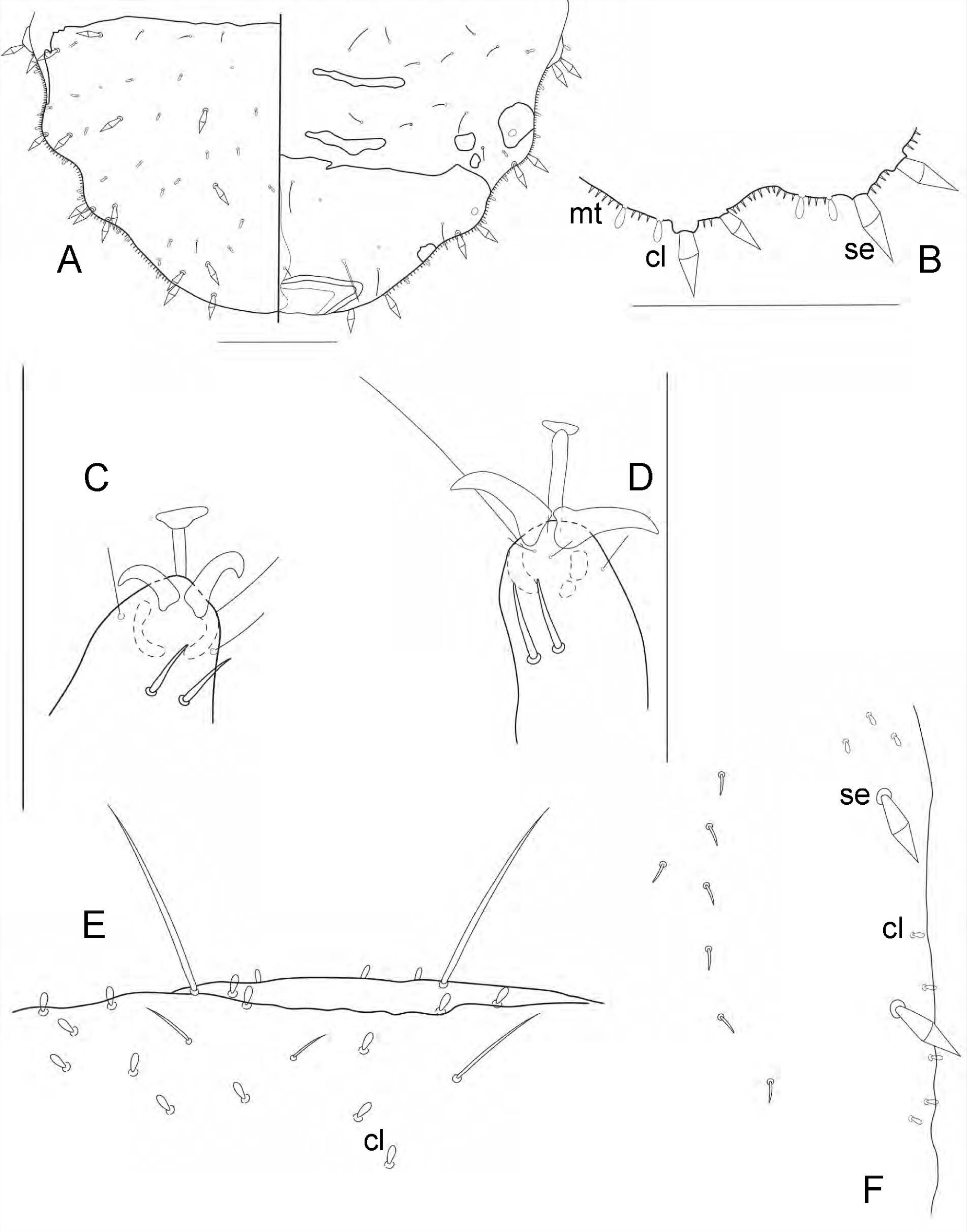
Fifth instar immature. A–C, *Melanastera simillima* sp. nov.; D, *M. eremanthi* sp. nov.; E, F, *M. xylopiae* sp. nov. A, caudal plate, dorsal (left) and ventral (right) views; B, margin of the caudal plate with microtrichia (mt), minute clavate setae (cl) and pointed sectasetae (se); C, D, tarsal arolium with claws, ventral view; E, margin of the head with minute clavate (cl); F, margin of the forewing pad with minute clavate setae (cl) and pointed sectasetae (se). Scale bars: 0.1 mm.

**FIGURE 38.**
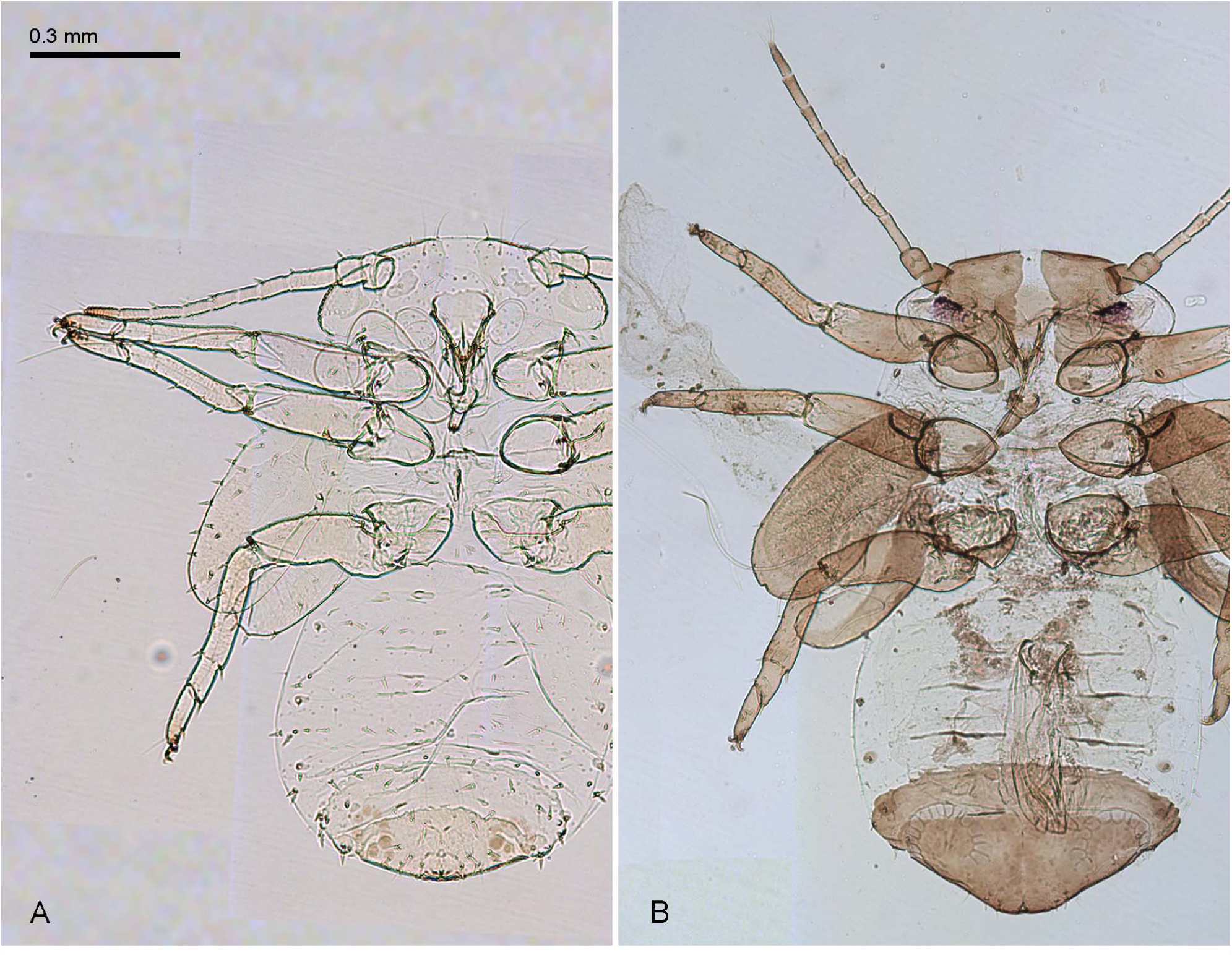
Fifth instar immature. A, *Klyveria crassiflagellata* (Burckhardt); B, *Melanastera smithi* (Burckhardt, Morais & Picanço).

**FIGURE 39.**
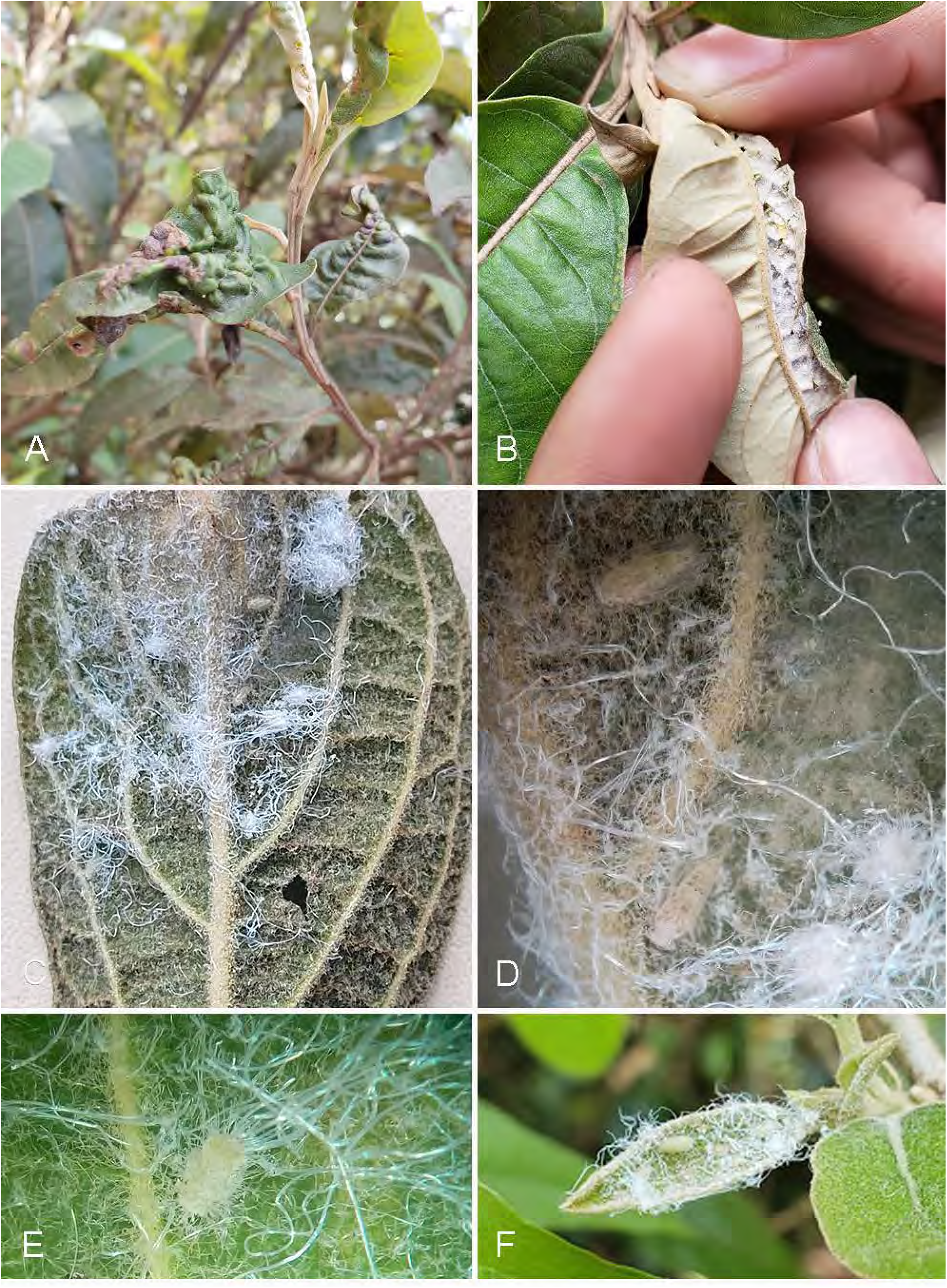
*Melanastera* spp. on Asteraceae. A, leaf deformations induced by *M. eremanthi* sp. nov. on *Eremanthus erythropappus*; B, same but abaxial aspect with open roll gall showing immatures; C–F, adult and immature *M. moquiniastri* sp. nov. on leaves of *Moquiniastrum polymorphum*.

**FIGURE 40.**
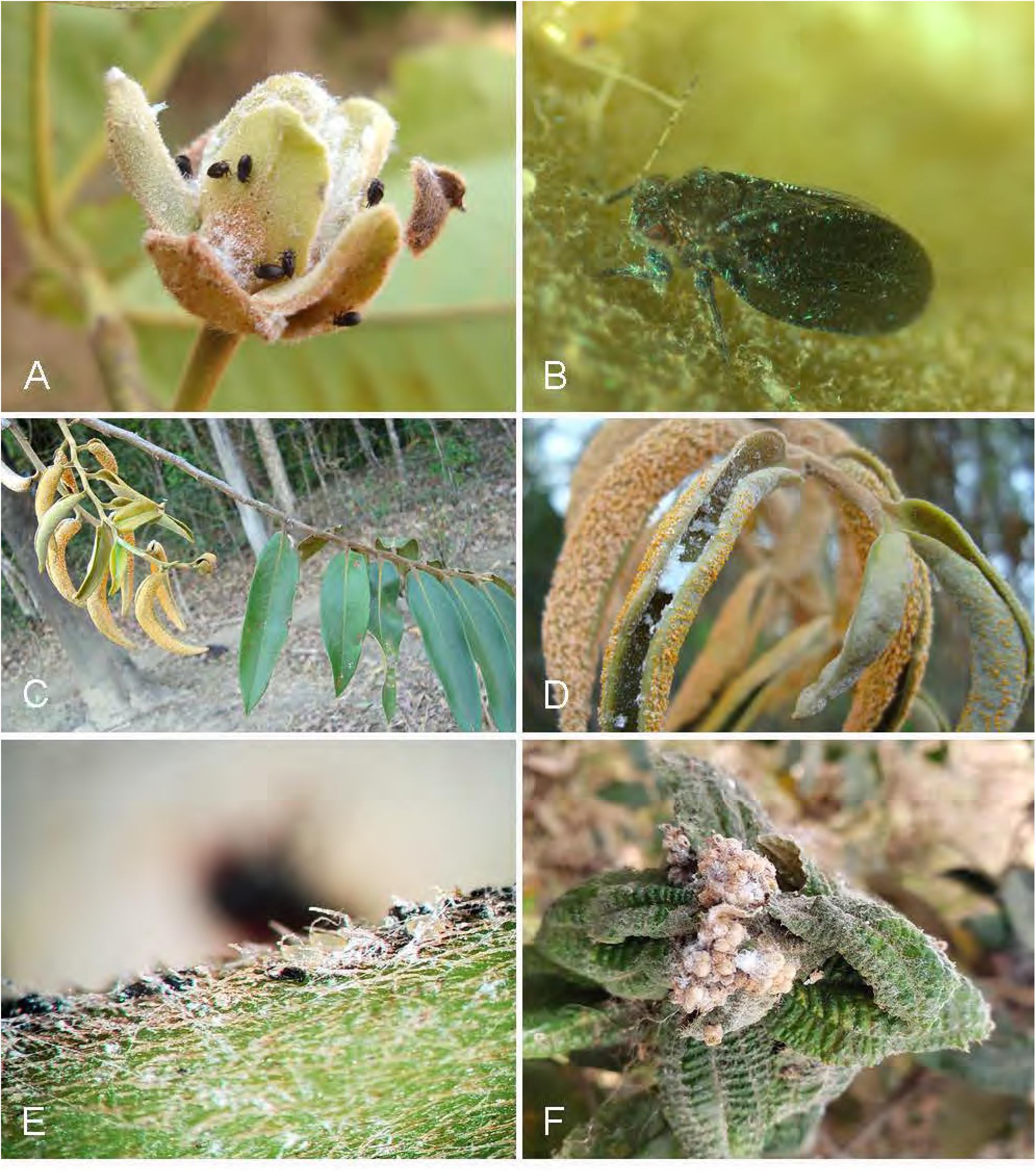
*Melanastera* spp. on Annonaceae and Melastomataceae. A, B, adult *M. olgae* sp. nov. on flowers of *Guatteria punctata*; C, D, immature *M. xylopiae* sp. nov. in galls induced by *Aecidium xylopiae* on *Xylopia aromatica*; E, white and black eggs of *M. melanocephala* sp. nov. on shoot of *Miconia albicans*; F, the feeding of immature and adult *M. melanocephala* sp. nov. on young leaves of *Miconia albicans* damages the leaf epidermis, causing the leaf to crumple.

## APPENDIX 1. GenBank accession numbers for specimens with DNA barcodes.

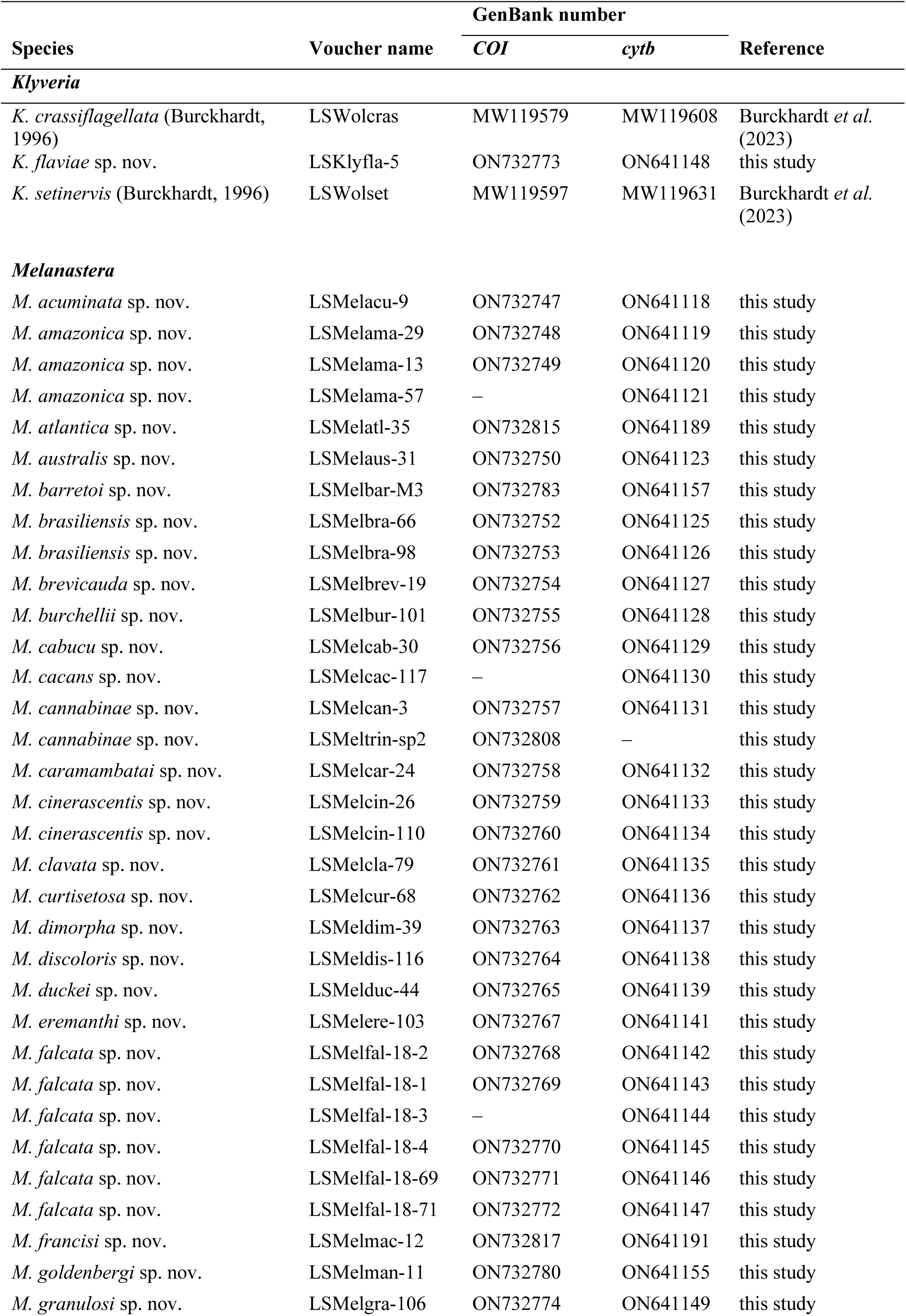
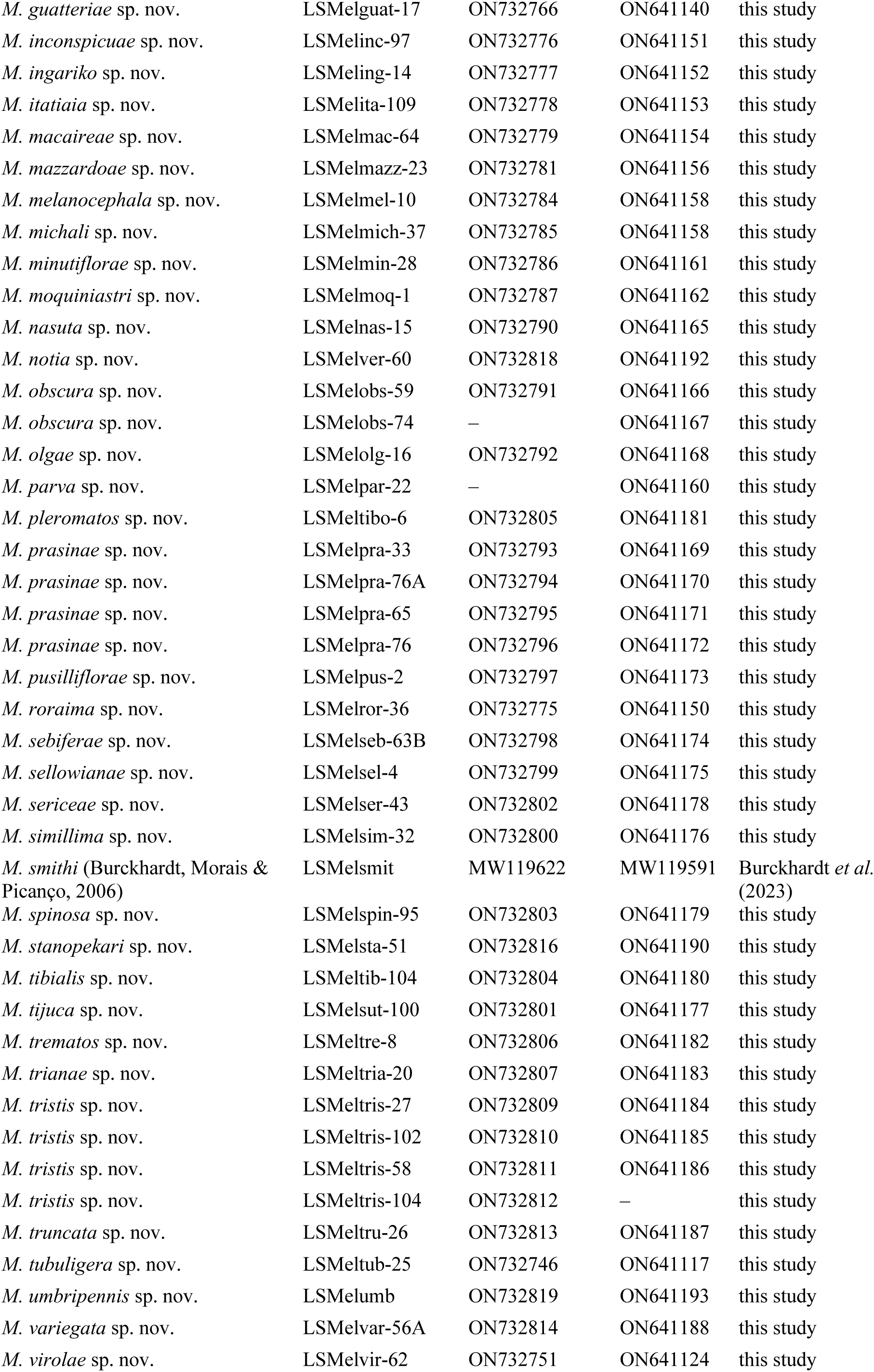
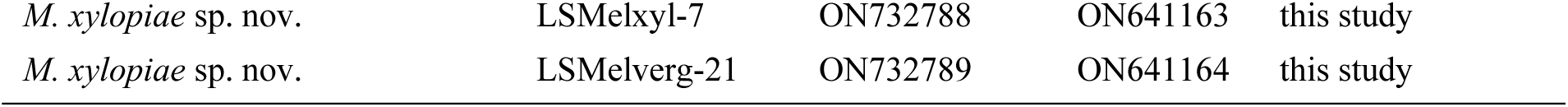

